# Diatom adhesive trail proteins acquired by horizontal gene transfer from bacteria serve as primers for marine biofilm formation

**DOI:** 10.1101/2023.03.06.531300

**Authors:** Jirina Zackova Suchanova, Gust Bilcke, Beata Romanowska, Ali Fatlawi, Martin Pippel, Alastair Skeffington, Michael Schroeder, Wim Vyverman, Klaas Vandepoele, Nils Kröger, Nicole Poulsen

**Author notes:** **Authors for correspondence:** Nicole Poulsen, Email/telephone, Nils Kröger, Email/telephone /.

## Abstract

- Biofilm-forming benthic diatoms are key primary producers in coastal habitats, where they frequently dominate sunlit submerged and intertidal substrata. The development of a unique form of gliding motility in raphid diatoms was a key molecular adaptation that contributed to their evolutionary success. Gliding motility is hypothesized to be driven by an intracellular actin-myosin motor and requires the secretion of polysaccharide- and protein-based adhesive materials. To date, the structure-function correlation between diatom adhesives utilized for gliding and their relationship to the extracellular matrix that constitutes the diatom biofilm is unknown.
- Proteomics analysis of the adhesive material from *Craspedostauros australis* revealed eight novel, diatom-specific proteins. Four of them constitute a new family of proteins, named Trailins, which contain an enigmatic domain termed Choice-of-Anchor-A (CAA). Immunostaining demonstrated that Trailins are only present in the adhesive trails required to generate traction on native substrata, but are absent from the extracellular matrix of biofilms. Phylogenetic analysis and Protein 3D structure prediction suggests that the CAA-domains in Trailins were obtained from bacteria by horizontal gene transfer, and exhibit a striking structural similarity to ice-binding proteins.
- Our work advances the understanding of the molecular basis for diatom underwater adhesion and biofilm formation providing evidence that there is a molecular switch between proteins required for initial surface colonization and those required for 3D biofilm matrix formation.

## Introduction

Many marine microorganisms are capable of adhering to surfaces underwater where they form taxonomically diverse, often photosynthetically active biofilm communities embedded in a three-dimensional (3D) matrix of self-produced extracellular polymeric substances (EPS). In fact biofilms are the most abundant microbial lifestyle on the planet, accounting for ~80% of all bacterial cells (Flemming & Wuertz, 2019). In shallow coastal habitats, diatoms and cyanobacteria rich biofilms contribute immensely to global primary productivity, being responsible for ~20% of marine carbon fixation even though they only occupy ~0.03% of the ocean’s surface area (Pinckney, 2018). The carbohydrate-rich EPS provides numerous advantages for the encapsulated cells, including protection against desiccation and mechanical stresses, thereby enhancing their ability to thrive in highly dynamic environments (Flemming & Wingender, 2010; Steele *et al*., 2014).

The evolution of a raphe system in pennate diatoms was a significant milestone that enabled the development of a unique mode of active gliding motility and the production of copious amounts of EPS, which facilitated the colonization of a wide range of benthic habitats (Ashworth, 2013; Nakov *et al*., 2018). Furthermore, the gliding of diatoms allows cells to actively optimize their position in the environment allowing them to move towards favorable light and nutrient conditions, while avoiding desiccation and toxic compounds. Diatom motility is critically dependent on the presence of a specialized slit in the silica cell wall, termed the raphe. It is hypothesized that the secretion of EPS strands through the raphe slit provides a continuous physical link between the plasma membrane and the substratum (Edgar & Pickett-Heaps, 1984). The EPS strands have remarkable properties: they (i) form intertwined long tethers that can extend a distance of up to 30 μm away from the cell body, (ii) exhibit high adhesive, elastic and tensile strength, and (iii) are adhesive underwater on a wide range of materials (Higgins *et al*., 2003; Holland *et al*., 2004; Dugdale *et al*., 2006; Gutierrez-Medina *et al*., 2022). One of the proposed models for gliding is based on the observation that two actin bundles are positioned immediately below the plasma membrane along the entire length of the raphe (Edgar & Zavortink, 1983; Edgar & Pickett-Heaps, 1984). It is hypothesized that the EPS strands are connected to the acto-myosin motor system via a continuum of biomolecules that span the plasma membrane, which is referred to as the adhesion motility complex (AMC). According to this model, the acto-myosin motor system translocate the EPS strands that are anchored to the substratum in a rearward direction, thus propelling the cell forwards (Edgar, 1983; Edgar & Zavortink, 1983; Edgar & Pickett-Heaps, 1984). This model was supported by inhibition studies with drugs specific for actin and myosin (Poulsen *et al*., 1999), but the EPS composition and the other components of the AMC remained unknown.

To investigate the proposed mechanism for diatom gliding we have embarked on identifying the components of the machinery. As diatoms glide across a surface, they often deposit behind them a trail of the EPS strands that are composed of acidic polysaccharides and proteins (Lind *et al*., 1997; Higgins *et al*., 2000; Poulsen *et al*., 2014). We have previously performed a proteomics study and identified 21 putative adhesion proteins from *Amphora coffeaeformis* (Lachnit *et al*., 2019). These proteins contain several features that also occur in adhesive proteins from other organisms, and a newly described, diatom-specific GDPH-domain. Immunolocalization of one GDPH-domain containing protein, Ac629, confirmed its presence in the raphe and EPS trails, which is consistent with a role in diatom adhesion (Lachnit *et al*., 2019). However, bioinformatics analysis revealed that not all motile diatoms possess proteins with a GDPH-domain (Lachnit *et al*., 2019), indicating that other protein domains must be able to mediate diatom adhesion. To address this question, we have pursued a proteomics analysis of the adhesive trails from *Craspedostauros australis*, which is a raphid diatom species lacking GDPH-domain bearing proteins. Through long-read genome sequencing and assembly, comparative genomics, phylogenetics, structural homology and immunolocalization studies we aimed to investigate the evolutionary history of diatom adhesive proteins and their role in the formation of diatom trails and biofilms.

## Materials and methods

### Cell cultures

*Craspedostauros australis* (Cox, CCMP 3328) was grown in artificial seawater medium (ESAW) (Harrison *et al*., 1980) at 18°C and 12 hours light/12 hours dark cycle with an intensity of 100 μmol photons m^-2^ s^-1^, using cool-white lamps.

### Isolation of diatom adhesive material

The *C. australis* adhesive material (AM) was purified according to a previously published procedure (Poulsen *et al*., 2014) and stored at −20°C. The freeze-dried AM was solubilized in 0.5 M borate buffer (pH 8.5) containing 2 M hydroxylamine (Sigma Aldrich) for 2 hours at 45°C and then desalted against 50 mM ammonium acetate using a PD MidiTrap G-10 column (size exclusion limit: 700 Da; GE Healthcare, Sweden) according to the manufacturer’s instructions. Detailed methods for the anhydrous hydrofluoric acid treatment and gel filtration chromatography are in the supplementary information.

### LC-MS/MS proteomics analysis

The solubilized AM was subjected to SDS-PAGE and after a short separation (~ 4 cm migration into the separating gel) visualized by Coomassie Blue staining. An entire gel lane was cut into four slabs, and each slab was subjected to in-gel proteolytic digestion with trypsin (Promega, Mannheim, Germany) overnight as described in (Shevchenko *et al*., 2006). Detailed LC-MS methods are provided in the supplementary information.

### PacBio genomic DNA sequencing

HMW genomic DNA (gDNA) was isolated using a CTAB and phenol-chloroform method. The Dresden-concept genome center (https://dresden-concept.de/genome-center/?lang=en) prepared the libraries for long read PacBio sequencing. Detailed methods are described in the supplementary information.

### Confirmation of gene models

RACE and RT-PCR were used to determine the full-length gene models of the *C. australis* adhesive trail proteins, detailed methods and primer sequences are provided in the supplementary information (Table S1).

### Functional annotation and homology searches

Proteins identified by mass spectrometry were manually annotated according to transcriptomic and genomic data (Poulsen, 2023) and the PacBio genome assembly generated in this study. Protein sequence homology analysis was performed using a BLASTx search against the non-redundant protein sequences in NCBI database with an E-value cutoff of 10^-10^. Parameters used for further bioinformatics searches are found in the supplementary information. The evolutionary origin of diatom CAA-domains was assessed by creating Hidden Markov Model (HMM) profiles which were used to search for homologous protein domains in the PLAZA Diatoms 1.0, MMETSP and the NCBI nr datasets (Keeling *et al*., 2014; Osuna-Cruz *et al*., 2020), as described in supplementary information.

### Structural analysis of diatom (Choice-of-Anchor-A) CAA-domains

Alphafold (version v2.0.1) was used to predict the structure of the CAA-domain and compared to experimentally-determined 3D protein structures in the PDB database (https://www.rcsb.org/). Detailed methods are provided in the supplementary information.

### Genetic transformation of C. australis

*C. australis* was genetically transformed using a previously developed protocol (Poulsen, 2023). In brief, 1×10^8^ cells were plated on an ESAW agar plate and five μg of plasmid DNA coated on W-microparticles (M17, Biorad) was delivered into the cells using the Biorad Biolistic Particle Delivery System (1550 psi, 28 mmHg vacuum). The cells were allowed to recover for 24 hours, then 5×10^6^ cells were plated on ESAW agar plates containing 450 μg mL^-1^ nourseothricin (Jena Bioscience), and incubated in constant light at 18 °C.

### Production of polyclonal antibodies

Rabbit polyclonal antibodies were obtained from GenScript (NJ, USA). Epitopes were chosen based on the manufacturer’s prediction algorithm results in regions that were covered by the protein sequencing: CaTrailin3 DEDLSKQNTGKTIN; CaTrailin2 EDNLDQIRIITESN; CaTrailin4 DDNVPYEETQRHTA. The antibodies were raised against selected protein sequences and purified by affinity chromatography using the antigen as a ligand. To obtain the IgG fraction, the antibodies were further affinity purified using Protein G mag sepharose (GE Healthcare).

### Immunofluorescence and Lectin labelling

For immunolabelling of adhesive trails, 5×10^3^ *C. australis* cells were incubated for 16 hours on a glass bottom chamber slide (μ-slide 8 well chamber, ibidi). All steps were performed at RT. The culture medium was aspirated and the cells incubated in 2% BSA in PBS (BS) for 1 hour. The samples were then incubated for 2 hours with primary antibodies or pre-immune serum (5 μg·mL^-1^) diluted in BS + 0.05% Tween 20. The samples were washed (1x 2 sec, 1x 5 min) with PBS + 0.05% Tween 20 (PBST), and then incubated in secondary antibody (Goat-anti-rabbit conjugated with AlexaFluor488, 1:3000 diluted in PBST; Life Technologies) for 1 hour in the dark. After washing twice with PBST and overlaying with ibidi mounting medium (Ibidi), confocal microscopy was performed using a Zeiss LSM780 inverted microscope equipped with a Zeiss Plan Apochromat 63x (1.4) Oil DIC M27 objective or C-Apochromat 40x/1.2 W Corr M27 objective. Two channels were used to separately monitor chloroplast fluorescence (emission at 654-693 nm) and AlexaFluor488 or Atto488 channel (emission at 480-515 nm). Images were analysed using the ZEN 3.1 software (Zeiss). Detailed methods for the lectin labelling and whole cell and biofilm immunolabelling are provided in the supplementary information.

## Results

### Identification of C. australis adhesive trail proteins

The biochemical characterization of biological adhesives is hampered by their insolubility in reagents typically used in biochemical studies. Recently, we demonstrated that hydroxylamine completely solubilizes the *A. coffeaeformis* AM, which enabled identification of the first diatom adhesive trail proteins (Lachnit *et al*., 2019). Following the same approach, we used hydroxylamine to solubilize the *C. australis* AM. SDS-PAGE in combination with Stains-All staining revealed that the *C. australis* AM, like *A. coffeaeformis*, is dominated by acidic, high molecular weight (HMW) components (>250 kDa) and a minor fraction of low molecular weight (LMW) components (<30 kDa) (Fig. S1A). In contrast to the *A. coffeaformis* LMW components, which can be stained with Coomassie blue, the *C. australis* LMW components can only be stained with silver, indicating clear differences in the chemical composition of the LMW components from both diatom species (Fig. S1A).

To identify proteins in the AM, we performed a proteomics analysis of two different samples, hydroxylamine solubilized adhesive material (i) before and (ii) after hydrofluoric acid (HF) treatment, which was used to remove glycan moieties that may impede protein sequencing. The proteomics analysis resulted in the identification of eight proteins (Fig. 1, Table S2 and S3). Among these was CaFAP1, which was recently shown to be a cell surface glycoprotein that is sloughed off the cell wall during gliding and remains associated with the adhesive trails (Poulsen, 2023). The presence of CaFAP1 in the AM was therefore expected and served as a positive control for the identification of adhesive trail proteins. Four of the eight proteins contain ‘choice-of-A-anchor’ (CAA) domains, and were named Trailins, because immunolabelling confirmed their presence in the adhesive trails (see below “Immunolocalization of Trailins”).

**Figure 1.**
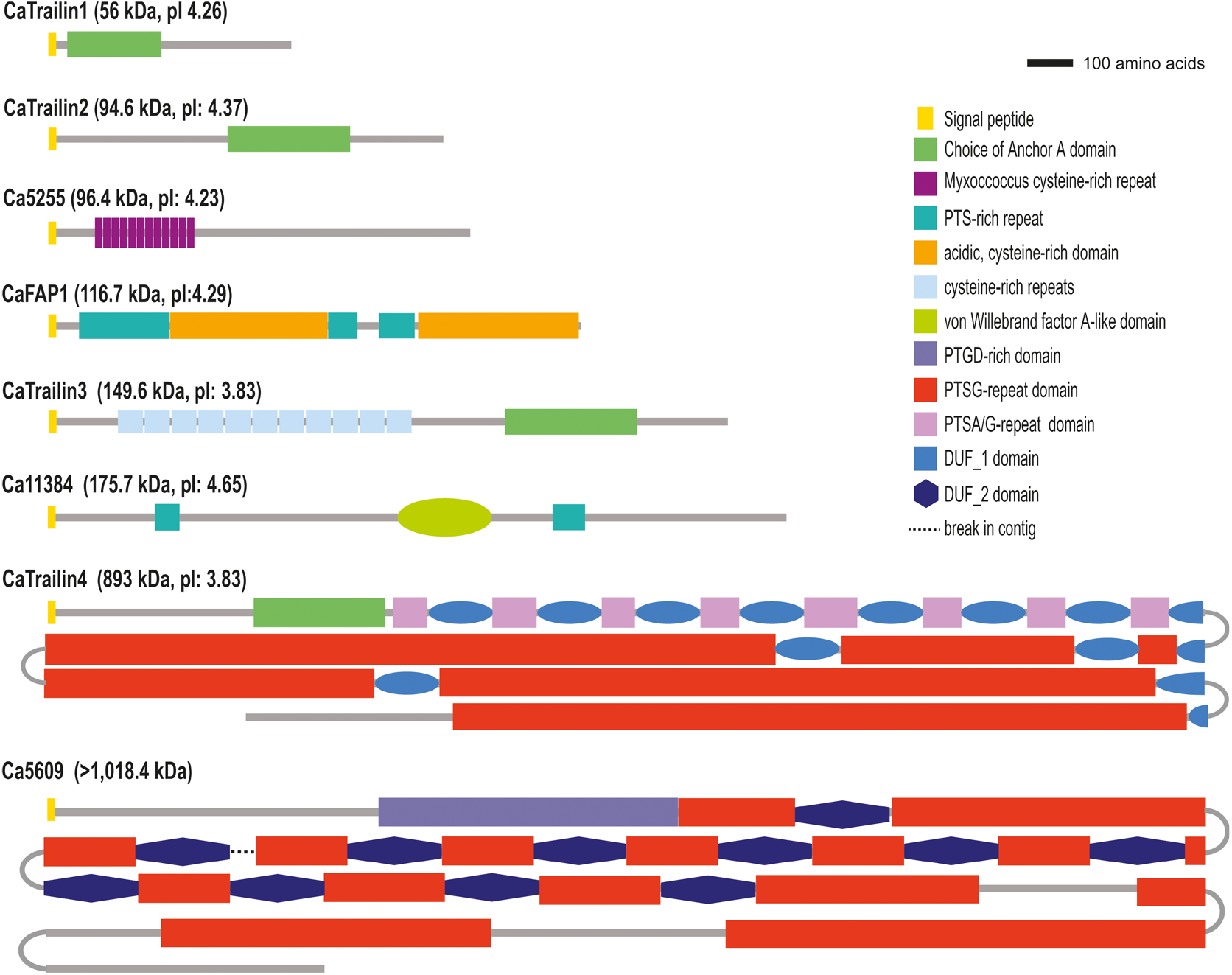
Schematic primary structures of the proteins identified in the *C. australis* adhesive trails. For reasons explained in the text, the gene model of Ca5609 is tentative. The molecular mass (in kDa) and isoelectric point (pI) of each protein were calculated from the polypeptide sequence lacking the predicted signal peptide. DUF = domain of unknown function.

Determining the full-length sequences of CaTrailin4 and Ca5609 from the short-read genome assembly (Poulsen, 2023) was not possible as the two genes were truncated at their 5’-ends within regions encoding long repetitive stretches. Therefore, we combined Pacific Biosciences (PacBio) Single Molecule Real-Time long-read sequencing with the Illumina reads to create a new high quality genome assembly, which spans 88 contigs with an N50 of 1,724,159 bp and a BUSCO completeness score of 97% (Fig. S2, Table S4 and S5). A single continuous 31 kb read allowed for the manual prediction of the CaTrailin4 gene model, which is 28,951 bp long and predicted to be devoid of introns (Fig. S3, Fig. S4). The Ca5609 gene model is split within the tandem repeat region across two different contigs, as not a single PacBio read fully spans the genomic region around this gene (Fig. S5). Therefore, the precise length of the tandem repeat region remains unknown.

To detect individual proteins in the AM, we raised peptide antibodies against CaTrailin-2, −3, and −4. Gel filtration chromatography revealed that the majority of the hydroxylamine solubilized AM exhibited molecular masses >450 kDa (i.e. the molecular mass of the largest standard protein; Fig. S1B). Dot blot analysis with the antibodies demonstrated that CaTrailins-2, −3, and −4 were each present in these very high molecular mass fractions (Fig. S1B, C), yet only CaTrailin4 (894 kDa) would be expected to be present in the >450 kDa fraction. This indicates that CaTrailin3 (95 kDa) and CaTrailin4 (163 kDa) contain either extensive post-translational modifications and/or are engaged in supramolecular complexes.

### Primary structures of the diatom adhesive trail proteins

Sequence analysis of the eight AM proteins (Fig. 1 and Table S2) revealed each to contain a predicted N-terminal signal peptide, as is expected for secreted proteins. None of the proteins contained a GDPH-domain, which was the most abundant diatom-specific sequence feature in the *A. coffeaeformis* adhesive proteins (Lachnit *et al*., 2019). Instead, other protein domains were present as described in the following.

#### Proline-Threonine-Serine (PTS)-rich domains

CaFAP1, CaTrailin4 and Ca5609 contain numerous PTS-repeats that constitute ~23%, ~70% and ~50%, respectively, of their sequence. PTS repeats are a hallmark feature of highly glycosylated extracellular proteins, such as mucins and hydroxyproline-rich-glycoproteins (Perez-Vilar & Hill, 1999; Mathieu-Rivet *et al*., 2020). CaFAP1 has already been shown to be glycoslyated (Chiovitti *et al*., 2003), Therefore, we expect CaTrailin4 and Ca5609 to be highly O-glycosylated within their PTS-rich domains. Previously we demonstrated that the organic component of *C. australis* crude AM contains ~70% carbohydrates and the most abundant amino acids were serine (22%), glycine (18%), threonine (12%), and the proline derivative dihydroxyproline (17%) (total PTSG = 69%) (Poulsen *et al*., 2014). This suggests that CaTrailin4 (78% PTSG) and Ca5609 (69% PTSG) might be the most abundant proteins in the adhesive trails, while CaFAP1 (40% PTSG) may be a minor component.

#### Cys-rich repeats

Regions rich in cysteine residues are present in three of the *C. australis* adhesive proteins (CaFAP1, CaTrailin3 and Ca5255). CaFAP1 has a modular structure of alternating PTS-rich and cysteine-rich domains that resembles mucin-like proteins (Poulsen et al., 2022). CaTrailin3 contains eleven imperfect cysteine-rich repeats, each with six conserved cysteine residues (Table S6). The cysteine-rich repeats of the CaTrailin3 do not match any known protein domains, but are found in several other diatoms (including centrics, which are non-motile) as well as some fungal species (Fig S6).

Ca5255 contains twelve *Myxococcus* cys-rich repeats/domain of unknown function (DUF4215; IPR011936) (Table S6). Although the exact function of this domain is unknown, Myxobacteria are biofilm-forming bacteria that exhibit gliding motility (Nan *et al*., 2010; Faure *et al*., 2016). Myxococcus cysteine-rich repeats are present in numerous bacterial and eukaryotic species, including the diatom *Nitzschia inconspicua*.

#### Other repetitive sequences

The two large PTS-rich proteins CaTrailin4 and Ca5609 also contain repetitive sequences that interspace some of the PTS-rich regions, termed domain of unknown function (DUF) 1 and 2, respectively (Fig. 1, Table S6). Both DUFs contain six cysteine residues that might form disulfide bonds to stabilize a particular fold. Previously the mechanical properties of diatom EPS have been measured using AFM measurements (Dugdale *et al*., 2006) and optical tweezers (Gutierrez-Medina *et al*., 2022), which revealed that the diatom adhesive strands are modular and elastic. Therefore, the interspacing of the repeat regions within the PTS-rich domains may play a role in the mechanical stability/self-healing/elasticity of the polymeric adhesive strands.

#### Choice-of-Anchor A (CAA) domain/pAdhesive_15 (Interpro: IPR026588; Pfam: PF20597)

CAA-domains are present in the four *C. australis* Trailins, and were previously identified in three *A. coffeaeformis* AM proteins (Lachnit *et al*., 2019). The term ‘choice-of-anchor’ refers to the fact that bacterial proteins possessing this domain are surface-exposed proteins, anchored to the cell membrane via one of three conserved transmembrane domain types (LPXTG cell wall anchor, PEP-CTERM domain, or type IX secretion system) (Xu *et al*., 2004). However, the CAA-domain itself is not the cell membrane anchor. The *C. australis* CAA-domain bearing proteins do not contain a membrane anchor domain.

Very recently, the predicted structure of a CAA-domain containing protein from *Bacillus anthracis* (BA_0871 UniProtKB accession no. A0A384LNE7) (Monzon & Bateman, 2022) was determined using Alphafold and described as highly similar to the structure of the ice-binding adhesive domain (IPR021884) and is the founding member a new Pfam family termed pAdhesive_15 (PF20597). However, the precise function of this domain remains unknown as not all bacterial species that possess this domain are aquatic or encountering ice structures.

#### Von Willebrand factor A (vWA)-like domain (IPR036465)

Ca11384 contains a vWA-like domain. This domain is typically involved in the formation of multi-protein complexes in a wide variety of biological processes, including adhesion and extracellular matrix assembly (Colombatti et al., 1993).

### Phylogenetic analysis of the adhesive trail proteins

Phylogenetic analysis revealed that outside the known protein domains (vWA-like, CAA, cysteine-rich repeats) and PTS-rich regions, the eight *C. australis* trail proteins are specific to diatoms (Table S7, Fig. S6) and four are found exclusively in *C. australis* (CaFAP1, Ca5255, CaTrailin1 and CaTrailin2). The C-terminal regions of two proteins are specific to raphid pennates (CaTrailin4 and Ca5609), and are conserved among the orders Bacillariales and Mastogloiales but absent from the Naviculales. Ca11384 shows a similar pattern of homology amongst raphid pennates. The N-terminal domain of CaTrailin3 lacks similarity to pennate sequences but has putative centric homologues. It is notable that none of the proteins share sequence homology along their entire length with proteins from other diatoms.

To investigate the evolutionary origin of the diatom CAA-domains, a hidden Markov Model (HMM) profile was constructed from the *C. australis* and *A. coffeaeformis* CAA-domains from the adhesive trail proteins thereby defining a ‘diatom CAA-domain’. Using this HMM profile, a comprehensive set of homologs from both eukaryotes and prokaryotes was compiled from three protein databases: PLAZA Diatoms 1.0 (Osuna-Cruz *et al*., 2020), NCBI non-redundant database, and the decontaminated MMETSP database (Keeling *et al*., 2014; Van Vlierberghe *et al*., 2021). Combining hits from these sources, we found that CAA-domains are largely restricted to bacteria, rotifers (aquatic invertebrates), and diatoms (Fig. 2A). Among the CAA-containing hits from diatoms, five genes (three from *Fragilariopsis cylindrus*, one from *Pseudo-nitzschia multistriata* and one from *Phaeodactylum tricornutum)* were assessed in a recent study, in which four were shown to be acquired through horizontal gene transfer (HGT) from bacteria (Vancaester *et al*., 2020). Indeed, phylogenetic analysis shows that diatom CAA-domains are nested within bacterial clades, supporting their bacterial origin (Fig. 2A). Notably, diatom CAA-domains cluster into two distinct clades, which we designated type-1 and type-2 (Fig. S7 and S8), suggesting two independent HGT events (Fig. 2A). Mapping the occurrence of these two CAA-domains to the diatom species tree (Fig. 2B) reveals that type-1 CAA-domains are distributed across the major clades of diatoms, suggesting that the domain was acquired by a common ancestor of diatoms. Meanwhile, type-2 CAA-domains likely have a more recent origin, as they are restricted to a subclade of raphid diatoms. Interestingly, dinoflagellate CAA-domains (*Kryptoperidinium foliaceum* and *Durinskia baltica*) are nested within the type-1 raphid clade (Fig. 2B). The origin of these genes could be explained by the fact that both species belong to the so-called dinotoms, a select group of dinoflagellates possessing a chloroplast obtained through tertiary endosymbiosis of a diatom (Imanian *et al*., 2010). Likewise, diatom type-2 CAA-domains were acquired by bdelloid rotifers belonging to the genera *Rotaria, Adineta* and *Didymodactylos* (Fig. S8).

**Figure 2.**
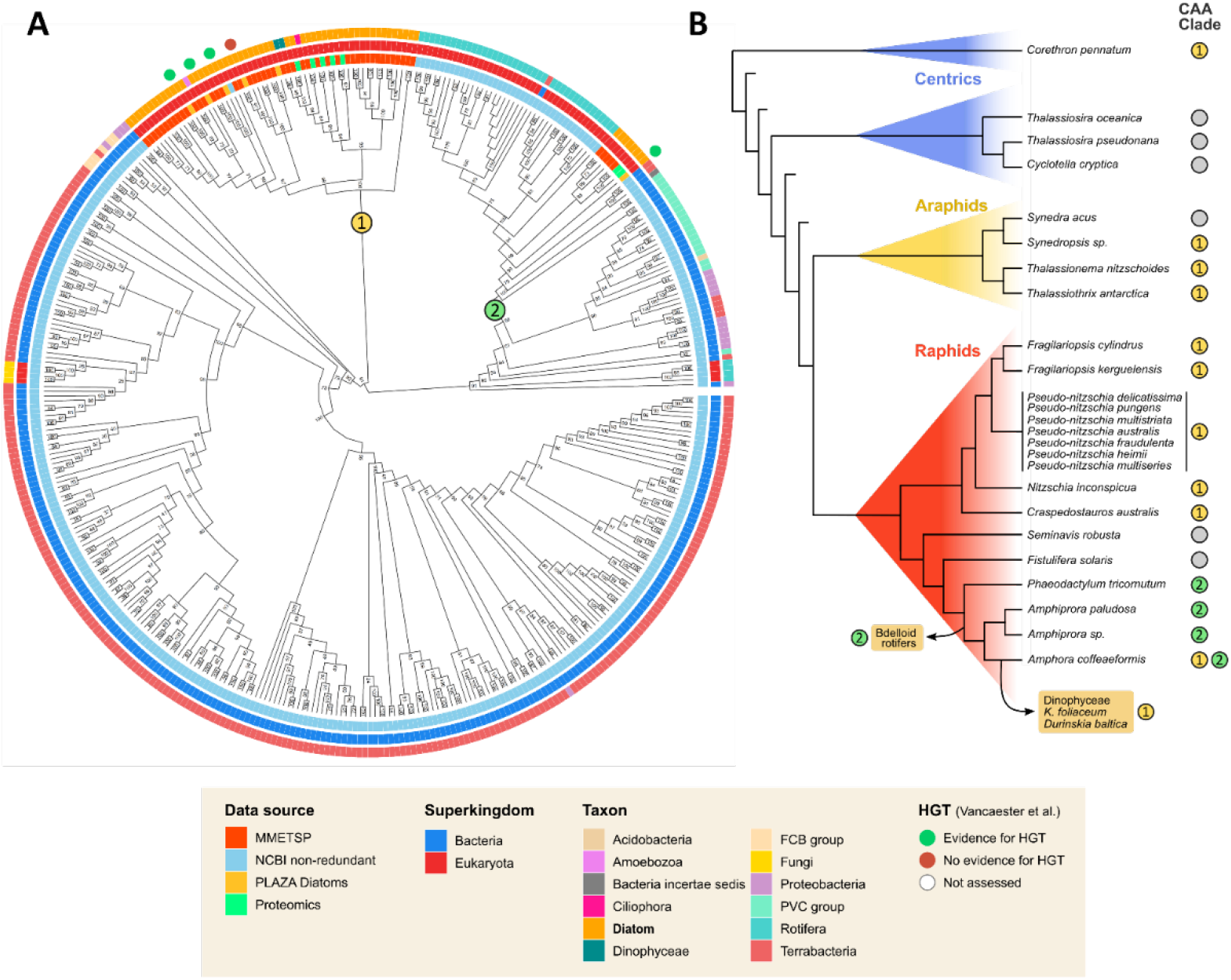
Evolutionary history of diatom CAA-domains. **(A)** Midpoint-rooted maximum-likelihood phylogenetic tree of homologs of the diatom CAA-domain across the tree of life. Node labels indicate bootstrap support. The coloured outer rings indicate the data source and taxonomic binning of each hit. The prediction of possible horizontal gene transfer (HGT) for selected genes from Vancaester et al. (2020) is shown by coloured circles outside the phylogeny. The two separate clades of diatom CAA-domains are indicated with a coloured circle on the branches. **(B)** Species tree showing the distribution of diatom CAA-domains throughout the major clades of diatoms (indicated with coloured triangles). To the right, the presence of CAA-domains in the proteome of each species is indicated, specifying the diatom CAA-domain clade (panel A). Arrows show the putative timing of secondary HGT events from diatoms to other taxa (bdelloid rotifers and dinoflagellates). Phylogenetic relationships of diatoms were parsed from Nakov et al. (2018).

Taken together, two independent transfers of CAA-domains occurred from bacteria into diatoms and were widely retained across diatoms, suggesting these genes are under purifying selection pressure and potentially confer a functional advantage. Coupling the abundance of CAA-domains in the adhesive material of benthic diatoms with their presumed role in adhesion in their bacterial ancestors suggests that the CAA-domain offers novel adhesive functionalities to diatoms. It is noteworthy that type-1 CAA-domains were also found in centric and araphid pennate diatoms (Fig. 2B), so their role is not restricted to gliding diatoms.

### 3D structure prediction of the diatom CAA-domain

Very recently, a large scale bioinformatics study identified 24 clusters of bacterial putative adhesive domains (Monzon & Bateman, 2022), wherein ‘cluster 10’ included a representative of a CAA-domain bearing protein from *Bacillus anthracis* (UniProt: BA_0871) (Xu *et al*., 2004). Using Alphafold, the predicted structure of the CAA-domain (now also termed *pAdhesive_15)* was shown to be similar to an ice-binding protein (mLeIBP) from the fungus *Leucosporidium* sp. (PDB ID 4NUH:A) (Monzon & Bateman, 2022). This is a puzzling result in the context of CAA-domain bearing adhesion proteins from *C. australis* as this diatom species is not found associated with ice in nature. To further investigate this, we submitted the CAA-domain from CaTrailin4 to Alphafold 2 (Jumper *et al*., 2021). The predicted structure of the CaTrailin4 CAA-domain is a β-helical fold consisting of two units (see Fig. 3, Fig. S9) held together by a long alpha helix. Out of the predicted positions of the 4230 atoms, 72% were of high quality and 20% of medium quality, indicating that the predicted structure of the CAA-domain is highly reliable and the predicted topology likely correct (Fig. S9). Only the exact position of a few loops (amino acids 40-50 and 70-75) is uncertain.

**Figure 3.**
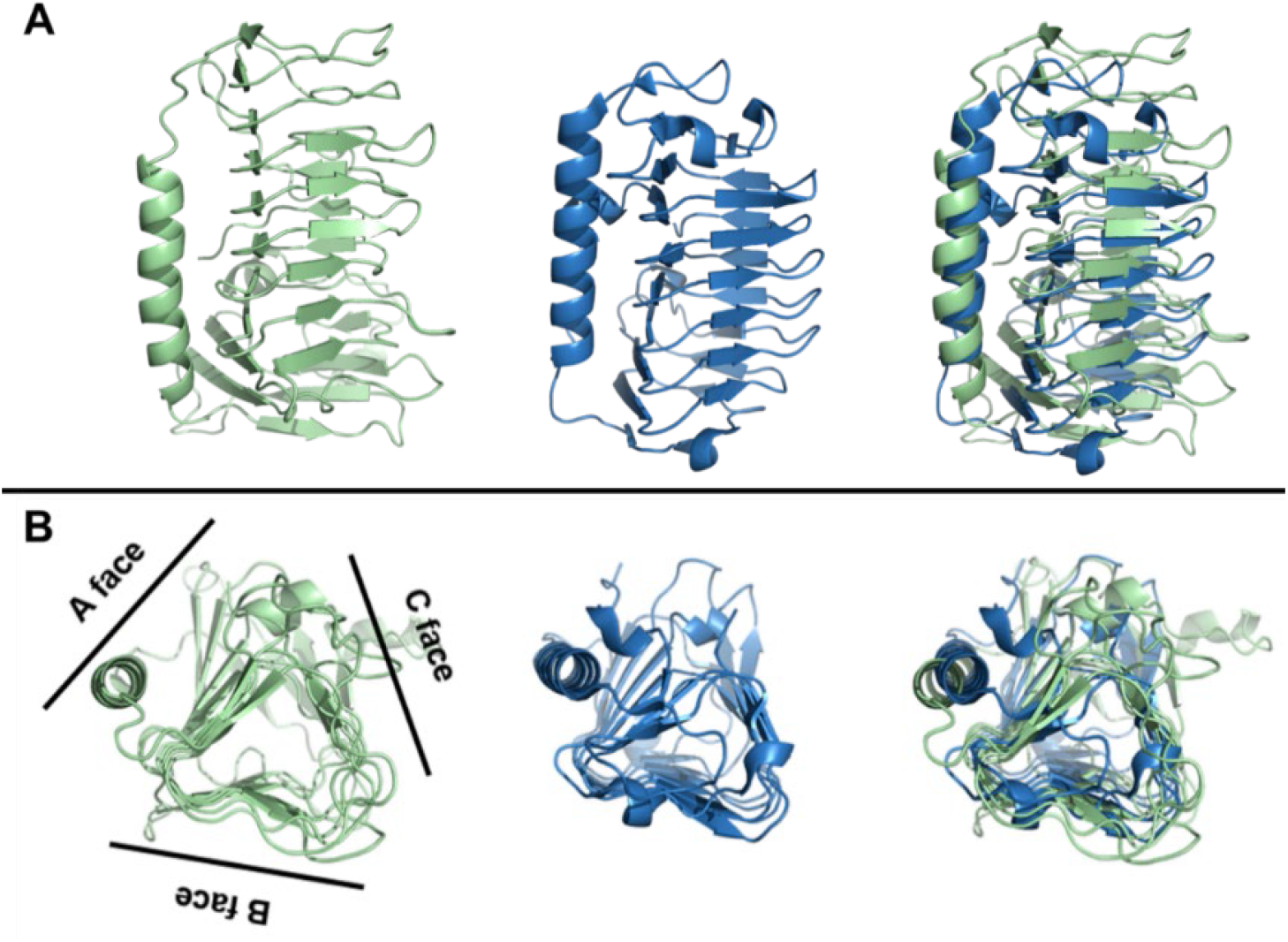
Relationship between 3D structures of CAA-domains and ice binding proteins. (**A**) Left: predicted structure of the CAA-domain from CaTrailin4; middle: experimentally determined structure of the ice-binding protein FfIBP (PDB ID 4NU2); right: Superposition of the CaTrailin4 CAA-domain (green) and FfIBP (blue). (**B**) The CaTrailin4 CAA-domain and FfIBP adopt a β-helical fold with three faces.

The top eight hits from the structural homology search against the experimentally determined 3D structures from the Protein Data Bank (PDB) are listed in Table S8, which align with 70-85% of their residues to the CaTrailin4 CAA with an overall RMSD between 4 and 6Å. The highest structural similarity is to the ice-binding protein FfIBP from the Antarctic bacterium *Flavobacterium frigoris* (see Fig 3) (Do *et al*., 2014). Both structures align with 85% of the residues and a root mean square deviation of 5.18Å. In both, the CaTrailin4 CAA-domain and FfIBP, the β-helical core has three faces and the long alpha helix packs against face A (see Fig 3B). A sequence representation of the structural alignment highlights the lack of sequence similarity (see Fig. S10). Yet, 85% of residues are structurally aligned (Fig. S10, shown in red) including the long helix as well as most of the strands making up the three faces. However, only 25 residues across the full sequences are identical. Interestingly, although seven out of these eight structures are ice-binding proteins from bacteria, fungi and diatoms living in cold environments, the one protein that is not ice-binding, ComZ, is from *Thermus thermophilus*, a bacterium with an optimal growth temperature of about 65°C. ComZ is thought to be located on the pilus tip and to play a role in DNA uptake (Salleh et al., 2019).

The strong structural similarity to ice-binding proteins begs the question as whether the diatom CAA-domains are capable of ice-binding. To address this question, we checked the CAA-domain of CaTrailin4 for the presence of ice-binding motifs (IBMs) (Do *et al*., 2014) and ordered surface carbons (OSCs) (Doxey *et al*., 2006), which are predictive of ice-binding regions. The tetrapeptide sequence T-A/G-X-T/N is present at three locations in FfIBP and regarded as an IBM (Do *et al*., 2014). The three locations align structurally well with the CaTrailin4 CAA-domain, but they are not conserved in sequence, indicating the absence of IBMs. To further corroborate this finding, we screened the CaTrailin4 CAA-domain for OSCs (Doxey *et al*., 2006), which are specific geometric constellations enabling binding to the highly structured and regular surface of an ice crystal. It was shown that the absolute and relative amount of the protein surface areas covered by OSCs is predictive of ice-binding with values of >325Å^2^ and >6%, respectively (Doxey *et al*., 2006). FfIBP meets these criteria, whereas the CAA-domain of CaTrailin4 does not. Therefore, we assume that CaTraillin4 is highly unlikely to serve the purpose of binding to ice. Instead, *C. australis* seems to have adapted this fold for proteins that mediate adhesion to other solid surfaces under water. This assumption is supported by the following observation. Diatoms living in polar habitats, like *Fragilariopsis cylindrus*, produce ice binding proteins (IBPs) that protect them from freezing and also serve to attach cells to ice (Raymond & Kim, 2012; Mock *et al*., 2017; Dorrell *et al*., 2023). Based on sequence analysis *F. cylindrus* contains three proteins with a type-1 CAA-domain (Fig. 2) and more than 50 IBPs (IPR021884) (Osuna-Cruz *et al*., 2020). In contrast, *C. australis* contains no proteins with an IBP domain, but encodes six CAA-domain bearing proteins, of which four were identified here in the adhesive material.

### Trailin Immunolocalization

To gain insight into the Trailins function(s), we analysed their localization on the cell surface, in the adhesive trails, and the biofilm matrix. Initial attempts to express the Trailins (CaTrailin2 and −3) as GFP fusion proteins were unsuccessful. The GFP-fusion proteins remained trapped inside the cell (Fig. S11), which is analogous to previous attempts to visualize putative *P. tricornutum* and *A. coffeaeformis* adhesion proteins (Buhmann *et al*., 2014; Willis *et al*., 2014; Lachnit *et al*., 2019). Therefore, we pursued immunolocalization studies with peptide antibodies raised against CaTrailin2, −3 and −4. As a control we have employed the fucose-specific lectin (*Aleuria aurantia*, AAL) that recognizes the *C. australis* adhesive trails (Fig. 4) (Neu & Kuhlicke, 2017). All three antibodies (αCaTrailin2, −3 and −4) showed specific labelling of adhesive trails (Fig. 4A). αCaTrailin2 and −3 also recognized numerous small particles on the substratum, which may arise from colloidal secreted material that settles from solution (Fig. 4A). The strongest labelling of the adhesive trails was obtained using αCaTrailin4. Control experiments performed using pre-immune serum and secondary antibody alone showed no labelling of the adhesive trails (Fig. S12).

**Figure 4.**
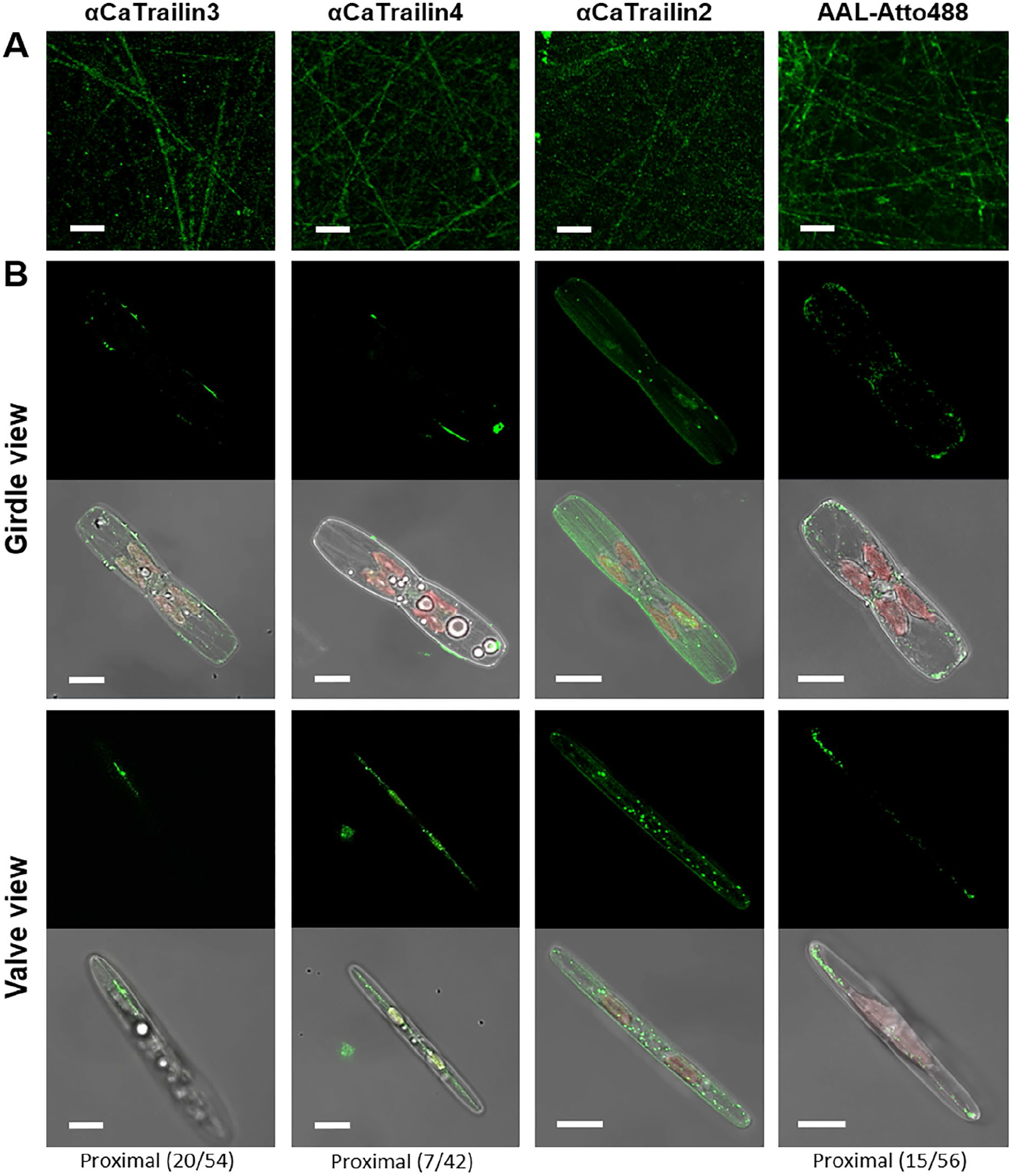
Localization of adhesive proteins. Immunolabelling of *C. australis* **(A)** adhesive trails and **(B)** chemically fixed cells with Trailin-specific antibodies αCaTrailin3, αCaTrailin4 and αCaTrailin2, as well as the fucose-specific lectin (AAL-Atto488). All images are presented as the maximum intensity Z-projection. For valve view only a part of all Z-stacks was used for maximum intensity projection, the position and number of Z-stacks is indicated below the image. The green colour represents the fluorescence signal from either the Atto488 dye (AAL-Atto488) or the AlexaFluor488 secondary antibody. The red colour represents chloroplast autofluorescence. Scale bars: 20 μm (A) and 10 μm (B).

To determine whether the Trailins are secreted together with the adhesive material through the raphe slit, their location on the cell surface was determined. Like the trail-specific lectin AAL, CaTrailin3 and −4 were located only at the raphe slits, whereas CaTrailin2 was present throughout the cell surface but notably absent from the raphe slits (Fig. 4B). The CaTrailin2 location is similar to the cell surface protein CaFAP1, which becomes secondarily associated with the trails after being secreted from the cell through a raphe-independent mechanism (Poulsen, 2023).

There is previous circumstantial evidence that the biomolecular composition of the primary adhesive trails and extracellular matrix of biofilms differ (Smith & Underwood, 1998; Wetherbee *et al*., 1998; Underwood *et al*., 2004; Tong & Derek, 2021). However, the direct visualization of proteins in the diatom biofilm matrix and how these change over time has not yet been accomplished. Here we used immunolabelling with the αCaTrailin2, −3, and −4 antibodies in combination with confocal laser scanning microscopy (CLSM) to analyse their distribution at different stages of biofilm development. To visualize the development of the biofilm, we also employed the fluorescently labelled lectin AAL, which binds to fucose-bearing biopolymers both in the adhesive trails (see Fig. 4A) and the biofilm matrix (Fig. 5). After day 1, AAL-Atto488 labelling revealed numerous, overlapping but discrete straight and curved lines, which are the primary adhesive trails deposited on the substratum by moving diatoms (Fig. 5A, B). The trails generate a meshwork of ~6 μm thickness with occasional patches of globular EPS material deposited on top of the trails or around the cells (Fig. 5B). After day 4, the density of the trail meshwork and the globular EPS material had markedly increased generating a ~10 μm thick layer (Fig. 5E, F). In many places globular EPS material piled up on top of this layer reaching Z-heights of up to 20 μm (Fig. 5E, F). After day 8, the morphology of the biofilm changed, and became dominated by a narrow meshwork of rather short fibres while the long adhesive trails seemed to be absent (Fig. 5I, J). Large EPS agglomerates were abundant throughout the biofilm, and most cells are found in the upper biofilm layer (Fig. 5I, J). At day 12, the biofilm was still composed of a narrow meshwork of short fibres and extensive EPS aggregates up to 30 μm in Z-height and tens of micrometer wide in the X-Y direction (Fig. 5M, N). Both cell-free and cell bearing EPS aggregates were observed (Fig. 5M, N).

**Figure 5.**
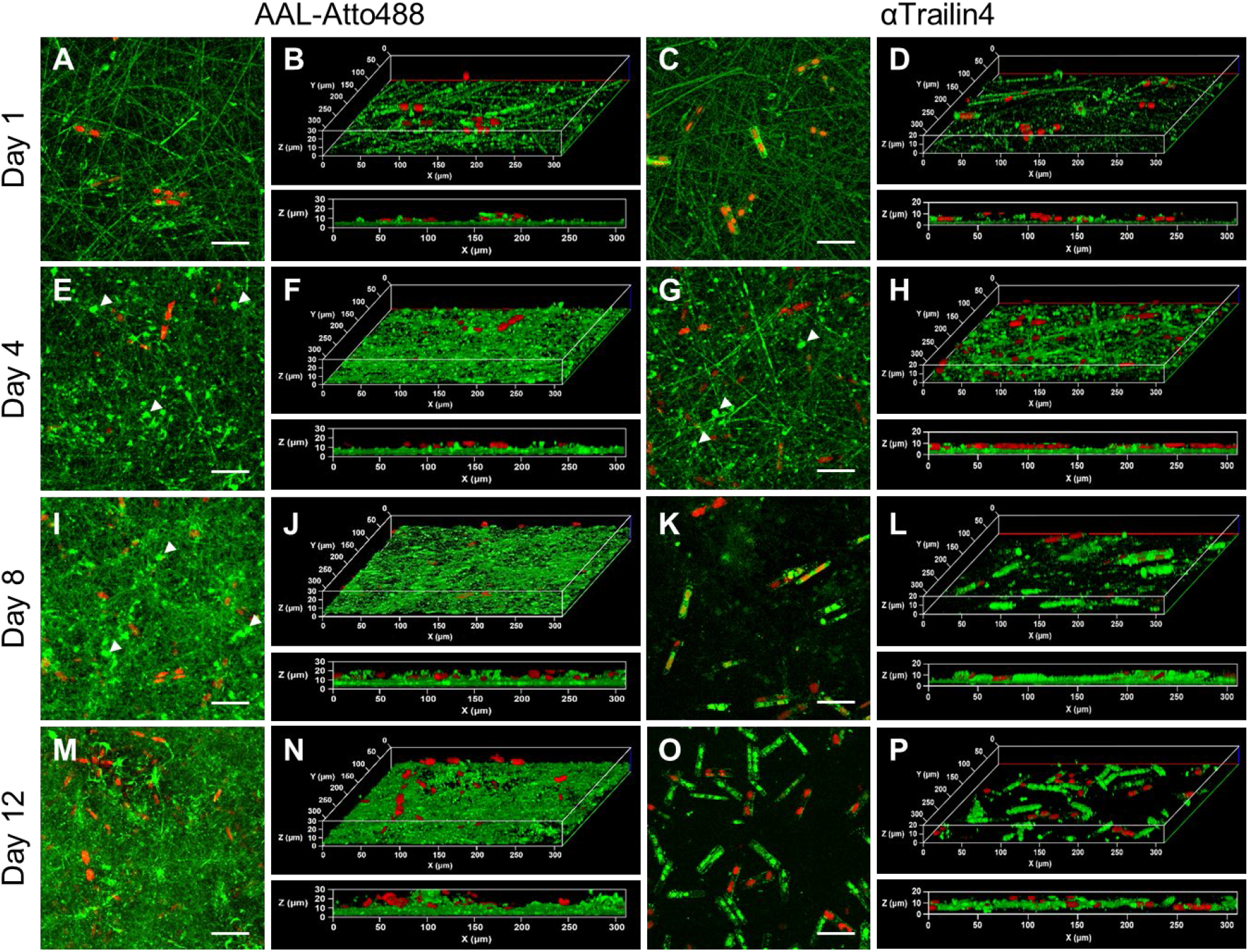
Development of the *C. australis* biofilm matrix. All images are from confocal microscopy and presented as the maximum intensity Z-projection. (**A, C, E, G, I, K, M, O**) top view and (**B, D, F, H, J, L, N, P**) oblique and side-on view of the confocal fluorescence images obtained by probing the biofilm with AAL-Atto488 (left) and αTrailin4 antibodies (right). The days indicated on the left margin indicate the time after seeding the cells on the surface, at which the labeling of the surfaces with the probes indicated on the top margin was performed. White arrow heads depict some of the agglomerates of adhesive material. Green color represents the fluorescence signal from AlexaFluor488 secondary antibody or Atto488 conjugate, and red colour chloroplast autofluorescence. Scale bar: 50 μm, 3D reconstruction: 300 μm (X) x 300 μm (Y) x 20 μm (Z).

The labelling with AAL-Atto488 showed that fucose bearing biopolymers are seemingly continuously produced at all stages of biofilm development. To investigate the contribution of Trailins to biofilm development, we performed separate immunolabelling experiments with αCaTrailin4 (Fig. 5), αCaTrailin2 and αCaTrailin3 (Fig. S13). This demonstrated that the primary adhesive trails were recognized by all three antibodies on days 1 and 4 (Fig. 5C, D, G, H, Fig. S13A-H). At day 4, the results with the αCaTrailin4 labelling was very similar to the AAL lectin, showing adhesive trails and globular agglomerates (Fig. 5G, H), whereas the labelling with αCaTrailin2 and αCaTrailin3 showed mainly adhesive trails (Fig. S13E-H). In striking contrast to the AAL-Atto488 labelling, none of the antibodies labelled the biofilm matrix on days 8 and 12 (Fig. 5K, L, O, P, Fig. S13I-P). Occasionally, trail-like labelling patterns were observed with αCaTrailin3 on day 8, but the labelling was much weaker and infrequent when compared to the previous days (Fig. S13K, L). We were concerned that the absence of antibody labelling on days 8 and 12 might be an artefact caused by the gelatinous nature of the biofilm, slowing down diffusion of the antibodies thereby preventing their binding to the trails on the underlying substratum. To investigate the ability of IgG antibody molecules to diffuse through the biofilm, Protein G beads (~1 μm) that bind IgG molecules were seeded together with the diatoms on a fresh substratum. After 12 days, the secondary antibody (i.e. labelled IgG molecules) were readily able to diffuse through the entire biofilm and bind to the Protein G beads on the substratum (Fig. S14). This demonstrated that the absence of CaTrailins2-4 in the days 8 and 12 biofilms is not caused by their restricted accessibility for the antibody. Instead, we conclude that after day 4 the production of CaTrailins decreases and ceases by day 8, and within the same time period, the proteinaceous component of the CaTrailin containing primary trails become proteolytically degraded. A final control experiment using the pre-immune serum revealed that the strong green fluorescence of cells seen in days 8 and 12 is a result of cell death and not due to binding of the primary or secondary antibody (Fig. S12). We observed that upon cell death the chloroplast autofluorescence shifts towards the blue spectrum and is visible in the green channel. These dying cells may therefore be the source of enzymes that degrade the proteinaceous component of the primary adhesive trails.

Taken together, these results indicate that diatom biofilm formation is a two-step process: (i) EPS secreted from the raphe-slit forms the initial adhesive material that allows for colonization of the underlying substratum, followed by (ii) continuous EPS production that expands the matrix three-dimensionally allowing for a permanent binding phase. Our results indicate that there is a switch in the biochemical composition of the initial adhesive to the biofilm matrix.

## Discussion

In this study, we describe the discovery, structure and localization of novel adhesive trail proteins involved in gliding motility of the model adhesion diatom *C. australis*. Combined with the recently described adhesive trail proteins from *A. coffeaeformis* (Lachnit *et al*., 2019) our studies reveal that each species contains a unique set of adhesive trail proteins that are modular, contain PTS-rich regions and the presence of CAA and/or GDPH-domains. Using long-read PacBio genome sequencing we discovered extremely large (>1 MDa), repetitive, PTS-rich proteins that we suspect to be highly glycosylated as the isolated crude adhesive material contains roughly 70% (w/w) carbohydrates (Poulsen *et al*., 2014). The occurrence of extremely large, repetitive proteins in underwater adhesives is not unique to diatoms. For example, Stewart and co-workers identified a ~650 kDa adhesive silk protein, H-fibroin, from caddisflies (Frandsen *et al*., 2019). The use of long read sequencing technologies in the latter study and our present work illustrate the potential of these technologies to identify and characterize high molecular weight, complex adhesive proteins that have evaded detection using traditional sequencing technologies.

Here we report the finding that diatoms have acquired CAA-domains from bacteria through horizontal gene transfer (HGT). Throughout their evolutionary history, HGT has allowed diatoms to expand their ability to adapt to different ecological niches. (Bowler *et al*., 2008; Vancaester *et al*., 2020; Dorrell *et al*., 2021). For example, the acquisition of bacterial genes and their integration in several metabolic pathways such as the ornithine-urea cycle and vitamin B12 uptake has contributed to their ability to outcompete other phytoplankton (Bowler *et al*., 2008; Vancaester *et al*., 2020; Dorrell *et al*., 2021). Our observation that the CAA-domain was acquired at least twice independently suggests that is it clearly beneficial for diatoms. As the CAA-domain is present in numerous, but not all raphid diatoms, it not essential for gliding motility. Recent studies suggest that HGT is continuously occurring in diatoms and that there is a high rate of HGT loss due constant environmental pressures and ecological niche adaption (Dorrell *et al*., 2021; Dorrell *et al*., 2023). Of particular note is the recent description of Antarctic vs Arctic clades of diatom IBPs that suggests two independent HGT events for these two Polar Regions (Dorrell *et al*., 2023). This genome plasticity may therefore help to explain our inability to identify a single ‘universal’ adhesive protein employed by motile diatoms as HGT acquisitions and losses occurring in their local habitat may have allowed for the evolution of a diverse set of species-specific adhesive proteins. In this context it is interesting to note that the diatom-specific GDPH-domain, which we previously identified in the *A. coffeaeformis* adhesive trail proteins (Lachnit *et al*., 2019), is present in many but not all raphid diatom species. The GDPH-domain is absent from raphid diatoms that contain proteins with a type-1 CAA-domain, while raphid diatoms that possess GDPH-domain bearing proteins contain proteins that have the type-2 CAA-domain or completely lack CAA-domain bearing proteins. This observation might indicate that diatom adhesion and gliding can be accomplished by three different types of protein assemblies that contain (i) type-2 CAA-domains and GDPH-domains, or (ii) type-1 CAA-domains, or (iii) only GDPH-domains. Furthermore, based on the current available databases the type-1 CAA-domain is also present in a single centric diatom (*Corethron*) and some araphid species. Therefore, we suggest that in non-motile species the CAA-domain may play a role in other adhesive functions such as mucilage pads, chain-formation or cell aggregation (Edgar & Pickett-Heaps, 1984; Hoagland *et al*., 1993; Thornton, 2002).

The protein structure prediction revealed that the CAA-domain shares structural but not sequence similarity with ice-binding proteins from bacteria and eukaryotes as well as the ComZ pilus protein from a thermophilic bacteria. It may seem paradoxical that this protein structure is present in both psychrophilic and thermophilic organisms as well as diatoms from temperate habitats, yet all these organisms share the ability to adhere to surfaces underwater. Interestingly, *F. cylindrus*, which is a sea-ice diatom, possesses both true ice-binding proteins as well as proteins bearing the CAA-domain (Krell *et al*., 2008; Mock *et al*., 2017). Although the exact ice-binding mechanism of proteins that possess this β-solenoid protein structure is unknown it has been suggested that they are able to order surface-associated water molecules, which are fine-tuned to merge with the water molecules at the ice-water interface (Yamauchi *et al*., 2020; Khan *et al*., 2021). Site-directed mutagenesis studies demonstrated the importance of threonine-rich repeats in ice-binding. Although the diatom CAA-domains exhibit an almost identical β-solenoid protein structure to ice-binding proteins, they are not predicted to be ice-binding as they lack the threonine-rich ice-binding sites on the flat surface of one of the β-sheets. Nevertheless, this highly conserved protein structure might endow these diatom proteins with the ability to order water molecules allowing for the alignment of protein surface and the interfacial water molecules on any surface underwater. Future studies using recombinant CAA-domains are needed to test the hypothesis that they play a role in the underwater adhesion of diatoms.

One of the most puzzling aspects of the current model for diatom gliding is that the adhesive material secreted through the raphe slits functions as both an adhesive and a transducer of the actomyosin-generated force from inside the cell to the substratum. To achieve this dual function, the adhesive material needs to reach from the plasma membrane through the raphe slit to the underlying substratum, which equals a distance ≥1 μm in *C. australis* and many other diatoms. It was previously proposed that proteins with high tensile strength and elasticity would be suitable to fulfil this function (Dugdale *et al*., 2006), prompting us to speculate that the extremely large, modular proteins in the *C. australis* adhesive trails, CaTrailin4 and Ca5609, might be such proteins. The repetitive PST-rich domains in these proteins are predicted to be intrinsically disordered (Table S4), and we regard it very likely that they are highly O-glycosylated, like many other PTS-rich extracellular proteins from algae (Hallmann, 2003; Tatli *et al*., 2018; Mathieu-Rivet *et al*., 2020). The glycan moieties may be highly negatively charged due to the presence of uronic acids, which are abundant in the *C. australis* adhesive material (Poulsen et al., 2014). A high negative charge density would, due to electrostatic repulsion, stretch out the PTS polypeptide regions, which in the case of CaTrailin4 could span ~2.0 μm (~6,500 aa, 3.5 Å per amino acid (Ainavarapu *et al*., 2007)). This distance is in fact very close to the measured length of EPS strands from *C. australis* (3.5 ±1.2 μm) (Higgins *et al*., 2003) and *Nitszschia communis* (Gutierrez-Medina *et al*., 2022). Recently, a 1.5 MDa adhesive protein (*Mp*IBP) from an arctic bacteria (*Marinomonas primoryensis*) was shown to be 600 nanometers long, with an exceptionally long extender region (RII) composed of ~120 tandem Ig-like domains that serves to project the adhesive regions of the protein into the medium (Guo *et al*., 2017). Furthermore, it has also been demonstrated that the ‘elastic reach’ of cell adhesion molecules (i.e. the distance between the two contact sites that the protein can be displaced without breaking) can depend on the number of tandem IgG repeats (Carl *et al*., 2001). The repetitive tandem repeats of in CaTrailin4 (DUF1) and Ca5609 (DUF2) might serve an analogous role enabling the reversible mechanical extensibility of these proteins. To investigate the structure-property relationship in CaTrailin4 and Ca5609, the native proteins need to be isolated from the solubilized adhesive material whose preparation was established in the present work.

From the moment a surface is submerged in an aquatic habitat, biofilm-forming organisms will attach and commence the production of a 3D extracellular matrix (Callow & Callow, 2011; de Carvalho, 2018). While such biofilms have important roles in biogeochemical cycling and ecological functions of benthic communities, the accumulation of microbial biofilms (termed biofouling) on manufactured structures (e.g. ship hulls, piping, aquaculture nets) continues to be a very costly problem. A ‘heavy’ biofilm, which describes the condition where the underlying paint colour is difficult or impossible to determine, increases the drag forces on a ship’s hull up to 18%, resulting in enhanced fuel consumption (Schultz *et al*., 2011). Mitigating the detrimental effects (e.g. corrosion, increased CO2 emissions) and costs associated with biofouling requires an understanding of the adhesive biomolecules to develop a targeted approach to prevent initial adhesion. In this study, we have demonstrated that there appears to be a molecular switch between the biomolecules required for initial adhesion to those present in the mature biofilm matrix. The components of the primary adhesives, particularly the widespread CAA domain, might be suitable targets for the development of anti-fouling coatings that prevent the adhesion of a wide variety of diatoms and possibly also bacteria. The discoveries described here are the fundament to achieve a detailed mechanistic understanding of how diatoms accomplish their remarkable underwater adhesion and establishing a biofilm community.

## Acknowledgments

We thank Jennifer Klemm for technical support, Stefan Görlich for his initial assistance with the structural protein analysis, Stefan Golfier for critically reading the manuscript, and the CMCB Technology Platform at TU Dresden (Light Microscopy Facility, Molecular Analysis/Mass Spectrometry Facility) and the Dresden-concept Genome Center. This work was supported by grants from the Deutsche Forschungsgemeinschaft (PO 2256/1-1) to N.P., (KR 1853/9-1) to N.K. G.B. is a postdoctoral fellow supported by Fonds Wetenschappelijk Onderzoek (FWO, 1228423N). K.V. and W.V. want to acknowledge the funding obtained by the BOF project GOA01G01323. The Molecular Analysis/Mass Spectrometry Facility was supported by a grants from the Deutsche Forschungsgemeinschaft (INST 269/731-1 FUGG), from the German Federal Ministry of Education and Research (#03Z22EB1), and the European Regional Development Fund (ERDF/EFRE, Contract #100232736).

## Author contributions

NP and NK conceived the project.

NP, NK, JZS designed research

NP, BR, JZS, GB, AF, AS, MP performed experiments

NP, JZS, BR, GB, AF, MS, MP, AS, WV, KS, NK analysed the data

NP, NK, JZS, GB, MS wrote the manuscript

All authors contributed to the corrections of the manuscript

## Data availability

The data that support the finding of this study are available in the Supporting Information of this article. The proteomics data are deposited on PRIDE (accession: PXD039465). The sequencing data of this study are available through NCBI (Bioproject:PRJNA850956)

## References

Ainavarapu RK, Brujic J, Huang HH, Wiita AP, Lu H, Li LW, Walther KA, Carrion-Vazquez M, Li HB, Fernandez JM. 2007. Contour length and refolding rate of a small protein controlled by engineered disulfide bonds. Biophysical Journal 92: 225–233.

Ashworth MP. 2013. Rock snot in the age of transcriptomes: Using a phylogenetic framework to identify genes involved in diatom extracellular polymeric substance-secretion pathways. PhD thesis, University of Texas at Austin Austin, Texas, USA.

Bowler C, Allen AE, Badger JH, Grimwood J, Jabbari K, Kuo A, Maheswari U, Martens C, Maumus F, Otillar RP, et al. 2008. The *Phaeodactylum* genome reveals the evolutionary history of diatom genomes. Nature 456: 239–244.

Buhmann MT, Poulsen N, Klemm J, Kennedy MR, Sherrill CD, Kröger N. 2014. A tyrosine-rich cell surface protein in the diatom *Amphora coffeaeformis* identified through transcriptome analysis and genetic transformation. PLoS ONE 9: e110369.

Callow JA, Callow ME. 2011. Trends in the development of environmentally friendly fouling-resistant marine coatings. Nature Commununications 2: 244.

Carl P, Kwok CH, Manderson G, Speicher DW, Discher DE. 2001. Forced unfolding modulated by disulfide bonds in the Ig domains of a cell adhesion molecule. Proceedings of the National Academy of Sciences of the United States of America 98: 1565–1570.

Chiovitti A, Bacic A, Burke J, Wetherbee R. 2003. Heterogeneous xylose-rich glycans are associated with extracellular glycoproteins from the biofouling diatom *Craspedostauros australis* (Bacillariophyceae). European Journal of Phycology 38: 351–360.

Colombatti A, Bonaldo P, Doliana R. 1993. Type A modules: interacting domains found in several non-fibrillar collagens and in other extracellular matrix proteins. Matrix 13: 297–306.

de Carvalho CCCR. 2018. Marine biofilms: A successful microbial strategy with economic implications. Frontiers in Marine Science 5: 126.

Do H, Kim SJ, Kim HJ, Lee JH. 2014. Structure-based characterization and antifreeze properties of a hyperactive ice-binding protein from the Antarctic bacterium *Flavobacterium frigoris* PS1. Acta Crystallographica Section D: Structural Biology 70: 1061–1073.

Dorrell RG, Kuo A, Fussy Z, Richardson EH, Salamov A, Zarevski N, Freyria NJ, Ibarbalz FM, Jenkins J, Pierella Karlusich JJ, et al. 2023. Convergent evolution and horizontal gene transfer in Arctic Ocean microalgae. Life Science Alliance 6: e202201833.

Dorrell RG, Villain A, Perez-Lamarque B, Audren de Kerdrel G, McCallum G, Watson AK, Ait-Mohamed O, Alberti A, Corre E, Frischkorn KR, et al. 2021. Phylogenomic fingerprinting of tempo and functions of horizontal gene transfer within ochrophytes. Proceedings of the National Academy of Sciences of the United States of America 118: e2009974118.

Doxey AC, Yaish MW, Griffith M, McConkey BJ. 2006. Ordered surface carbons distinguish antifreeze proteins and their ice-binding regions. Nature Biotechnology 24: 852–855.

Dugdale TM, Willis A, Wetherbee R. 2006. Adhesive modular proteins occur in the extracellular mucilage of the motile, pennate diatom Phaeodactylum tricornutum. Biophysical Journal 90: L58–L60.

Edgar LA. 1983. Mucilage Secretions of Moving Diatoms. Protoplasma 118: 44–48.

Edgar LA, Pickett-Heaps J. 1984. Diatom Locomotion. In: Round FE, Chapman DJ eds. Progress in Phycological Research. Bristol: Biopress Ltd, 47–88.

Edgar LA, Zavortink M. 1983. The Mechanism of Diatom Locomotion. II: Identification of Actin. Proceedings of the Royal Society of London. Series B, Biological Sciences 218: 345–348.

Faure LM, Fiche JB, Espinosa L, Ducret A, Anantharaman V, Luciano J, Lhospice S, Islam ST, Treguier J, Sotes M, et al. 2016. The mechanism of force transmission at bacterial focal adhesion complexes. Nature 539: 530–535.

Flemming HC, Wingender J. 2010. The biofilm matrix. Nature Reviews Microbiology 8: 623–633.

Flemming HC, Wuertz S. 2019. Bacteria and archaea on Earth and their abundance in biofilms. Nature Reviews Microbiology 17: 247–260.

Frandsen PB, Bursell MG, Taylor AM, Wilson SB, Steeneck A, Stewart RJ. 2019. Exploring the underwater silken architectures of caddisworms: comparative silkomics across two caddisfly suborders. Philosophical Transactions of the Royal Society B-Biological Sciences 374: 20190206.

Guo S, Stevens CA, Vance TDR, Olijve LLC, Graham LA, Campbell RL, Yazdi SR, Escobedo C, Bar-Dolev M, Yashunsky V, et al. 2017. Structure of a 1.5-MDa adhesin that binds its Antarctic bacterium to diatoms and ice. Science Advances 3: e1701440.

Gutierrez-Medina B, Pena Maldonado AI, Garcia-Meza JV. 2022. Mechanical testing of particle streaming and intact extracellular mucilage nanofibers reveal a role of elastic force in diatom motility. Physical Biology. 19: 056002.

Hallmann A. 2003. Extracellular matrix and sex-inducing pheromone in Volvox. International Review of Cytology 227: 131–182.

Harrison PJ, Waters RE, Taylor FJR. 1980. A broad-spectrum artificial seawater medium for coastal and open ocean phytoplankton. Journal of Phycology 16: 28–35.

Higgins MJ, Crawford SA, Mulvaney P, Wetherbee R. 2000. The topography of soft, adhesive diatom ‘trails’ as observed by atomic force microscopy. Biofouling 16: 133–139.

Higgins MJ, Molino P, Mulvaney P, Wetherbee R. 2003. The structure and nanomechanical properties of the adhesive mucilage that mediates diatom-substratum adhesion and motility. Journal of Phycology 39: 1181–1193.

Hoagland KD, Rosowski JR, Gretz MR, Roemer SC. 1993. Diatom extracellular polymeric substances - Function, fine-structure, chemistry, and physiology. Journal of Phycology 29: 537–566.

Holland R, Dugdale TM, Wetherbee R, Brennan AB, Finlay JA, Callow JA, Callow ME. 2004. Adhesion and motility of fouling diatoms on a silicone elastomer. Biofouling 20: 323–329.

Imanian B, Pombert JF, Keeling PJ. 2010. The complete plastid genomes of the two ‘Dinotoms’ *Durinskia baltica* and *Kryptoperidinium foliaceum*. PLoS ONE 5: e10711.

Jumper J, Evans R, Pritzel A, Green T, Figurnov M, Ronneberger O, Tunyasuvunakool K, Bates R, Zidek A, Potapenko A, et al. 2021. Highly accurate protein structure prediction with AlphaFold. Nature 596: 583–589.

Keeling PJ, Burki F, Wilcox HM, Allam B, Allen EE, Amaral-Zettler LA, Armbrust EV, Archibald JM, Bharti AK, Bell CJ, et al. 2014. The Marine Microbial Eukaryote Transcriptome Sequencing Project (MMETSP): illuminating the functional diversity of eukaryotic life in the oceans through transcriptome sequencing. PLoS Biology 12: e1001889.

Khan NMU, Arai T, Tsuda S, Kondo H. 2021. Characterization of microbial antifreeze protein with intermediate activity suggests that a bound-water network is essential for hyperactivity. Scientific Reports 11: 5971.

Krell A, Beszteri B, Dieckmann G, Glockner G, Valentin K, Mock T. 2008. A new class of ice-binding proteins discovered in a salt-stress-induced cDNA library of the psychrophilic diatom *Fragilariopsis cylindrus* (Bacillariophyceae). European Journal of Phycology 43: 423–433.

Lachnit M, Buhmann MT, Klemm J, Kroger N, Poulsen N. 2019. Identification of proteins in the adhesive trails of the diatom *Amphora coffeaeformis*. Philosophical Transactions of the Royal Society B-Biological Sciences 374: 20190196.

Lind JL, Heimann K, Miller EA, vanVliet C, Hoogenraad NJ, Wetherbee R. 1997. Substratum adhesion and gliding in a diatom are mediated by extracellular proteoglycans. Planta 203: 213–221.

Mathieu-Rivet E, Mati-Baouche N, Walet-Balieu ML, Lerouge P, Bardor M. 2020. N-and O-Glycosylation Pathways in the Microalgae Polyphyletic Group. Frontiers in Plant Science 11: 609993.

Mock T, Otillar RP, Strauss J, McMullan M, Paajanen P, Schmutz J, Salamov A, Sanges R, Toseland A, Ward BJ, et al. 2017. Evolutionary genomics of the cold-adapted diatom *Fragilariopsis cylindrus*. Nature 541: 536–540.

Monzon V, Bateman A. 2022. Large-scale discovery of microbial fibrillar adhesins and identification of novel members of adhesive domain families. Journal of Bacteriology 204: e0010722.

Nakov T, Beaulieu JM, Alverson AJ. 2018. Accelerated diversification is related to life history and locomotion in a hyperdiverse lineage of microbial eukaryotes (Diatoms, Bacillariophyta). New Phytologist 219: 462–473.

Nan B, Mauriello EM, Sun IH, Wong A, Zusman DR. 2010. A multi-protein complex from *Myxococcus xanthus* required for bacterial gliding motility. Molecular Microbiology 76: 1539–1554.

Neu TR, Kuhlicke U. 2017. Fluorescence lectin bar-coding of glycoconjugates in the extracellular matrix of biofilm and bioaggregate forming microorganisms. Microorganisms 5: 5.

Osuna-Cruz CM, Bilcke G, Vancaester E, De Decker S, Bones AM, Winge P, Poulsen N, Bulankova P, Verhelst B, Audoor S, et al. 2020. The *Seminavis robusta* genome provides insights into the evolutionary adaptations of benthic diatoms. Nature Communications 11: 3320.

Perez-Vilar J, Hill RL. 1999. The structure and assembly of secreted mucins. Journal of Biological Chemistry 274: 31751–31754.

Pinckney JL. 2018. A mini-review of the contribution of benthic microalgae to the ecology of the continental shelf in the south atlantic bight. Estuaries and Coasts 41: 2070–2078.

Poulsen N, Hennig H, Geyer VF, Diez S, Wetherbee R, Fitz-Gibbon S, Pellegrini M, Kröger N. 2023. On the role of cell surface associated, mucin-like glycoproteins in the pennate diatom *Craspedostauros australis* (Bacillariophyceae). Journal of Phycology 59:54–69.

Poulsen N, Kröger N, Harrington MJ, Brunner E, Paasch S, Buhmann MT. 2014. Isolation and biochemical characterization of underwater adhesives from diatoms. Biofouling 30: 513–523.

Poulsen NC, Spector I, Spurck TP, Schultz TF, Wetherbee R. 1999. Diatom gliding is the result of an actin-myosin motility system. Cell Motility and the Cytoskeleton 44: 23–33.

Raymond JA, Kim HJ. 2012. Possible role of horizontal gene transfer in the colonization of sea ice by algae. PLoS ONE 7: e35968.

Schultz MP, Bendick JA, Holm ER, Hertel WM. 2011. Economic impact of biofouling on a naval surface ship. Biofouling 27: 87–98.

Shevchenko A, Tomas H, Havlis J, Olsen JV, Mann M. 2006. In-gel digestion for mass spectrometric characterization of proteins and proteomes. Nature Protocols 1: 2856–2860.

Smith DJ, Underwood GJC. 1998. Exopolymer production by intertidal epipelic diatoms. Limnology and Oceanography 43: 1578–1591.

Steele DJ, Franklin DJ, Underwood GJC. 2014. Protection of cells from salinity stress by extracellular polymeric substances in diatom biofilms. Biofouling 30: 987–998.

Tatli M, Ishihara M, Heiss C, Browne DR, Dangott LJ, Vitha S, Azadi P, Devarenne TP. 2018. Polysaccharide associated protein (PSAP) from the green microalga *Botryococcus braunii* is a unique extracellular matrix hydroxyproline-rich glycoprotein. Algal Research-Biomass Biofuels and Bioproducts 29: 92–103.

Thornton DCO. 2002. Diatom aggregation in the sea: mechanisms and ecological implications. European Journal of Phycology 37: 149–161.

Tong CY, Derek CJC. 2021. The role of substrates towards marine diatom *Cylindrotheca fusiformis* adhesion and biofilm development. Journal of Applied Phycology 33: 2845–2862.

Underwood GJC, Boulcott M, Raines CA, Waldron K. 2004. Environmental effects on exopolymer production by marine benthic diatoms: Dynamics, changes in composition, and pathways of production. Journal of Phycology 40: 293–304.

Van Vlierberghe M, Di Franco A, Philippe H, Baurain D. 2021. Decontamination, pooling and dereplication of the 678 samples of the Marine Microbial Eukaryote Transcriptome Sequencing Project. BMC Research Notes 14: 306.

Vancaester E, Depuydt T, Osuna-Cruz CM, Vandepoele K. 2020. Comprehensive and functional analysis of horizontal gene transfer events in diatoms. Molecular Biology and Evolution 37: 3243–3257.

Wetherbee R, Lind JL, Burke J, Quatrano RS. 1998. The first kiss: Establishment and control of initial adhesion by raphid diatoms. Journal of Phycology 34: 9–15.

Willis A, Eason-Hubbard M, Hodson O, Maheswari U, Bowler C, Wetherbee R. 2014. Adhesion molecules from the diatom *Phaeodactylum tricornutum* (Bacillariophyceae): genomic identification by amino-acid profiling and in vivo analysis. Journal of Phycology 50(5): 837–849.

Xu Y, Liang XW, Chen YH, Koehler TM, Hook M. 2004. Identification and biochemical characterization of two novel collagen binding MSCRAMMs of Bacillus anthracis. Journal of Biological Chemistry 279: 51760–51768.

Yamauchi A, Arai T, Kondo H, Sasaki YC, Tsuda S. 2020. An ice-binding protein from an Antarctic ascomycete is fine-tuned to bind to specific water molecules located in the ice prism planes. Biomolecules 10: 759.

