## Supplementary file for "Diatom adhesive trail proteins acquired by horizontal gene transfer from bacteria serve as primers for marine biofilm formation"

### Supplementary Information

#### Supplementary Material & Methods

##### Gel filtration chromatography

The adhesive material (~5.0 mg) was solubilized in 1 mL of 0.5 M borate buffer (pH 8.5) containing 2 M hydroxylamine (Sigma Aldrich) and incubated for 3 h at 45°C. The solubilized material was centrifuged for 10 min at 20,000 xg and then loaded on a HiLoad 16/600 Superdex 200p column (GE Healthcare) that had been pre-equilibrated in 500 mM NaCl/50 mM Tris-HCl pH 7.5 at a flow rate of 1 mL/min. The elution of the adhesive components was monitored at 226 and 280 nm and 2.0 mL fractions were collected. The resulting fractions were pooled and concentrated 8-fold using an Amicon Ultra 4 Centrifugal Filter Unit and then 20 µL of each pooled fraction was analyzed on a 6% Schägger Gel and stained with Stains All. A size calibration was performed under the same conditions using protein molecular weight standards (Serva, Germany, cat. No. 39064).

##### Anhydrous hydrofluoric acid treatment

To deglycosylate the AM, the freeze-dried hydroxylamine solubilized material was resuspended in 1 mL freshly condensed anhydrous hydrofluoric acid (HF) and incubated on ice for 1 h. The HF was evaporated using a gentle nitrogen stream followed by a final drying step in a UNIVAPO vacuum concentrator centrifuge (25°C, -0.8 bar, 1 h). The dry samples were immediately dissolved in 50 mM ammonium acetate and when necessary, the pH was neutralized with ammonium hydroxide.

##### LC-MS/MS proteomics analysis

The desalted adhesive material was submitted to Proteomics Facility at Max Planck Institute of Molecular Cell Biology and Genetics in Dresden for reversed phase liquid chromatography nanospray tandem mass spectrometry (LC-MS/MS). The desalted AM was submitted as two samples: HF and non-HF treated. The recovered peptides were dissolved in 20 µL of 5% aqueous formic acid and a 5 µL aliquot was analyzed by GeLC–MS/MS on an Ultimate3000 nanoLC system (Dionex, Amsterdam, The Netherlands) interfaced on-line to a LTQ Orbitrap Velos hybrid tandem mass spectrometer (Thermo Fisher, Bremen, Germany) applying a 180 min elution program. Acquired spectra were searched against 3’-frame translated *C. australis* transcriptome (18699 entries) (Poulsen, 2023) using Mascot software (Matrix Science,

London, UK, v.2.2.04). The hits were evaluated by Scaffold program v.4.3.4 (Proteome Software Inc., Portland, OR) and identifications were accepted if two of more peptides could be established at greater probability than 95% (Peptide Prophet Algorithm) and 99% (Protein Prophet Algorithm) for peptides and proteins, respectively. False Discovery Rate (FDR) was below 1.5% as calculated by Scaffold.

The hydroxylamine-solubilized only AM was submitted to the Mass Spectrometry Facility at Joint Technology Platform of Center for Molecular and Cellular Bioengineering in Dresden. Collected protein material was separated by SDS gel electrophoresis and proteins were digested in gel either with trypsin (Promega, Mannheim, Germany) according (Shevchenko *et al.*, 2006). Digestion was performed in 10 mM ammonium bicarbonate buffer and enzyme concentration was 10 ng/µL. Extracted peptides were dried in vacuum and stored until analysis at -20°C. For analysis by HPLC-MS/MS protein digests were recovered in 3 µl 30% formic acid containing retention time peptide standard (25 fmol/µL, Thermo Pierce) and diluted by addition of 20 µL H2O. For analysis by HPLC-MS/MS 5 µl were injected into Ultimate3000 nanoLC system (Dionex, Amsterdam, The Netherlands) system equipped with a 75 µm x 2 cm trap column and 75 µm x 15 cm separation column (Acclaim C18 PepMap, Thermo Fisher, Bremen, Germany). Samples were separated in a linear gradient of solvent A (0.1% formic acid) and solvent B (60% acetonitrile, 0.1% formic acid) from 0% B to 55% B in 90 min. The flow rates for sample loading on the trap column was 3 µL/min while the flow rate for separation and analysis was 200 nl/min. The mass spectrometer (LTQ Orbitrap XL ETD, Thermo Fisher, Bremen, Germany) was operated in data dependent analysis (DDA) mode with selection of the 8 most intense ion signals of charge states +2,+3 and +4. MS1 spectra were acquired in the Orbitrap at a nominal resolution of 60000 at 400 Th and MS/MS in the ion trap using CID and a normalized collision energy (NCE) of 35. Acquired spectra were searched against 3'-frame translated *C. australis* transcriptome by Mascot software (Matrix Science, London, UK, v.2.6.0). The hits were evaluated by Scaffold program v.4.8.2 (Proteome Software Inc 2017; Craig & Beavis 2003; Nesvizhskii & Keller 2003) and accepted if two of more peptides were matched under 95% and 99% probability thresholds for peptides and proteins respectively. False Discovery Rate (FDR) was below 1.5% as calculated by Scaffold.

***C. australis* genomic DNA isolation**

HMW genomic DNA (gDNA) isolation, $4x10^8$ cells were resuspended in CTAB extraction solution (AppliChem) containing 2% β-mercaptoethanol (Merck) and incubated for 1 hour at

65°C. The cell debris was removed by centrifugation (3,200 g, 30 min) and the resulting supernatant was treated with RNase A (50 µg/mL, Fermentas) for 1 hour at 37°C and then extracted six times with phenol:chloroform:isoamyl alcohol (25:24:1) and twice with chloroform:isoamyl alcohol (24:1). To precipitate the DNA, 2.5 volumes of ice-cold ethanol (100%, Merck) and 0.1 volumes 1 M NaCl (Sigma Aldrich) were added and the solution was incubated overnight on ice. The precipitated gDNA was collected by centrifugation (5,000 xg, 10 min, 4°C) and washed twice with 70% ethanol. After drying, the gDNA was dissolved in TE buffer (AppliChem).

##### *C. australis* PacBio Genome Assembly

De novo genome assembly was performed with DAmar (https://github.com/MartinPippel/DAmar). This assembler is based on an improved MARVEL assembler (https://github.com/schloi/MARVEL, commit ID: 5e17326) [1] and the integration of parts from the DAZZLER, DALIGNER, DAMASKER, DASCRUBBER, DAZZ_DB, (https://github.com/thegenemyers; [2]) and the DACCORD (version: 0.0.635, [3]) code base. To assemble the genome, we performed the following steps: setup, PacBio read patching, assembly and error-polishing.

###### a) Setup Phase

PacBio reads were filtered by choosing only the longest read of each zero-mode waveguide (ZMW) and requiring subsequently a minimum read length of 4 kb. The resulting 333 thousand reads (69X coverage) were stored in a DAZZLER database. As the chloroplast and mitochondria reads were strongly overrepresented in the database both organelles were first fully assembled with the DAmar organelle pipeline (see 1e). Afterwards any PacBio read that aligns with the assemblies were filtered out. The reduced dazzler database consists of 263 thousand reads (54X coverage).

###### b) Read Patching

The patch phase detects and corrects read artefacts including missed adapters, polymerase strand jumps, chimeric reads, and long low-quality read segments that are the primary impediments to long contiguous assemblies. To this end, we first computed local alignments of all raw reads. Because local alignment computation is by far the most time- and storage-consuming part of the pipeline, we reduced runtime and storage by masking repeats in the reads as follows. First, low-complexity intervals, such as microsatellites or homopolymers, were masked with DBdust (https://github.com/thegenemyers/DAZZ_DB). Second, tandem repeats were masked by using datander and TANmask (https://github.com/thegenemyers/DAMASKER). Third, we used a read alignment step to detect repeats. To this end, we first split all reads into groups representing 1× read coverage. For each group, we then aligned all reads against all others in the same group with daligner (https://github.com/thegenemyers/DALIGNER) and masked all local regions in each read where ≥50 other reads aligned. The repeat masks were subsequently used to prevent k-mer seeding in repetitive regions when computing all local alignments between all reads. Then we applied LAfix to detect and correct read artefacts.

**c) *De novo* assembly**

In the assembly phase, we first calculated all overlaps between patched reads using the same alignment strategy of the patch phase. The subsequent steps of (i) computing a quality track for all reads, (ii) computing a detailed repeat mask, (iii) filtering overlap piles, (iv) computing the overlap graph, (v) touring the overlap graph to obtain primary contigs, and (vi) base error correction to create consensus sequences for the primary contigs follow the steps of the original MARVEL assembly pipeline [1], [4].

**d) Error Polishing**

The assembly and the organelle contigs were further polished by using the raw PacBio reads and applying two rounds of gcpp (https://github.com/PacificBiosciences/gcpp) polishing.

To further correct base errors and reduce remaining length errors in homopolymer regions we used merfin (https://github.com/arangrhie/merfin; [5]) based on an Illumina read set. Therefore, we ran gcpp a third time to create a variant (vcf) file. Only remaining homozygous variants (QUAL>1 && (GT="AA" || GT="Aa")), were used for the merfin polishing.

**e) Mitochondrion and Chloroplast assemblies**

The mitochondrion and chloroplast genomes were assembled with the DAmar organelle assembly pipeline based on the closely related references from *Durinskia baltica* (NCBI accession: NC_014287, ENA accession JN378735). The final circular mitochondrial genome has length of 66,255 bp and the chloroplast genome has a length of 133,148 bp.

The final *Craspedostauros australis* genome assembly was checked for contaminations with blobtoolkit (version 1.1) and an in-house pipeline which screens several blast databases. The base accuracy was estimated with merqury (version 1.3) [6] which reported a QV score of 38.7 and a k-mer completeness of 99.2. BUSCO scores were created based on the stramenopiles_odb10 lineage data set (version 5.2.2) [7].

#### Assembly sequence analysis of region CaTrailin_4 and Ca5609

##### a) CaTrailin_4

The genomic sequence of protein CaTrailin4 and the raw PacBio reads were aligned with minimap2 (version: 2.24-r1122) to the genome assembly. Tandem repeats were identified with the program Tandem Repeat Finder (TRF, version 4.09) and converted into a bed file format. CaTrailin4 unambiguously mapped to contig 00004_0 (uCraAus1_00004_0_1:344.489-373.438 (-) = 28.950bp). Alignments and repeat annotations were loaded into the Integrative Genomics Viewer (IGV, version 2.12.3) for a further manual inspection.

The mapping location of CaTrailin4 has a complex tandem repeat structure (Figure: CA195_Alignment – TandemRepeatTrack). When looking at the coverage profile (Figure: CA195_Alignment - PacBioCoverage), the alignments show a slight decrease in read coverage - 38.6X over the full contig versus 27.0X over the CaTrailin_4 region. But this is not unexpected for such a complicated repeat region. This is also reflected by a dot plot of the CaTrailin_4 self-alignment (Figure Dotplot).

There is no position in CaTrailin4 that is not supported by PacBio reads, in that case all alignments would break. Furthermore, a single PacBio read (highlighted in blue) fully spans the CaTrailin4 region (Figure: CaTrailin4_Alignment – PacBioAlignments). The long PacBio reads (>31Kb) highlighted in red and green, that could be anchored at both non-repetitive sites adjacent to the CaTrailin4 region, also support the different tandem repeat modules i.e. they span multiple different tandem repeat units (363Kb – 372Kb).

### b) Ca5609

The gene Ca5609 is split on two different contigs as not a single PacBio read fully spans the location. All the reads that map to contig uCraAus1_00018_0_1 that reach a bit further into contig uCraAus1_00034_0_1 are fully trapped in the repetitive part of the sequence and cannot be anchored outside the repetitive region.

#### Confirmation of gene models

*C. australis* cDNA and gDNA were prepared as described previously (Poulsen, 2023). Two nested RACE PCRs were performed as described previously (Poulsen & Kroger, 2004) using cDNA attached to the Oligo $(dT)_{25}$ Dynabeads and primer combinations listed in Table S1 using DreamTaq (Thermo Fisher Scientific). The resulting PCR products were cloned into pJet1.2 (Thermo Fisher Scientific) and transformed into the DH5α *E. coli* strain and subsequently sequenced by Eurofins Genomics (Ebersberg, Germany). To confirm the complete gene model and intron-exon boundaries a PCR was performed over the entire predicted coding region using cDNA and gDNA as a template. The PCR was performed using Q5 DNA polymerase (NEB) according to the manufacturer's instructions and using the GC enhancer buffer. The resulting PCR products were directly sequenced by Microsynth Seqlab (Göttingen, Germany).

**Table S1.** Primers used for determining the full-length gene models

| Gene | Primer Name | Sequence |
|---|---|---|
| **Ca807** | | |
| 3'RACE | Ca807_1st PCR | GGACGCCAACGGCCTCCCCTC |
| | Ca807_2nd PCR | GAGGACCTCAGCAAGCAGAAC |
| Full-length gene model | Ca807_FULL_1f | ACGAACCCCCACTTCCACTTC |
| | Ca807_FULL_2r | CTCCTCTGGGGAGCATGCTGG |
| **Ca1637** | | |
| 3'RACE | Ca1637_1st PCR | ATGCAGGTCGGCAACACCACC |
| | Ca1637_2nd PCR | CGGAGGCCACCTTCACCTAC |
| Full-length gene model | Ca1637_FULL_1f | ATTCGGCCAACCGCAACCCAC |
| | Ca1637_FULL_2r | CAGCCGTAGACACTGCACTTG |
| **Ca5255** | | |
| 3'RACE | Ca5255_1st PCR | GCTCTCACAAAGCAAGCGC |
| | Ca5255_2nd PCR | CGAACATGCCATGGACGGC |
| Full-length gene model | Ca5255_FULL_1f | CAAAGCAAGCGCAAATCAAACAC |
| | Ca5255_FULL_2r | GCATTCTACGTACTCTTCATGG |
| **Ca949** | | |
| Full-length gene model | Ca949_FULL_1f | GATCAAAGCCACGCACACACC |
| | Ca949_FULL_2r | ACGCAATAGTACTACGATATC |
| **Ca11384** | | |
| 3'RACE | Ca11384_3RACE_1f | GCACGCAAGCTCAGCGACAAG |
| | Ca11384_3RACE_2f | ATGGTGAACGTGTGCCATTAC |
| Full-length gene model | Ca11384_FULL_4r | GAATGGGTTCGGAAACAATGC |
| | Ca11384_FULL_3f | GAGAGAATAAGTAGGCACACAC |
| **Ca1945** | | |
| 3'RACE | Ca1945_1st PCR | GACTATTGTCGATCTGTGGTTC |
| | Ca1945_2nd PCR | CCCACCGTCCAGTACACATTC |

**Construction of genes encoding GFP-tagged CaTrailin2 and CaTrailin3**

The CaTrailin2 gene was PCR amplified from genomic DNA using the oligonucleotides 5'-GATC TAC GTA ATG ATC ATG AAG ATG CTG TCG-3' and 5'-AGGT TCT AGA TCC CTC GAC AGG TTC CGC ATC-3' that introduced a SnaBI (bold) and XbaI (italics) restriction sites. The resulting 2603-bp PCR product was digested with SnaBI and XbaI, ligated into the SnaBI and XbaI sites of pCafcp_GFP (Poulsen, 2023) and transformed into Dh5α competent *E. coli*. The sequence of the resulting plasmid pCafcp_CaTrailin_2_GFP was confirmed by DNA sequencing (Eurofins Genomics).

The CaTrailin3 gene was PCR amplified from genomic DNA using the oligonucleotides 5'-GATC TAC GTA ATG TTC AAG CAC ATC ACC GTC-3' and 5'-GAAT TCT AGA TGT GGA GGT TTG ACC CGG CGG-3' that introduced a SnaBI (bold) and XbaI (italics) restriction sites. The resulting 4,421-bp PCR product was cloned into pJet1.2 using SURE2 competent *E. coli* cells (Agilent) and sequenced. The resulting plasmid, pJet1.2/CaTrailin_3, was digested with with SnaBI and XbaI and introduced into the SnaBI and XbaI sites of pCafcp_GFP (Poulsen, 2023), generating the final plasmid pCafcp_CaTrailin_3_GFP.

**Bioinformatics analysis of AM proteins.**

Protein domains were identified using ScanProsite (de Castro *et al.*, 2006), Simple Modular Architecture Research Tool (Letunic & Bork, 2018), Pfam (Finn *et al.*, 2016), and InterPro (Finn *et al.*, 2017). Signal peptides were identified by the SignalP-5.0 program (Armenteros *et al.*, 2019). Intrinsically disordered regions were identified using IUPred3 (Erdos *et al.*, 2021). Homologs with other eukaryotic species were identified by performing a search against the MMETSP dataset (Keeling *et al.*, 2014), PLAZA Diatoms 1.0 (Osuna-Cruz *et al.*, 2020) and Phycocosm (Grigoriev *et al.*, 2021). The minimal e-value for homologs was set to $10^{-20}$. BLAST hits to *Tiarina fusa* (ciliophora) or the dinoflagellate *Karenia brevis* (myzozoa) are possibly due to (cross-)contamination of the transcriptomes and have therefore been excluded from further analysis (Marron *et al.*, 2016; Van Vlierberghe *et al.*, 2021).

**Phylogenetic analysis of diatom CAA domains**

A seed alignment was created using the amino acid sequences of the four *Craspedostauros australis* and three *Amphora coffeaeformis* proteins that contain a CAA-domain (see Table S2). After alignment with MAFFT v7.453 (Katoh & Standley, 2013), manual trimming was performed with MEGA X (Kumar *et al.*, 2018) to retain only the CAA domain, and a HMM profile was created with hmmbuild in HMMER v3.1b2. Next, hmmsearch was used for an initial homolog search against the full PLAZA Diatoms protein database, which contains 666,550 protein sequences of 10 diatom species and 16 other eukaryotic species (Osuna-Cruz *et al.*, 2020). An updated HMM profile including the six diatom hits with an E-value below 1e-20 was created by aligning the new protein sequences with the original seven CAA-domains with MAFFT, followed by manual trimming. Two extensive reference datasets were selected for further homology searches. Firstly, the NCBI non-redundant (nr) dataset v2022-02-05 was retrieved from ftp.ncbi.nlm.nih.gov/blast/db/FASTA/nr.gz. Secondly, 224 "clean" transcriptome samples were retrieved from the decontaminated Marine Microbial Eukaryote Transcriptome Sequencing Project (MMETSP) (Keeling *et al.*, 2014; Van Vlierberghe *et al.*, 2021). To translate the open-reading frames (ORF) of these nucleotide sequence datasets to amino acid sequences, Transdecoder v5.0.2 (available at github.com/TransDecoder) was used. To allow detection of the correct ORF based on homology, all possible translated ORFs were queried against the merged SwissProt v2022-02-09 and PLAZA Diatoms 1.0 proteome with Diamond v2.0.14 (Buchfink *et al.*, 2015) in --ultra-sensitive mode. Next, the hmmsearch function from HMMER v3.1b2 was used to search homologs, and only hits with an E-value below 1e-20 were retained. To reduce the number of bacterial sequences CD-HIT v 4.8.1 (Fu *et al.*, 2012) was used to select representative sequences, specifying a global sequence identity cutoff (-c) of 0.7. The NCBI taxonomy (downloaded on 2022-02-11) of the final hits was retrieved using Taxonkit v0.10.1 (Shen & Ren, 2021). Homologous CAA domains from all sources (proteomics, NCBI nr, PLAZA Diatoms and MMETSP) were combined and aligned using MAFFT v7.453, followed by automatic gap trimming with trimal v 1.4.1 (gap threshold 0.8). A maximum-likelihood phylogenetic tree was inferred using IQ-tree v 2.1.3 (Nguyen *et al.*, 2015) with automatic model selection from the following set of substitution models: JTT,LG,WAG,Blosum62,VT and Dayhoff, using empirical amino acid frequencies (-mfreq = F) and the FreeRate rate heterogeneity model (-mrate R) for model selection. Finally, phylogenetic trees were plotted with ggtree and ggtreeExtra in R (Yu *et al.*, 2017; Xu *et al.*, 2021). To construct the species tree in Fig. 2B, a maximum-likelihood 11-gene phylogeny consisting of 1151 taxa was retrieved from (Nakov *et al.*, 2018). The topology of the selected species was visualized with ggtree after pruning the tree using the drop.tip command from Ape v5.6-1 for R (Paradis & Schliep, 2019).

**Structural analysis of the Choice-of-Anchor A (CAA) domain**

The amino acid sequence of CaTrailin4 was submitted to Alphafold (version v2.0.1 with the full_dbs option) to predict its 3D structure (Jumper *et al.*, 2021). Reported quality scores are encoded in the predicted structures b-factors. To identify structural homologues of the CaTrailin_4 CAA domain we carried out a dedicated computational screen, as there are currently no tools to compare a predicted structure to all known structures. Firstly, the Uniprot database was downloaded on 22.11.2021 and 170,041 PDB IDs indexed in Uniprot were extracted. If a Uniprot entry had multiple PDB IDs associated with it, then a representative was selected. Criteria for selection were number of residues covered, resolution, and then method. After selecting representatives, 54,943 PDB IDs were remaining. The PDB IDs were then compared to the predicted structure of the CaTrailin_4 CAA domain using structural alignment (CE align) in Pymol. RMSD and percentage of aligned residues were recorded for each comparison. Overall, there were 223 structures with RMSD less than 10Å (see supplementary Table S8). Next, we sorted these structures by percentage-aligned residues. The top seven PDB IDs were 4NU2, 6QVI, 3UYU, 5B5H, 3WP9, 3VN3, 6A8K, 7BWX. All but 6QVI are ice-binding proteins. We aligned CaTrailin_4 CAA domain with 4NU2 using Pymol's CE align. We exported the sequence representation of the structural alignment and manually added secondary structure. To assess ice-binding, we used the tool AFPredictor (Doxey *et al.*, 2006).

**Immunolabelling of *C.australis* whole cells and biofilms**

Immunolabelling of *C.australis* cells was performed according to previously established method (Aumeier, 2015). If not stated otherwise, all procedures were performed at RT under constant rocking. Following cell counting, the cells were transferred to a 1.5 mL Eppendorf tube (1.8x$10^6$ cells/tube) and fixed with 1% glutaraldehyde + 1.5% paraformaldehyde in Actin Stabilizing Buffer (ASB; 50 mM Pipes, 10 mM EGTA, 5 mM $MgCl_2$, 50 mM KCl, 1% DMSO, pH 7.1) for 15 min. The cells were then rinsed twice with ASB (5 min each), treated with fresh sodium borohydride (2x 15 min) and washed again with ASB (3x 5 min). The non-specific antibody interactions were blocked by incubation with 0.1% Fish gelatin (Sigma Aldrich) and 1% BSA in PBS for 1 h. Following blocking, the cells were incubated with the primary polyclonal antibodies (5 µg/mL; GenScript) diluted in 1% BSA in PBS overnight at 4°C under constant rocking. The cells were then washed with PBS (2x 5 min), incubated for 1 h with the secondary antibody (Goat-anti-rabbit conjugated with AlexaFluor488, 1:3000, Life Technologies) diluted in 1% BSA in PBS and washed again with PBS (2x 5 min). Finally, the cells were resuspended in 200 µl of PBS and 50 µL ibidi mounting medium (Ibidi), mixed with 250 µl of 3% agarose (Biozym) and distributed on a glass bottom dish (µ-Dish 35 mm, ibidi).

To visualize biofilm formation, *C. australis* cells were incubated in a glass bottom chamber slide ($2.5x10^3$ cells/well, µ-slide 8 well chamber, ibidi) with a 12 h light/12 h dark cycle at 18°C for 1, 4, 8 or 12 days. All steps were performed at RT. The culture medium was aspirated and the cells incubated in 2% BSA in PBS (BS) for 1 hour. The samples were then incubated for 2 hours with primary antibodies or pre-immune serum (5 $\mu g \cdot mL^{-1}$) diluted in BS + 0.05% Tween 20. The samples were washed (1x 2 sec, 1x 5 min) with PBS + 0.05% Tween 20 (PBST), and then incubated in secondary antibody (Goat-anti-rabbit conjugated with AlexaFluor488, 1:3000 diluted in PBST; Life Technologies) for 1 hour in the dark. After washing twice with PBST and overlaying with ibidi mounting medium (Ibidi), confocal microscopy was performed using a Zeiss LSM780 inverted microscope equipped with a Zeiss Plan Apochromat 63x (1.4) Oil DIC M27 objective or C-Apochromat 40x/1.2 W Corr M27 objective. Two channels were used to separately monitor chloroplast fluorescence (emission at 654-693 nm) and AlexaFluor488 or Atto488 channel (emission at 480-515 nm). Images were analysed using the ZEN 3.1 software (Zeiss).

**Conjugation of fluorescent dyes to lectin**

Both lectins were labelled with fluorescent dyes via N-hydroxysuccinimide (NHS)-activated crosslinker. *Aleuria aurantia* lectin (AAL, Vector Laboratories) was dissolved in sodium bicarbonate buffer (0.2 M, pH 8.3) into final concentration 1 mg/ml and 7 nmol of AAL was used for conjugation to either Atto 488 NHS ester (Sigma Aldrich) (Thermofisher Scientific). Three times molar access of the dye was added into the solution and the sample was incubated for 3 hours at RT in the dark with gentle rotation. Then 100 µl of Tris-HCl was added (1 M, pH 8.5) to block the excess activated NHS groups and the unconjugated dye was removed by separation of the sample on the PD MiniTrap G-10 desalting column (GE Healthcare). Gravity protocol was used for removal of the free dye and washing steps were performed according to manufacturer's protocol. Briefly, PD MiniTrap G-10 column was prepared and equilibrated by washing with PBS (repeated 3 times) and then 0.5 ml of the sample was loaded onto the column. PBS was added to fill the column to 0.7 ml and the flow through was discarded. Finally, the sample was eluted with 0.2 ml PBS. The elution was repeated 3 times and eluates were kept separate. Eluates were tested on fluorescence microscope and the most effective one (eluate No.2) was aliquoted and frozen in -20°C.

**Lectin labelling**

The procedure for lectin labelling was similar to the immunolabelling. However, instead of antibodies the samples were incubated for 1 h with lectin from *Aleuria aurantia* (AAL: Vector Laboratories) conjugated with Atto488 (4 $\mu g \cdot mL^{-1}$). Confocal microscopy images were taken using a Zeiss LSM780 inverted microscope equipped with a Zeiss Plan Apochromat 63x (1.4) Oil DIC M27 objective or C-Apochromat 40x/1.2 W Corr M27 objective. Two channels were used to separately monitor chloroplast together with AlexaFluor647 fluorescence (emission 654-693 nm) and Atto488 channel (emission at 480-515 nm). Images were analyzed using the ZEN 3.1 software (Zeiss). The control experiments were performed at the same conditions without the labelling with lectins.

**Dot immunoblot analysis of the hydroxylamine solubilized adhesive material**

As attempts to transfer the HMW solubilized adhesive material for a western blot failed we have performed a dot immunoblot of the adhesive material separated by gel filtration chromatography. For the dot immunoblot assay, 4 µL of each pooled fraction was spotted onto a nitrocellulose membrane and allowed to air dry. The membrane was blocked for 90 min in Rotiblock (Carl Roth, Germany) and then incubated with the primary antibody (5 µg/mL) diluted in Rotiblock overnight at 4°C. The membrane was then washed three times for 10 min in PBS containing 0.05% Tween (PBST), and then incubated with the secondary antibody (anti-rabbit IgG-HRP, 1:10,000, Sigma Aldrich) diluted in Rotiblock for 1 h at RT. Finally, the membrane was washed three times for 10 min with PBST, incubated with Pierce SuperSignal West Pico chemiluminescent reagent (Thermo Scientific, Germany) for 5 min and then the signal was detected on a Chemidoc Imaging System (Bio-Rad, Germany).

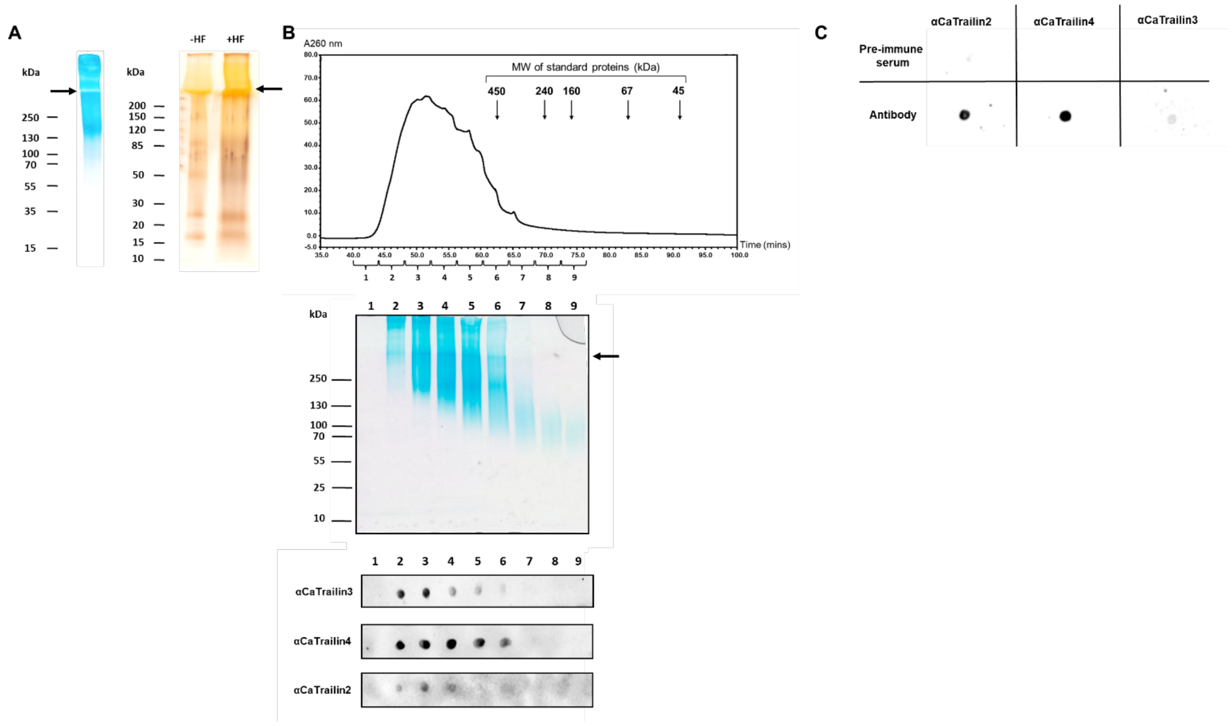

**Figure S1.** Biochemical characterization of the hydroxylamine solubilized adhesive trails from *C. australis*. (**A**) SDS-PAGE of adhesive material stained with Stains All (left) and Silver Stain (right). The interface between the stacking gel and resolving gel is highlighted with an arrow to indicate that much of the adhesive material is very high molecular weight and does not migrate into the resolving gel. **(B)** Gel filtration chromatography separation of hydroxylamine solubilized *C. australis* AM (upper panel). The arrows refer to the molecular weight of standard proteins (in kDa) using Ferritin (450 kDa), Catalase (240 kDa), Aldolase (160 kDa), Albumin bovine (6kDa), Albumin egg (45 kDa), Chymotrypsinogen A (17.8 kDa). The pooled fractions were separated on 16% Schägger gel with subsequent Stains All staining (middle panel). Each fractioned pool was spotted onto nitrocellulose membrane, air dried and then probed with the antibodies αCaTrailin3, αCaTrailin4 and αCaTrailin2. **(C)** Control dot blot probing the crude, adhesive material (9 nmol carbohydrates) using both the pre-immune serum and the polyclonal antibodies against CaTrailin2, CaTrailin3 and CaTrailin4.

**Table S3.** Nucleotide and protein sequences of putative adhesive proteins. The signal peptide is highlighted blue and peptide sequences used for antibody production are highlighted yellow and the Choice-of-Anchor A (CAA) domain is highlighted in purple.

| **Gene ID** | |
|---|---|
| **CaTrailin3** (*Ca807*) | |
| **gDNA/cDNA** | ATGTTCAAGCACATCACCGTCGCCGCCCTCGTGGCCTCAGCATCCCTGCTGATCGCCACCCCTGGCGTCATGGCCACGGACG<br>GCTGCGCCGTTGACACCTTTGGCGCCATCCTCGACAGCCTCCCCGACCAGACCGACAACCCGGCCGACATCCGCATCATCAC<br>CAAGACCACAGCCGAAGTCCCCGTCTGCGTCTACGACCCCGTTGTTGCTGGCTCCGTCAGCTTCCTCAGCGCTGCGCCAGCC<br>TCCACCTCGGCGCTCGAGAGCGACACCACCATGTACCTCTTCGCCGAGAAGACGGGACTCGAGTACACCTCCGCCCTAGGAT<br>CCGTGAATCTGGACGTCCTCGTCTCCGAGCTCGTGGGCGGCGGATCCCGCCGAAGCCTCTCTCTCACCGAGTTCTTCGGCTA<br>CACCGGCTCCAACTACTGCGGAACCGGCTGGATGGACACCTTTAACGCCTGCATGACCTCCAAGCCGTGCCCAGGAGGGCAG<br>AACAGCGAATGCGCCGACGGCCAGTCGTGCTTCGGAGGCATCAACTGCTCCGGCAACTCCTACGGCAGCACCGACCTCCCGT<br>ACTCAGGCCCCAAGTTCTGCGGCAGTTCTTGGACCGACGCATCCAACACCTGCGCCTCTGCCGCCCCGTGCCCAGGAGGACA<br>GGACAATGAGTGCCCTGATGGCCTCAAGTGCTTCTCTGACATCAACTGCGCCACCTCCACCACCTACACCCCTCCTGTGGAA<br>TCTTCGAGCTACTCAGGCCCCAACTTCTGCGGCAGCTCCTGGACCGGCGCGAAGAACGGCTGTGCCACTGGCGCCCCATGTC<br>CGGGAGGACAGAACGATGAGTGCCCCGACGGAGAGTCGTGCTTCGGAGGCATTACTTGCCCAACCCCCACCACCTACGGCGC<br>CGGAAGCACTAGCTCCTTCCTCGGTGACAGTGCCGGATCCTCCTCCTCGGCTTCGAGCTACACAGGCCCCAACTTCTGCGGC<br>AGCTCCTGGACCGGCGCGAATGACGGCTGTGCCACTGGTGCCCCGTGCCCAGGAGCACAGGACAGCGAATGTCCCGATGGCC<br>AGTCATGCTTCGGAGGCATCACCTGCCCAACCCCCACCACCTATGGCAGCACTAGCACCAGCGCAGCAACTTCCGGCTACAC<br>AGGCCCCAACTTCTGCGGTATCGACTGGATGAACGCAAAGAACATCTGCGCCGGGGGTGCTCCATGCCCAGGAGCACAGGAC<br>AGCGAATGTCCCGACGGCCTCAAGTGCTTCGGAGGCATCACCTGCGAAACCCCAACCACCTACGGCAGCACTAGCACCAGCA<br>CAGCAACTTCCAGCTACTCAGGCCCCAACTTCTGCGGCAGCTCCTGGACCAGCGCGAACGACGGCTGTGCCACTGGTGCCCC<br>GTGCCCAGGAGCACAGGATAGTGAGTGTCCCGACGGCCAGTCGTGCTTCGGGGGCATCACCTGCCCAACCCCAACCACCTAC<br>GGCGCCGGAAGCACTAGCTCCTTCCTCGATGACACTGCCGGATCCTCCTCCTCGACTTCGAGCTACACAGGTCCCAACTTCT<br>GTGGAACCTCCTGGACGGGCGCGAAGAACGGCTGTGCCACTGGCGCCCCGTGCCCAGGAGCACAAGACAGCGAATGTCCCGA<br>TGGCCAGTCGTGCTTCGGAGGCATCACCTGCCCAACTCCCACCACCTACGGCAGCACCAGCGCAGCAACTTCGGGCTACACA<br>GCCCCCAACTTCTGCGGCAGCTCCTGGACCGACGCGAACAACAACTGTGCCACTGGTGCCCCATGCCCAGGAGCACAGGACA<br>GTGAGTGTCCCGATGGCCAGTCGTGCTTCGGAGGTATCACCTGCCCAACTCCCACCACCTACGGCAGCGGAAGCACCAGCTC<br>CTTCCTCGGTGGCAGTGCTGGATCCTCCTCGGCTTCGGGCTACACAGCCCCCAACTTCTGTGGAAGCTCCTGGACCAACGCG<br>AACGACAACTGTGCCACTGGTGCCCCATGCCCAGGAGCACAGGATAGCGAATGTCCCGACGGCCAGTCATGCTTCGGAGGTA<br>TCACCTGCCCAACTCCCACCACCTACGGCGCCGGAAGCACCAGCTCCTTCCTCGACAACACCACACCATCGTACACCGGCCC<br>CTCGTTCTGTGGTGACAGCTGGTCGCTGGCGAATACCAACTGCGCCACTGGTGCTCCATGCCCAGGAGGAACCGACGGCGAA<br>TGTCCCGGCGGCCAGTCGTGCTTCGGAGGCATCACCTGCCCAACCCCCACCACCTACGGTACCGGCGCTGCCCCAGTTCCAA<br>ACCTGAACTTCTGCGGCCCTACCTACGATGACGCGGAGGCTAACAAGTGCGGCAACGGTGCCCAGGCATGTCCAGGTGGAAC<br>CGCAGACGAGTGCCCAGGCAGCCAAGACTGTTTCAAGGTCTCCGCGTGCCCCGGAGCCGCCTACCTCGCAACTGAGCAAGAA<br>TCCAACGCCTCCGTCAAGGCAGTCGGCTCATCCCCGGACGCCAGCTTCGGAAACCCTTCGTCCCTGTCCGCATCCCTCCCAG<br>ACGGCACCTACAGCTCATACCTTCTCCACGCCGACTCCCCAGCGCAGTCCACCATCACGGCACTCCTGACCGGATCCATCCG<br>CTTCCCAGGTCAAGAGGTGGTCGGAATCATGTTCTCCAGCGGCGCTCTCAGCTCCACCGACTCCATCTTCGGCCTCTCCTCC<br>ACGCAGTACGGCGGATCCAGCGCCAACCGCGGCTGGGAGGTCCAGAGCGAGTTCCAGGACTTCCTCACCGTCTCCGAATCGT<br>GCGAGGTCGACGTGTCCGCCCTCACCTGCCCTGCAACAACCGCCACCGAGTGTGCCATCACCGCCAACCTCGGATCTGAGAT<br>TGGCGCCGTCGGTGCCGCCTACTCCGGCGACGAAGTCCACGACGATCCATTTCCCAACATCAACACCGGCACCCCGCTCAAC<br>GGCCCCAACGGTCGCGACGACACCACCGCGTTCCTCGTGGGAGGCGACTACCTCTCCCAGATCGCAGCCGAAGTCGAGGGTG<br>GCATGGTCATCCTTGGCGACATGGACATCGACGCCACCGGTGCCAACTCCCTCGTCTACGCAGGATACGGTTCCGGAATCAT<br>CCCGCACGACGGAACGGACGTCATCAAGGTCGGTGGCAACATCAACCTCGACCGCGACGTCGCCGTCATGGACTTTGCCAGC<br>GGAGGTGGCATCATCCGTTACGGAGGATCGCTCACTGGAACCGGAACCTTCAAGATGGGCCCTCAGGCCCAGGCCATCCAAG<br>ATCCCAACCTCGACCTGTCCGCCTACGTCAAGGTCCTCGAAGACCTGAAGGCCAAGTCCGAGTACTGGGCGACCTTCACTCC<br>AAACGGTGACTTCGAGGGTTACGGATACGGCAATCAGGCCACATTCCAGGCCGGCGCAGACAATGCCTGCATCCAGGTCTTC<br>ACTATCGAGGCCGCCGAGCTCGACGCCAACTGGGGCATCAACGTGAAGTTCGACGCCAGCCTCGACGACAAGACCATCCTCA<br>TCAACGTCAACGCCCCACCGGGCGGAGTTGTCACCGTGAACAACCTCGGATTCATCGATGGCGCCGGCAATCTCAACTACGC<br>CTTCTCCTCGTCCGTGATCCAGAACATCATCTGGAACTTCCACGACGCCACCGTCCTCAACCTCGGCCTCTGCGATCCAACC<br>TCCAACCACTGCACCAGCGGAGAGTTCATGGGATCCATCATCGCGCCAAAGGCCAACATCGTCAACATGGCTCTCCCAGGAC<br>AGTCCGGACGCTTCGCCACCAGCGGAACAGTCGTGCACAAGTGGGGTGGATCCGAATTCCACAACTACCCATTCAACCCTCC<br>ATGCCCCCTGCCACCTCCAGACAACCTCCCAGTTCCTCCGGAGTGCACCACCCTGCCACCCGGCCAGACCGATGCCCCCACC<br>AGCTCACCAAAGCCACAGGGCTGCCCAACAGGAGTCACCCTGGTTGGACAGGTTGGTTCCTTGGGCTGGGTCTCAGGCGATG<br>GCAGCATCCCGATCGAGATCGTCTCTGCGGATGATACCACCGTCACCTTCACCGTGAGCTCCACCTGGAACACCGCCAACGT<br>TGACAAACTGTACACCCAGCTCGAGGAGGACGCCAACGGCCTCCCCTCCACCGCATGCTACCTCGAGGAGGCAGTCGGCATG<br>TCCAACGGGGAGACTTACACTGCCAAGTGCTCCATGTATTCCAAGACGGCCATTGTCGACATTTACGTCATCGATGAGGACC<br>TCAGCAAGCAGAACACAGGCAAGACAATCAACGTGCCTGATTGCTGCCACAGCGAGGCCGGCACGCCGCAGGCCATGCACTA<br>CGTGGCCATGCTGCAGTGCACACCGCAGCCATGCCCGCCGGGTCAAACCTCCACAAGACGTCTGGAGGGACGCGGAAACCTC<br>CGTGCCAACTAA |
| **Protein** | MFKHITVAALVASASLLIATPGVMATDGCAVDTFGAILDSLPDQTDNPADIRIITKTTAEVPVCVYDPVVAGSVSFLSAAPA<br>STSALESDTTMYLFAEKTGLEYTSALGSVNLDVLVSELVGGGSRRSLSLTEFFGYTGSNYCGTGWMDTFNACMTSKPCPGGQ<br>NSECADGQSCFGGINCSGNSYGSTDLPYSGPKFCGSSWTDASNTCASAAPCPGGQDNECPDGLKCFSDINCATSTTYTPPVE |

| | |
|---|---|
| | SSSYSGPNFCGSSWTGAKNGCATGAPCPGGQNDECPDGESCFGGITCPTPTTYGAGSTSSFLGDSAGSSSSASSYTGPNFCG<br>SSWTGANDGCATGAPCPGAQDSECPDGQSCFGGITCPTPTTYGSTSTSAATSGYTGPNFCGIDWMNAKNICAGGAPCPGAQD<br>SECPDGLKCFGGITCETPTTYGSTSTSTATSSYSGPNFCGSSWTSANDGCATGAPCPGAQDSECPDGQSCFGGITCPTPTTY<br>GAGSTSSFLDDTAGSSSSTSSYTGPNFCGTSWTGAKNGCATGAPCPGAQDSECPDGQSCFGGITCPTPTTYGSTSAATSGYT<br>APNFCGSSWTDANNNCATGAPCPGAQDSECPDGQSCFGGITCPTPTTYGSGSTSSFLGGSAGSSSASGYTAPNFCGSSWTNA<br>NDNCATGAPCPGAQDSECPDGQSCFGGITCPTPTTYGAGSTSSFLDNTTPSYTGPSFCGDSWSLANTNCATGAPCPGGTDGE<br>CPGGQSCFGGITCPTPTTYGTGAAPVPNLNFCGPTYDDAEANKCGNGAQACPGGTADECPGSQDCFKVSACPGAAYLATEQE<br>SNASVKAVGSSPDASFGNPSSLSASLPDGTYSSYLLHADSPAQSTITALLTGSIRFPGQEVVGIMFSSGALSSTDSIFGLSS<br>TQYGGSSANRGWEVQSEFQDFLTVSESCEVDVSALTCPATTATECAITANLGSEIGAVGAAYSGDEVHDDPFPNINTGTPLN<br>GPNGRDDTTAF**LVGGDYLSQIAAEVEGGMVILGDMDIDATGANSLVYAGYGSGIIPHDGTDVIKVGGNINLDRDVAVMDFAS**<br>**GGGIIRYGGSLTGTGTFKMGPQAQAIQDPNLDLSAYVKVLEDLKAKSEYWATFTPNGDFEGYGYGNQATFQAGADNACIQVF**<br>**TIEAAELDANWGINVKFDASLDDKTILINVNAPPGGVVTVNNLGFIDGAGNLNYAFSSSVIQNIIWNFHDATVLNLGLCDPT**<br>**SNHCTSGEFMGSIIAPKANIVNMALPGQSGRFATSGTVVHKWGGSEFHNYPFNP**PCPLPPPDNLPVPPECTTLPPGQTDAPT<br>SSPKPQGCPTGVTLVGQVGSLGWVSGDGSIPIEIVSADDTTVTFTVSSTWNTANVDKLYTQLEEDANGLPSTACYLEEAVGM<br>SNGETYTAKCSMYSKTAIVDIYVIDEDLSKQNTGKTINVPDCCHSEAGTPQAMHYVAMLQCTPQPCPPGQTSTRRLEGRGNL<br>RAN* |
| **CaTrailin2 *(Ca1637)*** | |
| **gDNA/cDNA** | ATGATCATGAAGATGCTGTCGCCGATGCTGCTGACGGTGGCGGCGCTCAGCGTGGGCACCGCCTCCGCGAACACCTGCATCA<br>ACGCTGGAAGCCTCTGCACGGAACCCGACACCCCGGACAACTGCTGCGAGGGCATGTACTGCAAGCAGTACTCGGTGTACTG<br>GAGCTCGTGCCTCCAAGGAACCGCGCCTCCGCCGCCTCTGGTCACCAACACGCTCTCCCTGCGCATGACCGTGCAGGAGGAC<br>AACCTGGACCAGATCCGCATCATCACGGAGTCCAACCTCGAGGTGCCGCAGTGCCTGTACGAGCCGGGCGTCGACGGCGAGA<br>TCACCCTGATCCCGGGACCATGGTCGTCGCAGATGAACAAGCTCGAGAGCAACAGCGGCCTCTTCATGTGGCAGGAGAAGTC<br>CGGCGTCGTGCTGCCCGGCGACCTCAAGGTGGACGTGCTCAAGACCTACGACCACACCAACGACTGCGAGGTCTACGAAGCC<br>CAGGACGTCGACTACACCGTCTCCCAGTGGTGCGGAACCTCGTGGGCCGAGGCCTCCGACAAGTGCAACGCCTCCTGCATGC<br>AGACCGGCTACGACCAGGAGTGCATCGACCAGCTGGGACCCGGCTCCAAGTGCTTCGGACAGATCAGCAACTGCGGCCGCTT<br>CAACCGCAAGCTGCAGGTCCCCCCACCGGACTTCTCCGACACCCTCGTGGCCGACGGAAAGCGCAAGTTCAACAGCTACATC<br>ATCCACGGCGACACCCTCACGGGCATCTCCGGCACCTACCAGGGATTCATCCGCTTCTTCGGCCAGACCATCGTGGCCGTGC<br>AGTACACCAAGGACGGACTCGACATCACCGATGGCCTGTTCGGCATGTCTGACACCCATCTACCCCACCAAGTCCCGGCGCC<br>GTGGGATCCACCGGCCGCGGCTTCGAGCTCGGCGGGCAACGGCAACGGAGACTTCTTCACGATCTGCCGCAACAAGTTCATC<br>ACCCTCCAGGGCCAGACCTGCCCTGAGACTGCGGCCAGCGCCTGCACCCAGACCGTCAACCTCGGCTACAACGTCGGACCGT<br>CCTCCGGCAACAACGCCTTCGGGGAGGACTACCCCATCACCGACCCCCTGGCTTCCATGTCCAGCCCCTTCGGACGCGACGA<br>CTCCATCGCCTTCCTCACCCTGCAGAGCTACCACTCCTTCCGGGCAGCCGAGGTGGAAGGAAAGATGGTCATCGGCAGAGAC<br>CTAATCATCAAGCCGGACGGTGCCAACTCCCTCGTGCAGGCCGGACACGGGTCCGGAATCATCCCCAACGTGGGAGAGGACA<br>TCATGATCGTCGGTGGCAACATCAAGGTGGAGAGAGACGTCTCCGTCATGGAGACCGGAACCCCGGGAGGCAACATCATCTA<br>CGCCGGAACCTACGAGGACTTCGGCACCGACGGCCTCAGACTCGGGTTCGGTGGACAGCGCATCAACAAGCCAAACCTAGAC<br>ATGGCACCGTACATCGAGATCATGGAGAACGTCCTCCTCAAGTCCGAGTACTGGGCCAGCCTCTCGCCAAATGGGGTCTTCC<br>TCCCGTACACCGAGGGACCAAACGACAACACCATCCTGTTCAAGGCTGGCCAGGACAACGCCTGCGTCCAGATCTTCCACGT<br>CGACCAGAAGGACTTGGGCAACCTCGCGTACGGTGTGAACGTCCAGTTCGATGCCTCCCTCGCCGGCAAGACCATCCTCATC<br>AACGTCGCGGCCGACCCAACCACCGGCCTTGCCAAGATCAGCCACCTGTCCAACTTCATCGACCCATTCGGCAACGGCGGAT<br>TCGACTTCACGTCGAAGACCACCGCCAGCATCATGTGGAACTTCTACGACGCAACTCACGTCGAGCTGGGTAACGACGAGAT<br>TTCCAACGGCCAGTTCCGCGGAACCGTCATCGTCCCCAAGGGAAGCCTCCGCATGACCATGCCTGGACAGTCCGGACGCATG<br>CTCGTCGCCGCGGACATCATGCACGACTTCGTCGGCAGCGAGTTCCACAACTACAAGTTCGACCCAGTCTGCGAGCTCCCAG<br>ACCCACCGACATTCCCAATTCCACAGGAGTGCCAGACGCCCGCACCATATGCCATGGCACCAGACACCGAGACCGGGGCACC<br>CACCATGGCGCCAACTCCGTACCAGTGCCCAGAGGACATCGAGCTGCTGATGCAGGTCGGCAACACCACCTGGACCAGCGCT<br>GACCTCCCCATCATCATCATCGAACAGAACACCACCAGCGTGACGTTCCGCATCAAGCAGGAGTGGGCCGAGGAGCTGTCCT<br>TCATGTACATCCTCTTCGATCAGGAGATCACCAGCGATGAAATCTGCTACACCCACGCGGACATCCCGCCGGAGGCCACCTT<br>CACCTACACCGCCATGTGCACGCACGAGAGCAAGGTCGCCATGGTGGAGATCTGGACCGCCGACGACAGCTTCCTCGTGGGG<br>ACCGACAACGCCGTCGTGCCAGAGTGCTGCCATTCCCCAGATGATGACACCTTCCCCAAGGTTCAGTACTGCTTCGAACTGC<br>ACTGCGTGACGCAGTGTCCCGATGCGGAACCTGTCGAGGGACGCAGACTTCGCGGGTTCAATTAA |
| **Protein** | MIMKMLSPMLLTVAALSVGTASANTCINAGSLCTEPDTPDNCCEGMYCKQYSVYWSSCLQGTAPPPPLVTNTLSLRMTVQED<br>NLDQIRIITESNLEVPQCLYEPGVDGEITLIPGPWSSQMNKLESNSGLFMWQEKSGVVLPGDLKVDVLKTYDHTNDCEVYEA<br>QDVDYTVSQWCGTSWAEASDKCNASCMQTGYDQECIDQLGPGSKCFGQISNCGRFNRKLQVPPPDFSDTLVADGKRKFNSYI<br>IHGDTLTGISGTYQGFIRFFGQTIVAVQYTKDGLDITDGLFGMSDTHLPHQVPAPWDPPAAASSSAGNGNGDFFTICRNKFI<br>TLQGQTCPETAASACTQTVNLGYNVGPSSGNNAFGEDYPI**TDPLASMSSPFGRDDSIAFLTLQSYHSFRAAEVEGKMVIGRD**<br>**LIIKPDGANSLVQAGHGSGIIPNVGEDIMIVGGNIKVERDVSVMETGTPGGNIIYAGTYEDFGTDGLRLGFGGQRINKPNLD**<br>**MAPYIEIMENVLLKSEYWASLSPNGVFLPYTEGPNDNTILFKAGQDNACVQIFHVDQKDLGNLAYGVNVQFDASLAGKTILI**<br>**NVAADPTTGLAKISHLSNFIDPFGNGGFDFTSKTTASIMWNFYDATHVELGNDEISNGQFRGTVIVPKGSLRMTMPGQSGRM**<br>**LVAADIMHDFVGSEFHNYKFD**PVCELPDPPTFPIPQECQTPAPYAMAPDTETGAPTMAPTPYQCPEDIELLMQVGNTTWTSA<br>DLPIIIIEQNTTSVTFRIKQEWAEELSFMYILFDQEITSDEICYTHADIPPEATFTYTAMCTHESKVAMVEIWTADDSFLVG<br>TDNAVVPECCHSPDDDTFPKVQYCFELHCVTQCPDAEPVEGRRLRGFN* |
| **CaTrailin1 *(Ca949)*** | |
| **gDNA/cDNA** | ATGAAGTTCCAAGGTATCGCAGCCATGTTCGGTGCCATCGTGGCCACCTTCGCGAGCACGGCCACAGCACAGGATGGCTCCA<br>CCTGCGAGGCCAACCTCGGAGACGCCATCGCCCCTCCAGATGGCGTGACCTACCCACCCCTCACCAACGGGATCACTGGCGC |

| | |
|---|---|
| | CAACGGCTTCGACAACAACCTTGCCTTCTTCGTCGGAGGTAACTTCATCTCCCGCATTGGATCCGAAGTCGAGGGCGCCATG<br>GTCATCTTGGGTGACATGGTGGTCGAGCCCAGCGGCCTCAACTCCCTCGTCCAGGTCGGACAGGGATCTGGCGTGTTCCCAA<br>ACGGAGGTGACATCCTTCGCATCGGTGGCTCAGTCACCCTCGAGCGCTTCGTCTCCATCGTCGCCTTCCCAGGACCTAACGG<br>AGCCGGAACCGTCACCCACAAGGGACCGGTCGTCGGCAGCGGAAGCTGGCAAATCGGCGCGGGAGGCACCATCGTGCATGAC<br>CCCAACCTCGACCTCACCGCATACAGCAACCTCATGTCCGAGCTCGACGTCAAGTCCCAGTACTACGGCAGCCTCCCAGGCA<br>ACGGAGTGTTCGTCCGCGAAGCTGGTGGATACTACAGACTCTCTGCCGGAGACGACGAATGCCTCCAGGTCTTCAACTTCCC<br>CGCGGATTTCCTCAACGGCGAAGCCTTCGGCTTCGTCATAGACCTCAACGCCAACCTTGCCGGCAAGACCATGATCTTCAAC<br>GTCGCCGCCTTCGACCCTACACCCCAGGAGCCACGCCGCGTCGCCCTCGTGGACAACATCGCCGGATTCTTGGACACCAACG<br>GCAATTTCGGCATCCAGCTCGACCCAGCCACCGGAGCCTCCATCTTGTGGAACTTCCCAGACGCTGAATACGTCGAGTTCGG<br>TAACAACGCCATTGGCAACGGACAGATCGCGGGAACCCTCCTTGCTCCAAGATCTGACCTTCGCTACTCCTTCCCTGGACAC<br>CAGGGACGCGTGATCGTCGGTGGTGACCTCATCCAGGAAAGACAGGGTACCGAGTTCCACAACTACCCATTCGACCCAGTTG<br>ACACCTGCCCACTCCCACCTCTTCCGCAATGCGACTGCCCAATCGAGAACAACGTCAAGTTCATCAAGCAGCAGGGACAGAC<br>CACGTGGACCGGCGAGCTCCCATTCGAGATCATCGAGCAGACCGGCAAGTCCGTCACCTTCAAGGTCAAGCACACCTGGGCC<br>ACACCAAGCATCGACTCGTACTACGTCTACCACGAGAACGGGCTCGACGCCGATGTCTGCGCCCTCAACGAGGACGTCCTTA<br>CCACCGATGAGTTCACGTACACCGCCTCGTGCATGAGCTGCAAGAACACCACTGTCGTTGATCTGTTCATCGCAGATGATGA<br>CTTCGACGCCGTCGCTGACGATGCCACCATCCCGCATTGCTGTCACCCAGAGCCAGAAGACATTGCACGCCCAACCGTGCAG<br>TACACGTTCATCCTGCACTGCGACGACCAGTGCGTGCCCGACAGCGACACGCCAGCAGAGGCGGTGCCTTCTTCATTCCGCA<br>AGCTTCGAGGGGCCTCCGCATAG |
| **Protein** | MKFQGIAAMFGAIVATFASTATAQDGSTCEANLGDAIAPPDGVTYPPLTNGITGANGFDN**NLAFFVGGNFISRIGSEVEGAM**<br>**VILGDMVVEPSGLNSLVQVGQGSGVFPNGGDILRIGGSVTLERFVSIVAFPGPNGAGTVTHKGPVVGSGSWQIGAGGTIVHD**<br>**PNLDLTAYSNLMSELDVKSQYYGSLPGNGVFVREAGGYYRLSAGDDECLQVFNFPADFLNGEAFGFVIDLNANLAGKTMIFN**<br>**VAAFDPTPQEPRRVALVDNIAGFLDTNGNFGIQLDPATGASILWNFPDAEYVEFGNNAIGNGQIAGTLLAPRSDLRYSFPGH**<br>**QGRVIVGGDLIQERQGTEFHNYPFD**PVDTCPLPPLPQCDCPIENNVKFIKQQGQTTWTGELPFEIIEQTGKSVTFKVKHTWA<br>TPSIDSYYVYHENGLDADVCALNEDVLTTDEFTYTASCMSCKNTTVVDLFIADDDFDAVADDATIPHCCHPEPEDIARPTVQ<br>YTFILHCDDQCVPDSDTPAEAVPSSFRKLRGASA* |
| **CaFAP1 (*Ca1769*)** | |
| **gDNA/cDNA** | ATGTTCAAGTTCAAGGCAGCCCTTTTTGGCGCGCTCCTCCTCGCCCATGCGGGTTCGACGGAGGCCGTGCTGCGTGGCAGCA<br>GAGCCGTCGGCCAGACCGAAGAGCTCCATGTCATGATGCAGGCCACCCGCACCCTGAAGGCTGGTAACGGCGACGGCGACAA<br>GCAAGACAAGGAGGGCGGCAAGAAGAGCAAAGACACGGAGGCGCCAACCAGCAACCCGACTGGCACCCCTGGCATGCCAACG<br>GCGGCACCGACCAGCGCCCCAACTTCGGCACCGACGTCGGCTCCGACCAGTGCCCCAACCGCCACGCCTGGCATGCCAACGG<br>CGGCACCCACGTCGGCTCCAACTTCCGCCCCAACTAGCGCCCCGACTTCGGCCCCAACTAGCGCCCCAACCTCGGCTCCGAC<br>ATCTGCGCCCACCAGCGCCCCAACCTCGGCTCCAACCTCCGCCCCAACCTCGGCTCCGACATCTGCACCAACCAGCGCACCA<br>ACCAGTGCCCCGACGTCGGCTCCAACCGCCAACCCAACTGCCACCCCAGGCATGCCAACACCGGCGCCAACCGCGGCCCCAA<br>CCGGCCCAACCGCGGCCCCCACATCTGCCCCAACCAGCGCCCCCACCTCGGCCCCAACGTCGGCACCAACCTCCGCACCAAC<br>CTCAGCTCCGACTTCGGCACCCACATCCGCACCAACCAGCGCCCCAACTTCCGGCCCGACTTCGGCCCCAACTTCCGCTCCC<br>ACGTCGGCTCCAACCTCCGTGCCTACCGCGGCCCCAACATCCGCACCAACCAGCGCCCCGACCGCGGCTCCAACATCTGCCC<br>CCACCGTCTTCGAAGCGTCTATCTCCGTCGCATGCAACCCAGATGTTATTCCAAGCAGCCCTTCGCTCCCACCTGAACTGTT<br>CAACACGAAGACTATCAGCTGCAACAAGGACAAGGATACGTGCCGTCAGGCCATCCGCGCAGTCCGCGACAACTCTGACCTC<br>AAAACGTTCTTCGGCTGCTACGGAAATCTGGGTGAGAATGAAGCACGCATCGGATTCAACACGTACGCATCCAATCCAGGTC<br>TGGCCCAGCAGCTCGAAATGGTCGACTCGGCTGGAACTGAGTTTGTCACTGCACTGCCAAAAGACATGAAGACGGCCTGCTC<br>CAACGCGAACGGCAGCCCCTGCTGCCAGCGCACGGACGGCTGTTCCTCATCCTCGAATTTCGGATTCATGACCGTGGCACTC<br>AACGCTTGCACCGGAATGGGTTCATGCAACAACGTGGGTCTGAATGCCCTCATCCTCGCCACTTCATGCACCGGGGACTATG<br>CGTGCGAAAACCTCAACCAAAACGACGACCCCAACACAGTCCAACGCGGAGTCACCTTCGTCGGTGAATCAGCCTGCACTCC<br>CGCAGAGTCATGCAAAAACGTCGGACAGTTCTCGTACGGTGACGTGACCATCCAGAACGGATCATGCAACGGTTCCAACTCT<br>TGCACAAACGCCGGCCGCCGTGCCATGCAGACGACAGCAGGACGAGTCGTGTCCGACATCAGCATCGGCGCGCAAGCCTGCT<br>CCCAAACCAATTCGTGCTTGGACGCGCACTCCTCCACTGACGCGGGAGCGAATGGAGGATCCAAGAACAGAGGAAGCCTCAG<br>CATTGAGGACGGAGCCTGCGATGGCACCAGTGCCTGCGAAGGCTTGGCCAACAGTGATTTCATGGAAACCGATGTCGTTGTT<br>GAGGAAGGCGACTGCAAAGGATTTGAGACCTGCATGGGATGCGCCCTGGAGAACTGCCCCTTCAATCAGGTTTACGTAATCA<br>TCAACGGACCTTTCCTCTGCGGCCCGGACAGCATGGGAAACCGAAACGGCGAAATCGACGAGGACTTCTGCGTCGCCAAAGC<br>CGACCGTCCAACCGAGTGCCCCACCTTCGCCCCCACCCCGCTCACGAGCGATATCCCATCCGTTCTGCCATCAGAAATGCCG<br>TCCATGGGACCAACCATGTCACCGTCAGCAGGCCCAACCGACGGGCCAACACGCCTCCCCTCCATCAGTCCATCCAACGTCC<br>CAAGCATCCTCCCATCCGTTGGACCTACCAGCACTCCAACGGCAGCCCCAACAGTCAAGAAGACCGCACTGCAGACCCTGTG<br>CCTCAGCGAGCCAGGAAACGTCAACTCTCCGTTTGTTATGAACAGGAGGCGTCTTCAGGAGACTGTGTCATACGGAGAAGAG<br>GGAAGAGAGCTCCAAGTTATCGTGCCGGTCACTGATGCTCCAACAGGGTCTGCCGCTCCAACGGAATCTGTGGCACCCTCGG<br>ATGTGCCCTCGGATGTGCCCTCAGATCTGCCCTCAGATCTGCCCTCGGATATTCCCTCAGACCTTCCGTCCGCACAGCCCTC<br>GGATGTGCCCTCAGACCTTCCGTCCGCACAGCCCTCGGATGTGCCCTCTATCGTACCAACCGACACAAGCGCCCCTACTAAA<br>TCATTCGCGCCGACAGCAACACCAACAATCGAAACAAACACTGGCGATGCTGTGATGGTGTTCTGCGATGATGACGTGGGTG<br>CGTGCACCGACTTCGTGAACCGAATCGAGGACCTCCCCGACAACAAGGACAGGCTCGTGAAGAGCATCTTCGGATGCGACAC<br>CGACACCATGTCCCTCACGACCGCTGAGGAAGCCCAGGCGCGCTTCGATCGCATCAAGAATGATCCCGACGCCCGCGCTTTG<br>ATCGAAAACGCCCTGATGGGAGACGGCGTCCTCCTCACTGGCAGCGGTGCTGATCTATTCACTGCATGCAGTGGGAAGGACT<br>GCTGTCCGACCGCCGCTGATTGCACGCCATCAGGCACCGCCCCCAAGGTGATCACCATTACGCCAGACTCCTGCGCAGGCAC<br>GGATTCTTGCGCAGGAATTAAGGAGTCGTCCATCATCGTTGGAAGCTGCCTTTCCGACAACTCTTGCGAGAATCTTGGAGCG<br>ATCACGAACGAACCTTCGAACGACGCCGTCACCGGATACACGATTGTGGGCCCATCAGCGTGCACGGGCAATTCCAACTGCA<br>ACAACCTTGGAACGGACGTATACGGCAATATTCGCATCGAGAACGCGGCCTGCAACGGCGCGAACAGCTGCACGGGCCTCGG<br>ACAAAAGGCCCCCCAAACCGCTGGAATTGTGACACCTGTTGTCGTTGACATCCTCATCCAGGAGAGGGCCTGCCAGGACCG<br>AGCTCATGCACCAATGTCCACGCCGACCTCACCAAGACCGCAGGAAGCATTGACATTGGAGAGAACGCATGCATGTCCAACA |

| | |
|---|---|
| | ACAGTTGTAATGGCGTTGCCATGCACCCCATGGCCATCGATGACCTCGTCATTAACCCCGGAAGATGTGTGCCAGCCAACAG<br>TTGCCAAAACTGTGTGAAGACTGACATGTGCCTAGGCGCCGGTGGCCAGACCTACACCATCGACGACGCATCCGAGGACTGC<br>ACACCGCCCTATCCGAAGGCTTGCTCCGACTAA |
| **Protein** | MFKFKAALFGALLLAHAGSTEAVLRGSRAVGQTEELHVMMQATRTLKAGNGDGDKQDKEGGKKSKDTEAPTSNPTGTPGMPT<br>AAPTSAPTSAPTSAPTSAPTATPGMPTAAPTSAPTSAPTSAPTSAPTSAPTSAPTSAPTSAPTSAPTSAPTSAPTSAPTSAP<br>TSAPTSAPTANPTATPGMPTPAPTAAPTGPTAAPTSAPTSAPTSAPTSAPTSAPTSAPTSAPTSAPTSAPTSGPTSAPTSAP<br>TSAPTSVPTAAPTSAPTSAPTAAPTSAPTVFEASISVACNPDVIPSSPSLPPELFNTKTISCNKDKDTCRQAIRAVRDNPDL<br>KTFFGCYGNLGENEARIGFNTYASNPGLAQQLEMVDSAGTEFVTALPKDMKTACSNANGSPCCQRTDGCSSSSNFGFMTVAL<br>NACTGMGSCNNVGLNALILATSCTGDYACENLNQNDDPNTVQRGVTFVGESACTPAESCKNVGQFSYGDVTIQNGSCNGSNS<br>CTNAGRRAMQTTAGRVVSDISIGAQACSQTNSCLDAHSSTDAGANGGSKNRGSLSIEDGACDGTSACEGLANSDFMETDVVV<br>EEGDCKGFETCMGCALENCPFNQVYVIINGPFLCGPDSMGNRNGEIDEDFCVAKADRPTECPTFAPTPLTSDIPSVLPSEMP<br>SMGPTMSPSAGPTDGPTRLPSISPSNVPSILPSVGPTSTPTAAPTVKKTALQTLCLSEPGNVNSPFVMNRRRLQETVSYGEE<br>GRELQVIVPVTDAPTGSAAPTESVAPSDVPSDVPSDLPSDLPSDIPSDLPSAQPSDVPSDLPSAQPSDVPSIVPTDTSAPTK<br>SFAPTATPTIETNTGDAVMVFCDDDVGACTDFVNRIEDLPDNKDRLVKSIFGCDTDTMSLTTAEEAQARFDRIKNDPDARAL<br>IENALMGDGVLLTGSGADLFTACSGKDCCPTAADCTPSGTAPKVITITPDSCAGTDSCAGIKESSIIVGSCLSDNSCENLGA<br>ITNEPSNDAVTGYTIVGPSACTGNSNCNNLGTDVYGNIRIENAACNGANSCTGLGQKAPQTAGIVTPVVVDILIQERACQGP<br>SSCTNVHADLTKTAGSIDIGENACMSNNSCNGVAMHPMAIDDLVINPGRCVPANSCQNCVKTDMCLGAGGQTYTIDDASEDC<br>TPPYPKACSD* |
| **CaTrailin4 (Ca1945)** | |
| **gDNA** | ATGAACACCAAGGCCTCCTTCAGAGGCATCATGCCGTCCCTCGGTTTCTTGGGACTCATCGCAGCAGCCATGCTGATGCCAA<br>CCGCGGACGCCACATGTACCCCCCTCAATACATGGTGCCCCAGCGGAAACGACAATTGCTGCGGAACCTCGGAATGCATCTT<br>CAGCAACGGCTGGGATCATTGCCAGACTCCTCCGGGCGAAACCTGTACCCGCCTGTGGGGCTTTCATTGCAACAACCCCGGC<br>ACCAAGAGCACCTGCTGCGACGGTTCGCACTGTGTGTACGGTGGCGTAGGCTGGAACTCCTGTCAGGCAGGTTTCGCGCCGA<br>GCGGCCCATCCGCCCCATCCCCCACCGAAGCCCCCACCTCCGCCCCCACCCAAGCACAGTGCGTCGCCGCCTGGGGTGGGTG<br>CGAACAGCCGGGTACCTACAGCAACTGCTGCGCCGGTTACTACTGTCAGTACGTCAACGAATACTGGAACGGCTGCTATCCA<br>GGCGACCCGCCAGTCCAGCCATCCGGATCCCCGCCGACCGACTCGCCGACCAAGACGCCAACCACCTCGCCCATCGACCCTG<br>GCAGCAGCACCCAGAACTTCTTTGGAAATTCGGGCAGCGGGGGCAGCTTCGCCTGCGCCCTTCCGGACAACCATGGTCTTTT<br>CACGGTCATCACCGAGGGAGATATGCTCATCGGATCCAAGAACATCTACAGTGGAGTGGCCGCAGGAGGTAACATCAACCGT<br>GTGAACAACAGCAACTCCTACAAATTCTGCAGCAACAAGGACGGAGAGGTGTTCGCCGGAGGTAGCATCACCAACAACAACG<br>TCAACTTCGATCCTCGCTGTGGTCCAGCCATCTCAAACGACAGCAATGCCTTGACCCGCGCCAAATGGGATCACTACGTCTG<br>GCTCACTCAAAACGTCAAGGAAGTCTCCACCCGTAAACAGTCCGACGGCAACGTCTTCGTCCGCACCACCGGTGGTGATGTC<br>ACGATGTGCAACTTCCTCGACGGCGACATCCAAGCAGAGGACAACGGAAGGACCATCATCTTCTTCAACACCAACGAGAAAA<br>TCACCGTGAAGAACTGCGGCAGGCAGTGGGGAGCGATCATCATCGCTCCCTTCGCCACAGTCGAAGTCGCTTCTGACATTCA<br>AGCCATGGACGGCCTCGTCGTCGCCAAGGAATTCTCCCAACCGACCGATGTCTCTGGTTCCCAGCTCCACGGTGACGCCTTC<br>GTCGGCTCCCTGACATGCAACCCACCCAGCGCGTCCGTCTGCGAAGACAATATCGGCGATGCCGTGGGCTCCGGATTCCCAT<br>ACGTCGCCGAGTACCCCAACCTCGATGACTGTCTCACCCCCACGGGATACGACAACAAGGTAGCCTTCCTCGTGGGAGGAAC<br>GTACACCTCCATCCAATCTGCTGAGATCGAAGGAAAGATCGTCATTGGTGGTGACTTTATCACCACTGGTGACCAGTCCATC<br>AACTCCGTCGTCTTCGCCGGTGTCGGTTCCTGTATCATCCCCGAGGGTGACCAGGATGTCTTCCTTGTCGGAGGTAACATGG<br>TCATCAACAGATCCATTGTGCTCGTCAACCCAAGCAACGCGTACGTCAGATACGGAGGAACCCTCTCCGGTTCAAACCCCAA<br>CGGTCTCGGCGGAATCGTCCAGCACACTCCCAACCTCGACCTGACCTATTACACTGATGTGATTGAGGCCCTCAAGATCAAG<br>TCTGCGTACTGGGCCACACTCCCAGCGAATGGTGTCTACACCGACCGTACCGGCAGAAACTACGTCGAGTTCATCGCAGGCG<br>CCGCCCCCAATGACGGCTGTCTTCAGGTCTTCGATCTGGATTACGACGACCTCTACTTCGGAGACTGGGGAACTGAAGTCTT<br>CTTCAAGAACCTTTCGGGCAAGACCATGCTCTTCAACGTCGGGTCCGACAGCAACGGCCACGTCAAGATCGCAAATCTCGCA<br>AAGTTCTACGATGGTGCAAAGAGCGACCAGGCCAGCCTCGGCGTCGACAAGGCCTTCAGCAAGGACATGGCCGCCGGCATGA<br>TCTGGAATTTCTACGATGCCACCTCCGTTGAGATCGGAGGAGGCTTGAACGGAAACGGAGAGTTCATCGGTACCATCTTGGC<br>GCCATTCGCAGACCTGGAGTTCACCGCCCCAGGTCAGAGTGGTCGTACCATCGTCGGAGGAGACGTCACTCACGATGCCTAC<br>GGAAGTGAATTCCACAACATCGATTTCGACCCGCCATGCGACCTTCCCATCCCACCGATTCCGGAATGCCCAAACACATCCC<br>CGACATCCGCCCCGACATCCGCGCCAACATCCGGGCCGATGTCCTCCCCGACATCCGCGCCGACGTCCTCCCCGACATCCGC<br>GCCAACATCCGCCCCGACATCCGCGCCGACGTCCGCGCCAACCGCCTCCCCAACATCCGCCCCGACATCCGCGCCAACGTCA<br>GCGCCGACGTCCGCGCCAACCGCCTCCCCAACATCCGCCCCGACATCCAGCATGGCACCAACCAACTGTGTCGATCCAAACG<br>GCGCGAAGTTCACCGGATCCACCGACCCCACTTTCACCGCTGATCCGCTCCCAATCGTCATCACCTTCGAGAGCGATGATGG<br>AAGCACCGTCGAGGTCGATGTTGTGCAGAACATTCTCAATGGGGAAATCAAACATGTCTTCATCAAGTACCGCAACCCAGAG<br>TCTGGAATGGACGACTGCGCCAAGCACGATGACGTCGAATTTGGCAAGAAGATCCCAATCACCGCTGAGTGTTACGGTCAGA<br>TCACCACCATCACGATTTACATTTACACCGGAGAAGACTTCCCCGCGGAGGAATGCGAGGCATGCGAGATCCCAGACGGCGA<br>CCCAGCTGACTTCGAGGCGTACAGCTTCGAGATTCCTTGCGACACTGAGTGCGGAGAACCAACCGCTTCGCCCACCAGCGCA<br>CCAACTTCTGGACCGACGGCCGAGCCTACCTCGGCTCCAACCAGCGGCCCAACATCTGCACCAACCAGCGATCCAACATCTG<br>CACCAACCAGCGACCCAACATCCGCTCCGACCAGCAAGCCAACCTCCGCTCCGACCAGCGACCCAACATCCGCGCCAACGAG<br>CGACCCAACATCCGCACCAACCAGCAAGCCAACTGGAGGACCAACATCCGCACCAACCAGCAAGCCAACTTCTGCTCCAACC<br>ACCAGCGCGGCACCAACCAACTGCGTTGATCCAAATGGAGCCAAGCTCACCAAGTCCACCGACCCTACATACACCGCCGACC<br>CACTTCCAATCGTCATCACCTTCGAGAGTGACGATGGAACCAGTGTCGAGTTCGACTTTGTGCAGAACCTCTCCACCGACAC<br>CATCCCTCATGTCTTCATCAAGTACCGCAACCAGGAGAGCGGAGACGAGGAGTGCGCCCAGTTCGACACTGTCACCCTTGGT<br>ACCACATTGCCAATCGCTGCCGAGTGCTTCGATGGAGTGACTGGAGTCACCATCTACATCTACACAGGAGAAGATTTCCCAA<br>GTGAGGAGTGCGAAGCTTGCAGCATCCCGGATGAAGGAGACTCTGGAAATCCGGACAACTTCGAGGCATACTACTTTGAGGT<br>TCCATGCCAGACCGAGTGTATCGATCCAACTGCGTCCCCAACTTCGGCGCCAACCTCAGGGCCAACTGCGGAACCAACATCT<br>GCACCAACCTCTGCTCCAACAAGCGATCCAACATCTGGACCTACCAGCGGCCCAACTTCGGCACCAACATCTGCACCAACCA<br>GCAATCCAACCGGAGGGCCAACATCCGCACCAACCAGCAAGCCAACTTCTGCTCCAACCACCAGCGCAGCACCAACCAACTG |

| | |
|---|---|
| | CGTTGATCCAAATGGAGCCGTCTTCAAGGAATCCACCGACCCTACTTACACTGCCGACCCACTTCCAATCGTCCTCACTTTC<br>GAGAGTGACGATGGAACCAGTGTTGAGTTCGACTTTGTGCAGAACATCATCAACGGAGTTATCCCACATGTCTTCATCAAGT<br>ACCGCAGTCAGGAGAGTGGAGACGAGGAGTGCGCCCAGTTCGACACTGTCACCCTTGGTACTACTTTGCCATTCGCTGCCGA<br>GTGCTTCGATGGAGTGACTGGAGTCACCATCTACATCTACACAGGAGAAGATTTCCCAAGTGAGGAGTGCGAAGCCTGCAGC<br>ATCCCGGATGAAGGGGACTCTGGAAATTCGGACAACTTCGAGGCATACCACTTTGAGGTTCCATGCCAGACCGAGTGTATCG<br>ATCCAACTGCGTCCCCAACTTCGGCGCCAACCTCAGGGCCAACTGCGGAACCAACATCTGCGCCAACCTCTGCCCCAACAAG<br>CGATCCAACCTCCGGACCTACCAGCGGCCCAACTTCGGCACCAACATCTGCCCCAACCAGCAATCCAACCGGAGGGCCAACA<br>TCCGCACCAACCAGCAAGCCAACGTCGGCACCAACATCCGGACCAACCAGTGCGCCAACTTCTGCTCCAACCACCAGCGCAG<br>CACCAACCAACTGCGTTGATCCAAACGGAGCCGTCTTCAAGGAATCTACCGACCCTACTTACACTGCCGACCCACTTCCAAT<br>CGTCCTCACTTTCGAGAGTGACGATGGAACCAGTGTTGAGTTCGACTTTGTGCAGAACATCATCAACGGAGTTATCCCACAT<br>GTCTTCATCAAGTACCGCAGTCAGGAGAGTGGAGACGAGGAGTGCGCCCAGTTCGACACTGTCACCCTTGGTACTACTTTGC<br>CATTCGCTGCCGAGTGCTTCGAAGGAGTGACTGGAGTCACCATCTACATCTACACAGGAGAAGATTTCCCAAGTGAGGAGTG<br>CGAAGCTTGCAGCATCCCGGATGAAGGAGACTCTGGTAACACGGACAACTTCGAGGCATACTACTTTGAGGTTCCATGCCAG<br>ACCGAGTGTATCGATCCAACTGCGTCCCCAACTTCGGCGCCAACCTCAGGGCCAACTGCGGAACCAACATCTGCACCAACCT<br>CTGCTCCAACCTCTGCTCCAACAAGCGGACCAACCTCTGCGCCAACCTCAGCACCAACAGGTGGCCCAACCGCGGCACCAAC<br>CTCCAAACCAACATCTGGACCTACCAGCGGACCCACATCTGCACCGACATCTGCGCCAACCTCAGGGCCAACTTCGGCACCG<br>ACATCTGCACCTACCGGAGGACCAACATCGGCACCAACTTCTGCACCAACCTCAGGGCCGACATCCGCACCAACCTCAGCAC<br>CAACTTCTGCTCCAACCACCAGCGCGGCACCAACCAACTGCGTTGATCCAAATGGAGCTATCTTCAAGGAATCTACTGACCC<br>TACTTACACTGCCGACCCACTTCCAATCGTCCTCACTTTCGAGAGTGACGATGGAACCAGTGTTGAGTTCGACTTTGTGCAG<br>AACATCATCAACGGAGTCATCCCTCATATCTTCATCAAGTACCGCAACCAAGAAACCGGAACAGAGGATTGTGCCCAGTTCG<br>ACAACGTCGAGACCGGCGCCACTTTGCCATTCGCTGCCGAGTGCTTCTCTGGTGTGACTGGAGTCACCATCTACATCTACAC<br>AGGAGAAGATTTCCCAAGTGAGGAGTGCGAAGCTTGCAGCATCCCAGATGAAGGGGACTCTGGTAACCCGGACAACTTCGAG<br>GCATACTACTTTGAAGTTCCATGCCAGACCGAGTGCATCGATCCAACCGCAGCGCCAACTTCAGCACCAACCAGTGGACCAA<br>CCTCCGGACCGACATCTGGACCTACTTCGGGCCCAACATCTGGACCAACCTCTGGACCTACTTCGGCACCAACTTCTGGACC<br>AACTTCTGGACCAACCAGCGACCCAACATCGGGACCAACATCGGGACCAACTTCTGCACCAACCTCAGGACCAACATCTGGA<br>CCAACCAGCGCGCCAACTTCTGCTCCAACCACCAGCGCAGCACCAACCAACTGCGTTGATCCAAATGGAGCCGTCTTCAAGG<br>AATCCACCGACCCTACTTACACTGCCGACCCACTTCCAATCGTCCTCACTTTCGAGAGTGACGATGGAACCAGTGTTGAGTT<br>CGACTTTGTGCAGAACATCATCAACGGAGTCATCCCTCATATCTTCATCAAGTACCGCAACCAAGAAACCGGAACAGAGGAT<br>TGCGCTCAGTTCGACACTGTCGGCCTTGGTGCTACATACCCACTCACTGCCGAGTGCTTCGAAGGAGTGACCTCAGTCACCA<br>TCTACATCTACACAGGAGAAGATTTCCCAAGTGAGGAGTGCGAAGCTTGCAGCATCCCGGATGGCGAAGGAGACTCTGGAAA<br>CCCGGACAACTTCGAGGCATACTACTTCGAGGTTCCATGCCAGACCGAGTGTATCGATCCAACCGCAGCGCCAACTTCAGCA<br>CCAACCAGTGGACCAACCTCCGGACCGACATCTGGACCTACTTCGGGCCCAACCTCTGGACCAACCTCTGGACCTACCAGCG<br>GCCCGACATCGGGACCAACTTCTGGACCAACCAGCGGCCCAACATCGGGACCAACATCGGGACCAACTTCTGCACCAACCTC<br>AGGACCAACATCTGGACCAACCAGCGCGCCAACTTCTGCTCCAACCACCAGCGCAGCACCAACCAACTGCGTTGATCCAAAC<br>GGAGCCGTCTTCAAGGAATCCACTGACCCTACTTACACTGCCGACCCACTTCCAATCGTCCTCACTTTCGAGAGTGACGATG<br>GAACCAGTGTTGAGTTCGACTTTGTGCAGAACATCATCAACGGAGTCATCCCACATGTCTTCATCAAGTACCGCAACCAAGA<br>AACCGGAACAGAGGATTGCGCTCAGTTCGATACTGTCGGCCTTGGTGCTACATACCCACTCACTGCCGAGTGCTTCGAAGGA<br>GTGACCTCAGTCACCATCTACATCTACACAGGAGAAGATTTCCCAAGTGAGGAGTGCGAAGCTTGCAGCATCCCGGATGGCG<br>AAGGAGACTCTGGAAACCCGGACAACTTCGAGGCATACTACTTCGAGGTTCCATGCCAGACCGAGTGTATCGATCCAACCGC<br>AGCGCCAACTTCAGCACCAACCAGTGGACCAACCTCCGGACCGACATCTGGACCTACTTCGGGCCCAACATCTGGACCAACC<br>TCTGGACCTACCAGCGGCCCGACATCGGGACCAACCTCGGGACCAACATCGGGCCCCACTTCTGGACCAACCAGCGGACCAA<br>CCAGCGGACCAACCTCAGGACCAACCTCAGGACCAACATCCGGACCAACCTCGGGACCAACTTCTGCTCCAACCACCAGCGC<br>GGCACCAACCAACTGCGTTGATCCAAACGGAGCCGTCTTCAAGGAATCCACTGACCCTACTTACACTGCCGACCCACTTCCA<br>ATCGTCCTCACTTTCGAGAGTGACGATGGAACCAGTGTTGAGTTCGACTTTGTGCAGAACATCATCAACGGAGTCATCCCAC<br>ATGTCTTCATCAAGTACCGCAACCAAGAAACCGGAACAGAGGATTGCGCTCAGTTCGACACTGTCGGCCTTGGTGCTACATA<br>CCCACTCACTGCCGAGTGCTTCGATGGAGTGACTGGAGTCACCATCTACATCTACACAGGAGAAGATTTCCCAAGTGAGGAG<br>TGCGAAGCTTGCAGCATCCCGGATGGCGAAGGAGACTCTGGAAACCCGGACAACTTCGAGGCATACTACTTCGAGGTTCCAT<br>GCCAGACCGAGTGTATCGATCCAACCGCAGCGCCAACTTCAGCACCAACCAGTGGACCAACCTCCGGACCGACATCTGGACC<br>TACTTCGGGCCCAACATCTGGACCAACCTCTGGACCTACTTCGGCACCAACTTCTGGACCAACTTCTGGACCAACCAGCGAC<br>CCAACATCGGGACCAACATCGGGACCAACTTCTGCACCAACCTCAGGACCAACATCTGGACCAACCAGCGCGCCAACTTCTG<br>CTCCAACCACCAGCGCAGCACCAACCAACTGCGTTGATCCAAATGGAGCCGTCTTCAAGGAATCCACCGACCCTACTTACAC<br>TGCCGACCCACTTCCAATCGTCCTCACTTTCGAGAGTGACGATGGAACCAGTGTTGAGTTCGACTTTGTGCAGAACATCATC<br>AACGGAGTCATCCCACATGTCTTCATCAAGTACCGCAACCAAGAAACCGGAACAGAGGATTGCGCTCAGTTCGACACTGTCG<br>GCCTTGGTGCTACATACCCACTCACTGCCGAGTGCTTCGATGGAGTGACTGGAGTCACCATCTACATCTACACAGGAGAAGA<br>TTTCCCAAGTGAGGAGTGCGAAGCTTGCAGCATCCCGGATGGCGAAGGAGACTCTGGAAACCCGGACAACTTCGAGGCATAC<br>TACTTCGAGGTTCCATGCCAGACCGAGTGTATCGATCCAACCGCAGCGCCAACTTCAGCACCAACCAGTGGACCAACCTCCG<br>GACCGACATCTGGACCTACTTCGGGCCCAACATCTGGACCAACCTCTGGACCTACCAGCGGCCCGACATCGGGACCAACCTC<br>GGGACCAACATCGGGCCCCACTTCTGGACCAACCAGCGGACCAACCAGCGGACCAACCTCAGGACCAACATCCGGACCAACG<br>TCCGGCCCAACATCCGGGCCAACCAGCGGACCAACCAGCGGACCAACTGGAGGACCAACATCTGCACCAACATCCGGCCCTA<br>CCAGCGGACCAACATCAGGCCCCACTTCTGGGCCAACTAGCGGACCGACCAGCGGACCGACATCTGGACCTACCAGCGGCCC<br>AACCAGTGGACCAACATCGGGCCCCACTTCTGGACCAACCAGCGGACCAACCAGTGGACCGACCAGTGGACCGACCTCTGGA<br>CCAACGTCCGGTCCAACGTCTGGACCAACATCTGGACCAACCAGCGGACCAACCGGAGGACCAACATCTGCACCAACATCCG<br>GCCCTACCAGCGGACCAACATCTGGACCAACCTCTGGACCAACCTCTGGACCAACCTCTGGACCAACCTCTGGACCAACCTC<br>TGGACCAACCTCTGGACCAACCTCTGGACCAACCTCTGGACCAACCTCTGGACCAACCTCTGGACCAACCTCTGGACCAACC<br>AGCGGACCAACCAGCGGACCAACCAGCGGACCAACCTCTGGACCAACCAGCGGACCAACATCTGGACCAACATCTGGACCGA<br>CCTCTGGACCAACGTCCGGCCCAACCTCGGGACCAACATCTGGACCAACCAGCGGACCAACTGGAGGACCAACATCTGCGCC<br>AACATCCGGCCCTACCAGCGGACCAACATCCGGTCCAACATCTGGACCAACCTCTGGACCAACCTCTGGCCCAACATCCGGC<br>CCCACTTCTGGACCAACCAGCGGACCCACCAGTGGACCGACTAGCGGCCCAACCTCTGGACCAACATCTGGACCAACATCTG<br>GACCAACCAGCGGCCCAACTGGAGGACCAACATCTGCGCCAACATCCGGCCCTACCAGCGGTCCAACATCGGGCCCAACATC |

| | |
|---|---|
| | TGGACCAACCTCTGGCCCAACATCTGGACCAACCTCTGGCCCAACATCTGGCCCAACATCTGGACCAACCTCTGGCCCAACA<br>TCTGGACCAACCAGCGGACCCACCAGTGGACCGACGAGCGGCCCAACATCTGGACCAACCAGTGGACCGACTTCTGGGCCGA<br>CCAGCGGCCCAACTGGGGGACCAACATCTGCGCCAACATCCGGCCCGACCAGTGGGCCCACCTCTGGACCAACATCCGGACC<br>AACTTCCGGCCCAACCTCTGGGCCAACTAGCGGACCAACCAGCGGCCCAACTTCGGGACCAACAAGCGGACCAACCAGCGGC<br>CCAACTTCGGGACCGACTTCCGGCCCAACCAGCGGCCCAACCAGCACACCTACTTCTGCTCCAACCACCAGCGCCGCGCCGA<br>CCAACTGCGTTGATCCCGATGGTGCCAAGTTGAAGGGATCAACCGACCCAGACTACACTGCTGACCCGAAGCCCATCGTCCT<br>GACATTCGAGAGCGACGACGGCACACTCGTTGAATTTGACTTTGTCCAGAATATCATCAACGGGGTCATTCCACATGTGTTT<br>ATTAAATACCGCAATGCCGAGACAGGAATCAATGAGTGTTCGGAGATCATGAATGCGGGCGAAGACTCCACGTTCTCGTACG<br>CTGCTGAGTGCATCGACCAAACAACCATGGTCACAATCTACATCTACACAGGCGATGACTTCGCTGGCGAAGATTGCGAAGC<br>ATGCAGTCTGCCAACCGACGATGGGGAGACCGGATCCGGCAACCCGGACAACTTTGAGGCGTACTACTTCGAACTGCCGTGC<br>AACACTGAGTGCATTGAGCCAACCGCATCACCCACATCCGCCCCAACCAGTGGACCAACATTCGGCCCAACCTCTGGACCAA<br>CCAGCGGACCAACTTCAGGACCTACTTCCGGCCCAACTAGCGGCCCAACTTCGGGACCAACTAGCGGACCAACATCTGGACC<br>AACATCTGGACCAACATCTGGACCAACCAGCGGACCAACCAGCGGACCAACCAGCGGACCAACCAGCGGACCAACTAGCGGG<br>CCAACCTCTGGACCAACATCCGGACCAACCAGCGGACCAACCTCTGGACCAACCTCTGGACCTACTTCTGGACCAACTTCGG<br>GACCTACTTCTGGACCAACTTCGGGACCAACCTCGGGACCAACTTCGGGACCAACCAGCGGCCCAACCAGCGGACCAACATC<br>TGGACCAACATCTGGACCTACTTCTGGACCAACCTCGGGACCAACCTCTGGACCAACTTCGGGACCAACATCCGGACCAACC<br>AGCGGACCAACCAGCGGACCAACATCTGGACCAACCAGCGGACCAACTTCGGGACCGACGTCTGGACCAACTAGCGGACCAA<br>CCTCTGGACCAACTAGCGGACCAACCTCTGGACCCACATCCGGCCCAACTTCGGGACCAACATCTGGACCAACGTCTGGACC<br>AACTAGCGGCCCAACCAGCGGGCCAACCTCGGGGCCAACTTCCGGACCTACTTCCGGACCAACCAGCGGACCAACTAGCGGC<br>CCAACCTCTGGACCAACATCCGGACCAACCAGCGGACCTACTTCCGGACCAACATCCGGACCAACATCTGGACCAACATCTG<br>GACCAACTAGCGGGCCAACCTCGGGACCAACTTCGGGACCAACCAGCGGCCCAACCAGCGGACCAACATCTGGACCAACATC<br>TGGACCTACTTCTGGACCAACCTCGGGACCAACCTCTGGACCAACTTCGGGACCAACATCCGGACCAACCAGCGGACCAACC<br>AGCGGACCAACATCTGGACCAACCAGCGGACCAACTTCGGGACCGACGTCTGGACCAACTAGCGGACCAACCTCTGGACCAA<br>CTAGCGGACCAACCTCTGGACCCACATCCGGCCCAACTTCGGGACCAACATCTGGACCAACGTCTGGACCAACTAGCGGCCC<br>AACCAGCGGGCCAACCTCGGGGCCAACTTCCGGACCTACTTCCGGACCAACCAGCGGACCAACTAGCGGCCCAACCTCTGGA<br>CCAACATCCGGACCAACCAGCGGACCTACTTCCGGACCAACATCCGGACCAACATCCGGACCAACATCTGGACCAACATCTG<br>GACCAACTAGCGGGCCAACCTCGGGACCAACTTCGGGACCAACCAGCGGCCCAACCAGCGGACCAACATCTGGACCAACATC<br>TGGACCAACATCTGGACCAACATCTGGACCAACATCTGGACCAACATCTGGACCAACATCTGGACCAACATCTGGACCAACC<br>AGCGGACCAACCAGCGGACCAACTAGCGGGCCAACCTCTGGACCAACGTCTGGACCAACATCCGGACCAACCTCTGGACCTA<br>CTTCTGGACCAACTTCGGGACCTACTTCTGGACCAACTTCGGGACCAACCAGCGGCCCAACCAGCGGACCAACATCTGGACC<br>AACCAGCGGACCAACTTCGGGACCGACGTCTGGACCAACTAGCGGACCAACGTCTGGACCAACGTCTGGACCAACCTCTGGA<br>CCCACATCCGGCCCAACTTCGGGACCAACATCTGGACCAACGTCTGGACCAACTAGCGGCCCAACCAGCGGGCCAACCTCGG<br>GGCCAACTTCCGGACCTACTTCCGGACCAACCAGCGGACCAACTAGCGGCCCAACCTCTGGACCAACATCCGGACCAACCAG<br>CGGACCTACTTCCGGACCAACATCCGGACCAACATCTGGACCAACATCTGGACCAACTAGCGGGCCAACCTCGGGACCAACT<br>TCGGGACCAACCAGCGGCCCAACCAGCGGACCAACATCTGGACCAACATCTGGACCAACATCTGGACCAACTTCCGGCCCAA<br>CCTCGGGACCAACCAGCGGACCGACGTCTGGACCAACTTCGGGACCAACATCTGGACCAACCAGCGGACCAACTTCGGGACC<br>AACATCTGGACCAACCAGCGGACCAACCTCTGGACCAACGTCTGGACCAACCTCTGGACCCACATCCGGCCCAACTTCGGGA<br>CCAACATCTGGACCAACGTCTGGACCAACTAGCGGCCCAACCAGCGGGCCAACCTCGGGGCCAACTTCCGGACCTACTTCCG<br>GACCAACCAGCGGACCAACTAGCGGCCCAACCTCTGGACCAACATCCGGACCAACCAGCGGACCTACTTCGGGACCAACATC<br>CGGACCAACCTCTGGACCAACCTCTGGACCAACTTCGGGACCAACATCCGGACCAACCAGCGGGCCAACCAGCGGACCAACA<br>TCTGGACCAACATCCGGACCAACCAGCGGACCTACTTCCGGACCAACATCTGGACCAACATCTGGACCAACTAGCGGGCCAA<br>CCTCGGGACCAACTTCGGGACCAACCAGCGGCCCAACCAGCGGACCAACATCTGGACCAACATCTGGACCAACTTCCGGCCC<br>AACCTCGGGACCAACCAGCGGACCGACGTCTGGACCAACTTCGGGACCAACATCTGGACCAACCAGCGGACCAACTTCGGGA<br>CCAACATCTGGACCAACCAGCGGACCAACCTCTGGACCAACGTCTGGACCAACCTCTGGACCCACATCCGGCCCAACTTCGG<br>GACCAACATCTGGACCAACGTCTGGACCAACTAGCGGCCCAACCAGCGGGCCAACCTCGGGGCCAACTTCCGGACCTACTTC<br>CGGACCAACCAGCGGACCAACTAGCGGCCCAACCTCTGGACCAACATCCGGACCAACCAGCGGACCTACTTCGGGACCAACA<br>TCCGGACCAACCTCTGGACCAACCTCTGGACCAACTTCGGGACCAACATCCGGACCAACCAGCGGGCCAACCAGCGGACCAA<br>CATCTGGACCAACATCCGGACCAACCAGCGGACCAACATCTGGACCAACATCCGGACCAACATCTGGACCAACATCTGGACC<br>AACTAGCGGGCCAACCTCGGGACCAACTTCGGGACCAACCAGCGGCCCAACCAGCGGACCAACATCTGGACCAACATCTGGA<br>CCAACATCCGGCCCAACTTCGGGACCAACTTCGGGACCAACGTCTGGACCAACTAGCGGCCCAACCAGCGGGCCAACCTCTG<br>GACCAACCTCGGGGCCAACTTCCGGACCTACTTCCGGACCAACCAGCGGACCAACTAGCGGCCCAACCTCTGGACCAACTTC<br>GGGACCAACATCCGGACCAACCAGCGGGCCAACCAGCGGACCAACATCTGGACCAACATCCGGACCAACCAGCGGACCTACT<br>TCCGGACCAACATCCGGACCAACATCTGGACCAACATCTGGACCAACTAGCGGCCCAACCTCTGGACCAACATCCGGACCAA<br>CCAGCGGACCTACTTCGGGACCAACATCCGGACCAACCTCTGGACCAACCTCTGGACCAACTTCGGGACCAACATCCGGACC<br>AACCAGCGGGCCAACCAGCGGACCAACATCTGGACCAACCTCGGGACCAACTTCCGGCCCAACCTCGGGACCAACTTCGGGA<br>CCAACATCCGGACCAACCAGCGGACCGACGTCTGGACCAACATCTGGACCAACCTCTGGACCAACCAGCGGACCAACATCCG<br>GCCCAACTTCGGGACCGACATCTGGACCAACCTCTGGACCAACCTCTGGACCAACCTCTGGACCAACCTCGGGACCAACCTC<br>GGGACCAACCTCTGGACCAACTTCGGGACCAACCAGCGGACCAACCTCTGGACCAACATCTGGACCAACCAGCGGACCAACA<br>TCCGGCCCAACTTCGGGACCGACGTCTGGACCAACCTCTGGACCAACATCCGGACCTACTTCGGGACCAACATCTGGGCCAA<br>CATCTGGACCAACGTCTGGACCAACTAGCGGGCCAACCTCTGGACCAACCTCGGGGCCAACCAGCGGACCAACTAGTGGACC<br>AACCAGCGGACCAACCAGCGGACCAACCAGCGGACCAACTAGTGGACCAACGTCTGGACCAACTTCCGGCCCAACCAGCGGA<br>CCAACATCCGGACCAACATCTGGACCAACATCTGGACCAACATCTGGACCAACATCTGGACCAACATCTGGACCAACATCTG<br>GACCAACCAGCGGACCAACGTCTGGACCAACCAGCGGACCAACGTCTGGACCAACCAGCGGACCAACGTCTGGACCAACCAG<br>CGGACCAACGTCTGGACCAACATCCGGACCGACGTCTGGACCCACTAGCGGACCAACATCCGGACCAACATCCGGACCAACA<br>TCTGGACCAACATCTGGACCAACTTCCGGCCCAACCAGCGGACCAACTTCGGGACCAACCTCTGGACCAACCAGCGGCCCAA<br>CTTCGGGACCAACATCCGGACCAACCAGCGGACCAACCTCTGGACCGACGTCTGGACCCACTAGCGGACCAACATCTGGACC<br>CACTAGCGGACCAACATCTGGACCAACCAGCGGACCAACATCCGGACCAACCAGCGGACCAACATCCGGACCAACATCCGGA<br>CCAACATCCGGACCAACATCTGGACCAACATCTGGACCAACTTCCGGCCCAACCAGCGGACCAACCAGCGGACCAACATCTG<br>GACCAACATCTGGACCAACATCTGGACCAACATCCGGCCCAACCAGCGGACCAACCAGCGGACCAACGTCTGGACCAACGTC |

| | |
|---|---|
| | TGGACCAACGTCTGGACCAACATCCGGACCGACGTCTGGACCCACTAGCGGACCAACCTCTGGACCAACATCTGGACCAACC<br>AGCGGCCCAACTTCGGGACCAACATCCGGACCAACCAGCGGACCAACCTCTGGACCAACCTCTGGACCTACTTCCGGCCCAA<br>CATCTGGACCAACCAGCGGACCAACCAGCGGACCAACGTCTGGACCAACTTCGGGACCAACGTCTGGACCAACTTCGGGACC<br>AACGTCTGGACCAACTTCGGGACCAACTTCGGGACCGACTTCTGGACCCACTAGCGGACCAACTAGCGGACCAACCTCGGGG<br>CCAACTTCCGGACCAACATCTGGACCAACCAGCGGACCAACCAGCGGACCAACTTCCGGCCCAACCAGCGGACCAACTAGCG<br>GACCAACTAGCGGCCCAACCTCTGGACCAACATCCGGACCAACCAGCGGACCTACTTCGGGACCAACATCTGGACCAACCAG<br>CGGACCAACCAGTGGGCCAACTTCGGGACCAACATCTGGACCAACATCTGGACCTACTAGCGGACCAACATCTGGACCTACT<br>TCGGGACCTACTTCAGGACCAACTAGCGGCCCAACCTCTGGACCAACCTCTGGACCAACATCTGGACCAACATCTGGACCAA<br>CATCCGGCCCAACCAGCGGACCAACCTCTGGACCAACCTCTGGACCAACGTCTGGACCAACCTCTGGACCAACGTCTGGACC<br>AACATCCGGACCGACGTCTGGACCCACTAGCGGACCAACCTCTGGACCAACATCTGGACCAACCAGCGGCCCAACTTCGGGA<br>CCAACATCCGGACCAACCAGCGGACCAACCAGCGGACCAACCAGCGGACCAACCAGCGGACCAACCAGCGGACCAACCAGCG<br>GACCAACCTCTGGACCTACTTCCGGCCCAACATCTGGACCAACCAGCGGACCAACGTCTGGACCAACTTCGGGACCAACGTC<br>TGGACCAACTTCGGGACCAACGTCTGGACCAACTTCGGGACCAACTTCGGGACCGACTTCTGGACCCACTAGCGGACCAACT<br>AGCGGACCAACCTCTGGACCAACATCTGGACCAACCAGCGGACCAACTTCCGGCCCAACCAGCGGACCAACTAGCGGACCAA<br>CTAGCGGCCCAACCTCTGGACCAACATCCGGACCAACCAGCGGACCTACTTCGGGACCAACATCTGGACCAACCAGCGGACC<br>AACCAGTGGGCCAACTTCGGGACCAACATCTGGACCAACATCCGGACCAACATCTGGACCTACTAGCGGACCAACATCTGGA<br>CCTACTTCGGGACCAACCTCTGGCCCAACCAGTGGACCTACTTCTGGACCAACCTCTGGCCCAACATCTGGGCCAACATCTG<br>GACCAACATCTGGCCCAACCAGCGGGCCAACCAGCGGCCCAACATCTGGACCAACCTCTGGGCCAACCTCTGGACCAACCAG<br>CGGACCAACATCCGGGCCAACTAGCGGGCCAACCTCGGGACCAACATCCGGACCAACCAGCGGACCAACTAGCGGACCAACC<br>AGCGGACCTACATCGGGACCAACATCTGGCCCAACCAGCGGACCTACTTCGGGACCAACCAGCGGACCAACATCCGGACCTA<br>CTTCGGGACCAACATCCGGCCCAACCTCAGGACCAACTTCCGGCCCAACCAGCGGCCCAACGTCTGGACCAACCAGCGGACC<br>AACCTCTGGACCAACTAGCGGACCAACTTCAGGACCAACCTCTGGCCCAACATCCGGACCAACCAGCGGACCAACATCTGGC<br>CCAACCAGCGGACCAACCTCTGGACCAACCTCTGGACCAACCTCTGGACCAACCTCTGGACCAACTAGCGGACCTACTTCGG<br>GACCAACATCTGGACCAACCTCCGGACCAACCTCCGGACCAACCTCCGGACCAACTTCGGGACCAACTAGCGGACCAACTTC<br>GGGACCAACCTCGGGACCAACTAGCGGACCCACATCCGGACCAACTAGCGGACCAACATCCGGCCCAACTCCACAGCCAACA<br>TCCCAGCCAACAGCCAGTGCTGCACCAACCAACTGCGTTGATCCAGATGGCGCCAAGTTCGTCAGGTCCACCGATCCTTCCT<br>ACACTGCAGACCCCAAGCCAATCATCCTCACATTCGAGAGCGACGATGCAACACTCGTTGAGTTTGACTTCATTCAGAACAT<br>CATCAACGGGGTCATCCCGCATGTCTTCATCAAGTACCGCAAGTCCGAGACTGGTATCAACGAATGCGCTGAGTTTGCAAAT<br>GTCGGAGAGAACACTCCGATCTCACTTTCTGCCGAGTGTGTGGACATGTCCACGAACGTGGTGATCTACATCTACACTGGCG<br>ATGACTTTGCCAGTGAGAACTGCGAGGCATGCAGCCTTCCAAGTGATGACGGCGAAGGTGGCTCAGGCAATCCAGACAACTT<br>TGCTGCGTACTACTTCGATGTTCCATGCAACACCGAGTGTATCGAGCCAACGGCTGCCCCAACTTCGGCGCCGACCTCTGGA<br>CCAACATCCGGACCAACAAGCGGCCCAACATCCGGACCAACGTCGGGACCAACCAGTGGACCAACATCCGGACCAACCTCTG<br>GACCAACCTCTGGACCAACTTCGGGACCAACCTCTGGACCAACCTCTGGACCAACTTCAGGACCAACATCTGGACCAACCTC<br>GGGACCAACCAGCGGACCAACTTCGGGACCAACCTCTGGACCAACATCTGGACCAACATCTGGCCCAACTAGCGGACCCACC<br>TCTGGACCAACCAGCGGACCAACCAGCGGACCAACCAGCGGACCAACATCTGGCCCAACCAGCGGACCAACATCTGGCCCAA<br>CTAGCGGACCAACATCTGGACCAACCAGCGGACCAACATCTGGCCCAACTTCCGGCCCAACCAGCGGACCAACCTCTGGACC<br>AACTAGCGGCCCAACATCTGGACCTACTTCGGGACCAACATCTGGACCAACATCTGGGCCAACATCCGGCCCAACCTCGGGA<br>CCAACCAGCGGACCAACATCCGGACCAACTTCAGGACCAACTAGCGGCCCAACCTCGGGACCAACCTCTGGACCAACCAGCG<br>GACCAACCAGCGGCCCAACATCTGGACCAACCAGTGGCCCAACCTCTGGGCCAACGTCTGGGCCAACGTCTGGGCCAACGTC<br>TGGACCAACTAGCGGACCAACCAGTGGCCCAACCTCTGGACCAACATCTGGACCAACCAGCGGACCAACATCCGGACCAACC<br>AGCGGACCTACTTCTGGACCAACCAGCGGACCTACTTCTGGACCAACCAGCGGACCAACATCCGGCCCAACCTCGGGACCAA<br>CCAGCGGACCAACCTCTGGACCAACCAGTGGACCAACCAGTGGACCAACCTCGGGACCTACTTCCGGCCCAACCAGCGGCCC<br>AACCTCGGGACCGACTTCTGGACCAACTAGCGGCCCAACCTCTGGCCCAACTTCGGGACCAACCAGTGGCCCAACCTCTGGA<br>CCAACTTCGGGACCCACATCTGGACCAACCTCTGGACCAACCAGCGGACCAACCAGCGGACCAACCAGCGGACCAACATCTG<br>GACCAACCAGCGGACCAACATCTGGCCCAACTTCCGGCCCAACCAGCGGACCAACCTCTGGACCAACCTCTGGACCAACTAG<br>CGGCCCAACATCTGGACCTACTTCGGGACCAACATCTGGACCAACATCCGGCCCAACCTCGGGACCAACCAGCGGACCAACA<br>TCCGGACCAACTTCAGGACCAACTAGCGGCCCAACCTCGGGACCAACCTCTGGACCAACCAGCGGACCAACCTCTGGACCAA<br>CATCCGGCCCAACATCTGGACCAACCAGTGGCCCAACCTCTGGACCAACTAGCGGACCAACCTCTGGACCAACCAGTGGACC<br>AACATCTGGACCAACTTCCGGCCCAACCTCGGGACCAACCTCTGGACCAACTTCGGGACCAACCAGCGGACCAACATCTGGA<br>CCGACGTCTGGACCCACTAGCGGACCAACTAGCGGACCAACCTCTGGACCAACATCTGGACCAACATCTGGACCAACTAGCG<br>GACCCACCTCTGGACCAACCAGCGGGCCAACGTCTGGACCAACCAGTGGACCTACTTCCGGCCCAACCTCTGGACCAACTAG<br>CGGACCAACATCTGGACCCACCTCTGGACCAACCAGCGGGCCAACGTCTGGACCAACCAGTGGACCTACTTCCGGACCAACA<br>TCCGGTCCAACTAGCGGACCAACATCTGGACCAACCAGCGGACCTACTTCTGGACCAACTAGCGGACCAACATCCGGACCAA<br>CTAGCGGCCCAACTTCGGGACCAACCAGCGGCCCAACATCTGGACCAACCTCGGGACCTACTTCGGGACCAACCTCTGGACC<br>AACTAGCGGACCAACCTCTGGGCCAACGTCTGGACCAACTAGCGGACCAACCAGTGGCCCAACCTCTGGACCAACTAGCGGA<br>CCAACATCTGGACCAACCAGCGGACCAACATCCGGACCAACCAGCGGACCTACTTCGGGACCTACTTCTGGACCAACTAGCG<br>GACCAACATCCGGACCAACATCCGGACCAACTAGCGGCCCAACTTCGGGACCAACCAGCGGACCAACATCTGGACCAACCAG<br>CGGACCAACATCCGGACCAACCAGCGGACCAACATCTGGACCAACCAGCGGACCAACATCTGGACCAACTAGCGGACCTACT<br>TCGGGACCAACCAGCGGACCCACTTCTGGACCAACCTCGGGACCTACTTCGGGACCAACATCTGGACCAACTTCTGGACCAA<br>CATCTGGACCAACTAGCGGCCCAACCTCTGGACCAACTTCAGGACCAACCTCGGGACCAACCTCTGGACCAACCAGCGGACC<br>AACATCCGGCCCAACTAGCGGCCCAACCTCGGGACCAACATCTGGACCAACCAGCGGACCAACATCCGGACCAACTTCAGGA<br>CCAACCTCGGGACCAACATCTGGACCAACCAGCGGACCAACATCCGGACCAACCTCTGGACCAACTAGCGGACCTACTTCGG<br>GACCAACATCCGGCCCAACCTCTGGACCAACTTCAGGACCAACTAGCGGCCCAACCTCGGGACCAACCAGCGGACCAACCAG<br>CGGACCAACCAGCGGACCAACCAGCGGACCAACCAGCGGACCAACCAGCGGACCAACTAGTGGACCAACCAGCGGACCAACC<br>AGCGGACCAACTAGTGGACCAACTTCCGGCCCAACATCGGGACCAACATCGGGACCAACCTCTGGCCCAACATCGGGACCAA<br>CCTCTGGACCAACCAGCGGCCCAACATCTGGACCTACTTCGGGACCAACATCCGGCCCAACTTCAGGACCAACCAGCGGACC<br>AACTAGCGGCCCAACCTCGGGACCAACATCTGGACCAACCAGCGGCCCAACCAGCGGCCCAACTTCGGGACCAACATCTGGA<br>CCTACTTCGGGACCTACTTCTGGACCAACATCGGGACCAACATCGGGACCAACCTCTGGCCCAACCAGCGGGCCAACCTCTG<br>GACCAACTAGCGGCCCAACCTCTGGCCCAACTTCGGGACCAACCAGCGGCCCAACATCTGGACCAACTTCGGGACCAACCAG |

| | |
|---|---|
| | CGGACCAACATCCGGACCAACCTCTGGACCAACTAGCGGACCTACTTCGGGACCAACATCCGGCCCAACCTCGGGACCAACT<br>AGCGGACCAACTAGCGGCCCAACCTCTGGACCAACATCTGGACCAACCAGCGGCCCAACTAGCGGACCAACTTCGGGACCAA<br>CATCTGGACCAACATCCGGACCCACATCTGGACCTACTAGCGGACCAACCTCTGGCCCAACCAGCGGGCCAACCTCTGGACC<br>AACTAGCGGCCCAACCTCTGGACCAACCAGTGGCCCAACCTCTGGACCAACTTCGGGACCAACCAGCGGACCAACATCCGGA<br>CCAACTAGCGGACCAACTTCGGGACCAACATCTGGACCAACATCCGGACCCACATCTGGACCTACTAGCGGACCAACATCGG<br>GACCAACCTCTGGACCAACCAGCGGGCCAACCTCTGGACCAACCTCTGGACCAACCTCTGGACCAACCTCTGGCCCAACTTC<br>GGGACCAACCAGCGGCCCAACATCTGGACCTACTTCGGGACCAACCAGCGGACCAACCTCTGGCCCAACTTCGGGACCAACT<br>TCGGGACCAACCAGCGGCCCAACCAGCGGCCCAACTTCGGGACCAACATCTGGACCAACATCCGGACCCACATCTGGACCTA<br>CTAGCGGACCAACATCGGGACCAACATCGGGACCAACATCGGGACCAACATCGGGACCAACATCGGGACCAACATCGGGACC<br>AACATCGGGACCAACATCGGGACCAACATCTGGCCCAACCAGCGGCCCAACCTCTGGACCAACTTCCGGCCCAACCAGCGGC<br>CCAACCTCTGGACCAACCTCTGGCCCAACTTCGGGACCAACCAGCGGACCAACCAGCGGCCCAACCTCTGGCCCAACTTCGG<br>GACCAACCAGCGGCCCAACATCTGGACCTACTTCGGGACCAACCAGCGGACCAACATCCGGACCTACATCGGGACCAACATC<br>TGGCCCAACCAGCGGACCAACCTCTGGCCCAACCAGCGGCCCAACCTCTGGACCAACTTCCGGCCCAACCAGCGGACCTACA<br>TCGGGACCAACATCTGGCCCAACCAGCGGACCTACTTCGGGACCAACCAGCGGACCAACATCCGGACCTACTTCGGGACCAA<br>CATCCGGCCCAACCTCAGGACCAACTTCCGGCCCAACCAGCGGCCCAACGTCTGGACCAACCAGCGGACCAACCTCTGGACC<br>AACTAGCGGACCAACTTCAGGACCAACCTCTGGCCCAACATCCGGACCAACCAGCGGACCAACATCTGGCCCAACATCTGGC<br>CCAACATCGGGACCAACCTCCGGACCAACCTCCGGACCAACCTCCGGACCAACCTCCGGACCAACCTCCGGACCAACCTCCG<br>GACCAACCTCCGGACCAACATCTGGACCAACATCTGGACCAACCTCTGGACCAACCTCTGGACCAACTAGCGGACCCACATC<br>CGGACCAACTAGCGGACCAACATCCGGCCCAACTCCACAGCCAACATCCCAGCCAACAGCCAGTGCTGCACCAACCAACTGC<br>GTTGATCCAGATGGCGCCAAGTTCGTCAAGTCCACCGATGCTTCCTACACTGCAGACCCCAAGCCAATCATCCTCACATTCG<br>AGAGCGACGATGCAACACTCGTTGAGTTTGACTTCATTCAGAACATCATCAACGGGGTCATCCCGCATGTCTTCATTCAGTA<br>CCGCAAGTCCGAGACTGGTATCAACGAATGCGCTGAGTTTGCAAATGTCGGAGAGAACACTCCAATCTCACTTTCTGCCGAG<br>TGTGTGGACATGTCCACGAACGTGGTGATCTACATCTACACTGGCGATGACTTTGCCAGTGAGAACTGCGAGGCATGCAGCC<br>TTCCAAGTGATGACGGCGAAGGTGGCTCAGGCAATCCAGACAACTTTGCTGCGTACTACTTCGATGTTCCATGCAACACCGA<br>GTGTATCGAGCCAACGGCTGCCCCAACTTCGGCGCCGACCTCTGGACCAACATCCGGACCAACAAGCGGCCCAACATCCGGA<br>CCAACGTCGGGACCAACCAGTGGACCAACATCCGGACCAACCTCTGGACCAACTTCAGGACCAACATCTGGACCAACCTCGG<br>GACCAACCAGCGGACCAACCTCTGGACCAACATCTGGACCAACATCTGGACCAACTAGCGGACCCACCTCTGGACCAACCAG<br>CGGGCCAACGTCTGGACCAACCAGTGGACCTACTTCCGGCCCAACCTCTGGACCAACTAGCGGACCAACATCTGGACCCACC<br>TCTGGACCAACCAGCGGGCCAACGTCTGGACCAACCAGTGGACCTACTTCCGGACCAACATCCGGTCCAACTAGCGGACCAA<br>CATCTGGACCAACCAGCGGACCTACTTCTGGACCAACTAGCGGACCAACATCCGGACCAACTAGCGGCCCAACTTCGGGACC<br>AACCAGCGGCCCAACATCTGGACCAACCTCGGGACCTACTTCGGGACCAACCTCTGGACCAACTAGCGGACCAACCTCTGGG<br>CCAACCTCTGGGCCAACGTCTGGACCAACTAGCGGACCAACCAGTGGCCCAACCTCTGGACCAACTTCCGGCCCAACCAGCG<br>GCCCAACCTCTGGACCAACCTCTGGCCCAACTTCGGGACCAACCAGCGGACCAACATCCGGACCAACCTCTGGACCAACCTC<br>GGGACCAACATCTGGACCAACTTCGGGACCAACCTCTGGACCAACCTCTGGACCAACCTCTGGACCAACCTCTGGACCAACC<br>TCTGGACCAACCTCTGGACCAACCAGCGGACCAACATCTGGACCGACGTCTGGACCCACTAGCGGACCAACCAGCGGACCAA<br>CCTCTGGACCAACATCTGGACCAACATCTGGACCAACTAGCGGACCCACCTCTGGACCAACCAGCGGGCCAACGTCTGGACC<br>AACCAGTGGACCTACTTCCGGCCCAACCTCTGGACCAACTAGCGGACCAACATCTGGACCCACCTCTGGACCAACCAGCGGG<br>CCAACGTCTGGACCAACCAGTGGACCTACTTCCGGACCAACATCCGGTCCAACTAGCGGACCAACATCTGGACCAACCAGCG<br>GACCTACTTCTGGACCAACTAGCGGACCAACATCCGGACCAACTAGCGGCCCAACTTCGGGACCAACCAGCGGCCCAACATC<br>TGGACCAACCTCGGGACCTACTTCGGGACCAACCTCTGGACCAACTAGCGGACCAACCTCTGGGCCAACGTCTGGACCAACT<br>AGCGGACCAACCAGTGGCCCAACCTCTGGACCAACTAGCGGACCAACATCTGGACCAACCAGCGGACCAACATCCGGACCAA<br>CCAGCGGACCTACTTCGGGACCTACTTCTGGACCAACTAGCGGACCAACATCCGGACCAACTAGCGGCCCAACTTCGGGACC<br>AACCAGCGGCCCAACTTCGGGACCAACCAGCGGCCCAACATCTGGACCAACCAGCGGACCAACATCTGGACCAACCAGCGGA<br>CCAACATCTGGACCAACTAGCGGACCAACATCTGGACCAACTAGCGGACCAACATCCGGACCAACCTCTGGACCAACCAGTG<br>GACCAACCTCGGGACCTACTTCCGGCCCAACCATCGGACCAACCTCGGGACCGACTTCTGGACCAACCAGCGGCCCAACCTC<br>TGGCCCAACTTCGGGACCAACCAGCGGCCCAACCTCTGGACCAACTTCGGGACCCACATCTGGACCAACCTCTGGCCCAACC<br>AGTGGACCTACTTCTGGACCAACATCTGGACCAACGTCTGGACCAACGTCTGGACCAACGTCTGGACCAACTAGCGGCCCAA<br>CCAGCGGGCCAACCTCTGGACCAACGTCTGGACCTACTTCCGGCCCAACTAGCGGACCAACCAGCGGACCAACCTCTGGACC<br>AACCTCTGGACCAACATCGGGACCAACCTCTGGCCCAACCTCTGGCCCAACCAGCGGACCAACCTCTGGACCAACCTCTGGA<br>CCAACTAGCGGACCAACTAGCGGACCAACTAGCGGACCAACCTCTGGACCAACCAGCGGCCCAACCTCTGGACCAACTAGCG<br>GACCAACCTCGGGACCAACCTCTGGACCAACCAGCGGACCAACATCTGGACCAACATCCGGCCCAACCAGCGGACCAACCAG<br>CGGACCAACCTCTGGACCAACCAGCGGCCCAACATCTGGACCAACCTCTGGACCAACCTCGGGACCAACATCTGGACCAACT<br>TCGGGACCAACCTCTGGACCAACTAGCGGCCCAACATCTGGACCTACTTCGGGACCAACATCTGGACCAACATCCGGCCCAA<br>CCTCGGGACCAACCAGCGGACCAACATCCGGACCAACTTCAGGACCAACTAGCGGCCCAACCTCGGGACCAACCTCTGGACC<br>AACCTCTGGACCAACCAGTGGACCAACCAGCGGACCAACCTCTGGACCAACTAGCGGACCAACCTCTGGACCAACATCCGGC<br>CCAACATCTGGACCAACCTCTGGACCAACATCCGGACCTACTTCGGGACCAACATCTGGACCAACATCTGGACCAACTAGCG<br>GCCCAACCAGCGGGCCAACCTCTGGACCAACGTCTGGACCTACTTCCGGCCCAACTAGCGGACCAACCAGCGGACCAACCTC<br>TGGACCAACCTCTGGACCAACCTCTGGACCAACATCGGGACCAACCTCTGGCCCAACCTCTGGCCCAACCAGCGGACCAACC<br>TCTGGACCAACTAGCGGACCAACTAGCGGACCAACTAGCGGACCAACTAGCGGACCAACTAGCGGACCAACTAGCGGACCAA<br>CCTCTGGACCAACCAGCGGCCCAACCTCTGGACCAACTAGCGGACCAACATCTGGACCAACTTCCGGCCCAACTTCCGGCCC<br>AACCTCGGGACCAACCTCTGGACCAACCAGCGGACCAACATCTGGACCAACATCCGGCCCAACCAGCGGACCAACCAGCGGA<br>CCAACCTCTGGACCAACCAGCGGCCCAACATCTGGACCAACTAGCGGACCAACCTCTGGACCAACATCCGGACCTACTTCGG<br>GACCAACCTCTGGACCAACGTCTGGACCAACTAGCGGCCCAACCAGCGGGCCAACCTCGGGGCCAACCAGCGGACCAACCAG<br>CGGACCAACCAGCGGACCAACCAGCGGACCAACTAGTGGACCAACCAGCGGACCAACTAGTGGACCAACTTCCGGCCCAACA<br>TCGGGACCAACATCGGGACCAACATCGGGACCAACCTCTGGCCCAACATCGGGACCAACCTCTGGACCAACCAGCGGCCCAA<br>CATCTGGACCTACTTCGGGACCAACATCCGGCCCAACTTCAGGACCAACCAGCGGACCAACATCTGGACCAACCTCGGGACC<br>AACATCTGGACCAACTAGTGGCCCAACCTCGGGACCAACTTCAGGACCAACATCTGGACCAACCTCTGGACCAACTTCGGGA<br>CCAACATCCGGCCCAACATCTGGACCAACTTCAGGACCAACTAGCGGCCCAACCTCTGGACCAACATCCGGCCCAACATCTG<br>GACCAACCAGCGGCCCAACATCTGGACCAACTAGCGGACCTACTTCGGGACCAACCAGCGGACCCACTTCTGGACCAACCTC |

| | |
|---|---|
| | GGGACCAACTTCAGGACCAACATCTGGACCAACATCTGGACCAACTTCGGGACCAACATCCGGACCAACTAGCGGACCAACC<br>TCTGGACCAACTAGCGGACCAACCTCTGGACCAACATCCGGCCCAACATCTGGACCAACCAGCGGCCCAACATCTGGACCAA<br>CTAGCGGACCTACTTCGGGACCAACCAGCGGACCCACTTCTGGACCAACCTCGGGACCAACTTCAGGACCAACATCTGGACC<br>AACCTCTGGACCAACTTCGGGACCAACATCCGGACCAACTAGCGGACCAACCTCTGGACCAACATCCGGCCCAACATCTGGA<br>CCAACCAGCGGCCCAACATCTGGACCAACTAGCGGACCTACTTCGGGACCAACCAGCGGACCCACTTCTGGACCAACCTCGG<br>GACCAACCTCTGGACCAACTAGCGGACCAACCTCTGGACCAACCAGCGGACCCACTTCTGGACCAACTAGCGGACCTACTTC<br>CGGACCAACCTCTGGCCCAACATCTGGACCAACTAGCGGACCAACTAGCGGACCAACTAGCGGACCAACCAGCGGCCCAACA<br>TCTGGACCTACTTCGGGACCAACCAGTGGACCAACATCCGGGCCAACATCTGGACCAACCAGCGGGCCAACATCTGGACCAA<br>CCAGCGGGCCAACATCGGGACCTACTTCCGGCCCAACCTCTGGACCAACATCGGGACCAACCAGTGGCCCAACATCTGGACC<br>AACATCTGGACCAACCAGCGGACCAACATCCGGACCAACTTCCGGCCCAACCTCTGGACCAACTTCGGGACCTACTAGTGGA<br>CCAACCTCTGGACCAACCAGTGGACCAACTAGCGGACCAACCAGCGGACCAACATCTGGACCAACCAGCGGACCAACATCTG<br>GACCAACCAGCGGCCCAACCAGCGGACCAACATCTGGACCAACTTCCGGCCCAACCTCGGGACCAACATCTGGACCAACAAG<br>CGGCCCAACTTCTGGACCAACCAGCGGACCAACATCCGGACCAACCAGCGGACCAACATCTGGACCAACAAGCGGCCCAACT<br>TCTGGACCTACTTCTGGACCAACCTCGGGACCAACTTCAGGCCCAACATCCGGACCAACGTCTGGACCAACTGAAACCGCCT<br>GCATCTACACTGTTCTCGATTTCGCGACCGACGCTGGAGGCAACCCAACAGGCAACGGATATGTGCCCTCGGATGTGTACTC<br>ACAATACGACATTGATGCCATCACAGTAGATGTCACCGGCAGCCCAGGATCCTACGGTGGACCAAACAACATGAGCGGCAGA<br>TACTGGGTCACTGGCAACCCGCTGGTGGGCACCGCCTTGATCATCCAAGGAGCGGAGGGACCCACCGCGGTTCCAAGCGACG<br>ATGGTGGTGATATCGTGTTCACCCTCGACGGAGCTTCAGGAGAAGAGATGCAGCTCACATTCTTCGATCTTGCGGAGGACGT<br>GACAATCACACTCAACACAGAGACTGACCCAGAGGCTGGCGGTACAAGCACGACTGTCCCTGTCGTCGCCGACCCAAGTGGT<br>GGGCTGCAGTCCATCGAGATCGATATCTCGCTCATCAAGCAGGTCATTGTTACATTCACAGGTCGCGGAGCCATCGCTTCCG<br>TTGGCATCTGTCGCGATCCAAACACTCCACCACCAATGGGCAACAACAGCCCGTCGGCAACTCTGGAGCCAACATCCATGCC<br>AACCACAAGCCCAGCACCAACCGAGACACCTGGATGCCCCGAAGACATCGTGTTGGTCGCCAGCGTGGGACACACCCTTCCT<br>CCAAGATTCGTCATCACGATCATCGAGCAAACAACCGACACTGTCACATTCGAAGTCACCCAAAAGTTCAACAAGACCGTCA<br>GTCGCGTCTTCACGCAGTACCACGAGACTCCAGCAGGAGAGACCGAGTGCTTCGAAGACGACAACGTCCCATACGAAGAGAC<br>TCAGCGTCACACCGCGTACTGTATGCACTGCGTACCGCTGACTATTGTCGATCTGTGGTTCTCCGATCCTACCCTCGAGCCA<br>ACCCTGGACAATGCCATCGTTCCTCCGTGCTGCCACCCGCCAACAAACGAGACCTACCCCACCGTCCAGTACACATTCAAGC<br>TGTACTGCGAGACCAAGTGTCCACCTGATGGAGAGCCGAACCAGTTCCAAGAGCGCCGCTTGTTGGAAGCCCCAGAAGAGCC<br>AGTCGAAGTCGAGGCCGAGGCCATCACAGACATCGCCCCAGCTCAAGAGGAGGAGGACGACTCCAAGCATGCGTGCTCCAGC<br>AAGGACCATCCATGCAACGGCGATATGGTCTACGTCTGCCACTACTCCAGCCGCCACGGATACCAGACCTACTGCGTGCCAG<br>AAGCGGATTCTGATATCGTCGAGTTCTATCCAAAGTCCTACTGCGGACGTTGTACCGGAGGACAGGGCAACTACATCAGAGA<br>GTAG |
| **Protein** | MNTKASFRGIMPSLGFLGLIAAAMLMPTADATCTPLNTWCPSGNDNCCGTSECIFSNGWDHCQTPPGETCTRLWGFHCNNPG<br>TKSTCCDGSHCVYGGVGWNSCQAGFAPSGPSAPSPTEAPTSAPTQAQCVAAWGGCEQPGTYSNCCAGYYCQYVNEYWNGCYP<br>GDPPVQPSGSPPTDSPTKTPTTSPIDPGSSTQNFFGNSGSGGSFACALPDNHGLFTVITEGDMLIGSKNIYSGVAAGGNINR<br>VNNSNSYKFCSNKDGEVFAGGSITNNNVNFDPRCGPAISNDSNALTRAKWDHYVWLTQNVKEVSTRKQSDGNVFVRTTGGDV<br>TMCNFLDGDIQAEDNGRTIIFFNTNEKITVKNCGRQWGAIIIAPFATVEVASDIQAMDGLVVAKEFSQPTDVSGSQLHGDAF<br>VGSLTCNPPSASVCEDNIGDAVGSGFPYVAEYPNLDDCLTPTGYDNKVA**FLVGGTYTSIQSAEIEGKIVIGGDFITTGDQSI**<br>**NSVVFAGVGSCIIPEGDQDVFLVGGNMVINRSIVLVNPSNAYVRYGGTLSGSNPNGLGGIVQHTPNLDLTYYTDVIEALKIK**<br>**SAYWATLPANGVYTDRTGRNYVEFIAGAAPNDGCLQVFDLDYDDLYFGDWGTEVFFKNLSGKTMLFNVGSDSNGHVKIANLA**<br>**KFYDGAKSDQASLGVDKAFSKDMAAGMIWNFYDATSVEIGGGLNGNGEFIGTILAPFADLEFTAPGQSGRTIVGGDVTHDAY**<br>**GSEFHNIDFD**PPCDLPIPPIPECPNTSPTSAPTSAPTSGPMSSPTSAPTSSPTSAPTSAPTSAPTSAPTASPTSAPTSAPTS<br>APTSAPTASPTSAPTSSMAPTNCVDPNGAKFTGSTDPTFTADPLPIVITFESDDGSTVEVDVVQNILNGEIKHVFIKYRNPE<br>SGMDDCAKHDDVEFGKKIPITAECYGQITTITIYIYTGEDFPAEECEACEIPDGDPADFEAYSFEIPCDTECGEPTASPTSA<br>PTSGPTAEPTSAPTSGPTSAPTSDPTSAPTSDPTSAPTSKPTSAPTSDPTSAPTSDPTSAPTSKPTGGPTSAPTSKPTSAPT<br>TSAAPTNCVDPNGAKLTKSTDPTYTADPLPIVITFESDDGTSVEFDFVQNLSTDTIPHVFIKYRNQESGDEECAQFDTVTLG<br>TTLPIAAECFDGVTGVTIYIYTGEDFPSEECEACSIPDEGDSGNPDNFEAYYFEVPCQTECIDPTASPTSAPTSGPTAEPTS<br>APTSAPTSDPTSGPTSGPTSAPTSAPTSNPTGGPTSAPTSKPTSAPTTSAAPTNCVDPNGAVFKESTDPTYTADPLPIVLTF<br>ESDDGTSVEFDFVQNIINGVIPHVFIKYRSQESGDEECAQFDTVTLGTTLPFAAECFDGVTGVTIYIYTGEDFPSEECEACS<br>IPDEGDSGNSDNFEAYHFEVPCQTECIDPTASPTSAPTSGPTAEPTSAPTSAPTSDPTSGPTSGPTSAPTSAPTSNPTGGPT<br>SAPTSKPTSAPTSGPTSAPTSAPTTSAAPTNCVDPNGAVFKESTDPTYTADPLPIVLTFESDDGTSVEFDFVQNIINGVIPH<br>VFIKYRSQESGDEECAQFDTVTLGTTLPFAAECFEGVTGVTIYIYTGEDFPSEECEACSIPDEGDSGNTDNFEAYYFEVPCQ<br>TECIDPTASPTSAPTSGPTAEPTSAPTSAPTSAPTSGPTSAPTSAPTGGPTAAPTSKPTSGPTSGPTSAPTSAPTSGPTSAP<br>TSAPTGGPTSAPTSAPTSGPTSAPTSAPTSAPTTSAAPTNCVDPNGAIFKESTDPTYTADPLPIVLTFESDDGTSVEFDFVQ<br>NIINGVIPHIFIKYRNQETGTEDCAQFDNVETGATLPFAAECFSGVTGVTIYIYTGEDFPSEECEACSIPDEGDSGNPDNFE<br>AYYFEVPCQTECIDPTAAPTSAPTSGPTSGPTSGPTSGPTSGPTSGPTSAPTSGPTSGPTSDPTSGPTSGPTSAPTSGPTSG<br>PTSAPTSAPTTSAAPTNCVDPNGAVFKESTDPTYTADPLPIVLTFESDDGTSVEFDFVQNIINGVIPHIFIKYRNQETGTED<br>CAQFDTVGLGATYPLTAECFEGVTSVTIYIYTGEDFPSEECEACSIPDGEGDSGNPDNFEAYYFEVPCQTECIDPTAAPTSA<br>PTSGPTSGPTSGPTSGPTSGPTSGPTSGPTSGPTSGPTSGPTSGPTSAPTSGPTSGPTSAPTSAPTTSAAPTNCVDPN<br>GAVFKESTDPTYTADPLPIVLTFESDDGTSVEFDFVQNIINGVIPHVFIKYRNQETGTEDCAQFDTVGLGATYPLTAECFEG<br>VTSVTIYIYTGEDFPSEECEACSIPDGEGDSGNPDNFEAYYFEVPCQTECIDPTAAPTSAPTSGPTSGPTSGPTSGPTSGPT<br>SGPTSGPTSGPTSGPTSGPTSGPTSGPTSGPTSGPTSGPTSGPTSGPTSAPTTSAAPTNCVDPNGAVFKESTDPTYTADPLP<br>IVLTFESDDGTSVEFDFVQNIINGVIPHVFIKYRNQETGTEDCAQFDTVGLGATYPLTAECFDGVTGVTIYIYTGEDFPSEE<br>CEACSIPDGEGDSGNPDNFEAYYFEVPCQTECIDPTAAPTSAPTSGPTSGPTSGPTSGPTSGPTSGPTSAPTSGPTSGPTSD<br>PTSGPTSGPTSAPTSGPTSGPTSAPTSAPTTSAAPTNCVDPNGAVFKESTDPTYTADPLPIVLTFESDDGTSVEFDFVQNII<br>NGVIPHVFIKYRNQETGTEDCAQFDTVGLGATYPLTAECFDGVTGVTIYIYTGEDFPSEECEACSIPDGEGDSGNPDNFEAY<br>YFEVPCQTECIDPTAAPTSAPTSGPTSGPTSGPTSGPTSGPTSGPTSGPTSGPTSGPTSGPTSGPTSGPTSGPTSGPTSGPT<br>SGPTSGPTSGPTSGPTGGPTSAPTSGPTSGPTSGPTSGPTSGPTSGPTSGPTSGPTSGPTSGPTSGPTSGPTSGPTSGPTSG<br>PTSGPTSGPTSGPTSGPTGGPTSAPTSGPTSGPTSGPTSGPTSGPTSGPTSGPTSGPTSGPTSGPTSGPTSGPTSGPTSGPT<br>SGPTSGPTSGPTSGPTSGPTSGPTSGPTSGPTSGPTSGPTSGPTSGPTGGPTSAPTSGPTSGPTSGPTSGPTSGPTSGPTSG |

| | |
|---|---|
| | PTSGPTSGPTSGPTSGPTSGPTSGPTSGPTSGPTGGPTSAPTSGPTSGPTSGPTSGPTSGPTSGPTSGPTSGPTSGPTSGPT<br>SGPTSGPTSGPTSGPTSGPTSGPTSGPTSGPTGGPTSAPTSGPTSGPTSGPTSGPTSGPTSGPTSGPTSGPTSGPTSGPTSG<br>PTSGPTSGPTSGPTSTPTSAPTTSAAPTNCVDPDGAKLKGSTDPDYTADPKPIVLTFESDDGTLVEFDFVQNIINGVIPHVF<br>IKYRNAETGINECSEIMNAGEDSTFSYAAECIDQTTMVTIYIYTGDDFAGEDCEACSLPTDDGETGSGNPDNFEAYYFELPC<br>NTECIEPTASPTSAPTSGPTFGPTSGPTSGPTSGPTSGPTSGPTSGPTSGPTSGPTSGPTSGPTSGPTSGPTSGPTSGPTSG<br>PTSGPTSGPTSGPTSGPTSGPTSGPTSGPTSGPTSGPTSGPTSGPTSGPTSGPTSGPTSGPTSGPTSGPTSGPTSGPTSGPT<br>SGPTSGPTSGPTSGPTSGPTSGPTSGPTSGPTSGPTSGPTSGPTSGPTSGPTSGPTSGPTSGPTSGPTSGPTSGPTSGPTSG<br>PTSGPTSGPTSGPTSGPTSGPTSGPTSGPTSGPTSGPTSGPTSGPTSGPTSGPTSGPTSGPTSGPTSGPTSGPTSGPTSGPT<br>SGPTSGPTSGPTSGPTSGPTSGPTSGPTSGPTSGPTSGPTSGPTSGPTSGPTSGPTSGPTSGPTSGPTSGPTSGPTSGPTSG<br>PTSGPTSGPTSGPTSGPTSGPTSGPTSGPTSGPTSGPTSGPTSGPTSGPTSGPTSGPTSGPTSGPTSGPTSGPTSGPTSGPT<br>SGPTSGPTSGPTSGPTSGPTSGPTSGPTSGPTSGPTSGPTSGPTSGPTSGPTSGPTSGPTSGPTSGPTSGPTSGPTSGPTSG<br>PTSGPTSGPTSGPTSGPTSGPTSGPTSGPTSGPTSGPTSGPTSGPTSGPTSGPTSGPTSGPTSGPTSGPTSGPTSGPTSGPT<br>SGPTSGPTSGPTSGPTSGPTSGPTSGPTSGPTSGPTSGPTSGPTSGPTSGPTSGPTSGPTSGPTSGPTSGPTSGPTSGPTSG<br>PTSGPTSGPTSGPTSGPTSGPTSGPTSGPTSGPTSGPTSGPTSGPTSGPTSGPTSGPTSGPTSGPTSGPTSGPTSGPTSGPT<br>SGPTSGPTSGPTSGPTSGPTSGPTSGPTSGPTSGPTSGPTSGPTSGPTSGPTSGPTSGPTSGPTSGPTSGPTSGPTSGPTSG<br>PTSGPTSGPTSGPTSGPTSGPTSGPTSGPTSGPTSGPTSGPTSGPTSGPTSGPTSGPTSGPTSGPTSGPTSGPTSGPTSGPT<br>SGPTSGPTSGPTSGPTSGPTSGPTSGPTSGPTSGPTSGPTSGPTSGPTSGPTSGPTSGPTSGPTSGPTSGPTSGPTSGPTSG<br>PTSGPTSGPTSGPTSGPTSGPTSGPTSGPTSGPTSGPTSGPTSGPTSGPTSGPTSGPTSGPTSGPTSGPTSGPTSGPTSGPT<br>SGPTSGPTSGPTSGPTSGPTSGPTSGPTSGPTSGPTSGPTSGPTSGPTSGPTSGPTSGPTSGPTSGPTSGPTSGPTSGPTSG<br>PTSGPTSGPTSGPTSGPTSGPTSGPTSGPTSGPTSGPTSGPTSGPTSGPTSGPTSGPTSGPTSGPTSGPTSGPTSGPTSGPT<br>SGPTSGPTSGPTSGPTSGPTSGPTSGPTSGPTSGPTSGPTSGPTSGPTSGPTSGPTSGPTSGPTSGPTSGPTSGPTSGPTSG<br>PTSGPTSGPTSGPTSGPTSGPTSGPTSGPTSGPTSGPTSGPTSGPTSGPTSGPTSGPTSGPTSGPTSGPTSGPTSGPTSGPT<br>SGPTSGPTSGPTSGPTSGPTSGPTSGPTSGPTSGPTSGPTSGPTSGPTSGPTSGPTSGPTSGPTSGPTSGPTSGPTSGPTSG<br>PTSGPTSGPTSGPTSGPTSGPTSGPTSGPTSGPTSGPTSGPTSGPTSGPTSGPTSGPTSGPTSGPTSGPTSGPTSGPTSGPT<br>SGPTSGPTSGPTSGPTSGPTSGPTSGPTSGPTSGPTSGPTSGPTSGPTSGPTSGPTSGPTSGPTSGPTSGPTSGPTSGPTSG<br>PTSGPTSGPTSGPTSGPTSGPTSGPTSGPTSGPTSGPTSGPTSGPTSGPTSGPTSGPTSGPTSGPTSGPTSGPTSGPTSGPT<br>SGPTSGPTSGPTSGPTSGPTSGPTSGPTSGPTSGPTSGPTSGPTSGPTSGPTSGPTSGPTSGPTSGPTSGPTSGPTSGPTSG<br>PTSGPTSGPTSGPTSGPTSGPTSGPTSGPTSGPTSGPTSGPTSGPTSGPTSGPTSGPTSGPTSGPTSGPTSGPTSGPTSGPT<br>SGPTSGPTSGPTSGPTSGPTSGPTSGPTSGPTSGPTSGPTSGPTSGPTSGPTSGPTSGPTSGPTSGPTSGPTSGPTSGPTSG<br>PTSGPTSGPTSGPTSGPTSGPTSGPTSGPTSGPTSGPTSGPTSGPTSGPTSGPTSGPTSGPTSGPTSGPTSGPTSGPTSGPT<br>SGPTSGPTSGPTSGPTSGPTSGPTSGPTSGPTSGPTSGPTSGPTSGPTSGPTSGPTSGPTSGPTSGPTSGPTSGPTSGPTSG<br>PTSGPTSGPTSGPTSGPTSGPTSGPTSGPTSGPTSGPTSGPTSGPTSGPTSGPTSGPTSGPTSGPTSGPTSGPTSGPTSGPT<br>SGPTSGPTSGPTSGPTSGPTSGPTSGPTSGPTSGPTSGPTSGPTSGPTSGPTSGPTSGPTSGPTSGPTSGPTSGPTSGPTSG<br>PTSGPTSGPTSGPTSGPTSGPTSGPTSGPTSGPTSGPTSGPTSGPTSGPTSGPTSGPTSGPTSGPTSGPTSGPTSGPTPQPT<br>SQPTASAAPTNCVDPDGAKFVRSTDPSYTADPKPIILTFESDDATLVEFDFIQNIINGVIPHVFIKYRKSETGINECAEFAN<br>VGENTPISLSAECVDMSTNVVIYIYTGDDFASENCEACSLPSDDGEGGSGNPDNFAAYYFDVPCNTECIEPTAAPTSAPTSG<br>PTSGPTSGPTSGPTSGPTSGPTSGPTSGPTSGPTSGPTSGPTSGPTSGPTSGPTSGPTSGPTSGPTSGPTSGPTSGPTSGPT<br>SGPTSGPTSGPTSGPTSGPTSGPTSGPTSGPTSGPTSGPTSGPTSGPTSGPTSGPTSGPTSGPTSGPTSGPTSGPTSGPTSG<br>PTSGPTSGPTSGPTSGPTSGPTSGPTSGPTSGPTSGPTSGPTSGPTSGPTSGPTSGPTSGPTSGPTSGPTSGPTSGPTSGPT<br>SGPTSGPTSGPTSGPTSGPTSGPTSGPTSGPTSGPTSGPTSGPTSGPTSGPTSGPTSGPTSGPTSGPTSGPTSGPTSGPTSG<br>PTSGPTSGPTSGPTSGPTSGPTSGPTSGPTSGPTSGPTSGPTSGPTSGPTSGPTSGPTSGPTSGPTSGPTSGPTSGPTSGPT<br>SGPTSGPTSGPTSGPTSGPTSGPTSGPTSGPTSGPTSGPTSGPTSGPTSGPTSGPTSGPTSGPTSGPTSGPTSGPTSGPTSG<br>PTSGPTSGPTSGPTSGPTSGPTSGPTSGPTSGPTSGPTSGPTSGPTSGPTSGPTSGPTSGPTSGPTSGPTSGPTSGPTSGPT<br>SGPTSGPTSGPTSGPTSGPTSGPTSGPTSGPTSGPTSGPTSGPTSGPTSGPTSGPTSGPTSGPTSGPTSGPTSGPTSGPTSG<br>PTSGPTSGPTSGPTSGPTSGPTSGPTSGPTSGPTSGPTSGPTSGPTSGPTSGPTSGPTSGPTSGPTSGPTSGPTSGPTSGPT<br>SGPTSGPTSGPTSGPTSGPTSGPTSGPTSGPTSGPTSGPTSGPTSGPTSGPTSGPTSGPTSGPTSGPTSGPTSGPTSGPTSG<br>PTSGPTSGPTSGPTSGPTSGPTSGPTSGPTSGPTSGPTSGPTSGPTSGPTSGPTSGPTSGPTSGPTSGPTSGPTSGPTSGPT<br>SGPTSGPTSGPTSGPTSGPTSGPTSGPTSGPTSGPTSGPTSGPTSGPTSGPTSGPTSGPTSGPTSGPTSGPTSGPTSGPTSG<br>PTSGPTSGPTSGPTSGPTSGPTSGPTSGPTSGPTSGPTSGPTSGPTSGPTSGPTSGPTSGPTSGPTSGPTSGPTSGPTSGPT<br>SGPTSGPTSGPTSGPTSGPTSGPTSGPTSGPTSGPTSGPTSGPTSGPTSGPTSGPTSGPTSGPTSGPTSGPTSGPTSGPTSG<br>PTSGPTSGPTSGPTSGPTSGPTSGPTSGPTSGPTSGPTSGPTSGPTSGPTSGPTSGPTSGPTSGPTSGPTSGPTSGPTSGPT<br>SGPTSGPTSGPTSGPTSGPTSGPTSGPTSGPTSGPTSGPTSGPTSGPTSGPTSGPTSGPTSGPTSGPTSGPTSGPTSGPTSG<br>PTSGPTSGPTSGPTSGPTSGPTSGPTSGPTSGPTSGPTSGPTSGPTSGPTSGPTSGPTSGPTSGPTSGPTSGPTSGPTSGPT<br>SGPTSGPTSGPTSGPTSGPTSGPTSGPTSGPTSGPTSGPTSGPTSGPTSGPTSGPTSGPTSGPTSGPTSGPTSGPTSGPTSG<br>PTSGPTSGPTSGPTSGPTSGPTSGPTSGPTSGPTSGPTSGPTSGPTSGPTSGPTSGPTSGPTSGPTPQPTSQPTASAAPTNC<br>VDPDGAKFVKSTDASYTADPKPIILTFESDDATLVEFDFIQNIINGVIPHVFIQYRKSETGINECAEFANVGENTPISLSAE<br>CVDMSTNVVIYIYTGDDFASENCEACSLPSDDGEGGSGNPDNFAAYYFDVPCNTECIEPTAAPTSAPTSGPTSGPTSGPTSG<br>PTSGPTSGPTSGPTSGPTSGPTSGPTSGPTSGPTSGPTSGPTSGPTSGPTSGPTSGPTSGPTSGPTSGPTSGPTSGPTSGPT<br>SGPTSGPTSGPTSGPTSGPTSGPTSGPTSGPTSGPTSGPTSGPTSGPTSGPTSGPTSGPTSGPTSGPTSGPTSGPTSGPTSG<br>PTSGPTSGPTSGPTSGPTSGPTSGPTSGPTSGPTSGPTSGPTSGPTSGPTSGPTSGPTSGPTSGPTSGPTSGPTSGPTSGPT<br>SGPTSGPTSGPTSGPTSGPTSGPTSGPTSGPTSGPTSGPTSGPTSGPTSGPTSGPTSGPTSGPTSGPTSGPTSGPTSGPTSG<br>PTSGPTSGPTSGPTSGPTSGPTSGPTSGPTSGPTSGPTSGPTSGPTSGPTSGPTSGPTSGPTSGPTSGPTSGPTSGPTSGPT<br>SGPTSGPTSGPTSGPTSGPTSGPTSGPTSGPTSGPTSGPTSGPTSGPTSGPTSGPTSGPTSGPTSGPTSGPTSGPTSGPTSG<br>PTSGPTSGPTSGPTSGPTSGPTSGPTSGPTSGPTSGPTIGPTSGPTSGPTSGPTSGPTSGPTSGPTSGPTSGPTSGPTSGPT<br>SGPTSGPTSGPTSGPTSGPTSGPTSGPTSGPTSGPTSGPTSGPTSGPTSGPTSGPTSGPTSGPTSGPTSGPTSGPTSGPTSG<br>PTSGPTSGPTSGPTSGPTSGPTSGPTSGPTSGPTSGPTSGPTSGPTSGPTSGPTSGPTSGPTSGPTSGPTSGPTSGPTSGPT<br>SGPTSGPTSGPTSGPTSGPTSGPTSGPTSGPTSGPTSGPTSGPTSGPTSGPTSGPTSGPTSGPTSGPTSGPTSGPTSGPTSG<br>PTSGPTSGPTSGPTSGPTSGPTSGPTSGPTSGPTSGPTSGPTSGPTSGPTSGPTSGPTSGPTSGPTSGPTSGPTSGPTSGPT<br>SGPTSGPTSGPTSGPTSGPTSGPTSGPTSGPTSGPTSGPTSGPTSGPTSGPTSGPTSGPTSGPTSGPTSGPTSGPTSGPTSG<br>PTSGPTSGPTSGPTSGPTSGPTSGPTSGPTSGPTSGPTSGPTSGPTSGPTSGPTSGPTSGPTSGPTSGPTSGPTSGPTSGPT<br>SGPTSGPTSGPTSGPTSGPTSGPTSGPTSGPTSGPTSGPTSGPTSGPTSGPTSGPTSGPTSGPTSGPTSGPTSGPTSGPTSG |

| | |
|---|---|
| | PTSGPTSGPTSGPTSGPTSGPTSGPTSGPTSGPTSGPTSGPTSGPTSGPTSGPTSGPTSGPTSGPTSGPTSGPTSGPTSGPT<br>SGPTSGPTSGPTSGPTSGPTSGPTSGPTSGPTSGPTSGPTSGPTSGPTSGPTSGPTSGPTSGPTSGPTSGPTSGPTSGPTSG<br>PTSGPTSGPTSGPTSGPTSGPTSGPTSGPTSGPTSGPTSGPTSGPTSGPTSGPTSGPTSGPTSGPTSGPTSGPTSGPTSGPT<br>SGPTSGPTSGPTSGPTSGPTSGPTSGPTSGPTSGPTSGPTSGPTSGPTSGPTSGPTSGPTSGPTSGPTSGPTSGPTSGPTSG<br>PTSGPTSGPTSGPTSGPTSGPTSGPTSGPTSGPTSGPTSGPTSGPTSGPTSGPTSGPTSGPTSGPTSGPTSGPTSGPTSGPT<br>SGPTSGPTSGPTSGPTSGPTSGPTETACIYTVLDFATDAGGNPTGNGYVPSDVYSQYDIDAITVDVTGSPGSYGGPNNMSGR<br>YWVTGNPLVGTALIIQGAEGPTAVPSDDGGDIVFTLDGASGEEMQLTFFDLAEDVTITLNTETDPEAGGTSTTVPVVADPSG<br>GLQSIEIDISLIKQVIVTFTGRGAIASVGICRDPNTPPPMGNNSPSATLEPTSMPTTSPAPTETPGCPEDIVLVASVGHTLP<br>PRFVITIIEQTTDTVTFEVTQKFNKTVSRVFTQYHETPAGETECFEDDNVPYEETQRHTAYCMHCVPLTIVDLWFSDPTLEP<br>TLDNAIVPPCCHPPTNETYPTVQYTFKLYCETKCPPDGEPNQFQERRLLEAPEEPVEVEAEAITDIAPAQEEEDDSKHACSS<br>KDHPCNGDMVYVCHYSSRHGYQTYCVPEADSDIVEFYPKSYCGRCTGGQGNYIRE* |
| **Ca11384** | |
| **gDNA** | ATGAACATCCTCGCAGCATCGCTGCTGGCGGCTCTCGGAGGCACGGCCGTCAGCGCTGCCCCTGAGTACGAGACCGTCCCGG<br>TCTCCAAGCCAACCTGCGCCAAGCCGGCCAGAAGCGGAACCTGCCACGCGGCTGGTCTCCTCAACGCCGGTGAGGAAGCCCT<br>CACTCTTGCCCCTGAGCAACCAGCCGACCTTGAGACCGTCCGCGTCGTCCTCAAGCAGAGCTTCTTCGAGGGACAGGACATC<br>GCCTTCGCCGGTGTGGTCATCCCCTCGATCTACACCAACGGCGAGCCGATCTGCATCGAGAACTTCGGACCCCTTGCCTTCG<br>GCGCCAACCTCGATCATGAAGGTCACCCATTCGACCTCATGTGCAATCCCGCCACCAACACCGCCAAGGTCTCCATCTACAT<br>CTCCGACGACAAGATCGACGCAGACAAGCTCGATGAGCTGGCCGAGGACTCGGACGGCCAATTCCCGGACGAGTGCGATCTC<br>GGCGAGTTCGTCCCCACCGAGATGTGCAGCGTGACTTACGAGATCCAGTGCGGATGCTCTGAGCACGGTGACGAAGAGGAGG<br>AGGAACTCGCCGACGCCGAGGATGGCATCGAGGATGACATTGAGGATGCTCATGCCGAAACCGAGGAAGCCATCGATGCCCA<br>CCTCGACGTGAGAGTCAATGCCGCTGCCGATGTTGTTGACATGGAGGATGGCGCCAATATCAAGCGCAAGCTCGATGCTCAG<br>CTCCGTAAGTTATCACGGCCGTCTTCATCTTCCAACACGCGCGAATGTGAAGCTTGTCTTGCATACTCGCACTGTGCTTTCG<br>TGATCCGGCTGTAACTAACCTATTCATTCGCTCCCTTCCTTCCTTCCTTCTTTCCTTCATTTGCCTCTTTTTCTACCACATT<br>ATCCACCGATGATCCACATTCCACGTGGCACACTTCGCGCCACCTTCTCCACGCGTTCCTCCGCCACATGCAGGCCCGACCC<br>CGGCACCGACCAGCGCCCCGACTAGCGCTCCAACACGCCCCAAGCCCACCAACCCGCCCAGCGTGAGCTTCGAGCCGACCAC<br>CAGTTTGGCACCGACCGAAGAGCAGGGACCAGGCTGCGACAGCATCACCTACCTTCCAGAGGGACACACCGGACCTCTTCCG<br>TTCTGCTACGATGGAAGCACGTACCATGGAACATACCGCGGCAACGGACGGGGTCGCGGCAAGGCTGACACCACCTGCTATG<br>TCGTCGAGGAGTCCGAATGCGGAATGATCGACTTCATGATCCAGGGCGCCAACTCGATGGACAACTACCGTTGCTCCGACGT<br>CTTTGACTGGTACATCCGCTGCGTGGCCTCCTACGGTTGCACTCCTGCCCATGACAGCACAGTCTACTTCGCCTACGGCAAG<br>CCAGGAACGCGAAGGGTGAGCTGGATCGGGGCCAGCAGCGGCCAGCCAGGATACTACCGATTCACTCTCGACGGAAATGGTG<br>TCTGGGGTGGTGCCTACAAGTGGAAATGTCCAAGCAAGGAAGCCGCGTACGGAATGAAGCACTTCACTGACGCCGTCTACAA<br>CACGAAGGTCCCTGGCGACTGGCGCCGTCTGTTCCAGACACCCGTCGGAACTTGTGAAAGCTCTGCCCCATCCGAGAGTCTG<br>GTCCCCACCTCTAGCCCTGCCCCGTCCCTCCCAGTCCCACCGGACTGCACATCGGACTTCGGATATCACTTCCCATACTCCA<br>AGCCACGAACAAGCAACCACGTCAACAGGAGAAGCCAGTCTGGTCCCAAGGGAAAGATATACATCGTCGTCAACAAGCACTG<br>CCAGTACTACATGTACGGCGCCGGTGTGCAAGGCACCAAGTACGTGAACAACATGGAAGAGATGGTAGAAGGCGTCGCCAAG<br>GCTTGGGGGGCGCAGATGGTCAACAACAACGCCAAGTTCAACCCTCTCGGGCGAATGAACCAAGGGATGATCATCTCCGGCA<br>ATTGCGGTAACCAGCTGTGGTGCACAAAGACTGCGTACGTCTCCTTCCGCAAGCACAGCGGTGTCGAGAACGAAGACAAGAT<br>CTTCCTCTTCGATTCGAGGACCTACGCCGACCGCTTCCGCCGACTGTTCTACGATACGGCCAAGAATGACAACGAGGTCTTC<br>TTCACCTTGGACGAGAACAACAAGTGCGGTGTGTCCCCAATGTGCACTGACGACGGGAAGACAGCATGCTACGTGGACACCA<br>AGAAGAGCGATTTTTGCGGCCCGTTCGACTTCGTCGTCGTGAACACTGAGGGCGACGAGCAGCTGAGGACCAACGATGTGTG<br>GGAATTCTTCACCGACTGCGCAGCGTCGTGGCCATCGTGCACCAAGTATGAATACGGAACCGTCCAGTTCGTCTACGGCAGA<br>TCTCCAAGAGTGCACAGCTGGAGAAGTGGTGGCAAGCGAATCGTCCGTCTCAAGGACAACGGCTTCAAGGGAGAGGTGCAGT<br>ACCACTGCACCAACAACGAGCAAGGTCTTGACCTGTACCACTTGGCCTACAAGATGTACTGCACTGATCTCTTCAAGAAGAA<br>GAAGAGCGAGTGCACACAAAGTCCAACCGACTCGCCAACTGCCTGCCAAGACAGCTTCCTCACCAAGCCACCGATCAACATC<br>AACATCGTCATGGATATTTCGTACTCCACCTACTACACCCCCTTCGGCGGTTCCTGGATCGGCGACCTCAACGGTGATGGCA<br>AGAGAAACTCCATCCTGGATGCAGAGATTGCCGCCGTCCTTCAATTGCTCAAGGTCATCGAGAACTCGGACGTCCTCGACAA<br>CACCAACGTCGATCTCGGTTTCGTGGTCTTCGAGACTTCTGGCTTCTACAAGGGCGAATACATGCACGCAGGACGTTACAAG<br>CCACTCAACAGCAATGGCAATGGCATCAACTCCGACCTCATCAAGGACCTCAAAGACATCCAGACTCTCAAGACGAACGCAA<br>CTGTCATCCAAAACAACAAGGGATTCACCAACTTCGACGATGCCTTGGACAAGTCCATCGAGTACTTCATGGAACGCGAGAA<br>TGACTCGACCATCAACTTCGACGAGTGGACCAACCTGATGGTGTTCCTTTCTGACGGAATCCCCAACATCCGCGGTGACGGA<br>GACAATGAGCCATTCTGCAAGCACGGCCTGAGCAATGTGAACTGCAACAACCCCAACGCAAGGAACAAGCCTGACACCGCCG<br>CAACCCTGCGATGGGAGAGCGGAGAGCTCTCCTTCTGCCAGTCTGGCGATACGGGATGTGCTGACAACAAGTACAAGCCATG<br>CGTCCTTGGACCCAAATGCGAGGGCGATGACTCCGTGAAGATGTTCGGGAGCGAGTTCAAGCGCCTTGACATGTTCGGGCGA<br>GACGGTGTGAAGCGAATCTCCATCGGTGTGGGAAAGAGCAGCGACGGCAGCATGGGAAGCGCCCTGTACTCCATCGACAACA<br>ACCCGCTGAAGAAAACCTTCCCCGATCTCCTTCCACCGGTTGTGACCACGACCGATGCCCTGGCCAACCTCCTCCGAAACTT<br>GTGTTACGAAAACACGGCAAACCCGACGGATCAGCCATCTGTCAGCCCGACATCCATCCCAACCACCTTACCGACTGGCTCC<br>CCAACTCACGAACCAACATCCGCACCAACCAGCAAGCCAACCAGCGCACCGACGTCCCCACCGACGGGCCTGCCAACCAGCG<br>CACCAACCGACGAGCCAACCGAGACACCGTCTTCCCAGCCCACCGACGACCTTTGCGACTACACCGTCCTCGACTTCGCCGG<br>CGAGACCGGCCCGGCGCTGCCGGACACTTGGACACCAACGTACGGAATGCGTATCATGTCCATGGTGTCGGACGTCGCAGGA<br>AACCTCGCGTACATCGACACGAACGATGCCGAGGCAGGAGTCGTCCTTGCTGTGCCAAAGGCAGATTCCGTGGTCGACGAGG<br>TGAGCCCGGACGGAGGATCCATCCTCATCAACCAGGACCTCACCCAAGACAAGTTCATGCGCATAAGTCTGTACAATGTCGA<br>AAATGAGTTCAAGATCATTCGCCGATTCCGGGAGATGAGCATGTTCGGCGAAGTCGTCGGAACTGTCGCCTCTGTTGATGAG<br>CACGATGCCTTCACCGGCGTCAAGGAGTTCAAGATCCCAGTCGACGGCATTTTTCAGGTCGAGGTCCAGTTCGAAGGACACG<br>GTAGCCTCTCCAAGCTCCAGATCTGCCGCAATCCCAACAGCACCCCGGCTCCATTCGGATTGTCCTCTAACGGATCCACGGA<br>CCCACCAACCGCAACCCCAACCACCAACACCGCTACCCCGACCATGTTGCCTACTGTGGACGACGGATGCCCGCCAGATGCT<br>GTCGTCATCAGCACCACTGGAAGCACCCCGTTCCCACAAAACCCCATTGAGGTCGTGTCTCAAGATGGAACGACTGTTACGT<br>TCAAGGTCAAGCAGACCTGGAAGGAGACCGTGGGATACCTGTACACCAACTACCAGACCTCCTTCTTCGACAGCGAGTGCAT<br>CTTGCTGGAGGAAGTGGAGAAGTACGCTAGCGAGACCTACACCGCCAAATGCATGGACTCTGTCGAGATCGCCGTGGTGGAC |

| | |
|---|---|
| | ATTTACGTCTCTGATCCCTCACTCAACAAGGTCTTGGACGACGCCGTCATCCCAATCTGCTGCCAGAAGCCGGACGTTGATA<br>GCCACCCCACCGTGAAATACACGTTCGAGGTCAAGTGCGTGGATCCATGCCCAACCCCGGCACGCAAGCTCAGCGACAAGCA<br>GGTTGTTGGCGACGAAGTCGACGCCAAGTACAGCAAGAAGACGATCATCAAGACCGACAAAGATGGCAAGGCCGAGACCGAG<br>ACCAGCAGCACAGACCACTACTGCAAGCCGAGCGAGCACCCGTGCAGTGACGATTCCAGCATGGTGAACGTGTGCCATTACT<br>CATCAAAGAAGGGCTTCCAGACTTACTGCATGACTGCCGCCGATTCTGACATCGTCTCGTACTATCCAAACGACTCCTGCGG<br>CCCTTGCCCGCCAGAGCTCTCGGAGAAGGCCAGCGTCAACGCCGAAGTTGCCAAGTAGGCGTGATGCGACGAGCCGATCGGA<br>CCGTCCGTCGCAGCGCAATTGA |
| **cDNA** | ATGAACATCCTCGCAGCATCGCTGCTGGCGGCTCTCGGAGGCACGGCCGTCAGCGCTGCCCCTGAGTACGAGACCGTCCCGG<br>TCTCCAAGCCAACCTGCGCCAAGCCGGCCAGAAGCGGAACCTGCCACGCGGCTGGTCTCCTCAACGCCGGTGAGGAAGCCCT<br>CACTCTTGCCCCTGAGCAACCAGCCGACCTTGAGACCGTCCGCGTCGTCCTCAAGCAGAGCTTCTTCGAGGGACAGGACATC<br>GCCTTCGCCGGTGTGGTCATCCCCTCGATCTACACCAACGGCGAGCCGATCTGCATCGAGAACTTCGGACCCCTTGCCTTCG<br>GCGCCAACCTCGATCATGAAGGTCACCCATTCGACCTCATGTGCAATCCCGCCACCAACACCGCCAAGGTCTCCATCTACAT<br>CTCCGACGACAAGATCGACGCAGACAAGCTCGATGAGCTGGCCGAGGACTCGGACGGCCAATTCCCGGACGAGTGCGATCTC<br>GGCGAGTTCGTCCCCACCGAGATGTGCAGCGTGACTTACGAGATCCAGTGCGGATGCTCTGAGCACGGTGACGAAGAGGAGG<br>AGGAACTCGCCGACGCCGAGGATGGCATCGAGGATGACATTGAGGATGCTCATGCCGAAACCGAGGAAGCCATCGATGCCCA<br>CCTCGACGTGAGAGTCAATGCCGCTGCCGATGTTGTTGACATGGAGGATGGCGCCAATATCAAGCGCAAGCTCGATGCTCAG<br>CTCCGCCCGACCCCGGCACCGACCAGCGCCCCGACTAGCGCTCCAACACGCCCCAAGCCCACCAACCCGCCCAGCGTGAGCT<br>TCGAGCCGACCACCAGTTTGGCACCGACCGAAGAGCAGGGACCAGGCTGCGACAGCATCACCTACCTTCCAGAGGGACACAC<br>CGGACCTCTTCCGTTCTGCTACGATGGAAGCACGTACCATGGAACATACCGCGGCAACGGACGGGGTCGCGGCAAGGCTGAC<br>ACCACCTGCTATGTCGTCGAGGAGTCCGAATGCGGAATGATCGACTTCATGATCCAGGGCGCCAACTCGATGGACAACTACC<br>GTTGCTCCGACGTCTTTGACTGGTACATCCGCTGCGTGGCCTCCTACGGTTGCACTCCTGCCCATGACAGCACAGTCTACTT<br>CGCCTACGGCAAGCCAGGAACGCGAAGGGTGAGCTGGATCGGGGCCAGCAGCGGCCAGCCAGGATACTACCGATTCACTCTC<br>GACGGAAATGGTGTCTGGGGTGGTGCCTACAAGTGGAAATGTCCAAGCAAGGAAGCCGCGTACGGAATGAAGCACTTCACTG<br>ACGCCGTCTACAACACGAAGGTCCCTGGCGACTGGCGCCGTCTGTTCCAGACACCCGTCGGAACTTGTGAAAGCTCTGCCCC<br>ATCCGAGAGTCTGGTCCCCACCTCTAGCCCTGCCCCGTCCCTCCCAGTCCCACCGGACTGCACATCGGACTTCGGATATCAC<br>TTCCCATACTCCAAGCCACGAACAAGCAACCACGTCAACAGGAGAAGCCAGTCTGGTCCCAAGGGAAAGATATACATCGTCG<br>TCAACAAGCACTGCCAGTACTACATGTACGGCGCCGGTGTGCAAGGCACCAAGTACGTGAACAACATGGAAGAGATGGTAGA<br>AGGCGTCGCCAAGGCTTGGGGGGCGCAGATGGTCAACAACAACGCCAAGTTCAACCCTCTCGGGCGAATGAACCAAGGGATG<br>ATCATCTCCGGCAATTGCGGTAACCAGCTGTGGTGCACAAAGACTGCGTACGTCTCCTTCCGCAAGCACAGCGGTGTCGAGA<br>ACGAAGACAAGATCTTCCTCTTCGATTCGAGGACCTACGCCGACCGCTTCCGCCGACTGTTCTACGATACGGCCAAGAATGA<br>CAACGAGGTCTTCTTCACCTTGGACGAGAACAACAAGTGCGGTGTGTCCCCAATGTGCACTGACGACGGGAAGACAGCATGC<br>TACGTGGACACCAAGAAGAGCGATTTTTGCGGCCCGTTCGACTTCGTCGTCGTGAACACTGAGGGCGACGAGCAGCTGAGGA<br>CCAACGATGTGTGGGAATTCTTCACCGACTGCGCAGCGTCGTGGCCATCGTGCACCAAGTATGAATACGGAACCGTCCAGTT<br>CGTCTACGGCAGATCTCCAAGAGTGCACAGCTGGAGAAGTGGTGGCAAGCGAATCGTCCGTCTCAAGGACAACGGCTTCAAG<br>GGAGAGGTGCAGTACCACTGCACCAACAACGAGCAAGGTCTTGACCTGTACCACTTGGCCTACAAGATGTACTGCACTGATC<br>TCTTCAAGAAGAAGAAGAGCGAGTGCACACAAAGTCCAACCGACTCGCCAACTGCCTGCCAAGACAGCTTCCTCACCAAGCC<br>ACCGATCAACATCAACATCGTCATGGATATTTCGTACTCCACCTACTACACCCCCTTCGGCGGTTCCTGGATCGGCGACCTC<br>AACGGTGATGGCAAGAGAAACTCCATCCTGGATGCAGAGATTGCCGCCGTCCTTCAATTGCTCAAGGTCATCGAGAACTCGG<br>ACGTCCTCGACAACACCAACGTCGATCTCGGTTTCGTGGTCTTCGAGACTTCTGGCTTCTACAAGGGCGAATACATGCACGC<br>AGGACGTTACAAGCCACTCAACAGCAATGGCAATGGCATCAACTCCGACCTCATCAAGGACCTCAAAGACATCCAGACTCTC<br>AAGACGAACGCAACTGTCATCCAAAACAACAAGGGATTCACCAACTTCGACGATGCCTTGGACAAGTCCATCGAGTACTTCA<br>TGGAACGCGAGAATGACTCGACCATCAACTTCGACGAGTGGACCAACCTGATGGTGTTCCTTTCTGACGGAATCCCCAACAT<br>CCGCGGTGACGGAGACAATGAGCCATTCTGCAAGCACGGCCTGAGCAATGTGAACTGCAACAACCCCAACGCAAGGAACAAG<br>CCTGACACCGCCGCAACCCTGCGATGGGAGAGCGGAGAGCTCTCCTTCTGCCAGTCTGGCGATACGGGATGTGCTGACAACA<br>AGTACAAGCCATGCGTCCTTGGACCCAAATGCGAGGGCGATGACTCCGTGAAGATGTTCGGGAGCGAGTTCAAGCGCCTTGA<br>CATGTTCGGGCGAGACGGTGTGAAGCGAATCTCCATCGGTGTGGGAAAGAGCAGCGACGGCAGCATGGGAAGCGCCCTGTAC<br>TCCATCGACAACAACCCGCTGAAGAAAACCTTCCCCGATCTCCTTCCACCGGTTGTGACCACGACCGATGCCCTGGCCAACC<br>TCCTCCGAAACTTGTGTTACGAAAACACGGCAAACCCGACGGATCAGCCATCTGTCAGCCCGACATCCATCCCAACCACCTT<br>ACCGACTGGCTCCCCAACTCACGAACCAACATCCGCACCAACCAGCAAGCCAACCAGCGCACCGACGTCCCCACCGACGGGC<br>CTGCCAACCAGCGCACCAACCGACGAGCCAACCGAGACACCGTCTTCCCAGCCCACCGACGACCTTTGCGACTACACCGTCC<br>TCGACTTCGCCGGCGAGACCGGCCCGGCGCTGCCGGACACTTGGACACCAACGTACGGAATGCGTATCATGTCCATGGTGTC<br>GGACGTCGCAGGAAACCTCGCGTACATCGACACGAACGATGCCGAGGCAGGAGTCGTCCTTGCTGTGCCAAAGGCAGATTCC<br>GTGGTCGACGAGGTGAGCCCGGACGGAGGATCCATCCTCATCAACCAGGACCTCACCCAAGACAAGTTCATGCGCATAAGTC<br>TGTACAATGTCGAAAATGAGTTCAAGATCATTCGCCGATTCCGGGAGATGAGCATGTTCGGCGAAGTCGTCGGAACTGTCGC<br>CTCTGTTGATGAGCACGATGCCTTCACCGGCGTCAAGGAGTTCAAGATCCCAGTCGACGGCATTTTTCAGGTCGAGGTCCAG<br>TTCGAAGGACACGGTAGCCTCTCCAAGCTCCAGATCTGCCGCAATCCCAACAGCACCCCGGCTCCATTCGGATTGTCCTCTA<br>ACGGATCCACGGACCCACCAACCGCAACCCCAACCACCAACACCGCTACCCCGACCATGTTGCCTACTGTGGACGACGGATG<br>CCCGCCAGATGCTGTCGTCATCAGCACCACTGGAAGCACCCCGTTCCCACAAAACCCCATTGAGGTCGTGTCTCAAGATGGA<br>ACGACTGTTACGTTCAAGGTCAAGCAGACCTGGAAGGAGACCGTGGGATACCTGTACACCAACTACCAGACCTCCTTCTTCG<br>ACAGCGAGTGCATCTTGCTGGAGGAAGTGGAGAAGTACGCTAGCGAGACCTACACCGCCAAATGCATGGACTCTGTCGAGAT<br>CGCCGTGGTGGACATTTACGTCTCTGATCCCTCACTCAACAAGGTCTTGGACGACGCCGTCATCCCAATCTGCTGCCAGAAG<br>CCGGACGTTGATAGCCACCCCACCGTGAAATACACGTTCGAGGTCAAGTGCGTGGATCCATGCCCAACCCCGGCACGCAAGC<br>TCAGCGACAAGCAGGTTGTTGGCGACGAAGTCGACGCCAAGTACAGCAAGAAGACGATCATCAAGACCGACAAAGATGGCAA<br>GGCCGAGACCGAGACCAGCAGCACAGACCACTACTGCAAGCCGAGCGAGCACCCGTGCAGTGACGATTCCAGCATGGTGAAC<br>GTGTGCCATTACTCATCAAAGAAGGGCTTCCAGACTTACTGCATGACTGCCGCCGATTCTGACATCGTCTCGTACTATCCAA<br>ACGACTCCTGCGGCCCTTGCCCGCCAGAGCTCTCGGAGAAGGCCAGCGTCAACGCCGAAGTTGCCAAGTAGGCGTGATGCGA<br>CGAGCCGATCGGACCGTCCGTCGCAGCGCAATTGA |

| | |
|---|---|
| **Protein** | MNILAASLLAALGGTAVSAAPEYETVPVSKPTCAKPARSGTCHAAGLLNAGEEALTLAPEQPADLETVRVVLKQSFFEGQDI<br>AFAGVVIPSIYTNGEPICIENFGPLAFGANLDHEGHPFDLMCNPATNTAKVSIYISDDKIDADKLDELAEDSDGQFPDECDL<br>GEFVPTEMCSVTYEIQCGCSEHGDEEEEELADAEDGIEDDIEDAHAETEEAIDAHLDVRVNAAADVVDMEDGANIKRKLDAQ<br>LRPTPAPTSAPTSAPTRPKPTNPPSVSFEPTTSLAPTEEQGPGCDSITYLPEGHTGPLPFCYDGSTYHGTYRGNGRGRGKAD<br>TTCYVVEESECGMIDFMIQGANSMDNYRCSDVFDWYIRCVASYGCTPAHDSTVYFAYGKPGTRRVSWIGASSGQPGYYRFTL<br>DGNGVWGGAYKWKCPSKEAAYGMKHFTDAVYNTKVPGDWRRLFQTPVGTCESSAPSESLVPTSSPAPSLPVPPDCTSDFGYH<br>FPYSKPRTSNHVNRRSQSGPKGKIYIVVNKHCQYYMYGAGVQGTKYVNNMEEMVEGVAKAWGAQMVNNNAKFNPLGRMNQGM<br>IISGNCGNQLWCTKTAYVSFRKHSGVENEDKIFLFDSRTYADRFRRLFYDTAKNDNEVFFTLDENNKCGVSPMCTDDGKTAC<br>YVDTKKSDFCGPFDFVVVNTEGDEQLRTNDVWEFFTDCAASWPSCTKYEYGTVQFVYGRSPRVHSWRSGGKRIVRLKDNGFK<br>GEVQYHCTNNEQGLDLYHLAYKMYCTDLFKKKKSECTQSPTDSPTACQDSFLTKPPININIVMDISYSTYYTPFGGSWIGDL<br>NGDGKRNSILDAEIAAVLQLLKVIENSDVLDNTNVDLGFVVFETSGFYKGEYMHAGRYKPLNSNGNGINSDLIKDLKDIQTL<br>KTNATVIQNNKGFTNFDDALDKSIEYFMERENDSTINFDEWTNLMVFLSDGIPNIRGDGDNEPFCKHGLSNVNCNNPNARNK<br>PDTAATLRWESGELSFCQSGDTGCADNKYKPCVLGPKCEGDDSVKMFGSEFKRLDMFGRDGVKRISIGVGKEAADGSMGSAL<br>YSIDNNPLKKTFPDLLPPVVTTTDALANLLRNLCYENTANPTDQPSVSPTSIPTTLPTGSPTHEPTSAPTSKPTSAPTSPPT<br>GLPTSAPTDEPTETPSSQPTDDLCDYTVLDFAGETGPALPDTWTPTYGMRIMSMVSDVAGNLAYIDTNDAEAGVVLAVPKAD<br>SVVDEVSPDGGSILINQDLTQDKFMRISLYNVENEFKIIRRFREMSMFGEVVGTVASVDEHDAFTGVKEFKIPVDGIFQVEV<br>QFEGHGSLSKLQICRNPNSTPAPFGLSSNGSTDPPTATPTTNTATPTMLPTVDDGCPPDAVVISTTGSTPFPQNPIEVVSQD<br>GTTVTFKVKQTWKETVGYLYTNYQTSFFDSECILLEEVEKYASETYTAKCMDSVEIAVVDIYVSDPSLNKVLDDAVIPICCQ<br>KPDVDSHPTVKYTFEVKCVDPCPTPARKLSDKQVVGDEVDAKYSKKTIIKTDKDGKAETETSSTDHYCKPSEHPCSDDSSMV<br>NVCHYSSKKGFQTYCMTAADSDIVSYYPNDSCGPCPPELSEKASVNAEVAK* |
| **Ca5609** | |
| **gDNA** | ATGATGATGGTAACCAAGCGAGTTACTCCCATCCCCCCGTGGGCGGGTCGGAGCGGACTTCTCTTGGTGCTATGTGCACTGG<br>TTTTGACAGGTGGTAAAGAGGCGGAGGGCCTCAAGGTGACCAGGGTTTCCTCTGGTCAGGGTTACCATGGCGGACCTTCCGA<br>TGGTGTCGATCACCAAAGGAAGTTGCTTGGCGATGCGAATCGCCATGGCAAGCTGCTTGGTGGAGAGGAGGAGAAAGATGAG<br>GCCGAAGGCAAACGCGAAGGCGGAGCCATGCGGTTCCTGTACGGTGCACAGGTGGATGTTGGAGCGAGAACGGGGAGTTTCT<br>ACTCCGCGTTGGGAAGGAATGTGAAGCAGGAGACTTTGGAGAAAGCTCAAGATTGTGATCCAGACAATAAGAGCACTTGGAA<br>GTTCTGCGCGCCTCACATGCTTCCCAATTCGAACTTCAACCCAAACTTGCCCCCCGAGATTGCGGCCCAGCGAAGCAACGAA<br>GTTGCGTACCTGGCTAAGCAAGCGAAGGGAGGTGGGCTCACCGGACCCAACAAAGGGAACGGCCTTGACAAGGCCAAACCCA<br>TCGTTGGCTCCGAGAGTCCTGGAAGAAGCGTCGACGCACAAAGCGACGGAGGAACCCTCTTGGGTGACTCCGAAGCGGAAAG<br>CGCCGATGCGACTGTCGGTTCTACGATCGGTGCGAAGGACGCTGGAGGATTTGGGAGTGACGGTGGAGTCAAGAACAAGAGA<br>ACCTCTCCTGGTAGCAATACTGTTGGAGAGGGTCTGGTTCGCGATGAAGCTGTCCCGACCAATGACTCGGTGTCCGCCCGCA<br>CCAGTCTTGACGAGCTGAGCAGCTTGCCAGCGATCTGCGTTCTAGATCCTGATCATCCTGCGTGCCAAGAGGCACGGGCCAG<br>GATGCTTCAGACCACGGACGCGGGGGGCGGCAGCTCTGCTCCCTCGCTTGCGAGTGCAACCGACAACAGCCCGGTCGAAGCC<br>CCCGTTTCAGCCGAAAACCCATATGCCTCGGACTGGCCTTCTGGTTCCAAAACCACCGAATCGCCAACCAAGGATATGTGTT<br>ACGGTAACATGGAAATAGCGAAGCGCAAGTTCTGCGAGTCCGAGATGCCCTCAGTTTCGTCTGAGCCATCTCAAACGCCGAC<br>AGCGTTCTCTCCAAGCCCAGAAAGCGGACAGACCATCCCAACCGCGACGACCCCCTCAGATCCTGGTTCACCATCTGGACAA<br>GGATCCGATGCCCCAAGCGACGTTCCCAGCGAAAGTCCAACGGGCGAGGGTGGAAGCCGTACCCGCCGTCCAGCGGACTCCC<br>CTGACAGCTTCAAGCCTCCTGGTAATTCTGGAGCTGGTAGCGACGCTCCAAGCGACGTTCCCAGCGAAAGTCCAACCGGCGA<br>GGGTGGAAGCCGCACCCGGCGTCCAGCGGACTCCCCTGACAGCTTCAAGCCTCCTGGTAATTCTGGAGCTGGTAGCGACGCT<br>CCAAGCGACGTTCCCAGCGAAAGTCCCACCGGTGAGGGCGGAAGCCGTTCCCGGCGGCCCTTCGGAATCCCTGGTACGAGTT<br>TCGATACCCCTGTGGGTACAGATCCTCCAACCAGCAGTTCTGCTCCAACTGACTGTTACGACAGTATCGAGCCAAAACTCGA<br>ACGCCAAGTTGGAGTAGACTTCCCATGGGCTGATGGCGTGGTCGATGTGCAGCATCAAACCGGAAATAGTGTCACGTTCCAA<br>ATCCTTCAGGTTTGGAGCCGAAGCATTCTCCCGATGCTTGCGGTGCACTACCACAACTCTCCTGGATCAACGGTTTGCGATA<br>TGAACGTCGATGTTCAATATCAGGATACCTTTGACTACACGGCTCATTGCTTCGGAAACTATACGGACATTCGCGTTTACCT<br>CTTCGTCGGCGACGACTTCGACGAGGAAAAATGCGAGGCGTGTCAAACGCCACCAGATGATTCCAATGACACAGTGGCCTAC<br>TTCCTTGAACTATCATGTCATGACTGCGTCGAAGTGACTGCTCCTCCGAGCGTCACAGCACCGTCGCAGCCCGTGATTTCAC<br>CGACACCGCCCACCGGCGACGTCGGGAGTGATGCCCCAAGTGACGTGCCGAGTGAAGCGCCTACCGGAGACGGAGGGAGCCG<br>CACTCGTCGCCCACAGTCACCACAGGTGCCTTCTGCACCAACTGGAGACGAGCCAACCGGAGACGAGCCAACTGGAGAAGTC<br>GGAAGTGATGCCCCAAGTGATGTGCCGAGTGAAGCGCCAACCGGAGACGGAGGGAGCCGCACTCGTCGCCCACAGTCACCAC<br>AGGTGCCTTCTGCACCAACTGGAGACGCACCAACTGGAGATGAACCAACTGGAGAAGTCGGAAGTGATGCCCCAAGTGACGT<br>GCCGAGTGAAGCGCCTACCGGAGACGGAGGGAGCCGCACTCGTCGCCCACAGTCACCACAGGTGCCTTCTGCACCAACTGGA<br>GACGCACCAACTGGAGACGAGCCAACTGGAGACGAGCCAACTGGAGAAGTCGGAAGTGATGCCCCAAGTGACGTGCCGAGTG<br>AAGCGCCAACCGGAGACGGAGGGAGTCGCACTCGCCGCCCACAGTCACCACAGGTGCCTTCTGCACCAACTGGAGACGAGCC<br>CACTGGAGACGAGCCAACTGGAGAAGTCGGAAGTGATGCCCCAAGTGATGTGCCGAGTGAGGCGCCAACCGGAGACGGAGGG<br>AGCCGCACTCGTCGCCCACAGTCACCACAGGTGCCTTCTGCACCAACTGGAGACGCACCAACTGGAGACGAGCCAACTGGAG<br>AAGTCGGAAGTGATGCCCCAAGTGACGTGCCGAGTGAAGCGCCTACCGGAGACGGAGGGAGCCGCACTCGTCGCCCACAGTC<br>ACCACAGGTGCCTTCTGCACCAACTGGAGACGAGCCCACTGGAGACGAGCCAACTGGAGAAGTCGGAAGTGATGCCCCAAGT<br>GATGTGCCGAGTGAGGCGCCAACCGGAGACGGAGGGAGCCGCACTCGTCGCCCACAGTCACCACAGGTGCCTTCTGCACCAA<br>CTGGAGACGCACCAACTGGAGACGAGCCAACTGGAGAAGTCGGAAGTGATGCCCCAAGTGACGTGCCGAGTGAAGCGCCTAC<br>CGGAGACGGAGGGAGTCGCACTCGCCGCCCACAGTCGCCACAGGTGCCTTCTGCGCCAACTGGAGACGAGCCCACTGGAGAC<br>GAGCCAACTGGAGAAGTCGGAAGTGATGCCCCAAGTGATGTGCCTAGTGAAGCGCCTACCGGAGACGGAGGGAGCCGCACTC<br>GCCGCCCACAGTCACCACAGGTGCCTTCTACACCAACTGGAGACGAGCCAACCGGAGACGAGCCAACCGGAGACGAGCCAAC<br>TGGAGACGAGCCAACTGGAGAAGTCGGAAGTGATGCCCCAAGTGACGTGCCGAGTGAGGCGCCTACCGGAGACGGAAGGACT<br>CGCACTCGCCGCCCACAGTCGCCACAGGTGCCTTCTGCGCCAAGTGCTGAGGAGCCCACTGGAGACGAGCCAACTGGAGACG<br>AGCCAACTGGAGATGAACCAACTGGAGAAGTCGGAAGTGATGCCCCAAGTGACGTGCCGAGTGAGGCGCCAACCGGAGACGG<br>AAGGACTCGCACTCGCCGCCCACAGTCGCCACAGGTGCCTTCTGCGCCAAGTGCCGAGGAGCCCACTGGAGACGAGCCAACT<br>GGAGACGAACCAACTGGAGACGAACCAACTGGAGACGAACCAACTGGAGACGAACCAACTGGAGACGAGCCAACTGGAGAAG<br>TCGGAAGTGATGCCCCAAGTGACGTGCCGAGTGAGGCGCCTACCGGAGACGGAAGGACTCGCACTCGCCGCCCACAGTCGCC |

| | |
|---|---|
| | ACAGGTGCCTTCTGCGCCAAGTGCTGAGGAGCCCACTGGAGACGAGCCAACTGGAGACGAGCCAACTGGAGACGAGCCAACT<br>GGAGACGAGCCAACTGGAGATGAACCAACTGGAGAAGTCGGAAGTGATGCCCCAAGTGACGTGCCGAGTGAGGCGCCTACCG<br>GAGACGGAAGGACTCGCACTCGCCGCCCACAGTCGCCACAGGTGCCTTCTGCGCCAAGTGCTGAGGAGCCCACTGGAGACGA<br>GCCAACTGGAGATGAACCAACTGGAGATGAACCAACTGGAGAAGTCGGAAGTGATGTCCCAAGTGACGTGCCGAGTGAGGCG<br>CCTACCGGAGACGGAGGGAGCCGCACTCGCCGCCCACAGTCGCCACAGGTGCCTTCTGTGCCAACATCCGGCCCAACTTCTG<br>GACCAACTAGCGGACCAACTGGAGGCCCAACCTCTGGACCTACTTCTGGACCAACCAGCGGACCAACAAGCGGACCAACATC<br>CGGACCAACTTCTGGACCGACTTCTGGACCAACCAGCGGACCAACTTCTGGACCAACTAGCGGACCAACTGGAGGCCCAACC<br>TCTGGACCTACTTCTGGACCAACCAGCGGACCAACATCCGGCCCAACATCTGGACCAACTAGCGGACCTACTTCTGGACCAA<br>CATCTGGACCAACATCCGGCCCAACTTCTGGACCAACTAGCGGACCAACATCCGGCCCAACATCTGGACCAACCAGCGGACC<br>TACTTCTGGACCAACATCTGGACCAACATCCGGCCCAACTTCTGGACCAACTAGCGGACCAACATCCGGCCCAACATCTGGA<br>CCAACCAGCGGACCTACTTCTGGACCAACATCTGGACCAACATCCGGCCCAACTTCTGGACCAACTAGCGGACCAACATCCG<br>GCCCTACATCTGGACCAACCAGCGGACCTACTTCTGGACCAACATCTGGACCAACATCCGGCCCAACTTCTGGACCAACTAG<br>CGGACCAACCAGCGGACCTACTTCTGGACCAACCTCTGGCCCAACCTCTGGACCAACCAGCCAGCCAACCTACGTCTGTAGC<br>ACTGAGGCCCGCCTCGTCGACAAGGACGGTGGCAACACCAAGTACGATGAAGGCCTTGATGACGAGATCACCGGCGACGATG<br>GTATTCCATTCGAGGTGAGCGAGCGTGAAGTCGACAGCGTGAAGATCTCCCTTGAGCCACTCCTTGATGCCATCGAGGCCCT<br>TCGCACCGGCGGTGCTGTAGGCGAGGACGGCGAGCTCGGATCTGTGGCTGTCGTGTTCGACAGAATCAGCACCGACAACGAC<br>GGCAACGAAGTGACCGAGAACGACTTCTGCACGGAGACATCCGTGACTGAGGCCACCGGCCCAGCCGAGGAGTACGTGGCAC<br>AGTGTGTCGAGGATGATGATTTCGCCGGAACCACTGTTGCGATTGTGTACGTTGTCGTCGCCGACAGCGCCAACACTCAGGA<br>GGATCCATCCGGTTCCTTGATCGGAGCGGCAGGTGGCGAGAGCGCCCTGGGCACAAGCGGATGCGGCGTTCCATTCGATGAG<br>GACACCGTTATTCCAGTGATCGTGTACAAGTACGTGGTTCCATGCCAATACTCATGTGATGGGGCACCAACATCATCGCCGA<br>CAGGAGAGCCGACAAGTGCGCCCACGGTATTGCCAGGGCATCCCACGGGATCGCCAGTCACTGCCACGCCAACTTCCACACC<br>AACGTCTCTTCCAACCAGCGGACCTACTTCTGCGCCAACTTATACACCAACATCCGGCCCAACTTCAGGACCAACCAGCGGA<br>CCTACTTCTGGGCCAACTAGCGGACCAACTGGAGGCCCAACCTCTGGACCTACTTCAGGACCAACAAGCGGACCAACAAGCG<br>GACCAACATCCGGCCCAACATCTGGACCAACAAGCGGACCTACTTCTGGACCAACATCCGGCCCAACTTCTGGACCAACTAG<br>CGGACCAACTTCTGGACCAACTTCTGGCCCAACATCCGGACCAACCTCTGGACCTACTTCTGGTCCAACCAGCGGACCAACC<br>TCTGGACCAACCAGCGGACCAACATCCGGCCCCACTTCTGGACCAACATCCGGTCCAACCAGCGGACCAACTGGAGGCCCAA<br>CCTCTGGACCAACTAGCGGACCAACCAGCGGACCAACATCCGGACCAACTTCCGGACCAACCAGCGGACCTACTTCTGGACC<br>AACATCTGGACCAACCTCTGGACCGACTTCTGGACCAACCAGCGGACCTACTTCTGGACCAACTTCCGGACCAACCAGCGGA<br>CCTACATCTGGACCAACATCTGGACCAACATCTGGACCAACATCCGGCCCAACTAGCGGACCAACCAGCGGACCAACATCCG<br>GACCAACTTCCGGACCAACCAGCGGACCTACTTCTGGACCAACCAGCGGACCAACCTCTGGACCAACATCCGGCCCAACTTC<br>TGGACCAACTAGCGGACCAACTTCCGGCCCAACTTCTGGACCAACATCCGGTCCAACCAGCGGACCAACTGGAGGCCCAACC<br>TCTGGACCAACTAGCGGACCAACCAGCGGACCAACATCCGGACCAACTTCCGGACCAACCAGCGGACCTACTTCTGGACCAA<br>CATCTGGACCAACATCCGGCCCAACTTCTGGACCAACTAGCGGACCAACATCCGGCCCAACATCTGGACCAACCAGCGGACC<br>TACTTCTGGACCAACATCTGGACCAACATCCGGCCCAACTTCTGGACCAACTAGCGGACCAACATCCGGCCCAACATCTGGA<br>CCAACCAGCGGACCTACTTCTGGACCAACATCTGGACCAACATCCGGCCCAACTTCTGGACCAACTAGCGGACCAACATCCG<br>GCCCAACATCTGGACCAACCAGCGGACCTACTTCTGGACCAACATCTGGACCAACATCCGGCCCAACTTCTGGACCAACTAG<br>CGGACCAACATCCGGCCCAACATCTGGACCAACCAGCGGACCTACTTCTGGACCAACATCTGGACCAACATCCGGCCCAACT<br>TCTGGACCAACTAGCGGACCAACATCCGGCCCAACATCTGGACCAACCAGCGGACCTACTTCTGGACCAACATCTGGACCAA<br>CATCCGGCCCAACTTCTGGACCAACTAGCGGACCAACATCCGGCCCTACATCTGGACCAACCAGCGGACCTACTTCTGGACC<br>AACATCTGGACCAACATCCGGCCCAACTTCTGGACCAACTAGCGGACCAACTGGAGGCCCAACCTCTGGACCTACTTCTGGA<br>CCAACCAGCGGACCAACAAGCGGACCAACATCCGGACCAACTTCTGGACCGACTTCTGGACCAACCAGCGGACCAACTTCTG<br>GACCAACTAGCGGACCAACTGGAGGCCCAACCTCTGGACCTACTTCTGGACCAACCAGCGGACCAACATCCGGCCCAACATC<br>TGGACCAACTAGCGGACCTACTTCTGGACCAACATCTGGACCAACATCCGGCCCAACTTCTGGACCAACTAGCGGACCAACA<br>TCCGGCCCAACATCTGGACCAACCAGCGGACCTACTTCTGGACCAACATCTGGACCAACATCCGGCCCAACTTCTGGACCAA<br>CTAGCGGACCAACATCCGGCCCAACATCTGGACCAACCAGCGGACCTACTTCTGGACCAACATCTGGACCAACATCCGGCCC<br>AACTTCTGGACCAACTAGCGGACCAACATCCGGCCCTACATCTGGACCAACCAGCGGACCTACTTCTGGACCAACATCTGGA<br>CCAACATCCGGCCCAACTTCTGGACCAACTAGCGGACCAACCAGCGGACCTACTTCTGGACCAACCTCTGGCCCAACCTCTG<br>GACCAACCAGCCAGCCAACCTACGTCTGTAGCACTGAGGCCCGCCTCGTCGACAAGGACGGTGGCAACACCAAGTACGATGA<br>AGGCCTTGATGACGAGATCACCGGCGACGATGGTATTCCATTCGAGGTGAGCGAGCGTGAAGTCGACAGCGTGAAGATCTCC<br>CTTGAGCCACTCCTTGATGCCATCGAGGCCCTTCGCACCGGCGGTGCTGTAGGCGAGGACGGCGAGCTCGGATCTGTGGCTG<br>TCGTGTTCGACAGAATCAGCACCGACAACGACGGCAACGAAGTGACCGAGAACGACTTCTGCACGGAGACATCCGTGACTGA<br>GGCCACCGGCCCAGCCGAGGAGTACGTGGCACAGTGTGTCGAGGATGATGATTTCGCCGGAACCACTGTTGCGATTGTGTAC<br>GTTGTCGTCGCCGACAGCGCCAACACTCAGGAGGATCCATCCGGTTCCTTGATCGGAGCGGCAGGTGGCGAGAGCGCCCTGG<br>GCACAAGCGGATGCGGCGTTCCATTCGAGAGCGGACCATCCAGCACTCCAGTGATCGTGTACAAGTACGTGGTTCCATGCCA<br>GTACTCTTGCGAAGGATCTCCAACGTCTGTCCCAACCGACGCACCAACCGGAGGCCCAACTTCTGGACCAACCAGCGGACCA<br>ACCTCTGGACCAACTTCTGGCCCAACATCCGGACCAACCTCTGGACCTACTTCTGGTCCAACCAGCGGACCAACCTCTGGAC<br>CAACCAGCGGACCAACAAGCGGACCAACATCCGGCCCAACTTCTGGACCGACTTCTGGACCAACCAGCGGACCAACTTCTGG<br>GCCAACTAGCGGACCAACTGGAGGCCCAACCTCTGGACCTACTTCTGGACCAACCAGCGGACCAACATCCGGCCCAACTTCT<br>GGACCAACATCCGGTCCAACCAGCGGACCAACTGGAGGCCCAACCTCTGGACCTACTTCTGGACCAACCAGCGGACCAACAT<br>CCGGCCCAACTTCTGGACCAACATCCGGTCCAACCAGCGGACCAACTGGAGGCCCAACCTCTGGACCAACTAGCGGACCAAC<br>CAGCGGACCAACATCCGGACCAACTTCCGGACCAACCAGCGGACCTACTTCTGGACCAACATCTGGACCAACATCCGGCCCA<br>ACTTCTGGACCAACTAGCGGACCAACATCCGGCCCAACATCTGGACCAACCAGCGGACCTACTTCTGGACCAACATCTGGAC<br>CAACATCCGGCCCAACTTCTGGACCAACTAGCGGACCAACCAGCGGACCTACTTCTGGACCAACCTCTGGCCCAACCTCTGG<br>ACCAACCAGCCAGCCAACCTACGTCTGTAGCACTGAGGCCCGCCTCGTCGACAAGGACGGTGGCAACACCAAGTACGATGAA<br>GGCCTTGATGACGAGATCACCGGCGACGATGGTATTCCATTCGAGGTGAGCGAGCGTGAAGTCGACAGCGTGAAGATCTCCC<br>TTGAGCCACTCCTTGATGCCATCGAGGCCCTTCGCACCGGCGGTGCTGTAGGCGAGGACGGCGAGCTCGGATCTGTGGCTGT<br>CGTGTTCGACAGAATCAGCACCGACAACGACGGCAACGAAGTGACCGAGAACGACTTCTGCACGGAGACATCCGTGACTGAG<br>GCCACCGGCCCAGCCGAGGAGTACGTGGCACAATGTGTCGAGGATGATGATTTCGCCGGAACCACTGTTGCGATTGTGTACG<br>TTGTCGTCGCCGACAGCGCCAACACTCAGGAGGATCCATCCGGTTCCTTGATCGGAGCGGCAGGTGGCGAGAGCGCCCTGGG |

| | |
|---|---|
| | CACAAGCGGATGCGGCGTTCCATTCGAGAGCGGACCATTCAGCACTCCAGTGATCGTGTACAAGTACGTGGTTCCATGCCAG<br>TACTCTTGCGAAGGATCTCCAACGTCTGTCCCAACCGACGCACCAACCGGAGGCCCAACTTCTGGACCAACCAGCGGACCAA<br>CCTCTGGACCAACTTCTGGCCCAACATCCGGACCAACCTCTGGACCTACTTCTGGTCCAACCAGCGGACCAACCTCTGGACC<br>AACCAGCGGACCAACAAGCGGACCAACATCCGGCCCAACTTCTGGACCGACTTCTGGACCAACCAGCGGACCAACTTCTGGG<br>CCAACTAGCGGACCAACTGGAGGCCCAACCTCTGGACCTACTTCTGGACCAACCAGCGGACCAACATCCGGCCCAACTTCTG<br>GACCAACATCCGGTCCAACCAGCGGACCAACTGGAGGCCCAACCTCTGGACCTACTTCTGGACCAACCAGCGGACCAACATC<br>CGGCCCAACTTCTGGACCAACATCCGGTCCAACCAGCGGACCAACTGGAGGCCCAACCTCTGGACCAACTAGCGGACCAACC<br>AGCGGACCAACATCCGGACCAACTTCCGGACCAACCAGCGGACCTACTTCTGGACCAACCTCTGGCCCAACCTCTGGACCAA<br>CCAGCCAGCCAACCTACGTCTGTAGCACTGAGGCCCGCCTCGTCGACAAGGACGGTGGCAACACCAAGTACGATGAAGGCCT<br>TGATGACGAGATCACCGGCGACGATGGTATTCCATTCGAGGTGAGCGAGCGTGAAGTCGACAGCGTGAAGATCTCCCTTGAG<br>CCACTCCTTGATGCCATCGAGGCCCTTCGCACCGGCGGTGCTGTAGGCGAGGACGGCGAGCTCGGATCTGTGGCTGTCGTGT<br>TCGACAGAATCAGCACCGACAACGACGGCAACGAAGTGACCGAGAACGACTTCTGCACGGAGACATCCGTGACTGAGGCCAC<br>CGGCCCAGCCGAGGAGTACGTGGCACAGTGTGTCGAGGATGATGATTTCGCCGGAACCACTGTTGCGATTGTGTACGTTGTC<br>GTCGCCGACAGCGCCAACACTCAGGAGGATCCATCCGGTTCCTTGATCGGAGCGGCAGGTGGCGAGAGCGCCCTGGGCACAA<br>GCGGATGCGGCGTTCCATTCGAGAGCGGACCATCCAGCACTCCAGTGATCGTGTACAAGTACGTGGTTCCATGCCAGTACTC<br>TTGCGAAGGATCTCCAACGTCTGTCCCAACCGACGCACCAACCGGAGGCCCAACTTCTGGACCAACCAGCGGACCAACCTCT<br>GGACCAACTTCTGGCCCAACATCCGGACCAACCTCTGGACCTACTTCTGGTCCAACCAGCGGACCAACCTCTGGACCAACCA<br>GCGGACCAACAAGCGGACCAACATCCGGCCCAACTTCTGGACCGACTTCTGGACCAACCAGCGGACCAACTTCTGGGCCAAC<br>TAGCGGACCAACTGGAGGCCCAACCTCTGGACCTACTTCTGGACCAACCAGCGGACCAACATCCGGCCCAACTTCTGGACCA<br>ACATCCGGTCCAACCAGCGGACCAACTGGAGGCCCAACCTCTGGACCTACTTCTGGACCAACCAGCGGACCAACATCCGGCC<br>CAACTTCTGGACCAACATCCGGTCCAACCAGCGGACCAACTGGAGGCCCAACCTCTGGACCAACTAGCGGACCAACCAGCGG<br>ACCAACATCCGGACCAACTTCCGGACCAACCAGCGGACCTACTTCTGGACCAACATCTGGACCAACATCCGGCCCAACTTCT<br>GGACCAACTAGCGGACCAACATCCGGCCCAACATCTGGACCAACCAGCGGACCTACTTCTGGACCAACATCTGGACCAACAT<br>CCGGCCCAACTTCTGGACCAACTAGCGGACCAACATCCGGCCCAACATCTGGACCAACCAGCGGACCTACTTCTGGACCAAC<br>ATCTGGACCAACATCCGGCCCAACTTCTGGACCAACTAGCGGACCAACCAGCGGACCTACTTCTGGACCAACCTCTGGCCCA<br>ACCTCTGGACCAACCAGCCAGCCAACCTACGTCTGTAGCACTGAGGCCCGCCTCGTCGACAAGGACGGTGGCAACACCAAGT<br>ACGATGAAGGCCTTGATGACGAGATCACCGGCGACGATGGTATTCCATTCGAGGTGAGCGAGCGTGAAGTCGACAGCGTGAA<br>GATCTCCCTTGAGCCACTCCTTGATGCCATCGAGGCCCTTCGCACCGGCGGTGCTGTAGGCGAGGACGGCGAGCTCGGATCT<br>GTGGCTGTCGTGTTCGACAGAATCAGCACCGACAACGACGGCAACGAAGTGACCGAGAACGACTTCTGCACGGAGACATCCG<br>TGACTGAGGCCACCGGCCCAGCCGAGGAGTACGTGGCACAGTGTGTCGAGGATGATGATTTCGCCGGAACCACTGTTGCGAT<br>TGTGTACGTTGTCGTCGCCGACAGCGCCAACACTCAGGAGGATCCATCCGGTTCCTTGATCGGAGCGGCAGGTGGCGAGAGC<br>GCCCTGGGCACAAGCGGATGCGGCGTTCCATTCGAGAGCGGACCATCCAGCACTCCAGTGATCGTGTACAAGTACGTGGTTC<br>CATGCCAGTACTCTTGCGAAGGATCTCCAACGTCTGTCCCAACCGACGCACCAACCGGAGGCCCAACTTCTGGACCAACCAG<br>CGGACCAACCTCTGGACCAACTTCTGGCCCAACATCCGGACCAACCTCTGGACCTACTTCTGGTCCAACCAGCGGACCAACC<br>TCTGGACCAACCAGCGGACCAACAAGCGGACCAACATCCGGCCCAACTTCTGGACCGACTTCTGGACCAACCAGCGGACCAA<br>CTTCTGGGCCAACTAGCGGACCAACTGGAGGCCCAACCTCTGGACCTACTTCTGGACCAACCAGCGGACCAACATCCGGCCC<br>AACTTCTGGACCAACATCCGGTCCAACCAGCGGACCAACTGGAGGCCCAACCTCTGGACCTACTTCTGGACCAACCAGCGGA<br>CCAACATCCGGCCCAACTTCTGGACCAACATCCGGTCCAACCAGCGGACCAACTGGAGGCCCAACCTCTGGACCAACTAGCG<br>GACCAACCAGCGGACCAACATCCGGACCAACTTCCGGACCAACCAGCGGACCTACTTCTGGACCAACATCTGGACCAACATC<br>CGGCCCAACTTCTGGACCAACTAGCGGACCAACATCCGGCCCAACATCTGGACCAACCAGCGGACCTACTTCTGGACCAACA<br>TCTGGACCAACATCCGGCCCAACTTCTGGACCAACTAGCGGACCAACATCCGGCCCAACATCTGGACCAACCAGCGGACCTA<br>CTTCTGGACCAACATCTGGACCAACATCCGGCCCAACTTCTGGACCAACTAGCGGACCAACCAGCGGACCTACTTCTGGACC<br>AACCTCTGGCCCAACCTCTGGACCAACCAGCCAGCCAACCTACGTCTGTAGCACTGAGGCCCGCCTCGTCGACAAGGACGGT<br>GGCAACACCAAGTACGATGAAGGCCTTGATGACGAGATCACCGGCGACGATGGTATTCCATTCGAGGTGAGCGAGCGTGAAG<br>TCGACAGCGTGAAGATCTCCCTTGAGCCACTCCTTGATGCCATCGAGGCCCTTCGCACCGGCGGTGCTGTAGGCGAGGACGG<br>CGAGCTCGGATCTGTGGCTGTCGTGTTCGACAGAATCAGCACCGACAACGACGGCAACGAAGTGACCGAGAACGACTTCTGC<br>ACGGAGACATCCGTGACTGAGGCCACCGGCCCAGCCGAGGAGTACGTGGCACAGTGTGTCGAGGATGATGATTTCGCCGGAA<br>CCACTGTTGCGATTGTGTACGTTGTCGTCGCCGACAGCGCCAACACTCAGGAGGATCCATCCGGTTCCTTGATCGGAGCGGC<br>AGGTGGCGAGAGCGCCCTGGGCACAAGCGGATGCGGCGTTCCATTCGAGAGCGGACCATCCAGCACTCCAGTGATCGTGTAC<br>AAGTACGTGGTTCCATGCCAGTACTCTTGCGAAGGATCTCCAACGTCTGTCCCAACCGACGCACCAACCGGAGGCCCAACTT<br>CTGGACCAACCAGCGGACCAACCTCTGGACCTACTTCTGGACCAACATCCGGACCAACCTCTGGACCTACTTCTGGTCCAAC<br>CAGCGGACCAACCTCTGGACCAACCAGCGGACCAACAAGCGGACCAACATCCGGCCCAACTTCTGGACCGACTTCTGGACCA<br>ACCAGCGGACCAACTTCTGGGCCAACTAGCGGACCAACTGGAGGCCCAACCTCTGGACCTACTTCTGGACCAACCAGCGGAC<br>CAACATCCGGCCCAACTTCTGGACCAACATCCGGTCCAACCAGCGGACCAACTGGAGGCCCAACCTCTGGACCAACTAGCGG<br>ACCAACCAGCGGACCAACATCCGGCCCAACTTCTGGACCAACATCCGGTCCAACCAGCGGACCAACTGGAGGCCCAACCTCT<br>GGACCAACTAGCGGACCAACCAGCGGACCAACATCCGGACCAACTTCCGGACCAACCAGCGGACCTACTTCTGGACCAACAT<br>CTGGACCAACATCCGGCCCAACTTCTGGACCAACTAGCGGACCAACATCCGGCCCAACATCTGGACCAACCAGCGGACCTAC<br>TTCTGGACCAACATCTGGACCAACATCCGGCCCAACTTCTGGACCAACTAGCGGACCAACCAGCGGACCTACTTCTGGACCA<br>ACATCTGGACCAACCAGCGGACCAACTGGAGGCCCAACCTCTGGACCAACTAGCGGCCCAACCTCTGGACCAACTAGCGGAC<br>CAACATCCGGCCCAACATCTGGACCAACCAGCGGACCTACTTCTGGACCAACATCTGGACCAACATCCGGCCCAACTTCTGG<br>ACCAACTAGCGGACCAACATCCGGCCCAACATCTGGACCAACCAGCGGACCTACTTCTGGACCAACATCTGGACCAACATCT<br>GGACCAACATCCGGCCCAACTTCTGGACCTACTTCTGGACCAACCAGCGGACCTACTTCTGGACCAACATCCGACCCAACTT<br>CTGGACCAACATCCGGCCCAACCTCTGGACCTACTTCTGGACCAACCAGCGGACCAACTTCCGGCCCAACATCTGGACCAAC<br>TGGCGGACCAACTAGTGGACCAACATCAGGACCAACCAGTGGACCAACATCAGGACCAACCAGTGGACCAACGTCTGGACCT<br>ACAAGCGGACCAACCAGCGGACCAACCAGCGGCCCTACATCTGGCCCAACATCTGGCCCAACATCTGGTCCAACTTCTGGAC<br>CAACCAGCGGACCAACCAGCGGGCCTACAAGTGGACCTACAAGTGGACCTACAAGTGGACCTACAAGTGGACCTACAAGTGG<br>ACCTACAAGTGGACCTACAAGTGGACCTACAAGTGGACCTACAAGTGGACCTACAAGTGGACCTACAAGTGGACCTACAAGT<br>GGACCTACAAGTGGACCTACAAGTGGACCTACAAGTGGACCCACAAGTGGGCCTACGTCCGGACCAACATTGGCGCCCACCA<br>GTAACGTTCCAAGTTCTGCCCCATCGAGAAACTGTGAATATGTTGTGGCCGACTTTGATGGGGACGATGTGAAGACTAATGC |

| | |
|---|---|
| | CTATCTTTTCCCAACAGCACTCATTGAATCTGGTATCTCGATCGAGGCATTCGGTTCTAATGGCACTGGGCACACCCCAAGC<br>GATATGGCCAGGGTCATGGAGACCCCCACGGAGGGGAATGCCATCTTCATTCAAGAATCGAATGTCAGAGATATCATTCCAA<br>ACGCTGAAGGAGGAGTCTTGTCGTTCACTTTCCAAGCCACTGTGGAAGAAGTGATTGGGCTGCAGCTTTGGGATGTTTCGGA<br>GGGCGGAATTGTTACCGTTCTCACAGAAGGTGGCACCATCGAGTCCTTCCCAATCATGGCATCTCCGAATGCCACCCAGACG<br>ATTGATATTGGTGTCTTTGAGGCTCTCGAAATGAACGTTACCTTTGGCGGGACATTGTCTGTGACGGCCATCGAAATTTGTT<br>TTGACGGTTCCACGACTCCTGGTCCAGTACGATTCGCGCCTCCATCCGAGACAAAGAGTCCCACTGCCATGCCAACAACAAG<br>CCCCGCACCAACGGGCGAGCCCACAGAAAGTTCAGCACCAACTGACTGTTACGATCAACTCCGTGCGAAACTTGTCGACCAG<br>GTAGGAACGGATTCTACTCCTGTACCAGAGGATGAAGAACTGATCGTCGTTACGTCCCAGAACGGCACACACGTTGGCTTTG<br>AAGTCAACCAGGTGTGGGGATCAGGGGCAAACATCAGCCTGATTGCTGTGAATCACCATGTGAACTTTACATACACCGAATG<br>CAGTCGCCAATACGATGTGACACCAGATTCATCGTCGTTGTATTTGGCGAAGTGTTTCGATGAGTGGGCTGATGTCACTGTG<br>TACATCTATACAGCAGACGACTTTTCGGAGGAAGAATGTGACATCTGCGAAACGCCTGAAGAAGATGATGAAGACATCATGG<br>TATTCCACTTCGAGATTCCATGCAACTCCGAGTGCGATCTCTCCGCCTCTCCGACTGGAGGACCAACCAGCGGACCAACCTC<br>CGGCCCAACTTCTGGACCAACCTCTGTACCCACCTCTGAACCAACTAGCGGACCAACCAGTGGACCAACCAGCGGACCAACA<br>TCCGGACCAACATCCGGCCCAACCAGCGGCCCAACTTCTGGACCGACTTCCGGACCCACCTCTGGACCAACCAGTGGACCAA<br>CCAGCGGACCAACCAGCGGACCAACCAGCGGACCAACCAGCGGACCAACATCTGGACCCACCTCCGGCCCAACATCTGGACC<br>AACCAGTGGACCAACCAGTGGACCAACCAGCGGACCAACCAGCGGACCAACTTCTGGACCAACCTCCGGCCCAACATCTGGA<br>CCAACCAGTGGACCAACATCTGGACCAACGTCCGGCCCAACATCTGGACCAACCAGCGGACCAACCAGCGGACCAACCTCCG<br>GCCCAACATCTGGACCAACCAGTGGACCAACCAGTGGACCAACTTCTGGACCTACCTCCGGCCCAACTTCTGGACCTACCTC<br>CGGCCCAACATCCGGCCCAACATCCGGACCAACCTCTGGACCAACCTCTGGACCAACCAGCGGACCAACCAGCGGACCAACC<br>AGCGGACCAACCAGCGGACCAACCTCCGGCCCAACATCTGGACCAACATCTGGACCAACGTCCGGCCCAACATCTGGACCAA<br>CCAGCGGACCAACTTCTGGACCTACCTCCGGCCCAACATCCGGACCAACCTCTGGACCAACCAGCGGACCAACATCTGGACC<br>TACCTCCGGACCAACCAGCGGACCAACCAGCGGACCAACTTCTGGACCAACCTCCGGCCCAACATCTGGACCAACCAGTGGA<br>CCAACTTCTGGACCAACCTCTGGACCAACCTCCGGCCCAACTTCTGGACCAACTAGCGGACCAACCAGCGGACCAACCAGCG<br>GACCAACCAGCGGACCAACATCTGGACCAACTAGCGGACCAACTAGCGGACCAACCAGCGGACCAACTTCTGGACCAACCTC<br>TGGACCAACCTCCGGCCCAACTTCTGGACCAACTAGCGGACCAACCAGCGGACCAACCAGCGGACCAACCAGCGGACCAACA<br>TCTGGACCAACTAGCGGACCAACTAGCGGACCAACCAGCGGACCAACTTCTGGACCTACCTCCGGACCAACATCTGGACCAA<br>CATCTGGACCAACTTCTGGACCAACCAGTGGACCAACCTCCGGACCAACCAGTGGACCAACCAGCGGACCAACCAGCGGACC<br>AACTTCTGGACCCACCTCCGGCCCAACATCTGGACCAACCAGTGGACCAACTTCTGGACCTACCTCCGGCCCAACATCTGGA<br>CCAACCTCCGGACCAACCTCCGGACCAACCTCCGGACCAACCTCCGGACCAACCTCCGGACCAACCAGTGGACCAACCTCCG<br>GACCAACCTCCGGACCAACCAGTGGACCAACCAGTGGACCAACCAGTGGACCAACCAGTGGACCAACCAGCGGCCCAACTTC<br>TGGACCTACCTCCGGCCCAACATCCGGACCAACCTCTGGACCAACCTCTGGACCAACATCCGGACCAACCAGCGGACCAACA<br>TCTGGACCAACTTCTGGACCAACCAGCGGACCAACCAGTGGACCAACCAGTGGACCAACCAGTGGACCAACCAGCGGACCAA<br>CCAGCGGACCAACCAGCGGACCAACCAGCGGACCAACCAGCGGACCAACCAGCGGACCAACCAGCGGACCAACCTCTGGACC<br>TACTTCAGGCCCAACCTCTGGACCAACTAGTGGACCAACTAGTGGACCAACATCCGGACCAACATCCGGACCAACATCCGGA<br>CCAACATCCGGCCCAACATCTGGACCAACTAGTGGACCAACATCCGGACCAACATCCGGCCCAACATCTGGACCAACATCCG<br>GACCAACCTCCGGACCAACCTCCGGCCCAACATCTGGACCAACCAGCGGACCAACCAGCGGACCAACTTCTGGACCCACCTC<br>CGGACCAACATCCGGACCAACCAGTGGACCAACTAGCGGACCAACCAGCGGACCAACCAGCGGACCAACTTCTGGACCTACC<br>TCCGGCCCAACATCCGGACCAACTTCTGGACCTACCTCCGGACCAACCTCTGGACCAACCTCCGGCCCAACATCCGGACCAA<br>CCAGCGGACCAACATCTGGACCAACATCTGGACCAACCTCCGGACCAACCAGCGGACCAACATCCGGACCAACATCTGGACC<br>AACTAGTGGACCAACCAGCGGACCAACATCTGGACCAACCTCCGGCCCAACATCTGGACCAACCAGTGGACCAACCAGCGGC<br>CCAACTTCTGGACCTACCTCCGGCCCAACATCCGGACCAACCTCTGGACCAACCTCCGGACCAACATCCGGACCAACCTCCG<br>GACCAACTAGTGGACCAACATCCGGACCAACATCTGGACCAACATCTGGACCAACCAGTGGACCAACCAGTGGACCAACCAG<br>CGGCCCAACTTCTGGACCAACTAGCGGACCAACATCTGGACCAACCTCCGGACCAACCTCCGGACCAACCAGCGGACCAACC<br>TCCGGACCAACATCTGGACCAACTAGTGGACCAACCAGCGGACCAACATCTGGACCAACCTCCGGCCCAACATCTGCACCAA<br>CTAGCGGACCAACTAGCGGACCAACCTCTGGACCAACATCCGGACCAACCTCCGGACCAACATCTGGACCAACCTCCGGACC<br>AACCTCTGGACCAACCAGCGGACCAACCTCCGGACCAACCAGTGGACCAACATCCGGACCAACTAGTGGACCAACCTCTGGA<br>CCAACCAGCGGACCAACCTCCGGACCAACCAGTGGACCAACATCCGGACCAACTAGTGGACCAACATCCGGACCAACTAGTG<br>GACCAACCAGCGGACCAACCTCCGGACCAACCTCCGGACCAACCTCCGGACCAACCAGTGGACCAACCTCCGGACCAACCTC<br>CGGACCAACCAGTGGACCAACCAGTGGACCAACCAGTGGACCAACCTCCGGACCAACCTCCGGACCAACCAGTGGACCAACC<br>AGTGGACCAACATCCGGACCAACATCCGGACCAACCTCCGGACCAACATCTGGACCAACTAGCGGACCAACCTCTGGACCAA<br>CCAGCGGACCAACATCCGGACCAACATCCGGACCAACCAGTGGACCAACATCCGGACCAACATCCGGACCAACATCCGGACC<br>AACCTCCGGACCAACATCTGGACCAACTAGCGGACCAACCTCTGGACCAACCAGCGGACCAACCTCCGGACCAACCAGTGGA<br>CCAACATCCGGACCAACTAGTGGACCAACCAGCGGACCAACATCCGGACCAACCTCCGGACCAACATCTGGACCCACCTCTG<br>GACCAACATCCGGACCAACCAGCGGACCAACTAGCGGACCTACCTCCGGACCAACAAGTGGACCAACTTCGGGGCCAACGTC<br>GGGACCGACATCTGGACCAACCTCAGGACCAACCTCTGGACCTACTTCTGGACCAACCAGCGGACCTACTTCCGGACCAACA<br>ACCAGCCCAGCACCTTCCGAGTGCATCGACCAGGTCACACCAAAACCAACGGGAACATACGGTGCTGACGTCCCTCTGCCAT<br>TGGACAGCGACCTCATTGTCATCACCGGTCTGAACGGCACCCATGTGTCGTTCGCTGTGAACCCAGCCATCGGCGAAGACAC<br>CGACTCCAACATCACTGCCGTGTCCATCGAGTACAGACCTAACGTGACCCATTCTGAGTGCAAGACCGAGTTCGATGTCGTC<br>CCTGAGGCGGCGTTCGACTTGGAAGCGGTGTGCTTCGATGGTTGGACAATGGTGACAGTCTACATCTACTTTGGAGAACCCT<br>TGGAGGATCCGGAGCAGTGCGAATACTGTGAGACACCTGATGAAGACAGTGACGACTCTGTTGCGTACGTGTTCGAGTTGCC<br>GTGCGAGACCGACTGTTATGACACTATCGCGCCAACATCGATGCCAACGTCGGGACCTACCAGTGGTCCAACCAGCGGCCCA<br>ACTGAGACCATTTGCATCTACACAGTTCTTGACTTCGCCGCTGACGGCGACAACCAGCCAACTGGCAATGGCTATGTTCCCT<br>CTGAGATATACGATATATTCGACATTGACTCGATCACCGTTGATCTTGGCGATAGTCCTTCGTCGTACGGCGGGCCCACTGG<br>GAAGAACGCAAGATACTGGGTCACTGGTGACTCAGAGGCAGGCACAGCTATCATCATCCAAGAGAGCAATATGAACGACGCT<br>GTTCCGAGCCCGACTGGTGGTGATATTGTGTTCACCTTCAACGGGCAATCAACCGAAACAATTGAACTTACTTTCTACGATC<br>TTGCCGCGGATGTGACAATCACAGCGAGAACTACGGCCGATCCATCCGAGGGTGGACTCGCACAGACCACGACAGCCAGCGC<br>CAACCCAGGTGGATTTACAAAGGTTGAGATTGACGTTTCAATGATGAATCAAGTCATCGTGACGTTCACAGCTCCGGGTGCC<br>ATCGCATCTATTGGAGTATGCCGTGACCCAAATACGCCACCCCCGCAAGGAGACAACAGCCCGTCAGCAACACTGGAACCGA<br>CGTCACAGCCAACGACAAGCCCAGCGCCGACAGGCGAGCCAACCACCAGTGCTGCTCCAACCGATTGCTACGACCAACTCCG |

| | |
|---|---|
| | CGCCAAGTTCGTTCGCCAGGAAGGTGCAAGCGTTTCCCTTCCTGAAGACAACGATTTGATCGTTGTCACTAGTCAGAACGGA<br>ACGCACGTGTCTTTCGAAGTCAATCCATCCATCGGCGAAGATACCGACTCCAACATCACGACTGTGTCAATCCAATACAGAG<br>CGAATGTTACCTACTCTGAGTGCAAGACCGAGTTCGATGTCGCCCCTGAGGCGGCGTTCGACTTGGAAGCGGTGTGCTTCGA<br>TGGTTGGACTATGGTGACAGTCTACATCTATTTTGGCGAGCCTTTGGAGGATCCGGAGCAGTGCGATTTCTGCGAAACTCCT<br>GATGAGGACAGTGACGACTCAGTCGCATATGTCTTCGAGGTGCCATGCACAGCAAACTGTGATTTGTCAGCCTCACCAACCA<br>GCGGGCCAACCTCTGGACCAACTTCTGGACCTACTTCTGGACCAACCTCCGGCCCAACTTCTGGACCTACAAGCGGACCAAC<br>ATCTGGACCAACATCCGGACCTACGAGCGGACCAACATCCGGACCAACCAGTGGACCAACATCTGGACCAACCAGCGGACCT<br>ACTTCTGGACCAACTAGTGGACCAACTAGCGGACCAACCTCCGGCCCAACTTCTGGACCTACAAGCGGACCAACTTCTGGAC<br>CCACCTCTGGACCAACTAGCGGACCTACAAGCGGACCAACTAGCGGACCAACTTCGGGGCCTACATCTGGACCAACCAGCGG<br>ACCCACCTCTGGACCAACATCCGGACCAACCAGCGGACCAACATCCGGACCAACCAGCGGACCAACCTCCGGACCAACATCT<br>GGACCAACTAGTGGACCAACCAGCGGACCAACATCCGGACCAACCAGCGGACCAACATCTGGACCAACCAGCGGACCAACAT<br>CTGGACCAACCAGCGGACCAACATCCGGACCAACCAGTGGACCAACCAGTGGACCAACATCTGGACCAACTTCCGGACCAAC<br>TAGTGGACCAACCAGCGGACCAACCAGCGGACCAACATCCGGACCAACTAGCGGACCAACCTCTGGACCAACTAGTGGACCA<br>ACCAGCGGACCAACCTCCGGACCAACATCTGGACCAACTAGTGGACCAACCTCCGGACCAACATCTGGACCAACCAGTGGAC<br>CAACATCCGGACCAACCAGCGGACCAACATCCGGCCCAACTTCTGGACCAACTAGCGGACCAACATCTGGACCAACCTCCGG<br>GCCAACCTCCGGACCAACCTCCGGCCCAACATCTGGACCAACCAGCGGACCAACCAGTGGACCAACCAGTGGACCAACCAGT<br>GGACCAACCTCCGGACCAACCTCCGGACCAACCTCCGGACCAACCAGTGGACCAACCTCCGGACCAACCAGTGGACCAACAT<br>CCGGACCAACATCCGGACCAACCTCCGGACCAACATCCGGACCAACTAGCGGACCAACCTCTGGACCAACCAGCGGACCAAC<br>CTCCGGACCAACCAGTGGACCAACCAGTGGACCAACCTCCGGACCAACCAGTGGACCAACATCCGGACCAACCAGTGGACCA<br>ACATCCGGACCAACATCCGGACCAACCTCCGGACCAACATCCGGACCAACTAGCGGACCAACCTCTGGACCAACCAGCGGAC<br>CAACCTCCGGACCAACCTCCGGACCAACATCCGGACCAACTAGTGGACCAACATCCGGACCAACTAGTGGACCAACCAGCGG<br>ACCAACCAGCGGACCAACCAGCGGACCAACCTCCGGACCAACCTCCGGACCAACCAGTGGACCAACCAGTGGACCAACCAGT<br>GGACCAACCAGTGGACCAACCTCCGGACCAACCTCCGGACCAACCTCCGGACCAACCTCCGGACCAACCAGTGGACCAACCA<br>GTGGACCAACCAGTGGACCAACCAGTGGACCAACCAGTGGACCAACCTCCGGACCAACCAGTGGACCAACATCCGGACCAAC<br>CAGTGGACCAACATCCGGACCAACTAGTGGACCAACATCCGGACCAACTAGTGGACCAACCAGCGGACCAACCAGCGGACCA<br>ACCAGCGGACCAACCTCCGGACCAACCTCCGGACCAACCTCCGGACCAACCTCCGGACCAACCTCCGGACCAACCTCCGGAC<br>CAACCTCCGGACCAACCTCCGGACCAACCAGTGGACCAACCTCCGGACCAACCAGTGGACCAACATCCGGACCAACATCCGG<br>ACCAACATCCGGACCAACCTCCGGACCAACATCCGGACCAACCTCCGGACCAACATCTGGACCAACCAGCGGACCAACCTCC<br>GGACCAACCAGTGGACCAACATCCGGACCAACTAGTGGACCAACCAGCGGACCAACATCCGGACCAACCTCCGGACCAACAT<br>CTGGACCCACCTCTGGACCAACATCCGGACCAACCAGCGGACCAACTAGCGGACCTACCTCCGGACCAACAAGTGGACCAAC<br>TTCGGGGCCAACGTCGGGACCGACATCTGGACCAACCTCAGGACCAACCTCTGGACCTACTTCTGGACCAACCAGCGGACCT<br>ACTTCCGGACCAACAACCAGCCCAGCACCTTCCGAGTGCATCGACCAGGTCACACCAAAACCAACGGGAACATACGGCGCTG<br>ACGTCCCTCTGCCATTGGACAGCGACCTCATTGTCATCACCGGTCTGAACGGCACCCATGTGTCGTTCGCTGTGAACCCAGC<br>CATCGGCGAAGACACCGACTCCAACATCACTGCCGTGTCCATCGAGTACAGACCTAACGTGACCCATTCTGAGTGCAAGACC<br>GAGTTCGATGTCGTCCCTGAGGCGGCGTTCGACTTGGAAGCGGTGTGCTTCGATAGTTGGACAATGGTGACAGTCTACATCT<br>ACTTTGGAGAACCCTTGGAGGATCCGGAGCAGTGCGAATACTGTGAGACACCTGATGAAGACAGTGACGACTCTGTTGCGTA<br>CGTGTTCGAGTTGCCGTGCGAGACCGACTGTTATGACACTATCGCGCCAACATCGATGCCAACGTCGGGACCTACCAGTGGT<br>CCAACTGCAAGCCCTGCGCCAACCGACTGCTACGAACAACTCTTGCCAAAACTTACGCACCAATCCGGCGCAGACGTGAATA<br>TTCCAGAGAACGACGATCTCATCGTCATCACCAGCCAAAATGGAACGCACGTGGGCATTGAAGTTCGACAAGTTATTGAGAA<br>CTCGGAAGACGGAGCCTCGATGCTTGCTGTGCAATACAGAAACTCGACAGCCGAATCCTGTGAGAAGAGCTCGGGAGTGACG<br>TTTGGAACAACTGTGTCCTTCATCGCCACCTGCTTCGACTCCTACGCGGAAGTGCGAGTCTTCCTTGTGTTGGGTAATGCCA<br>CTGAGGACGCCGAGGATTGCGACGCATGCAGGTCACCGGACGAAGACGACACAGACAAGGTGGTCTGGGTCTTTGAAGTCCC<br>ATGTGAGTCTGGATGTGATCTTTCGGATTCCCCGAGCCAAAGCCCAACAAGTTCTCCGACAACCAGCCCAGCACCTTCCGAG<br>TGCATCGACCAGGTCACACCAAAACCAACGGGAACATACGGTGCTGACGTCCCTCTGCCATTGGACAGCGACCTCATTGTCA<br>TCACCGGTCTGAACGGCACCCATGTGTCGTTCGCTGTGAACCCAGCCATCGGCGAAGACACCGACTCCAACATCACTGCCGT<br>GTCCATCGAGTACAGACCTAACGTGACCCATTCTGAGTGCAAGACCGAGTTCGATGTCGCCCCTGAGGCGGCGTTCGACTTG<br>GAAGCGGTGTGCTTCGATGGTTGGACAATGGTGACAGTCTACATCTACTTTGGAGAACCCTTGGAGGATCCGGAGCAGTGCG<br>AATACTGTGAGACACCTGATGAAGACAGTGACGACTCTGTTGCGTACGTGTTCGAGTTGCCGTGCGAGACCGACTGTTATGA<br>CACTATCGCGCCAACATCAGCACCAACCAATTGCTACGATCGCGTCAACGTCGCTCCGGTCAGAAAGGCTGGAGACGACATG<br>GCTATTCCGGAGGGTGCAGTGACCATTGTGGAGCATGATGGCGCCACTGTCAAGTTCAAGGTCACACAATTGTGGAAAGATG<br>TGGACGTGAAGATGTTGGCGTTCCAATATCGCGAAACCGCATCTGCGGAGAAGTGCTTCCCGGACACAACCGTTGACTTTGG<br>AACAACGTATGAACATACCGCTGTCTGCTTCGACGGTTTCGCGTCCGTTGGTATCTATCTGTACACCGGCGATGATTTCGAT<br>CCAACGACATGTGAAGCATGTAAACAGCCAACGGACATCGACGATGGCGTCGCACTGTTCTACTTCGAGATGCCATGCAACT<br>CTGAATGCGCACCACCAACCCCGACATCAGAGACAGAGGCATCGAACTGCTACGACGATGTGGTTCTGGATGACTCTTCTGG<br>AACGTGTGTGTATGATGCGAGCCCTGTGTCAATCGTATCGCAGCACGGCAGCACAGTCAAGTTCTCTGTGTCTCACACGTGG<br>GACATAGCACAGACTCATGCCTTGGGTGTCGTCCCAGCTGCGCTGAGCCAGATGTCGATCCGGTACAAGGCCCAGGGAGAAA<br>ATTCCATGGGTTGCACAACGAGACTCGAACCAGCCGCCGGATATTTGGACACATACACAGCTCAGTGCAACAGCAACGGGCT<br>TGCACAGGTCGAGATCTTCGTGCATGACAGTACCTTCCCCAAGGACAGCAGCGCGGAGATCCCGGACGAATGCGACCAACCC<br>ACCGATCCTGACCATACTTGCGGATACTCGTTTGTCATCCCATGTAACGCGGACATGATGTGCGACAGCAACCGCCGAATGC<br>AGAAGATCCGCAGCCACATGAAGGAAGTGAAGGAAGAGTCTTACGCGCCAGGCCTGAGCTCAAGCATTCTCATGGAATCGTT<br>CAAGAAGACCTCCTTGGAAACAGATTTGCCGGCTGAAGGTGATGAAGATCCATACTGCGCTGCCGAAGACTTCCCGTGTGAG<br>GGCGACTCGCCCAACATGGTGTACGTCTGCCATTACTCTGTCACGAAGGGATACCAAACTTTCTGCGTGCCAGAGGCAGACT<br>CCGACATCCTTCGTTTGTACTCTTCGGATTACTGCGGAAAGTGCGATGGCCAGTAA**XXXXXXXXXXXXXXXXXXXXXXXXXX**<br>**XXXXXX**TTCGAGGTGAGCGAGCGTGAAGTCGACAGCGTGAAGATCTCCCTTGAGCCACTCCTTGATGCCATCGAGGCCCTTC<br>GCACCGGCGGTGCTGTAGGCGAGGACGGCGAGCTCGGATCTGTGGCTGTCGTGTTCGACAGAATCAGCACCGACAACGACGG<br>CAACGAAGTGACCGAGAACGACTTCTGCACGGAGACATCCGTGACTGAGGCCACCGGCCCAGCCGAGGAGTACGTGGCACAG<br>TGTGTCGAGGATGATGATTTCGCCGGAACCACTGTTGCGATTGTGTACGTTGTCGTCGCCGACAGCGCCAACACTCAGGAGG<br>ATCCATCCGGTTCCTTGATCGGAGCGGCAGGTGGCGAGAGCGCCCTGGGCACAAGCGGATGCGGCGTTCCATTCGAGAGCGG<br>ACCATCCAGCACTCCAGTGATCGTGTACAAGTACGTGGTTCCATGCCAGTACTCTTGCGAAGGATCTCCAACGTCTGTCCCA |

| | |
|---|---|
| | ACCGACGCACCAACCGGAGGCCCAACTTCTGGACCAACCAGCGGACCAACCTCTGGACCAACTTCTGGCCCAACATCCGGAC<br>CAACCTCTGGACCTACTTCTGGTCCAACCAGCGGACCAACCTCTGGACCAACCAGCGGACCAACAAGCGGACCAACATCCGG<br>CCCAACTTCTGGACCGACTTCTGGACCAACCAGCGGACCAACTTCTGGGCCAACTAGCGGACCAACTGGAGGCCCAACCTCT<br>GGACCTACTTCTGGACCAACCAGCGGACCAACATCCGGCCCAACTTCTGGACCAACATCCGGTCCAACCAGCGGACCAACTG<br>GAGGCCCAACCTCTGGACCTACTTCTGGACCAACCAGCGGACCAACATCCGGCCCAACTTCTGGACCAACATCCGGTCCAAC<br>CAGCGGACCAACTGGAGGCCCAACCTCTGGACCAACTAGCGGACCAACCAGCGGACCAACATCCGGACCAACTTCCGGACCA<br>ACCAGCGGACCTACTTCTGGACCAACATCTGGACCAACATCCGGCCCAACTTCTGGACCAACTAGCGGACCAACATCCGGCC<br>CAACATCTGGACCAACCAGCGGACCTACTTCTGGACCAACATCTGGACCAACATCCGGCCCAACTTCTGGACCAACTAGCGG<br>ACCAACCAGCGGACCTACTTCTGGACCAACCTCTGGCCCAACCTCTGGACCAACCAGCCAGCCAACCTACGTCTGTAGCACT<br>GAGGCCCGCCTCGTCGACAAGGACGGTGGCAACACCAAGTACGATGAAGGCCTTGATGACGAGATCACCGGCGACGATGGTA<br>TTCCATTCGAGGTGAGCGAGCGTGAAGTCGACAGCGTGAAGATCTCCCTTGAGCCACTCCTTGATGCCATCGAGGCCCTTCG<br>CACCGGCGGTGCTGTAGGCGAGGACGGCGAGCTCGGATCTGTGGCTGTCGTGTTCGACAGAATCAGCACCGACAACGACGGC<br>AACGAAGTGACCGAGAACGACTTCTGCACGGAGACATCCGTGACTGAGGCCACCGGCCCAGCCGAGGAGTACGTGGCACAAT<br>GTGTCGAGGATGATGATTTCGCCGGAACCACTGTTGCGATTGTGTACGTTGTCGTCGCCGACAGCGCCAACACTCAGGAGGA<br>TCCATCCGGTTCCTTGATCGGAGCGGCAGGTGGCGAGAGCGCCCTGGGCACAAGCGGATGCGGCGTTCCATTCGAGAGCGGA<br>CCATTCAGCACTCCAGTGATCGTGTACAAGTACGTGGTTCCATGCCAGTACTCTTGCGAAGGATCTCCAACGTCTGTCCCAA<br>CCGACGCACCAACCGGAGGCCCAACTTCTGGACCAACCAGCGGACCAACCTCTGGACCAACTTCTGGCCCAACATCCGGACC<br>AACCTCTGGACCTACTTCTGGTCCAACCAGCGGACCAACCTCTGGACCAACCAGCGGACCAACAAGCGGACCAACATCCGGC<br>CCAACTTCTGGACCGACTTCTGGACCAACCAGCGGACCAACTTCTGGGCCAACTAGCGGACCAACTGGAGGCCCAACCTCTG<br>GACCTACTTCTGGACCAACCAGCGGACCAACATCCGGCCCAACTTCTGGACCAACATCCGGTCCAACCAGCGGACCAACTGG<br>AGGCCCAACCTCTGGACCTACTTCTGGACCAACCAGCGGACCAACATCCGGCCCAACTTCTGGACCAACATCCGGTCCAACC<br>AGCGGACCAACTGGAGGCCCAACCTCTGGACCAACTAGCGGACCAACCAGCGGACCAACATCCGGACCAACTTCCGGACCAA<br>CCAGCGGACCTACTTCTGGACCAACCTCTGGCCCAACCTCTGGACCAACCAGCCAGCCAACCTACGTCTGTAGCACTGAGGC<br>CCGCCTCGTCGACAAGGACGGTGGCAACACCAAGTACGATGAAGGCCTTGATGACGAGATCACCGGCGACGATGGTATTCCA<br>TTCGAGGTGAGCGAGCGTGAAGTCGACAGCGTGAAGATCTCCCTTGAGCCACTCCTTGATGCCATCGAGGCCCTTCGCACCG<br>GCGGTGCTGTAGGCGAGGACGGCGAGCTCGGATCTGTGGCTGTCGTGTTCGACAGAATCAGCACCGACAACGACGGCAACGA<br>AGTGACCGAGAACGACTTCTGCACGGAGACATCCGTGACTGAGGCCACCGGCCCAGCCGAGGAGTACGTGGCACAGTGTGTC<br>GAGGATGATGATTTCGCCGGAACCACTGTTGCGATTGTGTACGTTGTCGTCGCCGACAGCGCCAACACTCAGGAGGATCCAT<br>CCGGTTCCTTGATCGGAGCGGCAGGTGGCGAGAGCGCCCTGGGCACAAGCGGATGCGGCGTTCCATTCGAGAGCGGACCATC<br>CAGCACTCCAGTGATCGTGTACAAGTACGTGGTTCCATGCCAGTACTCTTGCGAAGGATCTCCAACGTCTGTCCCAACCGAC<br>GCACCAACCGGAGGCCCAACTTCTGGACCAACCAGCGGACCAACCTCTGGACCAACTTCTGGCCCAACATCCGGACCAACCT<br>CTGGACCTACTTCTGGTCCAACCAGCGGACCAACCTCTGGACCAACCAGCGGACCAACAAGCGGACCAACATCCGGCCCAAC<br>TTCTGGACCGACTTCTGGACCAACCAGCGGACCAACTTCTGGGCCAACTAGCGGACCAACTGGAGGCCCAACCTCTGGACCT<br>ACTTCTGGACCAACCAGCGGACCAACATCCGGCCCAACTTCTGGACCAACATCCGGTCCAACCAGCGGACCAACTGGAGGCC<br>CAACCTCTGGACCTACTTCTGGACCAACCAGCGGACCAACATCCGGCCCAACTTCTGGACCAACATCCGGTCCAACCAGCGG<br>ACCAACTGGAGGCCCAACCTCTGGACCAACTAGCGGACCAACCAGCGGACCAACATCCGGACCAACTTCCGGACCAACCAGC<br>GGACCTACTTCTGGACCAACATCTGGACCAACATCCGGCCCAACTTCTGGACCAACTAGCGGACCAACATCCGGCCCAACAT<br>CTGGACCAACCAGCGGACCTACTTCTGGACCAACATCTGGACCAACATCCGGCCCAACTTCTGGACCAACTAGCGGACCAAC<br>ATCCGGCCCAACATCTGGACCAACCAGCGGACCTACTTCTGGACCAACATCTGGACCAACATCCGGCCCAACTTCTGGACCA<br>ACTAGCGGACCAACCAGCGGACCTACTTCTGGACCAACCTCTGGCCCAACCTCTGGACCAACCAGCCAGCCAACCTACGTCT<br>GTAGCACTGAGGCCCGCCTCGTCGACAAGGACGGTGGCAACACCAAGTACGATGAAGGCCTTGATGACGAGATCACCGGCGA<br>CGATGGTATTCCATTCGAGGTGAGCGAGCGTGAAGTCGACAGCGTGAAGATCTCCCTTGAGCCACTCCTTGATGCCATCGAG<br>GCCCTTCGCACCGGCGGTGCTGTAGGCGAGGACGGCGAGCTCGGATCTGTGGCTGTCGTGTTCGACAGAATCAGCACCGACA<br>ACGACGGCAACGAAGTGACCGAGAACGACTTCTGCACGGAGACATCCGTGACTGAGGCCACCGGCCCAGCCGAGGAGTACGT<br>GGCACAGTGTGTCGAGGATGATGATTTCGCCGGAACCACTGTTGCGATTGTGTACGTTGTCGTCGCCGACAGCGCCAACACT<br>CAGGAGGATCCATCCGGTTCCTTGATCGGAGCGGCAGGTGGCGAGAGCGCCCTGGGCACAAGCGGATGCGGCGTTCCATTCG<br>AGAGCGGACCATCCAGCACTCCAGTGATCGTGTACAAGTACGTGGTTCCATGCCAGTACTCTTGCGAAGGATCTCCAACGTC<br>TGTCCCAACCGACGCACCAACCGGAGGCCCAACTTCTGGACCAACCAGCGGACCAACCTCTGGACCAACTTCTGGCCCAACA<br>TCCGGACCAACCTCTGGACCTACTTCTGGTCCAACCAGCGGACCAACCTCTGGACCAACCAGCGGACCAACAAGCGGACCAA<br>CATCCGGCCCAACTTCTGGACCGACTTCTGGACCAACCAGCGGACCAACTTCTGGGCCAACTAGCGGACCAACTGGAGGCCC<br>AACCTCTGGACCTACTTCTGGACCAACCAGCGGACCAACATCCGGCCCAACTTCTGGACCAACATCCGGTCCAACCAGCGGA<br>CCAACTGGAGGCCCAACCTCTGGACCTACTTCTGGACCAACCAGCGGACCAACATCCGGCCCAACTTCTGGACCAACATCCG<br>GTCCAACCAGCGGACCAACTGGAGGCCCAACCTCTGGACCAACTAGCGGACCAACCAGCGGACCAACATCCGGACCAACTTC<br>CGGACCAACCAGCGGACCTACTTCTGGACCAACATCTGGACCAACATCCGGCCCAACTTCTGGACCAACTAGCGGACCAACA<br>TCCGGCCCAACATCTGGACCAACCAGCGGACCTACTTCTGGACCAACATCTGGACCAACATCCGGCCCAACTTCTGGACCAA<br>CTAGCGGACCAACATCCGGCCCAACATCTGGACCAACCAGCGGACCTACTTCTGGACCAACATCTGGACCAACATCCGGCCC<br>AACTTCTGGACCAACTAGCGGACCAACCAGCGGACCTACTTCTGGACCAACCTCTGGCCCAACCTCTGGACCAACCAGCCAG<br>CCAACCTACGTCTGTAGCACTGAGGCCCGCCTCGTCGACAAGGACGGTGGCAACACCAAGTACGATGAAGGCCTTGATGACG<br>AGATCACCGGCGACGATGGTATTCCATTCGAGGTGAGCGAGCGTGAAGTCGACAGCGTGAAGATCTCCCTTGAGCCACTCCT<br>TGATGCCATCGAGGCCCTTCGCACCGGCGGTGCTGTAGGCGAGGACGGCGAGCTCGGATCTGTGGCTGTCGTGTTCGACAGA<br>ATCAGCACCGACAACGACGGCAACGAAGTGACCGAGAACGACTTCTGCACGGAGACATCCGTGACTGAGGCCACCGGCCCAG<br>CCGAGGAGTACGTGGCACAGTGTGTCGAGGATGATGATTTCGCCGGAACCACTGTTGCGATTGTGTACGTTGTCGTCGCCGA<br>CAGCGCCAACACTCAGGAGGATCCATCCGGTTCCTTGATCGGAGCGGCAGGTGGCGAGAGCGCCCTGGGCACAAGCGGATGC<br>GGCGTTCCATTCGAGAGCGGACCATCCAGCACTCCAGTGATCGTGTACAAGTACGTGGTTCCATGCCAGTACTCTTGCGAAG<br>GATCTCCAACGTCTGTCCCAACCGACGCACCAACCGGAGGCCCAACTTCTGGACCAACCAGCGGACCAACCTCTGGACCTAC<br>TTCTGGACCAACATCCGGACCAACCTCTGGACCTACTTCTGGTCCAACCAGCGGACCAACCTCTGGACCAACCAGCGGACCA<br>ACAAGCGGACCAACATCCGGCCCAACTTCTGGACCGACTTCTGGACCAACCAGCGGACCAACTTCTGGGCCAACTAGCGGAC<br>CAACTGGAGGCCCAACCTCTGGACCTACTTCTGGACCAACCAGCGGACCAACATCCGGCCCAACTTCTGGACCAACATCCGG<br>TCCAACCAGCGGACCAACTGGAGGCCCAACCTCTGGACCAACTAGCGGACCAACCAGCGGACCAACATCCGGCCCAACTTCT |

| | |
|---|---|
| | GGACCAACATCCGGTCCAACCAGCGGACCAACTGGAGGCCCAACCTCTGGACCAACTAGCGGACCAACCAGCGGACCAACAT<br>CCGGACCAACTTCCGGACCAACCAGCGGACCTACTTCTGGACCAACATCTGGACCAACATCCGGCCCAACTTCTGGACCAAC<br>TAGCGGACCAACATCCGGCCCAACATCTGGACCAACCAGCGGACCTACTTCTGGACCAACATCTGGACCAACATCCGGCCCA<br>ACTTCTGGACCAACTAGCGGACCAACCAGCGGACCTACTTCTGGACCAACATCTGGACCAACCAGCGGACCAACTGGAGGCC<br>CAACCTCTGGACCAACTAGCGGCCCAACCTCTGGACCAACTAGCGGACCAACATCCGGCCCAACATCTGGACCAACCAGCGG<br>ACCTACTTCTGGACCAACATCTGGACCAACATCCGGCCCAACTTCTGGACCAACTAGCGGACCAACATCCGGCCCAACATCT<br>GGACCAACCAGCGGACCTACTTCTGGACCAACATCTGGACCAACATCTGGACCAACATCCGGCCCAACTTCTGGACCTACTT<br>CTGGACCAACCAGCGGACCTACTTCTGGACCAACATCCGACCCAACTTCTGGACCAACATCCGGCCCAACCTCTGGACCTAC<br>TTCTGGACCAACCAGCGGACCAACTTCCGGCCCAACATCTGGACCAACTGGCGGACCAACTAGTGGACCAACATCAGGACCA<br>ACCAGTGGACCAACATCAGGACCAACCAGTGGACCAACGTCTGGACCTACAAGCGGACCAACCAGCGGACCAACCAGCGGCC<br>CTACATCTGGCCCAACATCTGGCCCAACATCTGGTCCAACTTCTGGACCAACCAGCGGACCAACCAGCGGGCCTACAAGTGG<br>ACCTACAAGTGGACCTACAAGTGGACCTACAAGTGGACCTACAAGTGGACCTACAAGTGGACCTACAAGTGGACCTACAAGT<br>GGACCTACAAGTGGACCTACAAGTGGACCTACAAGTGGACCTACAAGTGGACCTACAAGTGGACCTACAAGTGGACCTACAA<br>GTGGACCCACAAGTGGGCCTACGTCCGGACCAACATTGGCGCCCACCAGTAACGTTCCAAGTTCTGCCCCATCGAGAAACTG<br>TGAATATGTTGTGGCCGACTTTGATGGGGACGATGTGAAGACTAATGCCTATCTTTTCCCAACAGCACTCATTGAATCTGGT<br>ATCTCGATCGAGGCATTCGGTTCTAATGGCACTGGGCACACCCCAAGCGATATGGCCAGGGTCATGGAGACCCCCACGGAGG<br>GGAATGCCATCTTCATTCAAGAATCGAATGTCAGAGATATCATTCCAAACGCTGAAGGAGGAGTCTTGTCGTTCACTTTCCA<br>AGCCACTGTGGAAGAAGTGATTGGGCTGCAGCTTTGGGATGTTTCGGAGGGCGGAATTGTTACCGTTCTCACAGAAGGTGGC<br>ACCATCGAGTCCTTCCCAATCATGGCATCTCCGAATGCCACCCAGACGATTGATATTGGTGTCTTTGAGGCTCTCGAAATGA<br>ACGTTACCTTTGGCGGGACATTGTCTGTGACGGCCATCGAAATTTGTTTTGACGGTTCCACGACTCCTGGTCCAGTACGATT<br>CGCGCCTCCATCCGAGACAAAGAGTCCCACTGCCATGCCAACAACAAGCCCCGCACCAACGGGCGAGCCCACAGAAAGTTCA<br>GCACCAACTGACTGTTACGATCAACTCCGTGCGAAACTTGTCGACCAGGTAGGAACGGATTCTACTCCTGTACCAGAGGATG<br>AAGAACTGATCGTCGTTACGTCCCAGAACGGCACACACGTTGGCTTTGAAGTCAACCAGGTGTGGGGATCAGGGGCAAACAT<br>CAGCCTGATTGCTGTGAATCACCATGTGAACTTTACATACACCGAATGCAGTCGCCAATACGATGTGACACCAGATTCATCG<br>TCGTTGTATTTGGCGAAGTGTTTCGATGAGTGGGCTGATGTCACTGTGTACATCTATACAGCAGACGACTTTTCGGAGGAAG<br>AATGTGACATCTGCGAAACGCCTGAAGAAGATGATGAAGACATCATGGTATTCCACTTCGAGATTCCATGCAACTCCGAGTG<br>CGATCTCTCCGCCTCTCCGACTGGAGGACCAACCAGCGGACCAACCTCCGGCCCAACTTCTGGACCAACCTCTGTACCCACC<br>TCTGAACCAACTAGCGGACCAACCAGTGGACCAACCAGCGGACCAACATCCGGACCAACATCCGGCCCAACCAGCGGCCCAA<br>CTTCTGGACCGACTTCCGGACCCACCTCTGGACCAACCAGTGGACCAACCAGCGGACCAACCAGCGGACCAACCAGCGGACC<br>AACCAGCGGACCAACATCTGGACCCACCTCCGGCCCAACATCTGGACCAACCAGTGGACCAACCAGTGGACCAACCAGCGGA<br>CCAACCAGCGGACCAACTTCTGGACCAACCTCCGGCCCAACATCTGGACCAACCAGTGGACCAACATCTGGACCAACGTCCG<br>GCCCAACATCTGGACCAACCAGCGGACCAACCAGCGGACCAACCTCCGGCCCAACATCTGGACCAACCAGTGGACCAACCAG<br>TGGACCAACTTCTGGACCTACCTCCGGCCCAACTTCTGGACCTACCTCCGGCCCAACATCCGGCCCAACATCCGGACCAACC<br>TCTGGACCAACCTCTGGACCAACCAGCGGACCAACCAGCGGACCAACCAGCGGACCAACCAGCGGACCAACCTCCGGCCCAA<br>CATCTGGACCAACATCTGGACCAACGTCCGGCCCAACATCTGGACCAACCAGCGGACCAACTTCTGGACCTACCTCCGGCCC<br>AACATCCGGACCAACCTCTGGACCAACCAGCGGACCAACATCTGGACCTACCTCCGGACCAACCAGCGGACCAACCAGCGGA<br>CCAACTTCTGGACCAACCTCCGGCCCAACATCTGGACCAACCAGTGGACCAACTTCTGGACCAACCTCTGGACCAACCTCCG<br>GCCCAACTTCTGGACCAACTAGCGGACCAACCAGCGGACCAACCAGCGGACCAACCAGCGGACCAACATCTGGACCAACTAG<br>CGGACCAACTAGCGGACCAACCAGCGGACCAACTTCTGGACCAACCTCTGGACCAACCTCCGGCCCAACTTCTGGACCAACT<br>AGCGGACCAACCAGCGGACCAACCAGCGGACCAACCAGCGGACCAACCAGCGGACCAACATCTGGACCAACTAGCGGACCAA<br>CTAGCGGACCAACCAGCGGACCAACTTCTGGACCTACCTCCGGACCAACATCTGGACCAACATCTGGACCAACTTCTGGACC<br>AACCAGTGGACCAACCTCCGGACCAACCAGTGGACCAACCAGCGGACCAACCAGCGGACCAACTTCTGGACCCACCTCCGGC<br>CCAACATCTGGACCAACCAGTGGACCAACTTCTGGACCTACCTCCGGCCCAACATCTGGACCAACCTCCGGACCAACCTCCG<br>GACCAACCTCCGGACCAACCTCCGGACCAACCTCCGGACCAACCAGTGGACCAACCTCCGGACCAACCTCCGGACCAACCAG<br>TGGACCAACCAGTGGACCAACCAGTGGACCAACCAGTGGACCAACCAGCGGCCCAACTTCTGGACCTACCTCCGGCCCAACA<br>TCCGGACCAACCTCTGGACCAACCTCTGGACCAACATCCGGACCAACCAGCGGACCAACATCTGGACCAACTTCTGGACCAA<br>CCAGCGGACCAACCAGTGGACCAACCAGTGGACCAACCAGTGGACCAACCAGCGGACCAACCAGCGGACCAACCAGCGGACC<br>AACCAGCGGACCAACCAGCGGACCAACCAGCGGACCAACCAGCGGACCAACCTCTGGACCTACTTCAGGCCCAACCTCTGGA<br>CCAACTAGTGGACCAACTAGTGGACCAACATCCGGACCAACATCCGGACCAACATCCGGACCAACATCCGGCCCAACATCTG<br>GACCAACTAGTGGACCAACATCCGGACCAACATCCGGCCCAACATCTGGACCAACATCCGGACCAACCTCCGGACCAACCTC<br>CGGCCCAACATCTGGACCAACCAGCGGACCAACCAGCGGACCAACTTCTGGACCCACCTCCGGACCAACATCCGGACCAACC<br>AGTGGACCAACTAGCGGACCAACCAGCGGACCAACCAGCGGACCAACTTCTGGACCTACCTCCGGCCCAACATCCGGACCAA<br>CTTCTGGACCTACCTCCGGACCAACCTCTGGACCAACCTCCGGCCCAACATCCGGACCAACCAGCGGACCAACATCTGGACC<br>AACATCTGGACCAACCTCCGGACCAACCAGCGGACCAACATCCGGACCAACATCTGGACCAACTAGTGGACCAACCAGCGGA<br>CCAACATCTGGACCAACCTCCGGCCCAACATCTGGACCAACCAGTGGACCAACCAGCGGCCCAACTTCTGGACCTACCTCCG<br>GCCCAACATCCGGACCAACCTCTGGACCAACCTCCGGACCAACATCCGGACCAACCTCCGGACCAACTAGTGGACCAACATC<br>CGGACCAACATCTGGACCAACATCTGGACCAACCAGTGGACCAACCAGTGGACCAACCAGCGGCCCAACTTCTGGACCAACT<br>AGCGGACCAACATCTGGACCAACCTCCGGACCAACCTCCGGACCAACCAGCGGACCAACCTCCGGACCAACATCTGGACCAA<br>CTAGTGGACCAACCAGCGGACCAACATCTGGACCAACCTCCGGCCCAACATCTGCACCAACTAGCGGACCAACTAGCGGACC<br>AACCTCTGGACCAACATCCGGACCAACCTCCGGACCAACATCTGGACCAACCTCCGGACCAACCTCTGGACCAACCAGCGGA<br>CCAACCTCCGGACCAACCAGTGGACCAACATCCGGACCAACTAGTGGACCAACCTCTGGACCAACCAGCGGACCAACCTCCG<br>GACCAACCAGTGGACCAACATCCGGACCAACTAGTGGACCAACATCCGGACCAACTAGTGGACCAACCAGCGGACCAACCTC<br>CGGACCAACCTCCGGACCAACCTCCGGACCAACCAGTGGACCAACCTCCGGACCAACCTCCGGACCAACCAGTGGACCAACC<br>AGTGGACCAACCAGTGGACCAACCTCCGGACCAACCTCCGGACCAACCAGTGGACCAACCAGTGGACCAACATCCGGACCAA<br>CATCCGGACCAACCTCCGGACCAACATCTGGACCAACTAGCGGACCAACCTCTGGACCAACCAGCGGACCAACATCCGGACC<br>AACATCCGGACCAACCAGTGGACCAACATCCGGACCAACATCCGGACCAACATCCGGACCAACCTCCGGACCAACATCTGGA<br>CCAACTAGCGGACCAACCTCTGGACCAACCAGCGGACCAACCTCCGGACCAACCAGTGGACCAACATCCGGACCAACTAGTG<br>GACCAACCAGCGGACCAACATCCGGACCAACCTCCGGACCAACATCTGGACCCACCTCTGGACCAACATCCGGACCAACCAG<br>CGGACCAACTAGCGGACCTACCTCCGGACCAACAAGTGGACCAACTTCGGGGCCAACGTCGGGACCGACATCTGGACCAACC |

| | |
|---|---|
| | TCAGGACCAACCTCTGGACCTACTTCTGGACCAACCAGCGGACCTACTTCCGGACCAACAACCAGCCCAGCACCTTCCGAGT<br>GCATCGACCAGGTCACACCAAAACCAACGGGAACATACGGTGCTGACGTCCCTCTGCCATTGGACAGCGACCTCATTGTCAT<br>CACCGGTCTGAACGGCACCCATGTGTCGTTCGCTGTGAACCCAGCCATCGGCGAAGACACCGACTCCAACATCACTGCCGTG<br>TCCATCGAGTACAGACCTAACGTGACCCATTCTGAGTGCAAGACCGAGTTCGATGTCGTCCCTGAGGCGGCGTTCGACTTGG<br>AAGCGGTGTGCTTCGATGGTTGGACAATGGTGACAGTCTACATCTACTTTGGAGAACCCTTGGAGGATCCGGAGCAGTGCGA<br>ATACTGTGAGACACCTGATGAAGACAGTGACGACTCTGTTGCGTACGTGTTCGAGTTGCCGTGCGAGACCGACTGTTATGAC<br>ACTATCGCGCCAACATCGATGCCAACGTCGGGACCTACCAGTGGTCCAACCAGCGGCCCAACTGAGACCATTTGCATCTACA<br>CAGTTCTTGACTTCGCCGCTGACGGCGACAACCAGCCAACTGGCAATGGCTATGTTCCCTCTGAGATATACGATATATTCGA<br>CATTGACTCGATCACCGTTGATCTTGGCGATAGTCCTTCGTCGTACGGCGGGCCCACTGGGAAGAACGCAAGATACTGGGTC<br>ACTGGTGACTCAGAGGCAGGCACAGCTATCATCATCCAAGAGAGCAATATGAACGACGCTGTTCCGAGCCCGACTGGTGGTG<br>ATATTGTGTTCACCTTCAACGGGCAATCAACCGAAACAATTGAACTTACTTTCTACGATCTTGCCGCGGATGTGACAATCAC<br>AGCGAGAACTACGGCCGATCCATCCGAGGGTGGACTCGCACAGACCACGACAGCCAGCGCCAACCCAGGTGGATTTACAAAG<br>GTTGAGATTGACGTTTCAATGATGAATCAAGTCATCGTGACGTTCACAGCTCCGGGTGCCATCGCATCTATTGGAGTATGCC<br>GTGACCCAAATACGCCACCCCCGCAAGGAGACAACAGCCCGTCAGCAACACTGGAACCGACGTCACAGCCAACGACAAGCCC<br>AGCGCCGACAGGCGAGCCAACCACCAGTGCTGCTCCAACCGATTGCTACGACCAACTCCGCGCCAAGTTCGTTCGCCAGGAA<br>GGTGCAAGCGTTTCCCTTCCTGAAGACAACGATTTGATCGTTGTCACTAGTCAGAACGGAACGCACGTGTCTTTCGAAGTCA<br>ATCCATCCATCGGCGAAGATACCGACTCCAACATCACGACTGTGTCAATCCAATACAGAGCGAATGTTACCTACTCTGAGTG<br>CAAGACCGAGTTCGATGTCGCCCCTGAGGCGGCGTTCGACTTGGAAGCGGTGTGCTTCGATGGTTGGACTATGGTGACAGTC<br>TACATCTATTTTGGCGAGCCTTTGGAGGATCCGGAGCAGTGCGATTTCTGCGAAACTCCTGATGAGGACAGTGACGACTCAG<br>TCGCATATGTCTTCGAGGTGCCATGCACAGCAAACTGTGATTTGTCAGCCTCACCAACCAGCGGGCCAACCTCTGGACCAAC<br>TTCTGGACCTACTTCTGGACCAACCTCCGGCCCAACTTCTGGACCTACAAGCGGACCAACATCTGGACCAACATCCGGACCT<br>ACGAGCGGACCAACATCCGGACCAACCAGTGGACCAACATCTGGACCAACCAGCGGACCTACTTCTGGACCAACTAGTGGAC<br>CAACTAGCGGACCAACCTCCGGCCCAACTTCTGGACCTACAAGCGGACCAACTTCTGGACCCACCTCTGGACCAACTAGCGG<br>ACCTACAAGCGGACCAACTAGCGGACCAACTTCGGGGCCTACATCTGGACCAACCAGCGGACCCACCTCTGGACCAACATCC<br>GGACCAACCAGCGGACCAACATCCGGACCAACCAGCGGACCAACCTCCGGACCAACATCTGGACCAACTAGTGGACCAACCA<br>GCGGACCAACATCCGGACCAACCAGCGGACCAACATCTGGACCAACCAGCGGACCAACATCTGGACCAACCAGCGGACCAAC<br>ATCCGGACCAACCAGTGGACCAACCAGTGGACCAACATCTGGACCAACTTCCGGACCAACTAGTGGACCAACCAGCGGACCA<br>ACCAGCGGACCAACATCCGGACCAACTAGCGGACCAACCTCTGGACCAACTAGTGGACCAACCAGCGGACCAACCTCCGGAC<br>CAACATCTGGACCAACTAGTGGACCAACCTCCGGACCAACATCTGGACCAACCAGTGGACCAACATCCGGACCAACCAGCGG<br>ACCAACATCCGGCCCAACTTCTGGACCAACTAGCGGACCAACATCTGGACCAACCTCCGGGCCAACCTCCGGACCAACCTCC<br>GGCCCAACATCTGGACCAACCAGCGGACCAACCAGTGGACCAACCAGTGGACCAACCAGTGGACCAACCTCCGGACCAACCT<br>CCGGACCAACCTCCGGACCAACCAGTGGACCAACCTCCGGACCAACCAGTGGACCAACATCCGGACCAACATCCGGACCAAC<br>CTCCGGACCAACATCCGGACCAACTAGCGGACCAACCTCTGGACCAACCAGCGGACCAACCTCCGGACCAACCAGTGGACCA<br>ACCAGTGGACCAACCTCCGGACCAACCAGTGGACCAACATCCGGACCAACCAGTGGACCAACATCCGGACCAACATCCGGAC<br>CAACCTCCGGACCAACATCCGGACCAACTAGCGGACCAACCTCTGGACCAACCAGCGGACCAACCTCCGGACCAACCTCCGG<br>ACCAACATCCGGACCAACTAGTGGACCAACATCCGGACCAACTAGTGGACCAACCAGCGGACCAACCAGCGGACCAACCAGC<br>GGACCAACCTCCGGACCAACCTCCGGACCAACCAGTGGACCAACCAGTGGACCAACCAGTGGACCAACCAGTGGACCAACCT<br>CCGGACCAACCTCCGGACCAACCTCCGGACCAACCTCCGGACCAACCAGTGGACCAACCAGTGGACCAACCAGTGGACCAAC<br>CAGTGGACCAACCAGTGGACCAACCTCCGGACCAACCAGTGGACCAACATCCGGACCAACCAGTGGACCAACATCCGGACCA<br>ACTAGTGGACCAACATCCGGACCAACTAGTGGACCAACCAGCGGACCAACCAGCGGACCAACCAGCGGACCAACCTCCGGAC<br>CAACCTCCGGACCAACCTCCGGACCAACCTCCGGACCAACCTCCGGACCAACCTCCGGACCAACCTCCGGACCAACCTCCGG<br>ACCAACCAGTGGACCAACCTCCGGACCAACCAGTGGACCAACATCCGGACCAACATCCGGACCAACATCCGGACCAACCTCC<br>GGACCAACATCCGGACCAACCTCCGGACCAACATCTGGACCAACCAGCGGACCAACCTCCGGACCAACCAGTGGACCAACAT<br>CCGGACCAACTAGTGGACCAACCAGCGGACCAACATCCGGACCAACCTCCGGACCAACATCTGGACCCACCTCTGGACCAAC<br>ATCCGGACCAACCAGCGGACCAACTAGCGGACCTACCTCCGGACCAACAAGTGGACCAACTTCGGGGCCAACGTCGGGACCG<br>ACATCTGGACCAACCTCAGGACCAACCTCTGGACCTACTTCTGGACCAACCAGCGGACCTACTTCCGGACCAACAACCAGCC<br>CAGCACCTTCCGAGTGCATCGACCAGGTCACACCAAAACCAACGGGAACATACGGCGCTGACGTCCCTCTGCCATTGGACAG<br>CGACCTCATTGTCATCACCGGTCTGAACGGCACCCATGTGTCGTTCGCTGTGAACCCAGCCATCGGCGAAGACACCGACTCC<br>AACATCACTGCCGTGTCCATCGAGTACAGACCTAACGTGACCCATTCTGAGTGCAAGACCGAGTTCGATGTCGTCCCTGAGG<br>CGGCGTTCGACTTGGAAGCGGTGTGCTTCGATAGTTGGACAATGGTGACAGTCTACATCTACTTTGGAGAACCCTTGGAGGA<br>TCCGGAGCAGTGCGAATACTGTGAGACACCTGATGAAGACAGTGACGACTCTGTTGCGTACGTGTTCGAGTTGCCGTGCGAG<br>ACCGACTGTTATGACACTATCGCGCCAACATCGATGCCAACGTCGGGACCTACCAGTGGTCCAACTGCAAGCCCTGCGCCAA<br>CCGACTGCTACGAACAACTCTTGCCAAAACTTACGCACCAATCCGGCGCAGACGTGAATATTCCAGAGAACGACGATCTCAT<br>CGTCATCACCAGCCAAAATGGAACGCACGTGGGCATTGAAGTTCGACAAGTTATTGAGAACTCGGAAGACGGAGCCTCGATG<br>CTTGCTGTGCAATACAGAAACTCGACAGCCGAATCCTGTGAGAAGAGCTCGGGAGTGACGTTTGGAACAACTGTGTCCTTCA<br>TCGCCACCTGCTTCGACTCCTACGCGGAAGTGCGAGTCTTCCTTGTGTTGGGTAATGCCACTGAGGACGCCGAGGATTGCGA<br>CGCATGCAGGTCACCGGACGAAGACGACACAGACAAGGTGGTCTGGGTCTTTGAAGTCCCATGTGAGTCTGGATGTGATCTT<br>TCGGATTCCCCGAGCCAAAGCCCAACAAGTTCTCCGACAACCAGCCCAGCACCTTCCGAGTGCATCGACCAGGTCACACCAA<br>AACCAACGGGAACATACGGTGCTGACGTCCCTCTGCCATTGGACAGCGACCTCATTGTCATCACCGGTCTGAACGGCACCCA<br>TGTGTCGTTCGCTGTGAACCCAGCCATCGGCGAAGACACCGACTCCAACATCACTGCCGTGTCCATCGAGTACAGACCTAAC<br>GTGACCCATTCTGAGTGCAAGACCGAGTTCGATGTCGCCCCTGAGGCGGCGTTCGACTTGGAAGCGGTGTGCTTCGATGGTT<br>GGACAATGGTGACAGTCTACATCTACTTTGGAGAACCCTTGGAGGATCCGGAGCAGTGCGAATACTGTGAGACACCTGATGA<br>AGACAGTGACGACTCTGTTGCGTACGTGTTCGAGTTGCCGTGCGAGACCGACTGTTATGACACTATCGCGCCAACATCAGCA<br>CCAACCAATTGCTACGATCGCGTCAACGTCGCTCCGGTCAGAAAGGCTGGAGACGACATGGCTATTCCGGAGGGTGCAGTGA<br>CCATTGTGGAGCATGATGGCGCCACTGTCAAGTTCAAGGTCACACAATTGTGGAAAGATGTGGACGTGAAGATGTTGGCGTT<br>CCAATATCGCGAAACCGCATCTGCGGAGAAGTGCTTCCCGGACACAACCGTTGACTTTGGAACAACGTATGAACATACCGCT<br>GTCTGCTTCGACGGTTTCGCGTCCGTTGGTATCTATCTGTACACCGGCGATGATTTCGATCCAACGACATGTGAAGCATGTA<br>AACAGCCAACGGACATCGACGATGGCGTCGCACTGTTCTACTTCGAGATGCCATGCAACTCTGAATGCGCACCACCAACCCC<br>GACATCAGAGACAGAGGCATCGAACTGCTACGACGATGTGGTTCTGGATGACTCTTCTGGAACGTGTGTGTATGATGCGAGC |

| | |
|---|---|
| | CCTGTGTCAATCGTATCGCAGCACGGCAGCACAGTCAAGTTCTCTGTGTCTCACACGTGGGACATAGCACAGACTCATGCCT<br>TGGGTGTCGTCCCAGCTGCGCTGAGCCAGATGTCGATCCGGTACAAGGCCCAGGGAGAAAATTCCATGGGTTGCACAACGAG<br>ACTCGAACCAGCCGCCGGATATTTGGACACATACACAGCTCAGTGCAACAGCAACGGGCTTGCACAGGTCGAGATCTTCGTG<br>CATGACAGTACCTTCCCCAAGGACAGCAGCGCGGAGATCCCGGACGAATGCGACCAACCCACCGATCCTGACCATACTTGCG<br>GATACTCGTTTGTCATCCCATGTAACGCGGACATGATGTGCGACAGCAACCGCCGAATGCAGAAGATCCGCAGCCACATGAA<br>GGAAGTGAAGGAAGAGTCTTACGCGCCAGGCCTGAGCTCAAGCATTCTCATGGAATCGTTCAAGAAGACCTCCTTGGAAACA<br>GATTTGCCGGCTGAAGGTGATGAAGATCCATACTGCGCTGCCGAAGACTTCCCGTGTGAGGGCGACTCGCCCAACATGGTGT<br>ACGTCTGCCATTACTCTGTCACGAAGGGATACCAAACTTTCTGCGTGCCAGAGGCAGACTCCGACATCCTTCGTTTGTACTC<br>TTCGGATTACTGCGGAAAGTGCGATGGCCAGTAA |
| **Protein** | MMMVTKRVTPIPPWAGRSGLLLVLCALVLTGGKEAEGLKVTRVSSGQGYHGGPSDGVDHQRKLLGDANRHGKLLGGEEEKDE<br>AEGKREGGAMRFLYGAQVDVGARTGSFYSALGRNVKQETLEKAQDCDPDNKSTWKFCAPHMLPNSNFNPNLPPEIAAQRSNE<br>VAYLAKQAKGGGLTGPNKGNGLDKAKPIVGSESPGRSVDAQSDGGTLLGDSEAESADATVGSTIGAKDAGGFGSDGGVKNKR<br>TSPGSNTVGEGLVRDEAVPTNDSVSARTSLDELSSLPAICVLDPDHPACQEARARMLQTTDAGGGSSAPSLASATDNSPVEA<br>PVSAENPYASDWPSGSKTTESPTKDMCYGNMEIAKRKFCESEMPSVSSEPSQTPTAFSPSPESGQTIPTATTPSDPGSPSGQ<br>GSDAPSDVPSESPTGEGGSRTRRPADSPDSFKPPGNSGAGSDAPSDVPSESPTGEGGSRTRRPADSPDSFKPPGNSGAGSDA<br>PSDVPSESPTGEGGSRSRRPFGIPGTSFDTPVGTDPPTSSSAPTDCYDSIEPKLERQVGVDFPWADGVVDVQHQTGNSVTFQ<br>ILQVWSRSILPMLAVHYHNSPGSTVCDMNVDVQYQDTFDYTAHCFGNYTDIRVYLFVGDDFDEEKCEACQTPPDDSNDTVAY<br>FLELSCHDCVEVTAPPSVTAPSQPVISPTPPTGDVGSDAPSDVPSEAPTGDGGSRTRRPQSPQVPSAPTGDEPTGDEPTGEV<br>GSDAPSDVPSEAPTGDGGSRTRRPQSPQVPSAPTGDAPTGDEPTGEVGSDAPSDVPSEAPTGDGGSRTRRPQSPQVPSAPTG<br>DAPTGDEPTGDEPTGEVGSDAPSDVPSEAPTGDGGSRTRRPQSPQVPSAPTGDEPTGDEPTGEVGSDAPSDVPSEAPTGDGG<br>SRTRRPQSPQVPSAPTGDAPTGDEPTGEVGSDAPSDVPSEAPTGDGGSRTRRPQSPQVPSAPTGDEPTGDEPTGEVGSDAPS<br>DVPSEAPTGDGGSRTRRPQSPQVPSAPTGDAPTGDEPTGEVGSDAPSDVPSEAPTGDGGSRTRRPQSPQVPSAPTGDEPTGD<br>EPTGEVGSDAPSDVPSEAPTGDGGSRTRRPQSPQVPSTPTGDEPTGDEPTGDEPTGDEPTGEVGSDAPSDVPSEAPTGDGRT<br>RTRRPQSPQVPSAPSAEEPTGDEPTGDEPTGDEPTGEVGSDAPSDVPSEAPTGDGRTRTRRPQSPQVPSAPSAEEPTGDEPT<br>GDEPTGDEPTGDEPTGDEPTGDEPTGEVGSDAPSDVPSEAPTGDGRTRTRRPQSPQVPSAPSAEEPTGDEPTGDEPTGDEPT<br>GDEPTGDEPTGEVGSDAPSDVPSEAPTGDGRTRTRRPQSPQVPSAPSAEEPTGDEPTGDEPTGDEPTGEVGSDVPSDVPSEA<br>PTGDGGSRTRRPQSPQVPSVPTSGPTSGPTSGPTGGPTSGPTSGPTSGPTSGPTSGPTSGPTSGPTSGPTSGPTSGPTGGPT<br>SGPTSGPTSGPTSGPTSGPTSGPTSGPTSGPTSGPTSGPTSGPTSGPTSGPTSGPTSGPTSGPTSGPTSGPTSGPTSGPTSG<br>PTSGPTSGPTSGPTSGPTSGPTSGPTSGPTSGPTSGPTSGPTSGPTSGPTSGPTSGPTSGPTSGPTSGPTSGPTSQPTYVCS<br>TEARLVDKDGGNTKYDEGLDDEITGDDGIPFEVSEREVDSVKISLEPLLDAIEALRTGGAVGEDGELGSVAVVFDRISTDND<br>GNEVTENDFCTETSVTEATGPAEEYVAQCVEDDDFAGTTVAIVYVVVADSANTQEDPSGSLIGAAGGESALGTSGCGVPFDE<br>DTVIPVIVYKYVVPCQYSCDGAPTSSPTGEPTSAPTVLPGHPTGSPVTATPTSTPTSLPTSGPTSAPTYTPTSGPTSGPTSG<br>PTSGPTSGPTGGPTSGPTSGPTSGPTSGPTSGPTSGPTSGPTSGPTSGPTSGPTSGPTSGPTSGPTSGPTSGPTSGPTSGPT<br>SGPTSGPTSGPTSGPTSGPTSGPTGGPTSGPTSGPTSGPTSGPTSGPTSGPTSGPTSGPTSGPTSGPTSGPTSGPTSGPTSG<br>PTSGPTSGPTSGPTSGPTSGPTSGPTSGPTSGPTSGPTSGPTSGPTSGPTSGPTSGPTSGPTSGPTSGPTSGPTSGPTGGPT<br>SGPTSGPTSGPTSGPTSGPTSGPTSGPTSGPTSGPTSGPTSGPTSGPTSGPTSGPTSGPTSGPTSGPTSGPTSGPTSGPTSG<br>PTSGPTSGPTSGPTSGPTSGPTSGPTSGPTSGPTSGPTSGPTSGPTSGPTSGPTSGPTSGPTSGPTSGPTSGPTSGPTSGPT<br>SGPTSGPTSGPTSGPTSGPTSGPTSGPTSGPTSGPTSGPTSGPTSGPTSGPTSGPTSGPTSGPTSGPTSGPTGGPTSGPTSG<br>PTSGPTSGPTSGPTSGPTSGPTSGPTSGPTSGPTGGPTSGPTSGPTSGPTSGPTSGPTSGPTSGPTSGPTSGPTSGPTSGPT<br>SGPTSGPTSGPTSGPTSGPTSGPTSGPTSGPTSGPTSGPTSGPTSGPTSGPTSGPTSGPTSGPTSGPTSGPTSGPTSGPTSG<br>PTSGPTSGPTSGPTSGPTSGPTSGPTSGPTSQPTYVCSTEARLVDKDGGNTKYDEGLDDEITGDDGIPFEVSEREVDSVKIS<br>LEPLLDAIEALRTGGAVGEDGELGSVAVVFDRISTDNDGNEVTENDFCTETSVTEATGPAEEYVAQCVEDDDFAGTTVAIVY<br>VVVADSANTQEDPSGSLIGAAGGESALGTSGCGVPFESGPSSTPVIVYKYVVPCQYSCEGSPTSVPTDAPTGGPTSGPTSGP<br>TSGPTSGPTSGPTSGPTSGPTSGPTSGPTSGPTSGPTSGPTSGPTSGPTSGPTSGPTSGPTGGPTSGPTSGPTSGPTSGPTS<br>GPTSGPTSGPTGGPTSGPTSGPTSGPTSGPTSGPTSGPTSGPTSGPTSGPTSGPTSGPTSGPTSGPTSGPTSGPTSGPTSGP<br>TSGPTSGPTSGPTSGPTSGPTSGPTSQPTYVCSTEARLVDKDGGNTKYDEGLDDEITGDDGIPFEVSEREVDSVKISLEPLL<br>DAIEALRTGGAVGEDGELGSVAVVFDRISTDNDGNEVTENDFCTETSVTEATGPAEEYVAQCVEDDDFAGTTVAIVYVVVAD<br>SANTQEDPSGSLIGAAGGESALGTSGCGVPFESGPSSTPVIVYKYVVPCQYSCEGSPTSVPTDAPTGGPTSGPTSGPTSGPT<br>SGPTSGPTSGPTSGPTSGPTSGPTSGPTSGPTSGPTSGPTSGPTSGPTSGPTSGPTGGPTSGPTSGPTSGPTSGPTSGPTSG<br>PTSGPTGGPTSGPTSGPTSGPTSGPTSGPTSGPTSGPTSGPTSGPTSGPTSGPTSGPTSGPTSGPTSGPTSGPTSGPTSGPT<br>SGPTSGPTSGPTSGPTSGPTSQPTYVCSTEARLVDKDGGNTKYDEGLDDEITGDDGIPFEVSEREVDSVKISLEPLLDAIEA<br>LRTGGAVGEDGELGSVAVVFDRISTDNDGNEVTENDFCTETSVTEATGPAEEYVAQCVEDDDFAGTTVAIVYVVVADSANTQ<br>EDPSGSLIGAAGGESALGTSGCGVPFESGPSSTPVIVYKYVVPCQYSCEGSPTSVPTDAPTGGPTSGPTSGPTSGPTSGPTS<br>GPTSGPTSGPTSGPTSGPTSGPTSGPTSGPTSGPTSGPTSGPTSGPTSGPTGGPTSGPTSGPTSGPTSGPTSGPTSGPTSGP<br>TGGPTSGPTSGPTSGPTSGPTSGPTSGPTSGPTSGPTSGPTSGPTSGPTSGPTSGPTSGPTSGPTSGPTSGPTSGPTSGPTS<br>GPTSGPTSGPTSGPTSQPTYVCSTEARLVDKDGGNTKYDEGLDDEITGDDGIPFEVSEREVDSVKISLEPLLDAIEALRTGG<br>AVGEDGELGSVAVVFDRISTDNDGNEVTENDFCTETSVTEATGPAEEYVAQCVEDDDFAGTTVAIVYVVVADSANTQEDPSG<br>SLIGAAGGESALGTSGCGVPFESGPSSTPVIVYKYVVPCQYSCEGSPTSVPTDAPTGGPTSGPTSGPTSGPTSGPTSGPTSG<br>PTSGPTSGPTSGPTSGPTSGPTSGPTSGPTSGPTSGPTSGPTSGPTGGPTSGPTSGPTSGPTSGPTSGPTSGPTSGPTGGPT<br>SGPTSGPTSGPTSGPTSGPTSGPTSGPTGGPTSGPTSGPTSGPTSGPTSGPTSGPTSGPTSGPTSGPTSGPTSGPTSGPTSG<br>PTSGPTSGPTSGPTSGPTSGPTSGPTSGPTSGPTSGPTSGPTSQPTYVCSTEARLVDKDGGNTKYDEGLDDEITGDDGIPFE<br>VSEREVDSVKISLEPLLDAIEALRTGGAVGEDGELGSVAVVFDRISTDNDGNEVTENDFCTETSVTEATGPAEEYVAQCVED<br>DDFAGTTVAIVYVVVADSANTQEDPSGSLIGAAGGESALGTSGCGVPFESGPSSTPVIVYKYVVPCQYSCEGSPTSVPTDAP<br>TGGPTSGPTSGPTSGPTSGPTSGPTSGPTSGPTSGPTSGPTSGPTSGPTSGPTSGPTSGPTSGPTSGPTSGPTGGPTSGPTS<br>GPTSGPTSGPTSGPTSGPTSGPTGGPTSGPTSGPTSGPTSGPTSGPTSGPTSGPTSGPTSGPTSGPTSGPTSGPTSGPTSGP<br>TSGPTSGPTSGPTSGPTSGPTSGPTSGPTSGPTSGPTSGPTSGPTSGPTSGPTSGPTSGPTSGPTSGPTSGPTSGPTSGPTS<br>GPTSGPTSGPTSGPTSGPTSQXXXXXXXXXXXXXXXXXXXXXXXXXFEVSEREVDSVKISLEPLLDAIEALRTGGAVGEDGE<br>LGSVAVVFDRISTDNDGNEVTENDFCTETSVTEATGPAEEYVAQCVEDDDFAGTTVAIVYVVVADSANTQEDPSGSLIGAAG<br>GESALGTSGCGVPFESGPSSTPVIVYKYVVPCQYSCEGSPTSVPTDAPTGGPTSGPTSGPTSGPTSGPTSGPTSGPTSGPTS |

| | |
|---|---|
| | GPTSGPTSGPTSGPTSGPTSGPTSGPTSGPTSGPTSGPTGGPTSGPTSGPTSGPTSGPTSGPTSGPTSGPTGGPTSGPTSGP<br>TSGPTSGPTSGPTSGPTSGPTGGPTSGPTSGPTSGPTSGPTSGPTSGPTSGPTSGPTSGPTSGPTSGPTSGPTSGPTSGPTS<br>GPTSGPTSGPTSGPTSGPTSGPTSGPTSGPTSGPTSQPTYVCSTEARLVDKDGGNTKYDEGLDDEITGDDGIPFEVSEREVD<br>SVKISLEPLLDAIEALRTGGAVGEDGELGSVAVVFDRISTDNDGNEVTENDFCTETSVTEATGPAEEYVAQCVEDDDFAGTT<br>VAIVYVVVADSANTQEDPSGSLIGAAGGESALGTSGCGVPFESGPFSTPVIVYKYVVPCQYSCEGSPTSVPTDAPTGGPTSG<br>PTSGPTSGPTSGPTSGPTSGPTSGPTSGPTSGPTSGPTSGPTSGPTSGPTSGPTSGPTSGPTSGPTGGPTSGPTSGPTSGPT<br>SGPTSGPTSGPTSGPTGGPTSGPTSGPTSGPTSGPTSGPTSGPTSGPTGGPTSGPTSGPTSGPTSGPTSGPTSGPTSGPTSG<br>PTSGPTSQPTYVCSTEARLVDKDGGNTKYDEGLDDEITGDDGIPFEVSEREVDSVKISLEPLLDAIEALRTGGAVGEDGELG<br>SVAVVFDRISTDNDGNEVTENDFCTETSVTEATGPAEEYVAQCVEDDDFAGTTVAIVYVVVADSANTQEDPSGSLIGAAGGE<br>SALGTSGCGVPFESGPSSTPVIVYKYVVPCQYSCEGSPTSVPTDAPTGGPTSGPTSGPTSGPTSGPTSGPTSGPTSGPTSGP<br>TSGPTSGPTSGPTSGPTSGPTSGPTSGPTSGPTSGPTGGPTSGPTSGPTSGPTSGPTSGPTSGPTSGPTGGPTSGPTSGPTS<br>GPTSGPTSGPTSGPTSGPTGGPTSGPTSGPTSGPTSGPTSGPTSGPTSGPTSGPTSGPTSGPTSGPTSGPTSGPTSGPTSGP<br>TSGPTSGPTSGPTSGPTSGPTSGPTSGPTSGPTSGPTSGPTSGPTSGPTSGPTSGPTSGPTSGPTSGPTSQPTYVCSTEARLVDKD<br>GGNTKYDEGLDDEITGDDGIPFEVSEREVDSVKISLEPLLDAIEALRTGGAVGEDGELGSVAVVFDRISTDNDGNEVTENDF<br>CTETSVTEATGPAEEYVAQCVEDDDFAGTTVAIVYVVVADSANTQEDPSGSLIGAAGGESALGTSGCGVPFESGPSSTPVIV<br>YKYVVPCQYSCEGSPTSVPTDAPTGGPTSGPTSGPTSGPTSGPTSGPTSGPTSGPTSGPTSGPTSGPTSGPTSGPTSGPTSG<br>PTSGPTSGPTSGPTGGPTSGPTSGPTSGPTSGPTSGPTSGPTSGPTGGPTSGPTSGPTSGPTSGPTSGPTSGPTSGPTGGPT<br>SGPTSGPTSGPTSGPTSGPTSGPTSGPTSGPTSGPTSGPTSGPTSGPTSGPTSGPTSGPTSGPTSGPTSGPTSGPTSGPTSG<br>PTSGPTSGPTSGPTSGPTSGPTSGPTSGPTSGPTSGPTSGPTSQPTYVCSTEARLVDKDGGNTKYDEGLDDEITGDDGIPFE<br>VSEREVDSVKISLEPLLDAIEALRTGGAVGEDGELGSVAVVFDRISTDNDGNEVTENDFCTETSVTEATGPAEEYVAQCVED<br>DDFAGTTVAIVYVVVADSANTQEDPSGSLIGAAGGESALGTSGCGVPFESGPSSTPVIVYKYVVPCQYSCEGSPTSVPTDAP<br>TGGPTSGPTSGPTSGPTSGPTSGPTSGPTSGPTSGPTSGPTSGPTSGPTSGPTSGPTSGPTSGPTSGPTSGPTGGPTSGPTS<br>GPTSGPTSGPTSGPTSGPTSGPTGGPTSGPTSGPTSGPTSGPTSGPTSGPTSGPTGGPTSGPTSGPTSGPTSGPTSGPTSGP<br>TSGPTSGPTSGPTSGPTSGPTSGPTSGPTSGPTSGPTSGPTSGPTSGPTSGPTSGPTSGPTSGPTSGPTGGPTSGPTSGPTS<br>GPTSGPTSGPTSGPTSGPTSGPTSGPTSGPTSGPTSGPTSGPTSGPTSGPTSGPTSGPTSGPTSGPTSGPTSGPTSGPTSGP<br>TSDPTSGPTSGPTSGPTSGPTSGPTSGPTSGPTGGPTSGPTSGPTSGPTSGPTSGPTSGPTSGPTSGPTSGPTSGPTSGPTS<br>GPTSGPTSGPTSGPTSGPTSGPTSGPTSGPTSGPTSGPTSGPTSGPTSGPTSGPTSGPTSGPTSGPTSGPTSGPTSGPTSGP<br>TLAPTSNVPSSAPSRNCEYVVADFDGDDVKTNAYLFPTALIESGISIEAFGSNGTGHTPSDMARVMETPTEGNAIFIQESNV<br>RDIIPNAEGGVLSFTFQATVEEVIGLQLWDVSEGGIVTVLTEGGTIESFPIMASPNATQTIDIGVFEALEMNVTFGGTLSVT<br>AIEICFDGSTTPGPVRFAPPSETKSPTAMPTTSPAPTGEPTESSAPTDCYDQLRAKLVDQVGTDSTPVPEDEELIVVTSQNG<br>THVGFEVNQVWGSGANISLIAVNHHVNFTYTECSRQYDVTPDSSSLYLAKCFDEWADVTVYIYTADDFSEEECDICETPEED<br>DEDIMVFHFEIPCNSECDLSASPTGGPTSGPTSGPTSGPTSVPTSEPTSGPTSGPTSGPTSGPTSGPTSGPTSGPTSGPTSG<br>PTSGPTSGPTSGPTSGPTSGPTSGPTSGPTSGPTSGPTSGPTSGPTSGPTSGPTSGPTSGPTSGPTSGPTSGPTSGPTSGPT<br>SGPTSGPTSGPTSGPTSGPTSGPTSGPTSGPTSGPTSGPTSGPTSGPTSGPTSGPTSGPTSGPTSGPTSGPTSGPTSGPTSG<br>PTSGPTSGPTSGPTSGPTSGPTSGPTSGPTSGPTSGPTSGPTSGPTSGPTSGPTSGPTSGPTSGPTSGPTSGPTSGPTSGPT<br>SGPTSGPTSGPTSGPTSGPTSGPTSGPTSGPTSGPTSGPTSGPTSGPTSGPTSGPTSGPTSGPTSGPTSGPTSGPTSGPTSG<br>PTSGPTSGPTSGPTSGPTSGPTSGPTSGPTSGPTSGPTSGPTSGPTSGPTSGPTSGPTSGPTSGPTSGPTSGPTSGPTSGPT<br>SGPTSGPTSGPTSGPTSGPTSGPTSGPTSGPTSGPTSGPTSGPTSGPTSGPTSGPTSGPTSGPTSGPTSGPTSGPTSGPTSG<br>PTSGPTSGPTSGPTSGPTSGPTSGPTSGPTSGPTSGPTSGPTSGPTSGPTSGPTSGPTSGPTSGPTSGPTSGPTSGPTSGPT<br>SGPTSGPTSGPTSGPTSGPTSGPTSGPTSGPTSGPTSGPTSGPTSGPTSGPTSGPTSGPTSGPTSGPTSGPTSGPTSGPTSG<br>PTSGPTSGPTSGPTSGPTSGPTSGPTSGPTSGPTSGPTSGPTSGPTSGPTSGPTSGPTSGPTSGPTSGPTSGPTSGPTSGPT<br>SGPTSGPTSGPTSGPTSGPTSGPTSGPTSGPTSGPTSGPTSGPTSGPTSGPTSGPTSGPTSGPTSGPTSGPTSGPTSGPTSG<br>PTSGPTSAPTSGPTSGPTSGPTSGPTSGPTSGPTSGPTSGPTSGPTSGPTSGPTSGPTSGPTSGPTSGPTSGPTSGPTSGPT<br>SGPTSGPTSGPTSGPTSGPTSGPTSGPTSGPTSGPTSGPTSGPTSGPTSGPTSGPTSGPTSGPTSGPTSGPTSGPTSGPTSG<br>PTSGPTSGPTSGPTSGPTSGPTSGPTSGPTSGPTSGPTSGPTSGPTSGPTSGPTSGPTSGPTSGPTSGPTSGPTSGPTSGPT<br>SGPTSGPTSGPTSGPTSGPTSGPTSGPTSGPTSGPTSGPTSGPTSGPTSGPTSGPTSGPTSGPTTSPAPSECIDQVTPKPTG<br>TYGADVPLPLDSDLIVITGLNGTHVSFAVNPAIGEDTDSNITAVSIEYRPNVTHSECKTEFDVVPEAAFDLEAVCFDGWTMV<br>TVYIYFGEPLEDPEQCEYCETPDEDSDDSVAYVFELPCETDCYDTIAPTSMPTSGPTSGPTSGPTETICIYTVLDFAADGDN<br>QPTGNGYVPSEIYDIFDIDSITVDLGDSPSSYGGPTGKNARYWVTGDSEAGTAIIIQESNMNDAVPSPTGGDIVFTFNGQST<br>ETIELTFYDLAADVTITARTTADPSEGGLAQTTTASANPGGFTKVEIDVSMMNQVIVTFTAPGAIASIGVCRDPNTPPPQGD<br>NSPSATLEPTSQPTTSPAPTGEPTTSAAPTDCYDQLRAKFVRQEGASVSLPEDNDLIVVTSQNGTHVSFEVNPSIGEDTDSN<br>ITTVSIQYRANVTYSECKTEFDVAPEAAFDLEAVCFDGWTMVTVYIYFGEPLEDPEQCDFCETPDEDSDDSVAYVFEVPCTA<br>NCDLSASPTSGPTSGPTSGPTSGPTSGPTSGPTSGPTSGPTSGPTSGPTSGPTSGPTSGPTSGPTSGPTSGPTSGPTSGPTS<br>GPTSGPTSGPTSGPTSGPTSGPTSGPTSGPTSGPTSGPTSGPTSGPTSGPTSGPTSGPTSGPTSGPTSGPTSGPTSGPTSGP<br>TSGPTSGPTSGPTSGPTSGPTSGPTSGPTSGPTSGPTSGPTSGPTSGPTSGPTSGPTSGPTSGPTSGPTSGPTSGPTSGPTS<br>GPTSGPTSGPTSGPTSGPTSGPTSGPTSGPTSGPTSGPTSGPTSGPTSGPTSGPTSGPTSGPTSGPTSGPTSGPTSGPTSGP<br>TSGPTSGPTSGPTSGPTSGPTSGPTSGPTSGPTSGPTSGPTSGPTSGPTSGPTSGPTSGPTSGPTSGPTSGPTSGPTSGPTS<br>GPTSGPTSGPTSGPTSGPTSGPTSGPTSGPTSGPTSGPTSGPTSGPTSGPTSGPTSGPTSGPTSGPTSGPTSGPTSGPTSGP<br>TSGPTSGPTSGPTSGPTSGPTSGPTSGPTSGPTSGPTSGPTSGPTSGPTSGPTSGPTSGPTSGPTSGPTSGPTSGPTSGPTS<br>GPTSGPTSGPTSGPTSGPTSGPTSGPTSGPTSGPTSGPTSGPTSGPTSGPTSGPTSGPTSGPTSGPTSGPTSGPTSGPTSGP<br>TSGPTSGPTSGPTSGPTSGPTSGPTSGPTSGPTSGPTSGPTSGPTSGPTSGPTSGPTSGPTSGPTSGPTTSPAPSECIDQVT<br>PKPTGTYGADVPLPLDSDLIVITGLNGTHVSFAVNPAIGEDTDSNITAVSIEYRPNVTHSECKTEFDVVPEAAFDLEAVCFD<br>SWTMVTVYIYFGEPLEDPEQCEYCETPDEDSDDSVAYVFELPCETDCYDTIAPTSMPTSGPTSGPTASPAPTDCYEQLLPKL<br>THQSGADVNIPENDDLIVITSQNGTHVGIEVRQVIENSEDGASMLAVQYRNSTAESCEKSSGVTFGTTVSFIATCFDSYAEV<br>RVFLVLGNATEDAEDCDACRSPDEDDTDKVVWVFEVPCESGCDLSDSPSQSPTSSPTTSPAPSECIDQVTPKPTGTYGADVP<br>LPLDSDLIVITGLNGTHVSFAVNPAIGEDTDSNITAVSIEYRPNVTHSECKTEFDVAPEAAFDLEAVCFDGWTMVTVYIYFG<br>EPLEDPEQCEYCETPDEDSDDSVAYVFELPCETDCYDTIAPTSAPTNCYDRVNVAPVRKAGDDMAIPEGAVTIVEHDGATVK<br>FKVTQLWKDVDVKMLAFQYRETASAEKCFPDTTVDFGTTYEHTAVCFDGFASVGIYLYTGDDFDPTTCEACKQPTDIDDGVA<br>LFYFEMPCNSECAPPTPTSETEASNCYDDVVLDDSSGTCVYDASPVSIVSQHGSTVKFSVSHTWDIAQTHALGVVPAALSQM |

| | |
|---|---|
| | SIRYKAQGENSMGCTTRLEPAAGYLDTYTAQCNSNGLAQVEIFVHDSTFPKDSSAEIPDECDQPTDPDHTCGYSFVIPCNAD<br>MMCDSNRRMQKIRSHMKEVKEESYAPGLSSSILMESFKKTSLETDLPAEGDEDPYCAAEDFPCEGDSPNMVYVCHYSVTKGY<br>QTFCVPEADSDILRLYSSDYCGKCDGQ* |
| **Ca5255** | |
| **gDNA/cDNA** | ATGAACGTCAAGACTCTCCTGGGATCGCTCGTGTTGAGCCAGATCGGCAGCAATGCCGTCCATGGCATGTTCGACACCGTCT<br>CTGACGCTGGTGGATCCAACTCGGAGCCATGCCCGTTGGACTATCAGTTGCTCCGTTCCAGCGACTTCCAGTACTCGGGAGC<br>ACCAATCGAGATCGTGAGCCAGGACACCAGCTCCGTCTCCTTCAAGGTTTACAACGTGTGGCCACAAGACGGCCGCCCCACC<br>GCCGAGATCTACACCGAGTACTCCGAGAACAACAGCATCGGCGGTGTCAACTGCGAGATGTTCGCCAACGTCCCGGCCAACC<br>AGGACGTGGGCGAAACCTTCACTGCGCAGTGCTTCCAGACCAAGCCAAAATCGGTCGTCACCCTCTACGTGCGTGATCCCGA<br>CATCGACAAGACCATCGGAACTGGCAAAGTTCCGCAGTGCTGCCATGCCGACCTCTCTATCCAGTACGCAGTCCTCAAGTTC<br>ACCTACGTGCTCGAGTGCGTCCCAACCTGCGAAGATCCCACCCCATTCTCCAGCTGCGGTGACGGCACAGTGGATCCAGGTG<br>AAGAGTGCGATGATGGCAACAAGATCAACGACGATTCCTGCACCAACCAGTGCACGAGCGCCAAGTGCGGTGATGGCATCAT<br>GCAATCCGGCGAAGAGTGCGATGATGGCAACAAGGTCGACAACGATGGCTGCACCAACGGCTGCAAACTTCCAAAATGCGGT<br>GACGGTTTCATGCAGGCCGGCGAAGAATGCGACGACGGGAACGCAGTCAACAACGATGGCTGCACCAACGGCTGCAAGCTTC<br>CAACCTGCGGCGATGGAATTGTCCAGAACGGAGAAGAATGCGACGACGGCAACGCAGTCAATAACGATGGCTGCACCAACAC<br>CTGCAAGAACGCCAAGTGCGGTGACGGTATCATGCAGGCCGGCGAAGAATGCGACGACGGCAACGCAGTCAACAACGATGGC<br>TGCACCAACGGCTGCAAGCTTCCAACCTGCGGCGATGGTATCGTCCAGAACGGAGAAGAATGTGACGATGGCAACAACAGCA<br>ACACCGATTCCTGCACCAACAACTGCAAGAACGCCAAGTGCGGTGACGGTATCATGCAGGCCGGCGAAGAATGCGACGATGG<br>CAACGCAGTCAATAACGATGGCTGTACCAACGGCTGCAAGCTTCCAACCTGCGGCGATGGTATCGTCCAGAACGGAGAAGAA<br>TGTGACGACGGCAACAACAGCAACACCGATTCCTGCACCAACACCTGCAAGAACGCCAAGTGCGGTGACGGTTTCATGCAGG<br>CCGGCGAAGAGTGTGATGATGGCAACGCAGTCAACAACGATGGCTGCACCAACGGCTGCAAGCTTCCAACCTGCGGCGATGG<br>TATCGTCCAGAACGGAGAGGAATGCGACGACGGCAACAACAGCAACACTGATTCCTGCACCAACAACTGCAAGAACGCCAAG<br>TGTGGTGACGGTTTCATGCAGGCCGGCGAAGAGTGTGATGATGGTAACAACAGCAACACTGATGGCTGCACCAACGGCTGCA<br>AGCTTCCCAAATGCGGTGATGGTATCGTCCAGAACGGAGAAGAGTGTGACGATGGCAACAACAACAACAACGACAACTGCAC<br>CAACAATTGCGAAATCCCCGTTCCAAACACATGCGGTAATGGCATCGTCGATCCCGGCGAGGAATGCGACGAAGGAACTCGC<br>AATGACTACTCCGGCACTCGATGCAAGCCAGACTGCACGCTCCCACCCGGCGTGTGCCCGCTGATCACCCTCGACTTCAACG<br>ACCTTCCAGATGGTGGCTTCATCTACGACGAATGGTGGAACACGAAGGGTGTGAAGATCATCGCCGCCAGCTCGTCTTCCAG<br>GGCGTACACACCAAGACCCACATCCAAGTGCAAAACTGGTGAGCTTGGTTGCTACTGCAGCTTCAATCCCAATTACCCATTG<br>CAGTGCAGCAACTGGAGAAGCGACAAAAGCTACGGAGCCGCCCGTGTCTTCGACACCTCCAAACCGACTGGGTACAGAACAT<br>CAGACTCAAAGTATGCATTGTACAACAACCCCATGTGTGACCCAAGGAGAAGCGGCGACCACGGCGACCCAGATCTCGGAAG<br>CCCCAACGAAGGCTGCCCAAATCCAGGTCCTGGTGTCGGTCGCGAAGGTGCGCCATTGAAACGCAACGGAGGCGTGAACAAG<br>TTCGCCAACTGCCCCGCTACCCCCATCGGCAAGGTCCTCGTCATCCAGGAGGACAAGAAGACGTGCCCTGACGACATTGGCA<br>AGCCAGGAGGATGCATCTATTTCATCTTCAGATCGTCCATGCACGGAGCCGTGAAGATAGAGATGGGTCGATTCCTGGACAT<br>CGACAACAATGAGGACCGCCCAAAGATGCAGTTCTACCACCCGAACCGCAGCAGCCCCAACCAGTCCCTCACCAGGTATGCG<br>GACAAGAGCGGCCACAACGGCCTCACCGACCTGTGGCTCAGTCACTCGAACATCTACAAGACCAGCGTGTCATACAACGGTT<br>CCGGATGCATCGCCGAGCTTCGTTTCCGTCCAAGCAAATGCCCCTCGCGCCGTTTGAGAGGACAGTCGGCGTGGGACAGCGA<br>TTTGGCCGAGGATGAGCTGGAGGCGGAGTTCAACGAGGAGATTGACGAGATTGACGAGATCGAGGAGGAGTGGATGATGGCG<br>AATTTGGCTTGA |
| **Protein** | MNVKTLLGSLVLSQIGSNAVHGMFDTVSDAGGSNSEPCPLDYQLLRSSDFQYSGAPIEIVSQDTSSVSFKVYNVWPQDGRPT<br>AEIYTEYSENNSIGGVNCEMFANVPANQDVGETFTAQCFQTKPKSVVTLYVRDPDIDKTIGTGKVPQCCHADLSIQYAVLKF<br>TYVLECVPTCEDPTPFSSCGDGTVDPGEECDDGNKINDDSCTNQCTSAKCGDGIMQSGEECDDGNKVDNDGCTNGCKLPKCG<br>DGFMQAGEECDDGNAVNNDGCTNGCKLPTCGDGIVQNGEECDDGNAVNNDGCTNTCKNAKCGDGIMQAGEECDDGNAVNNDG<br>CTNGCKLPTCGDGIVQNGEECDDGNNSNTDSCTNNCKNAKCGDGIMQAGEECDDGNAVNNDGCTNGCKLPTCGDGIVQNGEE<br>CDDGNNSNTDSCTNTCKNAKCGDGFMQAGEECDDGNAVNNDGCTNGCKLPTCGDGIVQNGEECDDGNNSNTDSCTNNCKNAK<br>CGDGFMQAGEECDDGNNSNTDGCTNGCKLPKCGDGIVQNGEECDDGNNNNNDNCTNNCEIPVPNTCGNGIVDPGEECDEGTR<br>NDYSGTRCKPDCTLPPGVCPLITLDFNDLPDGGFIYDEWWNTKGVKIIAASSSSRAYTPRPTSKCKTGELGCYCSFNPNYPL<br>QCSNWRSDKSYGAARVFDTSKPTGYRTSDSKYALYNNPMCDPRRSGDHGDPDLGSPNEGCPNPGPGVGREGAPLKRNGGVNK<br>FANCPATPIGKVLVIQEDKKTCPDDIGKPGGCIYFIFRSSMHGAVKIEMGRFLDIDNNEDRPKMQFYHPNRSSPNQSLTRYA<br>DKSGHNGLTDLWLSHSNIYKTSVSYNGSGCIAELRFRPSKCPSRRLRGQSAWDSDLAEDELEAEFNEEIDEIDEIEEEWMMA<br>NLA* |

**Table S4: *C. australis* PacBio genome assembly statistics:**

| | |
|---|---|
| Coverage | 54 |
| Total Length | 74,631,628 |
| Number of contigs | 88 |
| Contig N50 | 1,724,159 |
| Contig N90 | 411,438 |
| GC content | 52.93% |
| Shortest contig | 53,658 |
| Longest contig | 3,871,151 |

**Table S5: *C. australis* PacBio genome completeness statistics. Values shown are BUSCO (version 5.2.2) scores for stramenopiles ODB10 data set.**

| | |
|---|---|
| Complete BUSCOs | 97 |
| Complete and single-copy | 94 |
| Complete and duplicated | 3 |
| Fragmented BUSCOs | 2 |
| Missing BUSCOs | 1 |
| Total groups searched | 100 |
| % complete | 97.0 |

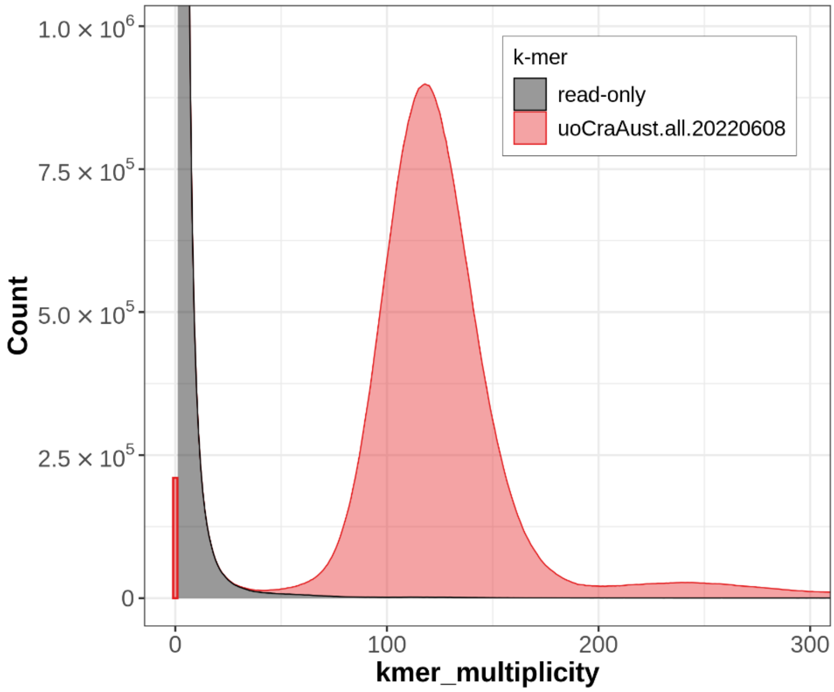

**Figure S2:** Merqury assembly spectrum plots for evaluating k-mer completeness.

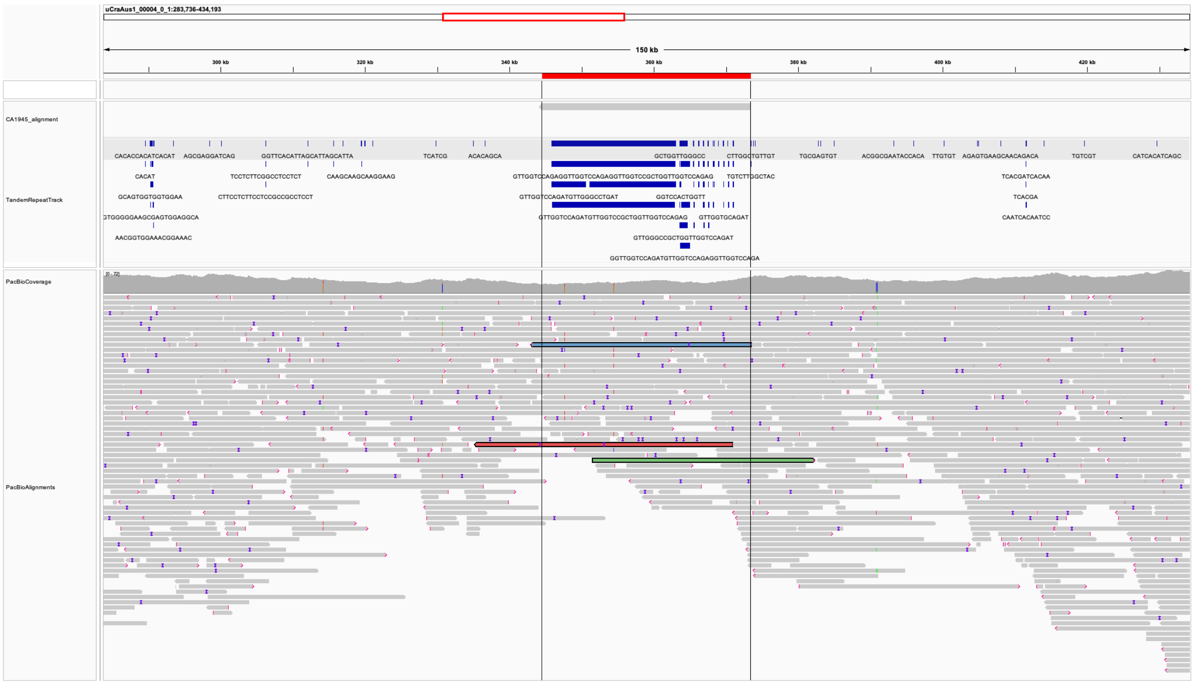

**Figure S3.** CaTrailin4 PacBio sequence alignment snapshot. Alignment snapshot of contig uCraAus1_00004_0_1 (283Kb – 434Kb) that includes the CaTrailin4 gene region highlighted in red and delimited by vertical black lines. Row CaTrailin4_alignment shows the alignment position of the CaTrailin4 gene sequence. The following row TandemRepeatTrack shows the tandem repeat annotation in blue blocks labelled with the tandem motifs that were reported by TRF. Rows PacBioCoverage and PacBioAlignments show the alignment coverage and the alignments of the raw PacBio reads that were mapped with minimap2.

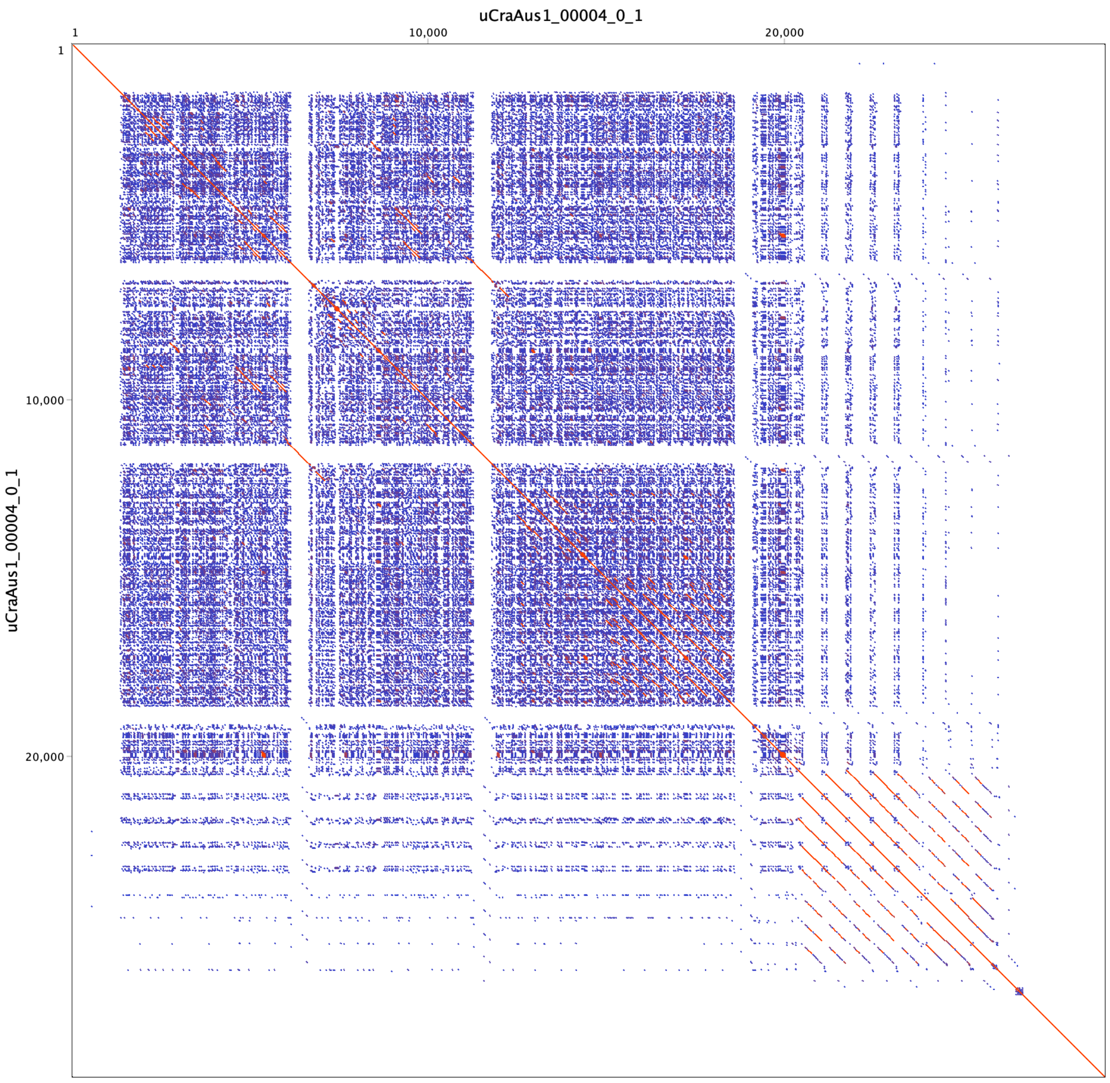

**Figure S4.** Dotplot: Dot plot of the CaTrailin4 region of contig uCraAus1_00004_0_1 (344kb – 373Kb). Created via Geneious (version 2020.0.5).

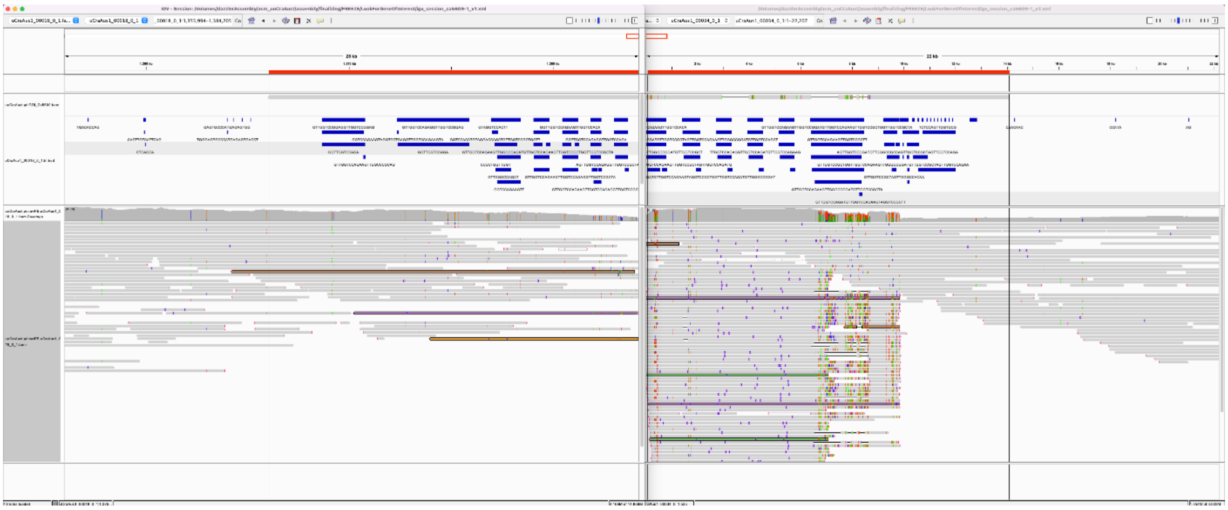

**Figure S5.** Ca5609 PacBio sequence alignment snapshot. Alignment snapshot of contig uCraAus1_00018_0_1 and uCraAus1_00034_0_1 that includes the Ca5609 gene region highlighted in red and is delimited by vertical black lines. The following row TandemRepeatTrack shows the tandem repeat annotation in blue blocks labelled with the tandem motifs that were reported by TRF. Rows PacBioCoverage and PacBioAlignments show the alignment coverage and the alignments of the raw PacBio reads that were mapped with minimap2.

**A.**

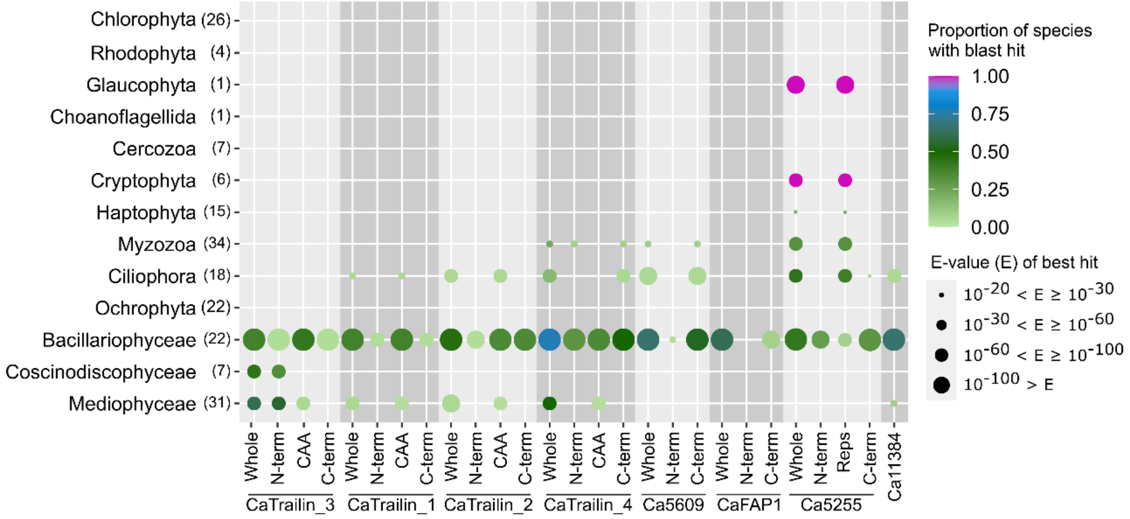

**B.**

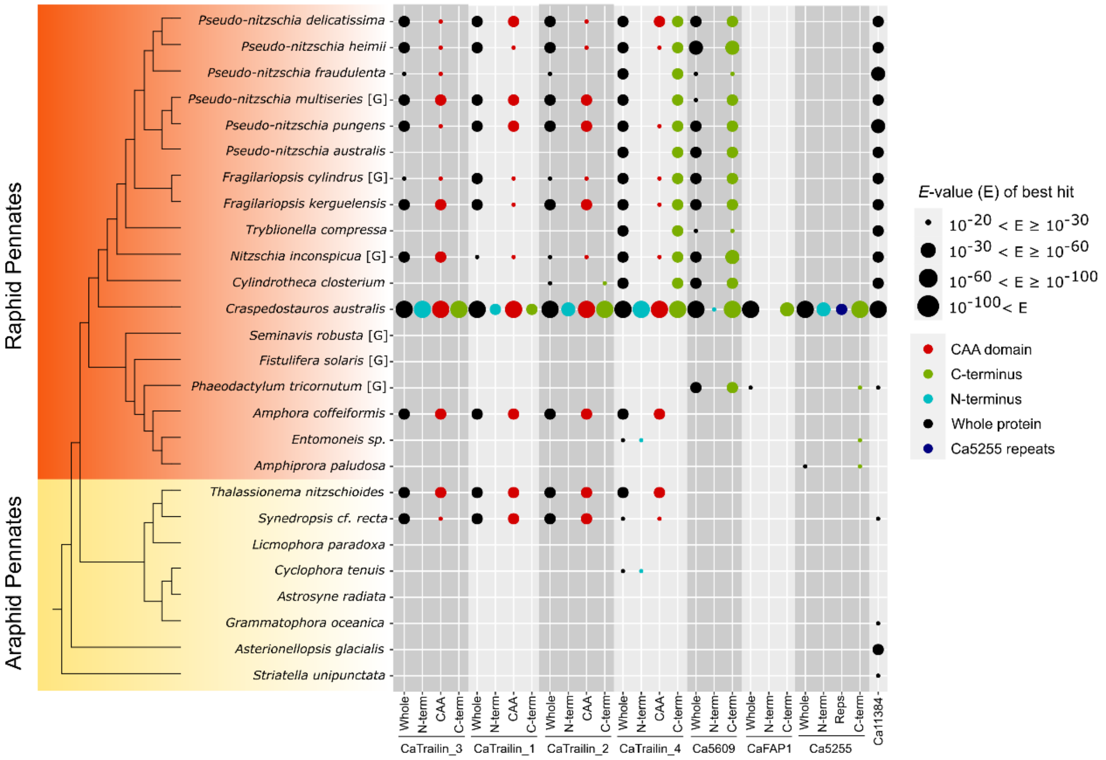

**Figure S6** (**A**) Distribution of blast hits of *C. australis* adhesive proteins across the Eukaryotic tree of life represented in the MMETSP transcriptomes. Either the entire protein sequence or partial sequence (non-repetitive N-terminal or C-terminal sections) were queried against the MMETSP transcriptome database (all other species) using tblastn with low complexity filtering. The size of the circles indicates e-value of the best blast hit, while the color indicates the proportion of species in the taxonomic group that contains a hit at a minimum e-value of 1E-20. The number of species in the taxonomic group is indicated next to the group name in brackets. (**B**) Distribution of blast hits of *C. australis* adhesive proteins across the Bacillariophyceae phylogenetic tree. Either the entire protein sequence or partial sequence (non-repetitive N-terminal or C-terminal sections) were queried against Phycosm genomic database (species labelled [G]) or the MMETSP transcriptome database (all other species) using tblastn with low complexity filtering. The size of the circles indicates e-value of the best blast hit.

**Table S6**. Clustal Omega sequence alignment of **(A)** CaTrailin3 Cysteine rich repeats, **(B)** Ca5255 Cysteine rich repeats, **(C)** CaTrailin4 NCVD repeats and Ca5609 QPTY-repeats. The consensus symbols represent: an * (asterix) fully conserved residue; a : (colon) indicates conservation between groups of strongly similar properties; a . (period) indicates conservation between groups of weakly similar properties.

| Panel | Repeat | Sequence | Position |
|---|---|---|---|
| **A.** CaTrailin3 Cys repeat | R1 | CGTGWMDTFNA-CMTSKP-CPGGQNSECADGQSCFGGINCSGNSYGSTDLPYSGPKF | |
| | R2 | CGSSWTDASNT-CASAAP-CPGGQDNECPDGLKCFSDINCATSTTYTPPVESSSYSGPNF | |
| | R3 | CGSSWTGAKNG-CATGAP-CPGGQNDECPDGESCFGGITCPTPTTYGAGSTSSFLGDSAGSSSSASSYTGPNF | |
| | R4 | CGSSWTGANDG-CATGAP-CPGAQDSECPDGQSCFGGITCPTPTTYGSTSTSAATSGYTGPNF | |
| | R5 | CGIDWMNAKNI-CAGGAP-CPGAQDSECPDGLKCFGGITCETPTTYGSTSTSTATSSYSGPNF | |
| | R6 | CGSSWTSANDG-CATGAP-CPGAQDSECPDGQSCFGGITCPTPTTYGAGSTSSFLDDTAGSSSSTSSYTGPNF | |
| | R7 | CGTSWTGAKNG-CATGAP-CPGAQDSECPDGQSCFGGITCPTPTTYGSTSAATSGYTAPNF | |
| | R8 | CGSSWTDANNN-CATGAP-CPGAQDSECPDGQSCFGGITCPTPTTYGSGSTSSFLGGSAGSSSASGYTAPNF | |
| | R9 | CGSSWTNANDN-CATGAP-CPGAQDSECPDGQSCFGGITCPTPTTYGAGSTSSFLDNTTPSYTGPSF | |
| | R10 | CGDSWSLANTN-CATGAP-CPGGTDGECPGGQSCFGGITCPTPTTYGTGAAPVPNLNF | |
| | R11 | CGPTYDDAEANKCGNGAQACPGGTADECPGSQDCFKVSACPGAAYLATEQESNASVKAVGSSPDASFGNPSSLSASL | |
| | | ** * *** ** ** * | |
| **B.** Ca5255 Cys repeat | R1 | CDDGNKINDDSCTNQCTSAK---CGDGIMQSGEE | |
| | R2 | CDDGNKVDNDGCTNGCKLPK---CGDGFMQAGEE | |
| | R3 | CDDGNAVNNDGCTNGCKLPT---CGDGIVQNGEE | |
| | R4 | CDDGNAVNNDGCTNTCKNAK---CGDGIMQAGEE | |
| | R5 | CDDGNAVNNDGCTNGCKLPT---CGDGIVQNGEE | |
| | R6 | CDDGNNSNTDSCTNNCKNAK---CGDGIMQAGEE | |
| | R7 | CDDGNAVNNDGCTNGCKLPT---CGDGIVQNGEE | |
| | R8 | CDDGNNSNTDSCTNTCKNAK---CGDGFMQAGEE | |
| | R9 | CDDGNAVNNDGCTNGCKLPT---CGDGIVQNGEE | |
| | R10 | CDDGNNSNTDSCTNNCKNAK---CGDGFMQAGEE | |
| | R1 | CDDGNNSNTDGCTNGCKLPK---CGDGIVQNGEE | |
| | R12 | CDDGNNNNNDNCTNNCEIPVPNTCGNGIVDPGEE | |
| | | ***** * *** * ** * *** | |
| **C.** CaTrailin4 DUF1 | NCVD_1 | NCVDPNGAKFTGSTDPTFTADPLPIVITFESDDGSTVEVDVVQNILNGEIKHVFIKYRNP | 60 |
| | NCVD_2 | NCVDPNGAKLTKSTDPTYTADPLPIVITFESDDGTSVEFDFVQNLSTDTIPHVFIKYRNQ | 60 |
| | NCVD_8 | NCVDPNGAVFKESTDPTYTADPLPIVLTFESDDGTSVEFDFVQNIINGVIPHVFIKYRNQ | 60 |
| | NCVD_9 | NCVDPNGAVFKESTDPTYTADPLPIVLTFESDDGTSVEFDFVQNIINGVIPHVFIKYRNQ | 60 |
| | NCVD_6 | NCVDPNGAVFKESTDPTYTADPLPIVLTFESDDGTSVEFDFVQNIINGVIPHIFIKYRNQ | 60 |
| | NCVD_7 | NCVDPNGAVFKESTDPTYTADPLPIVLTFESDDGTSVEFDFVQNIINGVIPHVFIKYRNQ | 60 |
| | NCVD_5 | NCVDPNGAIFKESTDPTYTADPLPIVLTFESDDGTSVEFDFVQNIINGVIPHIFIKYRNQ | 60 |
| | NCVD_3 | NCVDPNGAVFKESTDPTYTADPLPIVLTFESDDGTSVEFDFVQNIINGVIPHVFIKYRSQ | 60 |
| | NCVD_4 | NCVDPNGAVFKESTDPTYTADPLPIVLTFESDDGTSVEFDFVQNIINGVIPHVFIKYRSQ | 60 |
| | NCVD_10 | NCVDPDGAKLKGSTDPDYTADPKPIVLTFESDDGTLVEFDFVQNIINGVIPHVFIKYRNA | 60 |
| | NCVD_11 | NCVDPDGAKFVRSTDPSYTADPKPIILTFESDDATLVEFDFIQNIINGVIPHVFIKYRKS | 60 |
| | NCVD_12 | NCVDPDGAKFVKSTDASYTADPKPIILTFESDDATLVEFDFIQNIINGVIPHVFIQYRKS | 60 |
| | | *****:** : *** :**** **::******.: **.*.:**: .. * *:**:**. | |
|  | NCVD_1 | ESGMDDCAKHDDVEFGKKIPITAECYGQITTITIYIYTGEDFPAEECEACEIPD------ | 114 |
| | NCVD_2 | ESGDEECAQFDTVTLGTTLPIAAECFDGVTGVTIYIYTGEDFPSEECEACSIPD---EGD | 117 |
| | NCVD_8 | ETGTEDCAQFDTVGLGATYPLTAECFDGVTGVTIYIYTGEDFPSEECEACSIPD--GEGD | 118 |
| | NCVD_9 | ETGTEDCAQFDTVGLGATYPLTAECFDGVTGVTIYIYTGEDFPSEECEACSIPD--GEGD | 118 |
| | NCVD_6 | ETGTEDCAQFDTVGLGATYPLTAECFEGVTSVTIYIYTGEDFPSEECEACSIPD--GEGD | 118 |
| | NCVD_7 | ETGTEDCAQFDTVGLGATYPLTAECFEGVTSVTIYIYTGEDFPSEECEACSIPD--GEGD | 118 |
| | NCVD_5 | ETGTEDCAQFDNVETGATLPFAAECFSGVTGVTIYIYTGEDFPSEECEACSIPD---EGD | 117 |
| | NCVD_3 | ESGDEECAQFDTVTLGTTLPFAAECFDGVTGVTIYIYTGEDFPSEECEACSIPD---EGD | 117 |
| | NCVD_4 | ESGDEECAQFDTVTLGTTLPFAAECFEGVTGVTIYIYTGEDFPSEECEACSIPD---EGD | 117 |
| | NCVD_10 | ETGINECSEIMNAGEDSTFSYAAECIDQTTMVTIYIYTGDDFAGEDCEACSLPTDDGETG | 120 |
| | NCVD_11 | ETGINECAEFANVGENTPISLSAECVDMSTNVVIYIYTGDDFASENCEACSLPSDDGEGG | 120 |
| | NCVD_12 | ETGINECAEFANVGENTPISLSAECVDMSTNVVIYIYTGDDFASENCEACSLPSDDGEGG | 120 |
| | | *:* ::*:: . . :*** * :.******:** .*:****.:* | |
| | NCVD_1 | -GDPADFEAYSFEIPCDTECGEP | 136 |
| | NCVD_2 | SGNPDNFEAYYFEVPCQTECIDP | 140 |
| | NCVD_8 | SGNPDNFEAYYFEVPCQTECIDP | 141 |
| | NCVD_9 | SGNPDNFEAYYFEVPCQTECIDP | 141 |
| | NCVD_6 | SGNPDNFEAYYFEVPCQTECIDP | 141 |
| | NCVD_7 | SGNPDNFEAYYFEVPCQTECIDP | 141 |
| | NCVD_5 | SGNPDNFEAYYFEVPCQTECIDP | 140 |
| | NCVD_3 | SGNSDNFEAYHFEVPCQTECIDP | 140 |
| | NCVD_4 | SGNTDNFEAYYFEVPCQTECIDP | 140 |
| | NCVD_10 | SGNPDNFEAYYFELPCNTECIEP | 143 |
| | NCVD_11 | SGNPDNFAAYYFDVPCNTECIEP | 143 |
| | NCVD_12 | SGNPDNFAAYYFDVPCNTECIEP | 143 |
| | | *: :* ** *::**:*** :* | |

**D. Ca5609 DUF2**

```
Ca5609_R9       QPTYVCSTEARLVDKDGGNTKYDEGLDDEITGDDGIPFEVSEREVDSVKISLEPLLDAIE 60
Ca5609_R2       QPTYVCSTEARLVDKDGGNTKYDEGLDDEITGDDGIPFEVSEREVDSVKISLEPLLDAIE 60
Ca5609_R1       QPTYVCSTEARLVDKDGGNTKYDEGLDDEITGDDGIPFEVSEREVDSVKISLEPLLDAIE 60
Ca5609_R3       QPTYVCSTEARLVDKDGGNTKYDEGLDDEITGDDGIPFEVSEREVDSVKISLEPLLDAIE 60
Ca5609_R5       QPTYVCSTEARLVDKDGGNTKYDEGLDDEITGDDGIPFEVSEREVDSVKISLEPLLDAIE 60
Ca5609_R4       QPTYVCSTEARLVDKDGGNTKYDEGLDDEITGDDGIPFEVSEREVDSVKISLEPLLDAIE 60
Ca5609_R6       -------------------------------------FEVSEREVDSVKISLEPLLDAIE 23
Ca5609_R7       QPTYVCSTEARLVDKDGGNTKYDEGLDDEITGDDGIPFEVSEREVDSVKISLEPLLDAIE 60
Ca5609_R8       QPTYVCSTEARLVDKDGGNTKYDEGLDDEITGDDGIPFEVSEREVDSVKISLEPLLDAIE 60
Ca5609_R10      QPTYVCSTEARLVDKDGGNTKYDEGLDDEITGDDGIPFEVSEREVDSVKISLEPLLDAIE 60
Ca5609_R11      QPTYVCSTEARLVDKDGGNTKYDEGLDDEITGDDGIPFEVSEREVDSVKISLEPLLDAIE 60
                ************************************************************

Ca5609_R9       ALRTGGAVGEDGELGSVAVVFDRISTDNDGNEVTENDFCTETSVTEATGPAEEYVAQCVE 120
Ca5609_R2       ALRTGGAVGEDGELGSVAVVFDRISTDNDGNEVTENDFCTETSVTEATGPAEEYVAQCVE 120
Ca5609_R1       ALRTGGAVGEDGELGSVAVVFDRISTDNDGNEVTENDFCTETSVTEATGPAEEYVAQCVE 120
Ca5609_R3       ALRTGGAVGEDGELGSVAVVFDRISTDNDGNEVTENDFCTETSVTEATGPAEEYVAQCVE 120
Ca5609_R5       ALRTGGAVGEDGELGSVAVVFDRISTDNDGNEVTENDFCTETSVTEATGPAEEYVAQCVE 120
Ca5609_R4       ALRTGGAVGEDGELGSVAVVFDRISTDNDGNEVTENDFCTETSVTEATGPAEEYVAQCVE 120
Ca5609_R6       ALRTGGAVGEDGELGSVAVVFDRISTDNDGNEVTENDFCTETSVTEATGPAEEYVAQCVE 83
Ca5609_R7       ALRTGGAVGEDGELGSVAVVFDRISTDNDGNEVTENDFCTETSVTEATGPAEEYVAQCVE 120
Ca5609_R8       ALRTGGAVGEDGELGSVAVVFDRISTDNDGNEVTENDFCTETSVTEATGPAEEYVAQCVE 120
Ca5609_R10      ALRTGGAVGEDGELGSVAVVFDRISTDNDGNEVTENDFCTETSVTEATGPAEEYVAQCVE 120
Ca5609_R11      ALRTGGAVGEDGELGSVAVVFDRISTDNDGNEVTENDFCTETSVTEATGPAEEYVAQCVE 120
                ************************************************************

Ca5609_R9       DDDFAGTTVAIVYVVVADSANTQEDPSGSLIGAAGGESALGTSGCGVPFESGPFSTPVIV 180
Ca5609_R2       DDDFAGTTVAIVYVVVADSANTQEDPSGSLIGAAGGESALGTSGCGVPFESGPSSTPVIV 180
Ca5609_R1       DDDFAGTTVAIVYVVVADSANTQEDPSGSLIGAAGGESALGTSGCGVPFESGPSSTPVIV 180
Ca5609_R3       DDDFAGTTVAIVYVVVADSANTQEDPSGSLIGAAGGESALGTSGCGVPFESGPSSTPVIV 180
Ca5609_R5       DDDFAGTTVAIVYVVVADSANTQEDPSGSLIGAAGGESALGTSGCGVPFESGPSSTPVIV 180
Ca5609_R4       DDDFAGTTVAIVYVVVADSANTQEDPSGSLIGAAGGESALGTSGCGVPFESGPSSTPVIV 180
Ca5609_R6       DDDFAGTTVAIVYVVVADSANTQEDPSGSLIGAAGGESALGTSGCGVPFESGPSSTPVIV 143
Ca5609_R7       DDDFAGTTVAIVYVVVADSANTQEDPSGSLIGAAGGESALGTSGCGVPFESGPSSTPVIV 180
Ca5609_R8       DDDFAGTTVAIVYVVVADSANTQEDPSGSLIGAAGGESALGTSGCGVPFESGPSSTPVIV 180
Ca5609_R10      DDDFAGTTVAIVYVVVADSANTQEDPSGSLIGAAGGESALGTSGCGVPFESGPSSTPVIV 180
Ca5609_R11      DDDFAGTTVAIVYVVVADSANTQEDPSGSLIGAAGGESALGTSGCGVPFESGPSSTPVIV 180
                ***************************************************** ******

Ca5609_R9       YKYVVPCQYSCEGSPTSVPTDAPTGG  206
Ca5609_R2       YKYVVPCQYSCEGSPTSVPTDAPTGG  206
Ca5609_R1       YKYVVPCQYSCEGSPTSVPTDAPTGG  206
Ca5609_R3       YKYVVPCQYSCEGSPTSVPTDAPTGG  206
Ca5609_R5       YKYVVPCQYSCEGSPTSVPTDAPTGG  206
Ca5609_R4       YKYVVPCQYSCEGSPTSVPTDAPTGG  206
Ca5609_R6       YKYVVPCQYSCEGSPTSVPTDAPTGG  169
Ca5609_R7       YKYVVPCQYSCEGSPTSVPTDAPTGG  206
Ca5609_R8       YKYVVPCQYSCEGSPTSVPTDAPTGG  206
Ca5609_R10      YKYVVPCQYSCEGSPTSVPTDAPTGG  206
Ca5609_R11      YKYVVPCQYSCEGSPTSVPTDAPTGG  206
                **************************
```

**Table S7.** BLASTp alignments of *C. australis* adhesive proteins highlighting regions of sequence similarity

**CaTrailin3**

>TPX53846.1 chitinase [Chytriomyces confervae]
MLSSRAILVLLALTQILAMPTLPLINARDGASTRCGKSWADANGRCGQACPSGQNSQCSNGETCFKDLAT
ASCGGGSPAPAPAPGNTGGSTRCGKSWADANGKCGQSCPTGQNSQCSNGETCFKDLATASCGGGSPAPAP
APAPAPAPGNTGGSARCGKSWADANGKCGQSCPGGQNSECSNGETCFKDLATAPCGGSSPAPAPAPAPAP
APGNTGGSTRCGKSWADANGKCGQSCPGGQNSECSNGETCFKDLATASCGGGSPAPAPAPAPAPGNTG
GSTRCGKSWADANGKCGQSCPGGQNSECSNGETCFKDLATASCGGSSPAPAPAPAPAPGNTGGSIRCGKS
WADANGKCGQSCPGGQNSECSNGETCFKDLATASCGNSPAPGPAPAPGPQPNDPSNPTSIQSLISESKFN
SALQTCGISKSGLYQSLVKGFTAPLANLRELALLVGNTAHESGAFIYVEEIACAGVTSPTGNCPYGLYHG
RGYIQLSWDYNYRAAASALNRPDIFSNPWVVQQDEATNWSTVQWYWTTAVQPALKANGYTLAASVRAING
GLECGGNPIAAKRIQFVHCFEQQFTGTQSGQTSC

Yellow: region with sequence homology to CaTrailin3
Grey: chitinase domain

>WP_017559785.1 choice-of-anchor A family protein [Nocardiopsis baichengensis]
MGTNLDVASDRKRRSTGRRAVVLLGTGAAGAAVLAGALALSSVAATLPGGLGPCVPGDCPDPYPEVDNGP
IAGRDNGINVFVGGDMLVREGAAEAEGKVVVLGDFDMDKREGASQIYNVGVAGVGSRVPPDDGTDFLTAG
GNVTVADGQRVLAEEGSVHGVVRYGGELQGTVVPEEVQDPDAAAPYADLRSELAAASDCYAYPHGEQREA
TGTADNQGHTTVFTGDGTSSLQVFNVDFDLTNTDGGQQGFSFVDIPTDATVLINVTGDRRTINTYMGELP
DGLRERILWNFPDATEVNLKGTGQFQGSVLVGEPGSTTRLSLPGTNGRFFTTGTLSHESDGGSGGQELHA
YPFNGDLPDCRNLGPSPTPTPSPSPSVSPTPSPEPTPSPTLEPSPSPSPEPSPSPEPSPSPEPSPTPTPE
PTDSPGPTPEPTVSPEPTDSPGPTDEPTDGPTPEPTDSPEPTDGPTDGPTPGPDGGGGDGGHGDAGDDAP
APDRGGLPLTGPEIAGLVAAALVLLAFGGLAVYASWRRRADGDGPG

Grey: CAA domain
Underlined: region with sequence homology to CaTrailin3

**CaTrailin1**

>WP_084461253.1 choice-of-anchor A family protein [Curvibacter gracilis]
MTAAFSHPGFPILLSLSALLSTPSYAGSGLGAETGPCLGPDCPSHYGPPAACRKLGAEGRDEAYSLVVGG
HFHVLQGSEAEGRIFVGGDLVLRTRGYYNLVQVGAGACVVPPDADQASSPPHLVVGGQIQPVDPSAVLAV
GQPNLRSTALIGGARVGAGQVQARGGVRYQTSPTPPPVDFNSLAQRSAYWATLAPTGTVRGQHPMVLQGD
GQSALQVFKLSADLPSGGLRLKGIPPGATVLINHPGTPTVSLQFYDMTDPAGHGGFQFDTELTRRMLWNF
PQASLVKLGGGAQWQGSVLVARGALYNALPGLNGRVLVAGDLTQDSQGSEFHNYDFKGELPDPPTDDPPA
TEPGHDTSPPEPPPEVPPEPPPESPPPGLMQAALNIQVRYDGDALRITDAPLLQIESQCERSGTQSVRIT
PPAAGWIEGLQTDESCLLVAKLERAAQLPRGWQWPSPQPGLWVGDNPRVITPDTNPVELLLQIRPVMNAS
LRVLPRYLGNHRAVQTHPRIELNLLCSHSGEQSLLLQAPQVAQSGLLQDGEACEVLARRITGAVLQEGHT
LSLPDTGWWSPGSTLVAGDTPDPIPLDISILAASPGDESEPPPAPPPVPPSESGPSTLNSIPLLTPWGLL
GLASGLGLLSLGTWMGPAIDRRKRRTASDKVNESRGCAGSSRPGSGPAAH

Grey: CAA domain
Underlined: region with sequence homology to CaTrailin1

**CaTrailin2**

>WP_084461253.1 choice-of-anchor A family protein [Curvibacter gracilis]
MTAAFSHPGFPILLSLSALLSTPSYAGSGLGAETGPCLGPDCPSHYGPPAACRKLGAEGRDEAYSLVVGG
HFHVLQGSEAEGRIFVGGDLVLRTRGYYNLVQVGAGACVVPPDADQASSPPHLVVGGQIQPVDPSAVLAV
GQPNLRSTALIGGARVGAGQVQARGGVRYQTSPTPPPVDFNSLAQRSAYWATLAPTGTVRGQHPMVLQGD
GQSALQVFKLSADLPSGGLRLKGIPPGATVLINHPGTPTVSLQFYDMTDPAGHGGFQFDTELTRRMLWNF
PQASLVKLGGGAQWQGSVLVARGALYNALPGLNGRVLVAGDLTQDSQGSEFHNYDFKGELPDPPTDDPPA
TEPGHDTSPPEPPPEVPPEPPPESPPPGLMQAALNIQVRYDGDALRITDAPLLQIESQCERSGTQSVRIT
PPAAGWIEGLQTDESCLLVAKLERAAQLPRGWQWPSPQPGLWVGDNPRVITPDTNPVELLLQIRPVMNAS
LRVLPRYLGNHRAVQTHPRIELNLLCSHSGEQSLLLQAPQVAQSGLLQDGEACEVLARRITGAVLQEGHT
LSLPDTGWWSPGSTLVAGDTPDPIPLDISILAASPGDESEPPPAPPPVPPSESGPSTLNSIPLLTPWGLL
GLASGLGLLSLGTWMGPAIDRRKRRTASDKVNESRGCAGSSRPGSGPAAH

Grey: CAA domain
Underlined: region with sequence homology to CaTrailin2

**Ca5255**

>KAG7347470.1 cold-active serine alkaline protease [Nitzschia inconspicua]
*MRHRSFFPFWLLSLHSSRYLVVHG*EKSRKSNLRNSKHSSRIPATNGVVNFLPFLVENAGRTLGNGRKLHK
NDSDVGTEEQKGKRMSTQARIGNGFERGHNRSIQQEKPLLNKRKVEVRNLTFQRCRNGGTQFVVQCTTGS
EEHCYRELARANAVIVNELPNSDFFVVCVHTAEEKKLLNELTDVVDMEEDCIRTLSYLPELTKPVDRREL
QSGQQIPYGATMVNAPQFWQTKGDKGGSAKVCIIDTGLNIGHEDIQGAVFSGSTDSSVVADWDSDDAGHG
THVAGTIAAVDNNIGVVGVAPEAELLIVKVFEGANSQFTASSLVSALEECRKGGANIINMSLGGPSSSVV
ERNKVNQLANQGIQLIAASGNSGDTSNPIEYPASYDKVISVAAVDEDRHIAIFSTHNNEVDVAAPGVDIL
SLTNACSTCYGLYSGTSMATPHVAGVFALLMSKYPTKSISQIREAIQESASDSGACGIDRMFGHGIVDVM
AAAVYLESGVSASEQNNCINTKITVTTDKWGSETSYVIRNSDGEVVYKNGPYSNQIKTYTDVFQLPDDCY
EFELLDSYGDGICCEEGSGSFKVEYDGVEEVYNDSFLNGNSVKASFGCGSGGDVGSGSPPSCGNGILEDI
EECDDGNQNNNDSCTNECKIAVCGDGIVGPGEQCDDGNNVNDDSCRNDCTSALCGDGVVQSGEECDDGNN
SNNDTCTNACKNARCGDGFLQPGEECDDGNGDNGDGCSSDCKVETISTSCQAGEAEVELLFQSDAYSYNE
NELYFYQSTDQTSVGDIDFIWIGTKQGIESNKKYEMSACVDETKCYKFFFFDSYGDGLWGNDGLRLSWNG
VEVLSVAPYEVGPKWDGGPTIYWTEDLGACS

*italics*: signal peptide
red: peptidase domain
Green: Myxococcus cysteine-rich repeat
Underlined: region with sequence homology to Ca5255

>MCB0319749.1 DUF4215 domain-containing protein [Bdellovibrionales bacterium]
*MMKNAFSRLSVRLVAFFCLGLICTSSA*RAQTYPPECQPLLDGSVLFVAAECLNDSSALISDSTTSDSNKW
QYSFDSNSDGVNGSNVGGTAYEIYGIAIRELNTEVWVVINSNIPVTGVPSASAAGGSVSWGDLFWNFTGN
DFEAADTTSNHFAIRFVTSNESGVPSLGVYREVSAKSVTAQNVGYASWDVYAAFVASHGQDASLGDFGLN
QTYYGGHLSLNVIDIGTFVGPITFLSPAELLSAGFDDSLFDGTTTIAFRFPKSYLAQYPDTDGDGQDDCI
DPCGNGKIESPEECDDGNFVNTDECSNLCTLPRCGDAIVQSAKGEECDDGNQVNDDSCSNACTNPKCGDG
ILQSGEQCDDGNTNNTDSCSNGCTTAVCGDGIVQTDEQCDDGNQIDSDECTNLCTTPKCGDGILQAGEEC
DDGNTSDSDSCTSTCTNARCGDGILQDGEECDDGNTSNSDSCSNTCTTPVCGDEIIQSGEECDDGNSVDD
DGCTNSCALPKCGDGILQGGEQCDDGNSNNSDSCTNACTTPKCGDGVVQAGESCDDGNGNNSDACTTSCE
VAACGDGFVQPGELCDDGNTIIDDECTNQCTLPNCGDGIVQSGEECDDGNSVNSDTCTNVCTVANCGDGI
KADSEQCDDGNQINTDACSNDCTLPVCGDGIIQGGELCDDGNLVDDDDCTSFCTLPTCGDGIVQAGEECD
DGNSVNNDSCTDQCTSARCGDGIVQAPEECDDGNEVDNDSCSVSCKVIDVVPEDCKEYNIEPSQFILDQG
VKLQEAFINLQILKKYVRLGGSKKYVADARAEAHTLQTEGWVLSWSVASQGVICTSSTENCVTSTANGDM
LAQYTERTEGLFDLTKQALRKLRKVGGSKKLIRTLRKRALKLLNRNLSERDKIPTNSTECTTS

*italics*: signal peptide
Green: Myxococcus cysteine-rich repeat
Underlined: region with sequence homology to Ca5255

**Ca11384**

>MMETSP1070 (*Minutocellus polymorphus)*
VTFEYIVADGNGATDSGIVTLTLFGQDTFAPSSSPTELFVCPDQTVNAANQGTISGSNVSGGTTSQTLQFDLF
ASSFTSNGMACVDGLIGIKPPYNILFVIDVSGSTRNNFSGTAVGDVNGDGRSNTILDAQIDTVLKAIEAIINT
PSLTNDNVNIGIVTFSTSGNYVGNWSPTDENNPDAIDPALETTLKSLRSGGYTNFDDALDKAIVYFEGSDSSK
GGAPDVSSRTNRMYFLSDGLPNQCGDGDPNTAEDYCGEDEIYNDAAGATVFASELQSLELYSVAIYAIGVGEA
SDVSAGSGLDKIDNTENPFTGEKAVQVTTTDALTSNILESPVYAEVFDFKLKVNGAIVPGVDETSLVTDISGY
VLGVNEISGLDPTNGASNTIIATVLLDFDGDSATPDDQLEISTIVSIIGAAGLPV*

Teal: von Willebrand factor type A (vWFA) domain
Underlined: region with sequence homology to Ca11384

>VEU43960.1 unnamed protein product [Pseudo-nitzschia multistriata]
MNRFFLSIAAASLALVSADVNFKEMHERKLAMEEDHRVMEEYLSTKSGTKSGTKSDTKSGRQLDASFFQT
PSPSASPSKVPTQTPSASPTTSEPSKQPTTSPSAEPSSIPSRLSEEYGGLPWPSASPSNFYEPAPKCGTI
ETKDGSERLLGPLPDENYYCNGDSYAWYRRTFTHYDYEFELSQYCEYKFSSPETGSSVIMRDVNHWFEQA
ATSLKGWDEVGYKASPEEYVFSGYAVPEFDDIEYESDIADAECPLNYCIRITGPSVGLNTNEELSDGSAE
PYLLYAFSSWEGYSTFEGFLSSTMYSWNAHHIFMTNHYLKCSCPSFCIDEEFYVQDFKPLATCYVEDICG
VKTFQIAFSDDYPETQRVSWENTGTTHVMEWFSECANVMHGCPQLASDFSPARFSGVGHFSSIRGPDVYT
YDEYESATNVYGFGGWRGFGWDYYSQPALLWECPSPEHLERLVEFGDLISDYSFFRHDSIDTSVPETCPE
PPLIEPEDPIVFDPEGPPCTEVENEGLFCFELKHATCYIRDTHECSEFEFVIEGVLDGAEDMWVTTDVEV
WDSWCHFDGIGCGIPSYWDEQPPYTYENSPWAFVSWSDYQHQWLEFQSGSSGRPSAIGGGNSYRLWNCDD
ATQAYLARDFTDAVAASSSGDLYQVPPLDCTESPTASPSMGPTSSIQPSNSPTMSPTICGESYLNSESLN
IDLAIDLSYSTYLYLFSSEVDIGDVNGDGKANTILDAQVQAIEELLVAILESETLGNRNCEINLISFHTD
ATNHGTFLPLSEDESSINTRLMHYIKTELRAPTSDLEVQLTNNGFTNFDAALDLAGDYFEYEATPDRLNL
LVFLSDGEPNVRGDGDDEGYCADTAGVWNSNPTNGFPAEVQCADLDLEAGVRHTFCRADDPECVPRNPYQ
ECVRGMTKCINSPAVTQYESEINRLTELRVERLAIGVGDASNVAEGSALWMIDNNPGKDLGVLPIQALDL
EALSNALKSLCILNTDPPTAEPSHQPSDQPTSSTAPSVIPTDVPSSEPTDVPSSSPTEEPEFTIETPSPT
KSPTQTPTKTPTKTPTASPSQEPSTSEPTGSPTAFPTRLPSASPTSEPTTAEPSAAPSQYPSGSFYPSSA
PTESPTESPAPSQSPTDAPTTSPTVSPTVSPTLSPTASPTKSPTASPTSSPTNKPTEIPTTSPTDSPTSS
VAPTFLLPECYDYPKLVKKDSSDTGICFFSEDMIFIEQQNTTEVGLRINNVWSKTLPQDVILFSHSNGVN
SVRGGNGFECDSEDGNAIDLAGSDEIMVQCHSEGDGEPYLAVVDLMIIDSNIPVNDVNHPCNPNEVIPNA
CSWRMVIPCEEDVMCTPEPTDSPSVYPTYVPTFGESTESPTTLTTFHSESPTTGTTFHEKTESPTHELDR
HTNDDESNDIVYESECPEDIVLLEQEGVTEFPEGGLYVVGREDGTVTVKLTQTFTDATVDSLFYQYQIGR
FSNKCFEDKDVPVTDSVDITILCTEHSKIALLELWVVDALEHNLLSESDNSVVPECCHPTHPEGTPATKY
LFEIKCLTVCADAVE

Teal: von Willebrand factor type A (vWFA) domain
Underlined: region with sequence homology to Ca11384

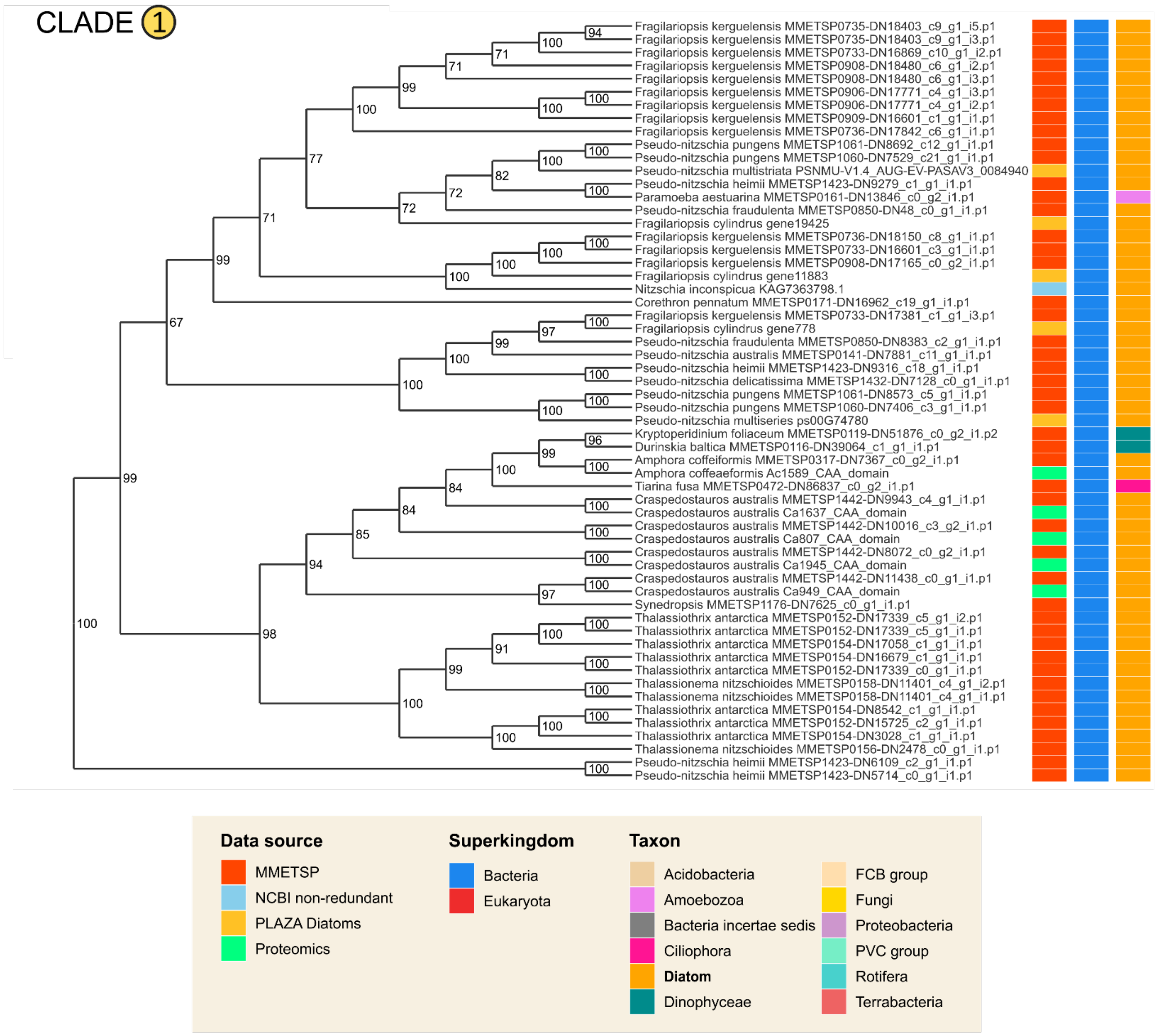

**Figure S7.** Phylogeny of homologs of the Diatom CAA-like domain belonging to Clade 1. For each hit, the species name and gene identifier is shown, as well as coloured boxes indicating the data source and taxonomic position.

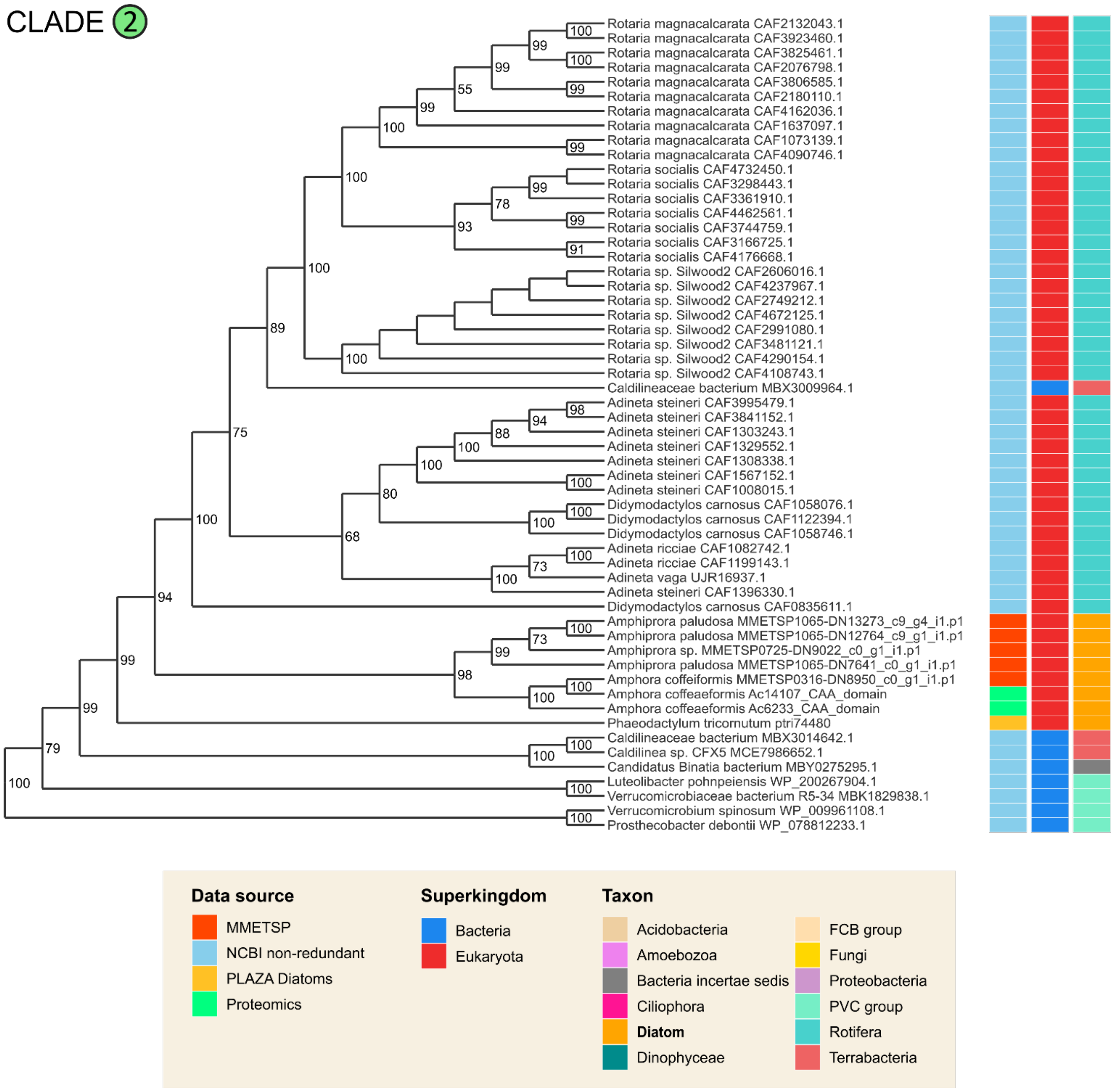

**Figure S8.** Phylogeny of homologs of the Diatom CAA-like domain belonging to Clade 2. For each hit, the species name and gene identifier is shown, as well as colored boxes indicating the data source and taxonomic position.

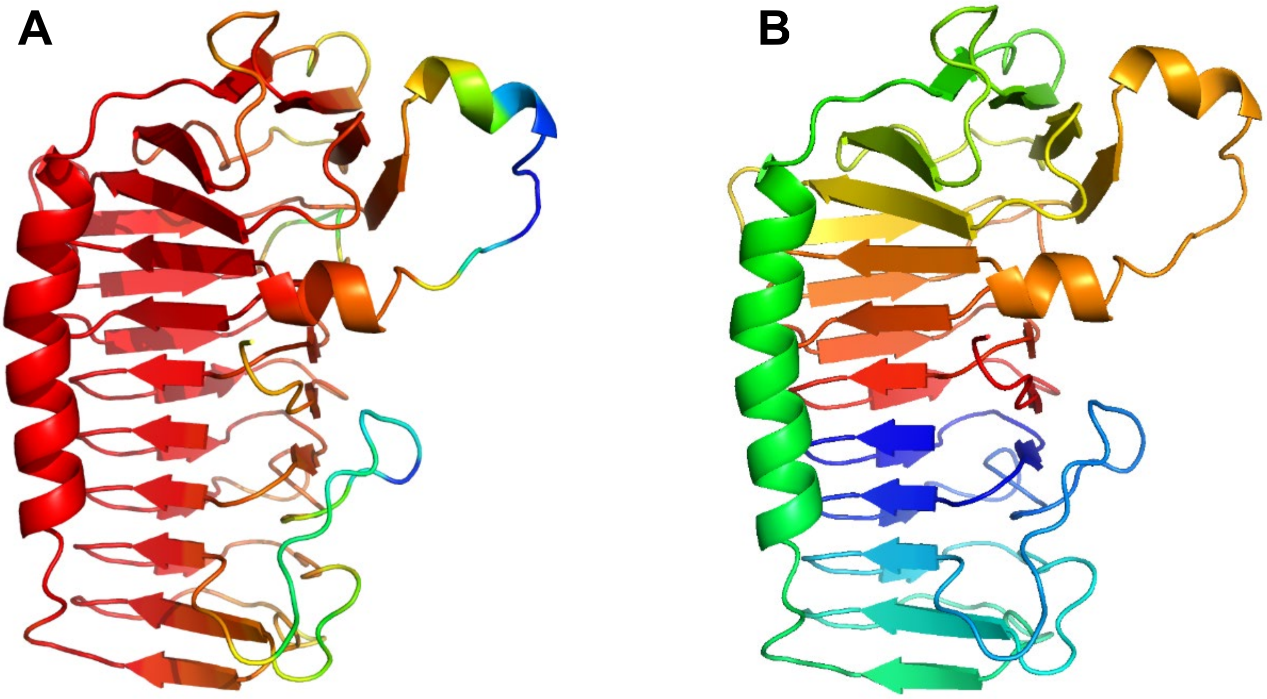

**Figure S9.** Alphafold structural prediction of the CaTrailin4 CAA domain. **(A)** Quality of CaTrailin4 model. Red indicates high quality and blue low quality. **(B)** CaTrailin4 CAA domain model consists of two beta helical cores (red/orange and blue, respectively) held together by an alpha helix (green).

**Table S8.** The top eight predictions from the structural homology search

| Name | Species | | Uniprot ID | Uniprot Accession | PDB ID | Chain | Method | Resolution | RMSD | Aligned residues | Ice-binding |
|---|---|---|---|---|---|---|---|---|---|---|---|
| FfIBP | *Flavobacterium frigoris* | Bacterium | IBP_FLAFP | H7FWB6, I3QNZ8 | 4NU2 | A | X-ray | 2.10A | 5,18 | 85% | Yes |
| ComZ | *Thermus thermophilus* | Bacterium | Q72JC1_THET2 | Q72JC1 | 6QVI | A | X-ray | 2.72A | 5,49 | 80% | No |
| AFP | *Leucosporidium* sp. (Arctic yeast) | Fungus | IBP_LEUSY | C7F6X3 | 3UYU | A | X-ray | 1.57A | 4,73 | 80% | Yes |
| K1-A | *Typhula ishikariensis* (Gray snow mold fungus) | Fungus | IBPKA_TYPIS | Q76CE8 | 5B5H | A | X-ray | 1.00A | 5,06 | 75% | Yes |
| ColAFP | *Colwellia* sp. | Bacteria | IBP_COLSX | A5XB26 | 3WP9 | A | X-ray | 1.60A | 5,82 | 73% | Yes |
| K3-B1 | *Typhula ishikariensis* (Gray snow mold fungus) | Fungus | IBPKB_TYPIS | Q76CE6 | 3VN3 | A | X-ray | 0.95A | 4,97 | 72% | Yes |
| fcIBP | *Fragilariopsis cylindrus* | Diatom | D0FHA3_9STRA | D0FHA3 | 6A8K | A | X-ray | 1.40A | 5,27 | 72% | Yes |
| Anp | *Antarctomyces psychrotrophicus* | Fungus | A0A2Z6DSM4_ANTPS | A0A2Z6DSM4 | 7BWX | A | X-ray | 1.90A | 4,21 | 70% | Yes |

```
              SS                                               S   SSS    SSS
CA1945        FL-----------------------------------------------VGGTYTSIQSAEIE
FfIBP         SLSVANSTYETTALNSQKSSTDQPNSGSKSGQTLDLVNLGVAANFAILSKTGITDVYKSAITG
                                                                  SSSS   SSS

                     SSS   SS                            SSS   SS           S

CA1945        GK-----IVIGGDFITTGDQSINSVVFAGVGSCIIPEGDQDVFLVGGNMVINRSIVLVNPSNA
FfIBP         DVGASPITGAAILLK----CDEVTGTI---------------FSVDAAGP------------A
              SS     HHHHH  H    HH SS   S               SS

              SSSS                        HHHHHHHHHHHHHH  HHH    S    SSSS
CA1945        YVRYGGTLSGSNPNGLGGIVQHTPNLDLTYYTDVIEALKIKSAYWATLPANG----VYTDRTG
FfIBP         CKITDAS-------------------RLTTAVGDMQIAYDNA--AGRLNPDFLNLGAGTIGGK
                SSHHH                   HHHHHHHHHHHHHHHH  HH     S    HHH

                              SSSSSS  S             SSS     SSSSSSS     SSSS
CA1945        RNYVEFIAGAAPNDGCLQVFDL--DYDDLYFGDWGTEVFFKNLSGKTMLFNVGSDSNGHVKIA
FfIBP         T-------------LTPGLYKWTSTLNIPTDI------TISGSSTDVWIFQVAG----NLNMS
              S             S  SSSSS   SSS   S      SSS      SSSSS        SSS

                 SSSSS HH              HHHHSSSS     SSSS            S SS   SS
CA1945        NLAKFYDGAKSDQASLGVDKAFSKDMAAGMIWNFYDATSVEIGGGLNGNGEFIGTILAPFADL
FfIBP         SAVRITLAGGA--------------QAKNIFWQ--TAGAVTLG----STSHFEGN-ILSQTGI
                  SSSS                 HHH SSSS  S   SSS        SSS S SSS   S

              S S   SSS  SSS   S SS    SS
CA1945        E-FTAPGQSGRTIVGGDVTHDAYGSEFHNIDFD
FfIBP         NMKTAASINGRMMAQTAV-TLQMNTVTIPQ---
              S     SSS  SSS   S SSS  SSS
```

**Figure S10.** Sequence representation of structural alignment of the CaTrailin4 CAA domain and FfIBP. Structurally aligned residues are colored in red, sequence identity is highlighted in bold. Identical secondary structure (S for strand and H for helix) is emphasized in bold. Three ice-binding motifs (T-A/G-X-T/N) are underlined in the sequence of FfIBP.

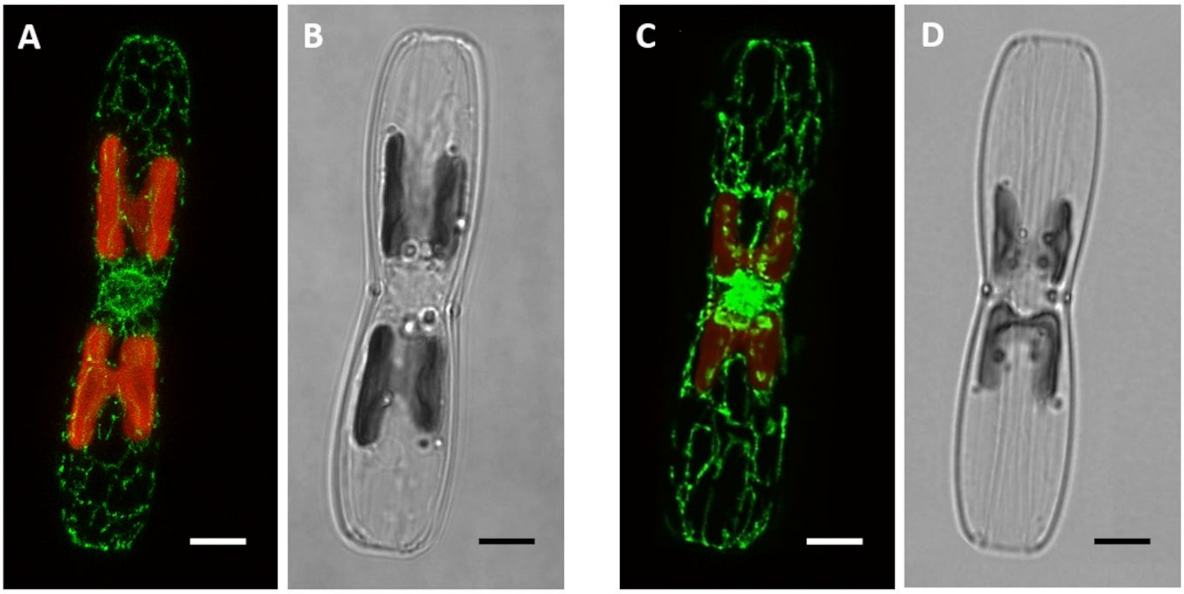

**Figure S11.** Confocal microscopy (z-projection) images of cell lines expressing the AM proteins CaTrailin2-GFP and CaTrailin3-GFP transformants. Localization of CaTrailin2-GFP **(A)** and CaTrailin3-GFP **(C)** is shown in green, whereas the red color depicts chloroplast autofluorescence. **(B, D)** The corresponding light micrographs. Scale bar: 5 μm.

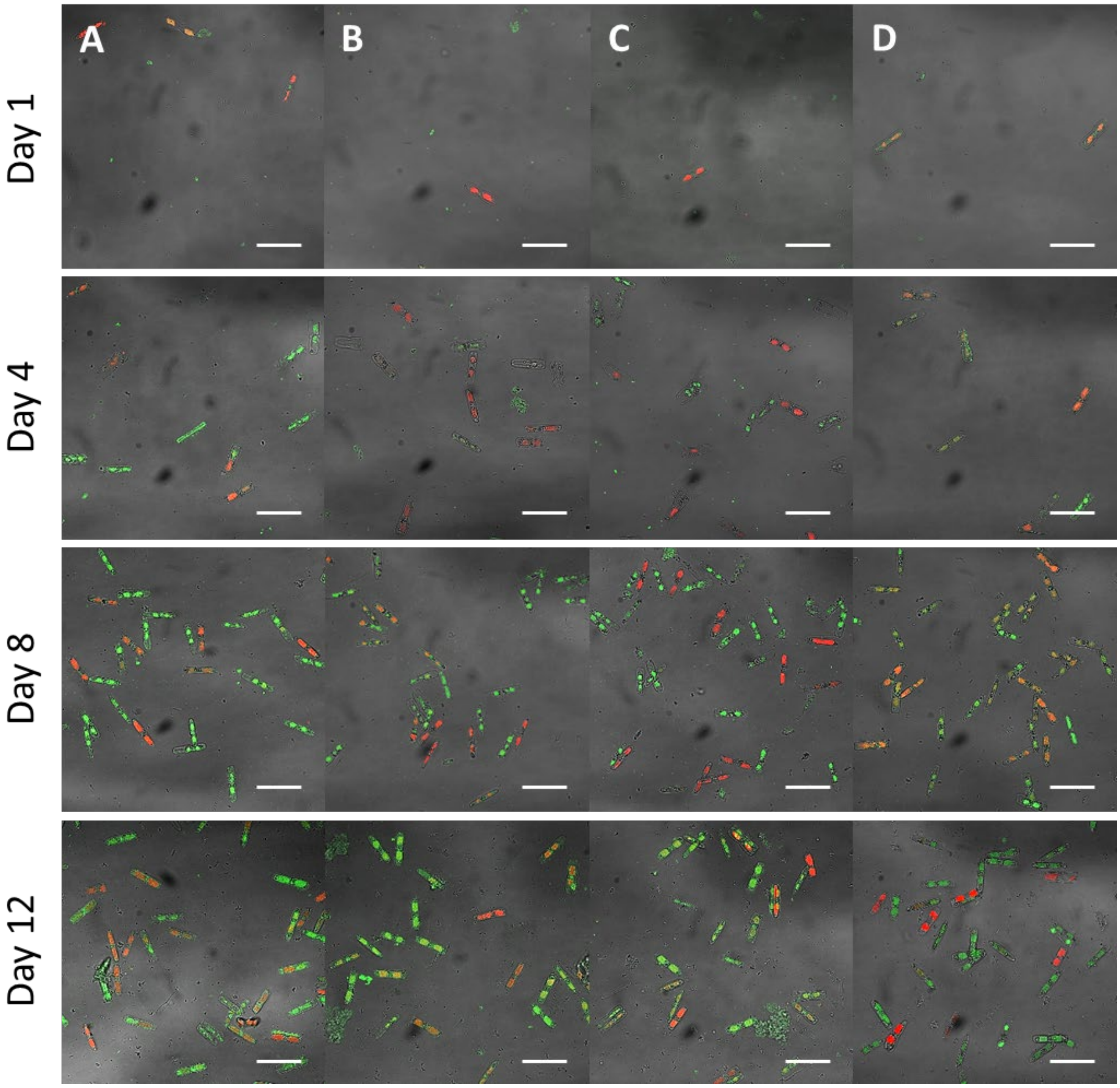

**Figure S12. Immunofluorescence control experiments**. Confocal microscopy images of *C. australis* adhesive material after 1, 4, 8 and 12 days. Samples were incubated with IgG purified pre-immune rabbit serum substituted for primary antibody αCaTrailin3 (**A**), αCaTrailin4 (**B**) and αCaTrailin2 (**C**). As a negative control, the cells were incubated only in blocking solution without the presence of primary antibody (**D**). All confocal images are presented as a merge of green, red and brightfield channel after the maximum intensity Z-projection. The red color represents chloroplast autofluorescence. Scale bar: 50 µm.

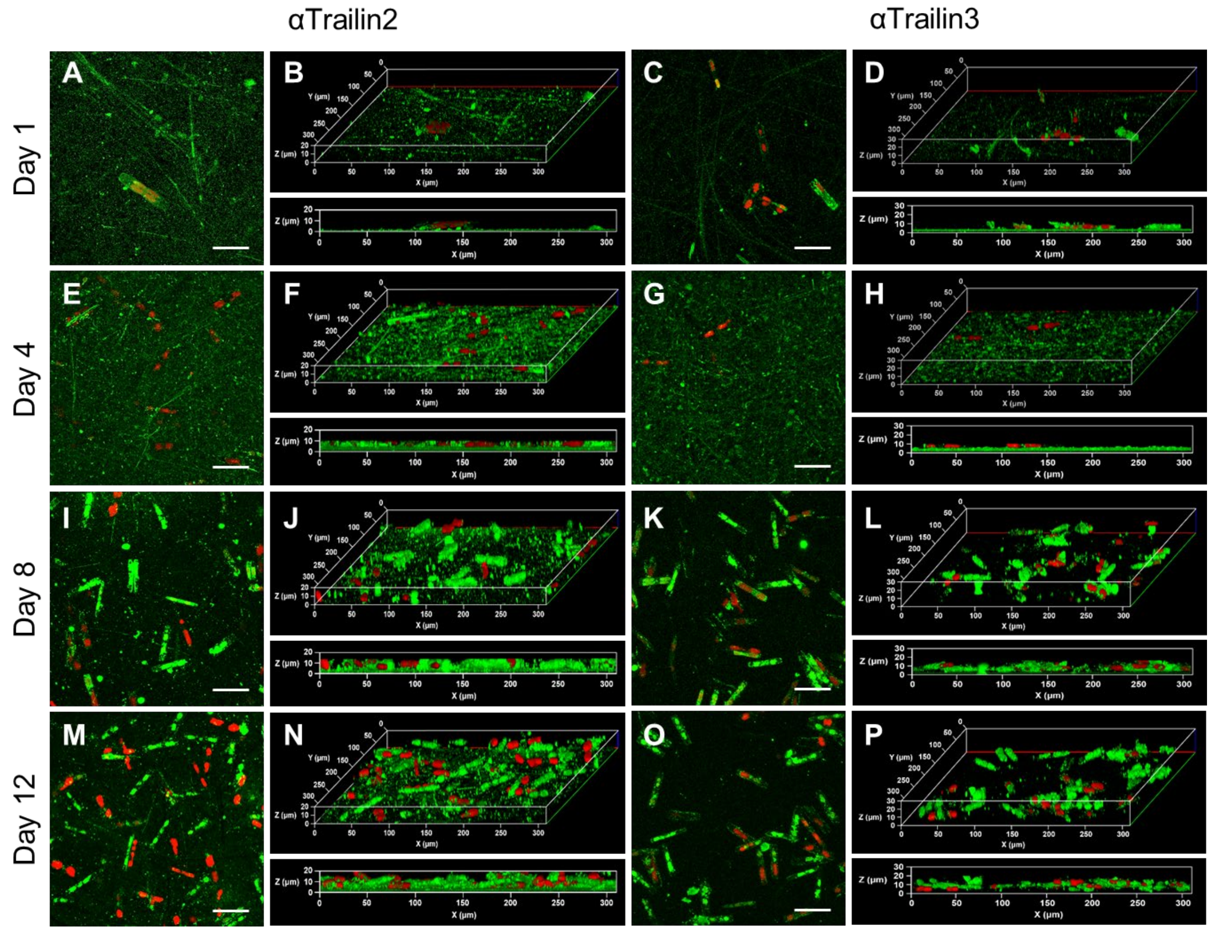

**Figure S13.** Monitoring the presence of adhesive proteins within biofilm by immunolabelling. Immunolocalization of adhesive proteins CaTrailin2 **(left)** and CaTrailin3 **(right)** in *C. australis* adhesive material after 1, 4, 8 and 12 days. The 3D reconstruction of the confocal images clearly shows the increase in height of adhesive material for both CaTrailin2 and CaTrailin3. All confocal images are presented as the maximum intensity Z-projection. The green color represents the fluorescence signal from AlexaFluor488 secondary antibody. The red color represents chloroplast autofluorescence. Scale bar: 50 µm, 3D reconstruction: 300 µm (X) x 300 µm (Y) x 30 µm (Z).

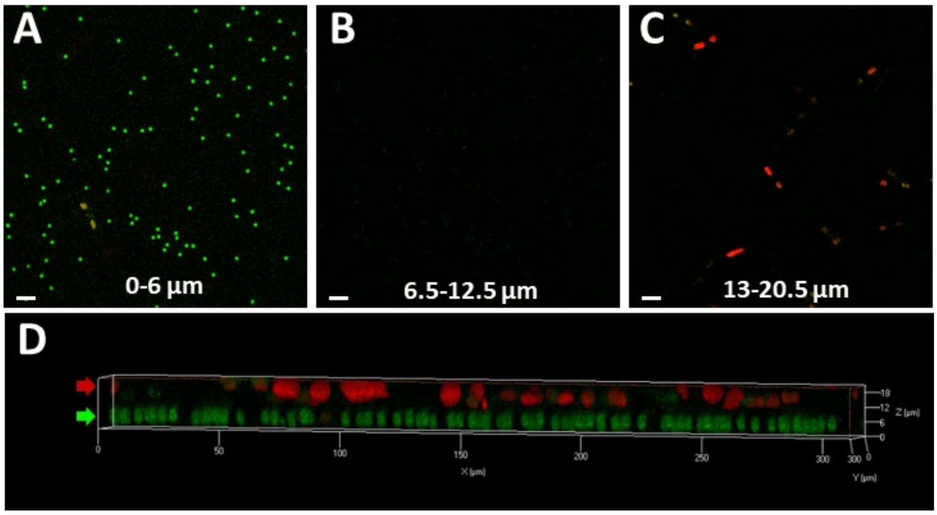

**Figure S14. Control live cell imaging experiments.** Confocal microscopy images of *C. australis* cells and Protein G beads after 12 days. Cells were visualized without any labelling, fluorescence signal is given only from autofluorescence of chloroplasts. **(A-D)** Green color shows the Protein G beads labelled with the secondary antibody, the red color represents the autofluorescence **(A)** maximum intensity Z-projection from the surface (0 µm) to 6 µm **(B)** maximum intensity Z-projection from 6.5 to 12.5 µm **(C)** maximum intensity Z-projection from 13µm to the top of the biofilm (20.5 µm) **(D)** The 3D reconstruction of the confocal images depicts the location of the Protein G beads located underneath the biofilm. 3D reconstruction: **45 µm (X) x 45 µm (Y) x 10 µm (Z)**. Scale bar: 10 µm

#### References

**Armenteros JJA, Tsirigos KD, Sønderby CK, Petersen TN, Winther O, Brunak S, von Heijne G, Nielsen H. 2019.** SignalP 5.0 improves signal peptide predictions using deep neural networks. *Nature Biotechnology* **37**: 420-423.

**Aumeier C. 2015.** *The Cytoskeleton of Diatoms : Structural and Genomic Analysis.* PhD thesis, Rheinische Friedrich-Wilhelms-Universität Bonn, Bonn, Germany.

**Buchfink B, Xie C, Huson DH. 2015.** Fast and sensitive protein alignment using DIAMOND. *Nature Methods* **12**: 59-60.

**de Castro E, Sigrist CJ, Gattiker A, Bulliard V, Langendijk-Genevaux PS, Gasteiger E, Bairoch A, Hulo N. 2006.** ScanProsite: detection of PROSITE signature matches and ProRule-associated functional and structural residues in proteins. *Nucleic Acids Research* **34**: W362-365.

**Doxey AC, Yaish MW, Griffith M, McConkey BJ. 2006.** Ordered surface carbons distinguish antifreeze proteins and their ice-binding regions. *Nature Biotechnology* **24**: 852-855.

**Erdos G, Pajkos M, Dosztanyi Z. 2021.** IUPred3: prediction of protein disorder enhanced with unambiguous experimental annotation and visualization of evolutionary conservation. *Nucleic Acids Research* **49**: W297-W303.

**Finn RD, Attwood TK, Babbitt PC, Bateman A, Bork P, Bridge AJ, Chang HY, Dosztanyi Z, El-Gebali S, Fraser M, et al. 2017.** InterPro in 2017-beyond protein family and domain annotations. *Nucleic Acids Research* **45**: D190-D199.

**Finn RD, Coggill P, Eberhardt RY, Eddy SR, Mistry J, Mitchell AL, Potter SC, Punta M, Qureshi M, Sangrador-Vegas A, et al. 2016.** The Pfam protein families database: towards a more sustainable future. *Nucleic Acids Research* **44**: D279-285.

**Fu LM, Niu BF, Zhu ZW, Wu ST, Li WZ. 2012.** CD-HIT: accelerated for clustering the next-generation sequencing data. *Bioinformatics* **28**: 3150-3152.

**Grigoriev IV, Hayes RD, Calhoun S, Kamel B, Wang A, Ahrendt S, Dusheyko S, Nikitin R, Mondo SJ, Salamov A, et al. 2021.** PhycoCosm, a comparative algal genomics resource. *Nucleic Acids Research* **49**: D1004-D1011.

**Jumper J, Evans R, Pritzel A, Green T, Figurnov M, Ronneberger O, Tunyasuvunakool K, Bates R, Zidek A, Potapenko A, et al. 2021.** Highly accurate protein structure prediction with AlphaFold. *Nature* **596**: 583-589.

**Katoh K, Standley DM. 2013.** MAFFT multiple sequence alignment software version 7: improvements in performance and usability. *Molecular Biology and Evolution* **30**: 772-780.

**Keeling PJ, Burki F, Wilcox HM, Allam B, Allen EE, Amaral-Zettler LA, Armbrust EV, Archibald JM, Bharti AK, Bell CJ, et al. 2014.** The Marine Microbial Eukaryote Transcriptome Sequencing Project (MMETSP): illuminating the functional diversity of eukaryotic life in the oceans through transcriptome sequencing. *PLoS Biology* **12**: e1001889.

**Kumar S, Stecher G, Li M, Knyaz C, Tamura K. 2018.** MEGA X: Molecular Evolutionary Genetics Analysis across Computing Platforms. *Molecular Biology and Evolution* **35**: 1547-1549.

**Letunic I, Bork P. 2018.** 20 years of the SMART protein domain annotation resource. *Nucleic Acids Research* **46**: D493-D496.

**Marron AO, Ratcliffe S, Wheeler GL, Goldstein RE, King N, Not F, de Vargas C, Richter DJ. 2016.** The evolution of silicon transport in eukaryotes. *Molecular Biology and Evolution* **33**: 3226-3248.

**Nakov T, Beaulieu JM, Alverson AJ. 2018.** Accelerated diversification is related to life history and locomotion in a hyperdiverse lineage of microbial eukaryotes (Diatoms, Bacillariophyta). *New Phytologist* **219**: 462-473.

**Nguyen LT, Schmidt HA, von Haeseler A, Minh BQ. 2015.** IQ-TREE: A Fast and Effective Stochastic Algorithm for Estimating Maximum-Likelihood Phylogenies. *Molecular Biology and Evolution* **32**: 268-274.

**Osuna-Cruz CM, Bilcke G, Vancaester E, De Decker S, Bones AM, Winge P, Poulsen N, Bulankova P, Verhelst B, Audoor S, et al. 2020.** The *Seminavis robusta* genome provides insights into the evolutionary adaptations of benthic diatoms. *Nature Communications* **11**: 3320.

**Paradis E, Schliep K. 2019.** ape 5.0: an environment for modern phylogenetics and evolutionary analyses in R. *Bioinformatics* **35**: 526-528.

**Poulsen N, Hennig H, Geyer VF, Diez S, Wetherbee R, Fitz-Gibbon S, Pellegrini M, Kröger N. 2023.** On the role of cell surface associated, mucin-like glycoproteins in the pennate diatom *Craspedostauros australis* (Bacillariophyceae). *Journal of Phycology* **59:**54-69.

**Poulsen N, Kröger N. 2004.** Silica morphogenesis by alternative processing of silaffins in the diatom *Thalassiosira pseudonana*. *Journal of Biological Chemistry* **279**: 42993-42999.

**Shen W, Ren H. 2021.** TaxonKit: A practical and efficient NCBI taxonomy toolkit. *Journal of Genetics and Genomics* **48**: 844-850.

**Shevchenko A, Tomas H, Havlis J, Olsen JV, Mann M. 2006.** In-gel digestion for mass spectrometric characterization of proteins and proteomes. *Nature Protocols* **1**: 2856-2860.

**Van Vlierberghe M, Di Franco A, Philippe H, Baurain D. 2021.** Decontamination, pooling and dereplication of the 678 samples of the Marine Microbial Eukaryote Transcriptome Sequencing Project. *BMC Research Notes* **14**: 306.

**Xu SB, Dai ZH, Guo PF, Fu XC, Liu SS, Zhou L, Tang WL, Feng TZ, Chen MJ, Zhan L, et al. 2021.** ggtreeExtra: Compact visualization of richly annotated phylogenetic data. *Molecular Biology and Evolution* **38**: 4039-4042.

**Yu GC, Smith DK, Zhu HC, Guan Y, Lam TTY. 2017.** GGTREE: an R package for visualization and annotation of phylogenetic trees with their covariates and other associated data. *Methods in Ecology and Evolution* **8**: 28-36.
