## Supplemental Table 2 for "Diatom adhesive trail proteins acquired by horizontal gene transfer from bacteria serve as primers for marine biofilm formation"

**Table S2.** Tabular overview of identified protein hits. NCBI BLAST results are shown for protein hits that returned an E-value less than 1E-20. Top hits are given for both the top diatom and non-diatom species. A hypen (-) indicates the absence of the attribute.

| Transcript ID | Alternative transcript ID | Protein name | Size (aa) | Top NCBI BLAST hit | Species | Accession number | Identity | E-value | Bit Score | Non-Diatom NCBI BLAST hit | Species | Accession number | Identity | E-value | Bit Score | Interpro | Intrisinically disordered IPRed3.0 score>0.5 | Signal Peptide | #1 # peptides | #2 # peptides | #3 # peptides |
|---|---|---|---|---|---|---|---|---|---|---|---|---|---|---|---|---|---|---|---|---|---|
| Ca807 | | CaTrailin_3 | 1479 | hypothetical protein THAOC_34954 | *Thalassiosira oceanica* | EJK46375.1 | 29.35 | 1.00E-57 | 234 | chitinase | *Chytriomyces confervae* | TPX53846.1 | 34 | 3.00E-33 | 150 | IPR026588: Choice of anchor A domain (aa: 996-1284) | 6% | Yes | 9 | 11 | 13 |
| Ca949 | | CaTrailin_1 | 526 | hypothetical protein FRACYDRAFT_240947 | *Fragilariopsis cylindrus* | OEU14408.1 | 31.52 | 2.00E-33 | 145 | choice-of-anchor A family protein | *Curvibacter gracilis* | WP_084461253.1 | 31.85 | 3.00E-24 | 119 | IPR026588: Choice of anchor A domain (aa: 61-353) | 0% | Yes | 2 | 2 | 2 |
| Ca1637 | Ca4552 | CaTrailin_2 | 868 | hypothetical protein FRACYDRAFT_240947 | *Fragilariopsis cylindrus* | OEU14408.1 | 35.16 | 4.00E-37 | 160 | choice-of-anchor A family protein | *Curvibacter lanceolatus* | WP_084679514.1 | 30.06 | 1.00E-28 | 135 | IPR026588: Choice of anchor A domain (aa: 369-677) | 6% | Yes | 2 | 5 | 1 |
| Ca1769 | | CaFAP1 | 1158 | only across PTS rich regions | - | - | - | - | - | only across PTS rich regions | - | - | - | - | - | - | 44% | Yes | 10 | 6 | 5 |
| Ca1945 | | CaTrailin_4 | 9649 | glycoside hydrolase family protein | *Nitzschia inconspicua* | KAG7360667.1 | 43.25 | 7.00E-39 | 176 | - | - | - | - | - | - | IPR026588: Choice of anchor A domain (aa: 460-748); PTHR36489: PROTEIN-COUPLED RECEPTOR GPR1, PUTATIVE-RELATED (aa: 969-1078; 1412-1511; 1640-1754; 1893-1578; 2569-2654; 2960-3101; 3154-3309; 5596-5745; 7310-7455; 7603-7757; 9051-9204) | 81% | Yes | 3 | 4 | 3 |
| Ca5255 | | | 905 | - | - | - | - | - | - | DUF4215 domain-containing protein | *Minicystis rosea* | WP_146725001.1 | 63.87 | 2.00E-158 | 509 | IPR011936: Myxococcus cysteine-rich repeat (aa: 182-552) | 7% | | 6 | 7 | 6 |
| Ca5609 | | | >8655 | hypothetical protein IV203_019591 | *Nitzschia inconspicua* | KAG7371021.1 | 27.46 | 8.00E-75 | 295 | only across PTGD rich regions | - | - | - | - | - | PTHR36489: PROTEIN-COUPLED RECEPTOR GPR1, PUTATIVE-RELATED (1423-1547; 1503-1635; 2425-2573; 2772-2907; 2883-3011; 3217-3383; 3583-3727; 3736-3861; 4059-4197; 4206-4331; 4528-4663; 4813- | 82% | Yes | 1 | 2 | 2 |
| Ca11384 | | | 1609 | unnamed protein product | *Pseudo-nitzschia multistriata* | VEU43960.1 | 30.79 | 3.00E-53 | 219 | - | - | - | - | - | - | IPR036465: von Willebrand factor A-like domain superfamily (aa: 774-1099) | 21% | Yes | 4 | 4 | 1 |
| **Proteins with less than 2 peptides in each experiment** | | | | | | | | | | | | | | | | | | | | | |
| Ca444 | | | 488 | predicted protein | *Phaeodactylum tricornutum* | XP_00217909.1 | 35.47 | 7.00E-10 | 74.7 | PEP-CTERM sorting domain-containing protein | *Leptolyngbya sp.* | NEQ46153.1 | 31 | 4.00E-05 | 57.8 | - | n.d. | n.d. | 6 | 1 | 1 |
| Ca15548 | Ca9058 | | 399 | actin | *Fragilaria crotonensis* | KAI2503016.1 | 99 | 0 | 734 | n.d. | - | - | - | - | - | IPR004000: Actin family | n.d. | n.d. | 5 | 0 | 0 |
| Ca2624 | | | 440 | ribulose-1,5-bisphosphate carboxylase/oxygenase large subunit | *Hantzschia amphioxys var. major* | AEB39371.1 | 96 | 0 | 865 | n.d. | - | - | - | - | - | IPR033966: RuBisCO | n.d. | n.d. | 4 | 0 | 0 |
| Ca54 | | | 683 | protein heat shock protein Hsp70 | *Phaeodactylum tricornutum* | XP_002177351.1 | 93 | 0 | 1120 | n.d. | - | - | - | - | - | IPR013126: Heat shock protein 70 family | n.d. | n.d. | 4 | 0 | 0 |
| Ca2183 | | | 476 | Na_Pi_cotrans-domain-containing protein | *Fragilariopsis cylindrus* | OEU13556.1 | 67 | 0 | 530 | Sodium-dependent phosphate transport protein 2B | *Lamellibrachia satsuma* | KAI0242724.1 | 45 | 4.00E-73 | 252 | IPR003841: Sodium-dependent phosphate transport protein | n.d. | n.d. | 2 | 0 | 0 |
| Ca433 | | | 621 | pyruvate kinase | *Fragilaria crotonensis* | KAI2490519.1 | 79 | 0 | 835 | n.d. | | | | | | IPR001697: Pyruvate kinase | n.d. | n.d. | 2 | 1 | 0 |
| Ca5196 | | | 412 | predicted protein | *Thalassiosira pseudonana* | XP_002297601.1 | 37 | 2.00E-05 | 60.5 | proprotein convertase P-domain-containing protein | *Anaerolineae bacterium* | MBK8837112.1 | 29 | 2.00E-04 | 57.8 | - | n.d. | n.d. | 2 | 1 | 0 |
| Ca1003 | | | 596 | - | - | - | - | - | - | - | - | - | - | - | - | - | n.d. | n.d. | 3 | 0 | 0 |
| Ca888 | | | 458 | translation elongation factor, EF-1, alpha subunit | *Phaeodactylum tricornutum* | XP_002186303.1 | 95 | 0 | 848 | n.d. | - | - | - | - | - | IPR004539: Translation elongation factor EF1A, eukaryotic/archaeal | n.d. | n.d. | 2 | 1 | 0 |
| Ca77 | | | 596 | Eukaryotic-type carbonic anhydrase | *Nitzschia inconspicua* | KAG7343373.1 | 50 | 0 | 541 | n.d. | - | - | - | - | - | IPR023561: Carbonic anhydrase, alpha-class | n.d. | n.d. | 2 | 0 | 0 |
| Ca670 | | | 174 | ubiquitin-domain-containing protein | *Fragilariopsis cylindrus* | OEU20687.1 | 94 | 4.00E-80 | 244 | n.d. | - | - | - | - | - | IPR000626:Ubiquitin-like domain; IPR002906: Ribosomal protein S27a | n.d. | n.d. | 2 | 0 | 0 |
| Ca781 | | | 682 | predicted protein | *Phaeodactylum tricornutum* | XP_002178097.1 | 54 | 0 | 649 | NAD nucleotidase | *Teredinibacter haidensis* | WP_075188388.1 | 39 | 2.00E-131 | 412 | IPR006179:5'-Nucleotidase/apyrase | n.d. | n.d. | 2 | 0 | 0 |
| Ca365 | | | 1105 | cation-transporting P-type ATPase | *Nitzschia inconspicua* | KAG7371055.1 | 70 | 0 | 1418 | n.d. | - | - | - | - | - | IPR001757: P-type ATPase | n.d. | n.d. | 2 | 0 | 0 |
| Ca667 | | | 237 | predicted protein | *Phaeodactylum tricornutum* | XP_002176613.1 | 58 | 6.00E-62 | 204 | hypothetical protein NSK_000944 | *Nannochloropsis salina* | TFJ87593.1 | 38 | 3.00E-22 | 102 | | n.d. | n.d. | 2 | 0 | 0 |
| Ca1241 | | | 341 | hypothetical protein FRACYDRAFT_242426 | *Fragilariopsis cylindrus* | OEU14072.1 | 30 | 4.00E-17 | 94.7 | - | - | - | - | - | - | | n.d. | n.d. | 2 | 0 | 0 |
| Ca8018 | | | | False Mascot hit - stop codon | - | - | - | - | - | - | - | - | - | - | - | - | n.d. | n.d. | 0 | 0 | 0 |
| Ca4387 | | | | False Mascot hit - stop codon | - | - | - | - | - | - | - | - | - | - | - | - | n.d. | n.d. | 0 | 0 | 0 |
| Ca2442 | | | | False Mascot hit - stop codon | - | - | - | - | - | - | - | - | - | - | - | - | n.d. | n.d. | 0 | 0 | 0 |
| Ca3575 | | | | False mascot hit | - | - | - | - | - | - | - | - | - | - | - | - | n.d. | n.d. | 0 | 0 | 0 |
